# Learning interpretable rules of protein binding from a trillion interactions

**DOI:** 10.1101/2025.10.14.682342

**Authors:** Matthew J. Styles, Aditya N. Singh, Shannon S. Lu, Cheng Frank Gao, Jingzhou Yang, Aditya Nandy, Christopher Basile, Joshua A. Pixley, Eddy Pineda, Siyuan Tao, Aaron R. Dinner, Suriyanarayanan Vaikuntanathan, Bryan C. Dickinson

**Affiliations:** Department of Chemistry, University of Chicago, 5735 S. Ellis Ave., Chicago, IL 60637; Department of Chemistry, Biochemistry, and Physics, South Dakota State University, 1055 Campanile Ave., Brookings, SD 57007; Department of Pathology, University of Chicago, 5735 S. Ellis Ave., Chicago, IL 60637; Chan Zuckerberg Biohub, Chicago, IL 60642

## Abstract

Protein binders are invaluable as research tools and therapeutics, yet why some targets readily yield binders while others resist discovery remains poorly understood. Here we quantify this variability through more than 1,300 independent selections against G6 protein targets, experimentally surveying over one trillion protein–protein interactions. We find that binder density—the probability that a random protein variant binds a target—is reproducible, varies by at least five orders of magnitude, and depends primarily on target identity rather than library design. To uncover the molecular determinants of binding, we develop an energy-based machine-learning model that learns pairwise interactions between target and binder residues from protein language model embeddings, and introduce a decomposition that resolves these interactions into interpretable feature–feature couplings. Strikingly, the features associated with binders are extensively shared across targets, whereas those associated with targets are largely unique. These features both recover established molecular recognition mechanisms and identify previously unrecognized interactions in natural proteins. Several of the newly predicted interactions are validated experimentally. Our findings establish a quantitative framework for binder discovery and reveal a shared, interpretable, and transferable molecular vocabulary for predicting protein–protein interactions.

---

Protein-protein interactions (PPIs) underlie virtually every cellular process^1–3^. Beyond native biology, proteins engineered to bind a protein of interest, termed binders, are invaluable as research tools and therapeutics^4–7^. Natural *de novo* interactions between two proteins typically arise via systems that mutate one interacting partner, such as through somatic hypermutation^8^ or diversity generating retroelements^9^. Mirroring this approach, a number of selection methods^10–14^, high-throughput screening^15–17^, and computational design ^18–21^ techniques have been developed for synthetic *de novo* binder discovery; these methods generally have highly variable and unpredictable rates of success.

In this study, we sought to address two questions concerning the emergence of *de novo* PPIs: 1) what is the *a priori* probability that a random protein variant binds a given target; and 2) to what extent is this probability governed by binder characteristics (e.g., scaffold type and randomization strategy, or “design”) versus target characteristics (i.e., how intrinsically “bindable” the target is)? Methods for mapping the sequence-function, or fitness, landscape across a broad array of targets and libraries are key to answering these questions. Current methods, however, are hampered by tradeoffs in fidelity and throughput capacity^11^. Structure-based co-folding and generative design methods ^18–21^ have made substantial progress in enabling *de novo* binder discovery but often require screening of hundreds or thousands of designs and offer limited insight into the sequence and structural determinants that govern recognition, making it difficult to diagnose why a given design succeeds or fails.

We recently developed a high fidelity, high-throughput screening platform, Phage-Assisted Non-Continuous Selection of Binders (PANCS-Binders)^14^, which is capable of rapidly assessing 10^10+^ PPIs across dozens of targets in parallel, providing a new opportunity for large-scale high-resolution PPI fitness landscape mapping. Such maps can potentially be used to derive new heuristics and empirical tools for understanding how likely a selection is to succeed, how library designs can be modified to improve success rate, and what sequences and/or structural features, such as binding motifs, contact geometry, and structural context, underlie *de novo* binding.

Here, we systematically map PPI fitness landscapes by performing rapid, multiplexed de novo binder screens against 96 diverse target proteins using affibody, a compact, thermostable three-helix bundle scaffold widely employed in binder engineering^14,22–24^. We find that binder density is reproducible, is set mainly by the target and by the frequency of a short binding motif, and provides a quantitative baseline against which binder-design methods can be judged. Furthermore, we develop an energy-based machine-learning model that learns pairwise interactions between target and binder residues from protein language model embeddings. We also introduce a decomposition that resolves these interactions into interpretable feature–feature couplings and show how these features recover established molecular recognition mechanisms and, in some cases, identify previously unrecognized interactions in natural proteins. Finally, we experimentally verify some of the newly predicted interactions. Our findings point to a quantitative framework for binder discovery and reveal a shared, interpretable, and transferable molecular vocabulary for predicting protein– protein interactions.

## Quantifying binder density

The potential sequence space for a 58-amino acid protein interacting with a single, invariant target includes 10^75^ variants, too many to exhaustively search. *De novo* binder discovery adopts the heuristics that a well-folded protein scaffold is more likely to bind a target and that randomization should be focused on a few select residues to present a coherent surface for binding^25–27^. In practice, 10-15 positions are randomized, yielding 10^13–19^ theoretical variants (Fig. 1a). Binder density is equivalent to the total number of binders in this sequence space normalized to the total sequence space. However, experimental selection methods can only sample a subset of this total sequence space. Binder density can be experimentally determined from the number of binders present in a sampling from the sequence space and the size of the sampling: if binders are consistently identified from a sampling of 10^9^ unique variants, then the binder density is at least 10^-9^. In a manner analogous to estimating how common planets are by surveying progressively larger numbers of stars until one is reliably detected, we infer binder density from the sampling size at which a binder is consistently identified ^28^ (Fig. 1b) to determine the sampling size required to consistently identify variants that bind to a target (Fig. 1c). This method of assessing binder density depends on the fidelity of the selection method – if a single binder exists in a sampling, the selection must identify it; similarly, if no binder exists, the selection should not have false positives. Our recently reported high-fidelity selection platform, however, is uniquely suited to measure binder density across a wide range of densities (down to 1 in 10 billion).

**Fig. 1:**
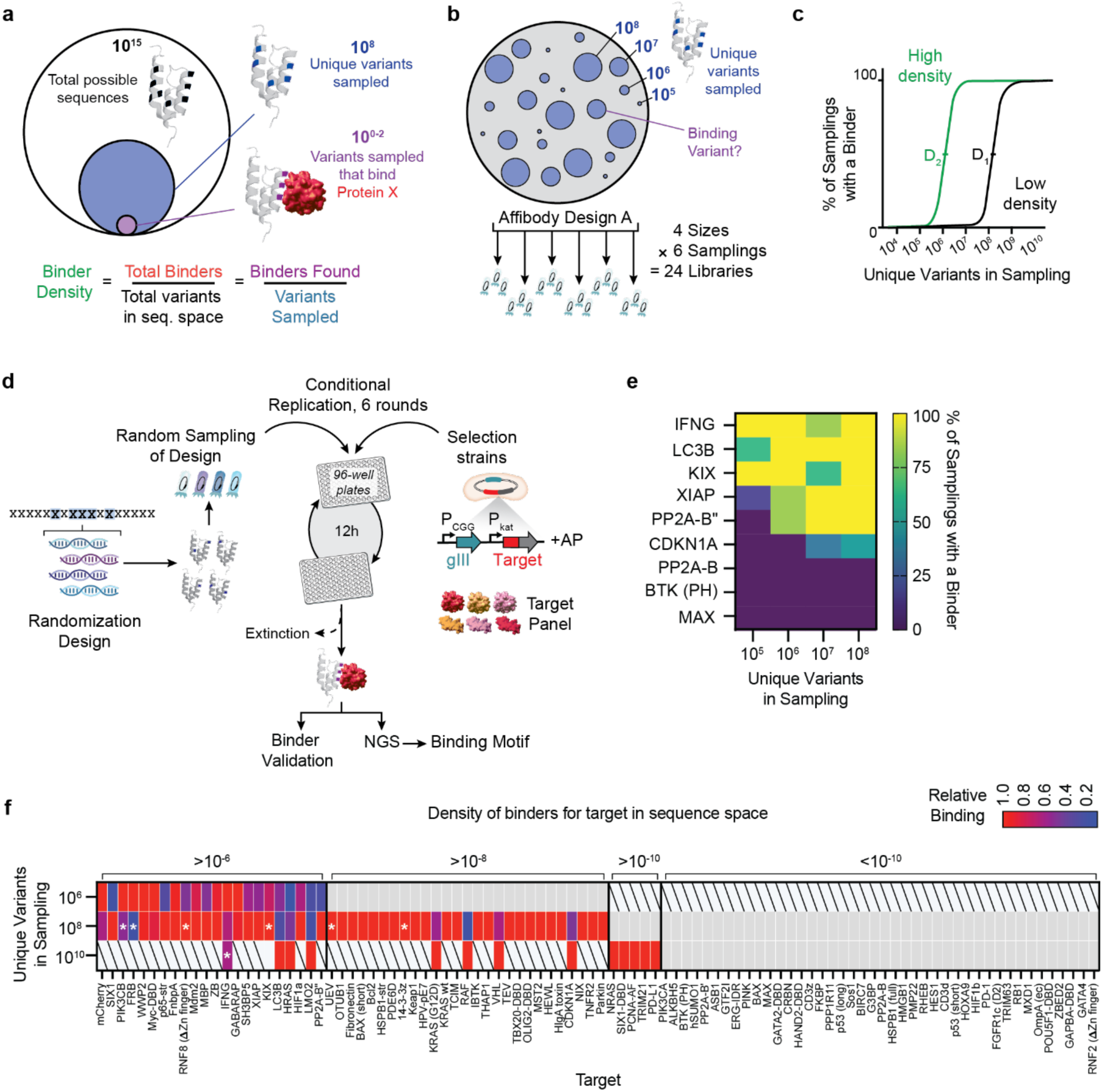
Quantifying binder density. **a**, A library sequence space includes all possible sequences based on the randomization design; for affibody library Design A (Supplementary Fig. 1), this includes >10^15^ possible sequences (white circle). A subset of this sequence space can be encoded into a physical random sampling of the overall sequence space (blue circle). Finally, using selections, the binding variants within the sampling can be identified for a given target protein (purple circle). “Binder Density,” the probability of finding a binder in a randomization design, is defined as the total number of binders in a sequence space over the size of the sequence space. In theory, this parameter can be determined experimentally by identifying how many binders are present in a random sampling of a library design. In practice, identifying whether a binder exists in a random sampling is easier than quantifying the exact number of binders in a sampling. **b,c**, Strategy for quantifying Binder Density. Different sized samplings are generated in replicate (b) and selections are performed (c) to determine what sampling size is required to identify a binder in 50% of samplings. **d**, PANCS-Binder selections: defined libraries of affibody fused to the N-terminal portion of a split-RNA polymerase (RNAP) are encoded in replication deficient phage (one unique variant encoded per phage genome) are passaged on *E. coli* selection strains that express a target protein fused to the C-terminal portion of a split RNAP. PPI-dependent reconstitution of the RNAP drives the expression of *gIII*, allowing phage replication. The replication rate of an individual phage is dependent on the affinity of the interaction of its encoded affibody for the target protein; therefore, endpoint phage populations represent a rank ordering of the binding affinity of affibody variants. After six 12-hour rounds of selection, the titer of the endpoint population is measured, and populations with titers >1 × 10^7^ PFU/mL are submitted for next-generation sequencing (NGS), and the top variant is subcloned and tested for binding activity. **e**, Percent of replicates (*n* = 6) which enriched a binder (endpoint titer >10^7^ (Supplementary Fig. 2) and dominant variant bound in validation luciferase assay (Supplementary Fig. 4); see Methods for further details of selection assessment) with different quantities of unique variants in a sampling on a 9-target panel (Supplementary Table 1). **f**, Selections were performed on a 96-target panel (Supplementary Table 1) with a single 10^8^ unique variant sampling size. Targets without a binder in this 10^8^ selection, were subject to a 10^10^ sampling size selection; targets with a binder in the 10^8^ selection were subject to a 10^6^ sampling size selection (several previously reported 10^10^ selections were also included in this dataset). After determining the endpoint titer (Supplementary Fig. 5) and collecting NGS, we isolated the dominant variant for validation in our luciferase assay (Supplementary Fig. 6). Heat map displays the relative luciferase signal for these dominant variants. The relative luciferase activity of binder per target was calculated by subtracting the background (negative control signals) from those of each binder, then normalizing to the best binder discovered for a target. Grey boxes: no binder was discovered; slashed boxes: not tested; white asterisks: selections where a variant less than three-fold abundant compared to the top variant was tested in the binding assay. Based on the sampling size required to identify a binder to each target, we split the 96-target panel into four categories of binder density with respect to design A: >10^-6^, >10^-8^, >10^-10^, and <10^-10^.

PANCS-Binders (Fig. 1d), which uses *in vivo* selection rather than separation-based selection, can reproducibly enrich a single binder from a pool of 10^10^ non-binders with near-perfect fidelity. We generated 24 samplings of the affibody scaffold containing different numbers of unique variants (six independent samplings of 10^5^, 10^6^, 10^7^, and 10^8^ unique variants, Fig. 1b) using a previously identified randomization design (Supplementary Fig. 1)^29^. We selected nine target proteins (see Supplementary Table 1 for target details) and performed PANCS-Binder selections with these 24 samplings (Fig. 1d). At the end of each PANCS-Binder selection, phage titer was used to determine which selections enriched binder variants. Next-generation sequencing (NGS) of high-titer selections was performed (Supplementary Figs. 2 and 3 and Supplementary Data 1) and the dominant variant was subcloned into an analogous split RNAP complementation binding assay to validate binding activity of the most enriched variant (Supplementary Fig. 4). We observed a wide range of affibody-target binder densities from >10^-5^ to <10^-8^ (Fig. 1e, Supplementary Table 2). For targets with intermediate binder density (between 10^-5^ and 10^-8^), we observed an expected transition from no binders identified to binders always being identified as the number of unique variants in a sampling was increased (XIAP, PP2A-B’’, CDKN1A). These results demonstrate that binder density is a reproducible and quantifiable feature of *de novo* PPI sequence landscapes.

We next expanded to 96 targets of varying size, structure, disorder, function and localization (Supplementary Table 1). To estimate binder density efficiently, we first performed one selection per target with a 10^8^-variant sampling, at which most of the original targets (6 of 9) yielded binders. Out of the 96 selections, 49 had binders (Fig. 1f; titers in Supplementary Fig. 5, NGS in Supplementary Data 2, and lux binding assay in Supplementary Fig. 6). To further stratify these targets by affibody binder density, we performed two subsequent sets of selections: for targets with binders in the 10^8^ sampling, we performed a selection with a smaller, independent 10^6^ unique variant sampling, and for targets that failed to enrich binders with the 10^8^ sampling, we performed selection with a larger 10^10^ unique variant sampling (Fig. 1f; titers in Supplementary Fig. 5, NGS in Supplementary Data 2, and binding assay in Supplementary Fig. 6). We separated targets into four categories based on selection outcomes: high density (22 targets with binder densities >10^-6^), moderate density (27 targets with a binder density between 10^-8^ and 10^-6^), low density (5 targets with a binder density between 10^-10^ and 10^-8^), and undetectable density (42 targets with binder densities <10^-10^). These results suggest that binder density varies across at least five orders of magnitude from at least one in 10^5^ to less than one in 10^10^. To the extent that this 96-target panel captures a reasonable representation of the broader human proteome, these results suggest that a large fraction (∼25%) of the proteome has a high binder density and >50% have easily discoverable binder densities (i.e., binders can discovered from accessible library sizes).

## Motif frequency sets binder density

Contacts between the target protein and the binder protein mediate binding. These contacts can be formed from amino acid side chains that were randomized in the library design (library positions) as well as from those that were not randomized in the library design (scaffold positions). Here, the originally randomized positions that mediate binding are referred to as the binding motif (Fig. 2a), and the frequency of these motifs can be calculated based on the specific motif identity and randomization strategy (scaffold positions can contribute to overall binding motif, but do not impact motif frequency). We hypothesized that binder density is primarily determined by the minimal number of library positions that must be specified to create a minimal binding motif for a target; i.e., binder density should be approximately equal to binder motif frequency (Fig. 2a). This relationship should constrain the size of discoverable binding motifs (Fig. 2b) to much fewer specified amino acids than are typically randomized in a library.

**Fig. 2:**
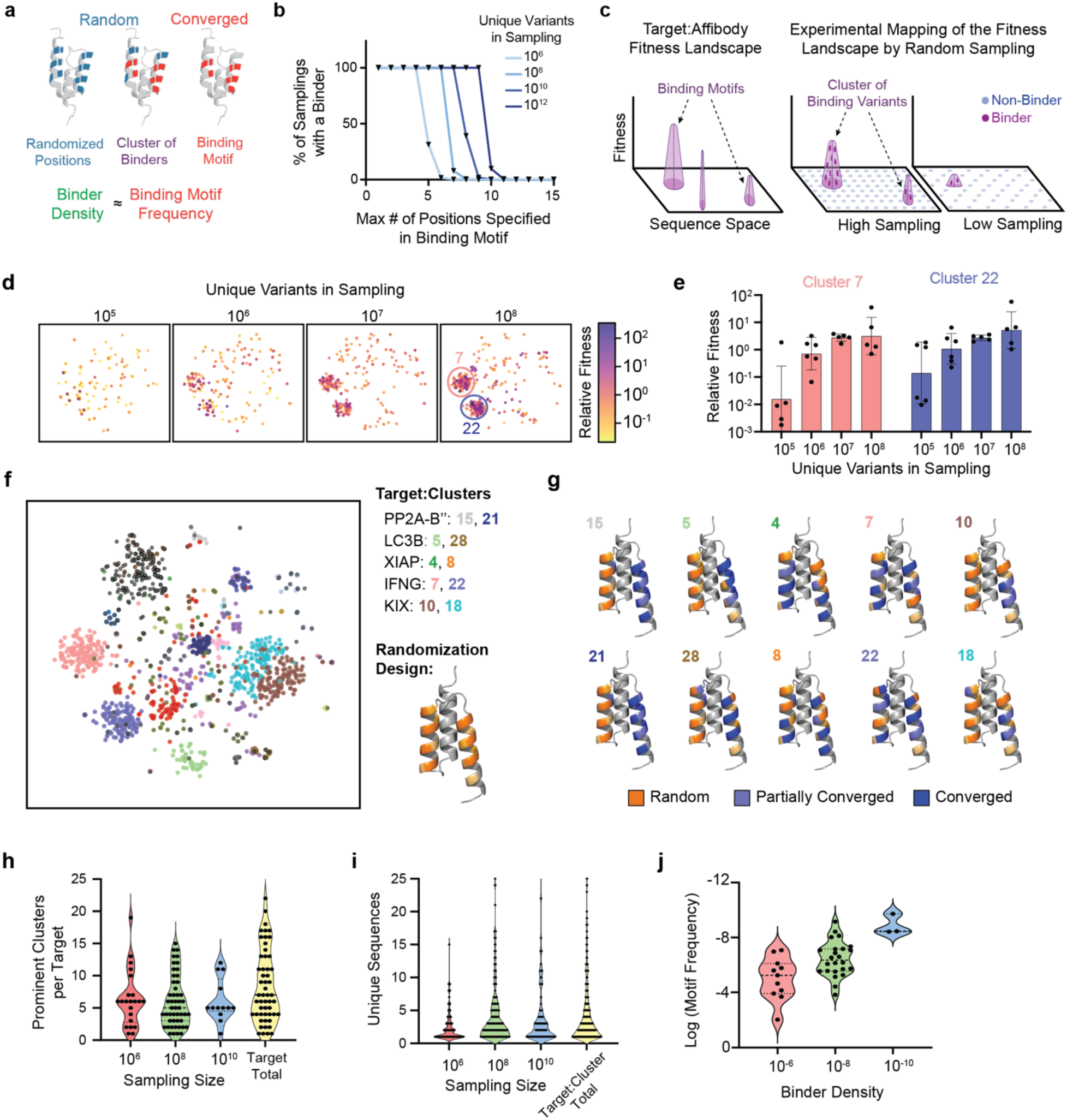
Binder density is determined by motif frequency. **a**, Within the randomized positions in our affibody library (left, blue), several positions will mediate binding to the target (right, red) for binding variants. At the end of a selection, there are likely to be several variants with the binding motif. The originally randomized variants that are not part of the binding motif should remain essentially random amongst the cluster of variants that contain the binding motif. **b**, The minimal number of amino acids required to specify a binding motif determines the frequency of the binding motif, the number of binding motifs within a random sampling of variants, and the chance of success for a given sampling size – as sampling depth increases, more complex binding motifs become discoverable. NNK randomization (all amino acids) is assumed in this calculation of how likely a binder is to be present in a sampling given different numbers of specified positions. **c**, A representation of the possible underlying binding motifs for a PPI fitness landscape. Random sampling can be used to determine the underlying topology by identifying variants a motif. Lower sampling sizes will change which motifs are sampled. **d**, Scatter plots on a t-SNE embedding showing all variants with >0.1% of NGS reads from all samplings that produced a binder for IFNG at each sampling size; colour indicates relative fitness (see Methods). **e**, The relative fitness (see Methods for calculation) of the two most dominant clusters (by % NGS) across sampling size, coloured by cluster identity. Each bar represents the geometric mean (*n*=6, except for 10^7^ which has *n* =5) and standard deviation (SD). **f**, t-SNE embeddings for all 10^8^-unique variant-sized samplings across all targets tested, coloured by cluster assignments. **g**, For select clusters (indicated on right in **f**), the convergence (blue) of randomized positions (orange) on the affibody scaffold (grey) are shown to indicate the position of binding motifs (darker colour indicates greater convergence or randomization; see Supplementary Fig. 10 for convergence analysis). The randomized positions for the design are indicated on the affibody structure on the left. **h,i**, For the 96-target panel selections (Fig. 1f) binding motif analysis (Supplementary Table 3), the number of prominent clusters (>1% of NGS reads for a target) per target (h) and the number of unique variants per target-cluster pair for each set of sampling sizes and then the total across sampling sizes for a target (i). **j**, Frequency of the top motif for targets in each binder density category.

The relationship between fitness and sequence can be represented as a landscape with high and low fitness represented by peaks and valleys, respectively (Fig. 2c)^30,31^. Here, the peaks represent distinct binding motifs, and all variants within a peak contain the same binding motif. We mapped *de novo* PPI fitness landscapes using our selections and endpoint sequencing results. Sampling size should dictate how well the landscape is mapped, with higher sampling sizes having the potential to identify rarer binding motifs. In selections with sampling sizes much larger than the binder density, many variants with that motif should be enriched (Fig. 2c). In principle, we should be able to identify the number of binding motifs (observable at the sampling depth) on the landscape through hierarchical clustering of binders identified in a selection. If enough variants are identified for a cluster, then we should be able to decipher the identity of the amino acids that comprise the binding motif, allowing comparisons of binder density to motif frequency.

To test this general approach, we began with our 9-target panel selection results (Fig. 1e). We pooled all variants that enriched to >0.1% of the population across all successful selections for all targets and performed hierarchical clustering (Methods, Supplementary Data 3). To visualize the clusters in two dimensions, we plotted sequence variation on a t-SNE embedding^32^. Using IFNG as a representative target, we observed sparse sequences at 10^5^ sampling size that coalesced into discrete, well-separated clusters at 10^8^ sampling size (Fig. 2d). The fitness landscape for each target behaved similarly: several clusters became increasingly populated after sampling above the binder density (Supplementary Fig. 7), each sampling produced a similar distribution of variants from each cluster, and the fitness of variants increased with increasing sampling size within each cluster (Fig. 2e, Supplementary Fig. 8). We plotted the binders for all targets on a single t-SNE plot to visualize the distinct clustering of variants in a target- and motif-specific manner (Fig. 2f). Because each target has several clusters that are poorly populated (even when sampled much higher than the binder density), we focused on “prominent clusters,” those with variants that comprise >0.5% of sequencing reads (thus setting a minimum fitness for the cluster).

Each cluster contained a varying number of sequences, with clusters from higher binder density targets having more sequences per cluster (Supplementary Fig. 9). Each target had at least two clusters with enough unique sequences (>10) to determine the underlying binding motif for these prominent clusters (except for CDKN1A clusters). Using the individual variants, we analysed the convergence at each randomized position (Supplementary Fig. 10) to identify binding motif underlying each prominent cluster and map them onto the affibody structure (Fig. 2g). While the exact binding motif varied between targets, similar positional motifs were observed across targets: binders to KIX, LC3B, and PP2A-B’’ have motifs that act as single helix binders while binding motifs for XIAP and IFNG utilize both helices on one face of the affibody. We observed strong concordance between the frequency of binding motifs within the randomization design (Supplementary Fig. 11) to the observed experimental binder density (Fig. 1e). KIX and IFNG had binder densities ≥10^-5^ and have motif frequencies of 10^-2^ and ∼5 × 10^-6^, respectively. Additionally, the binder motif for PP2A-B’’ has a frequency of 10^-6^ and observed binder densities of 10^-6^. Therefore, based on the targets for which we were able to most accurately assess both binder density and motif frequency, we observed that binder density mirrored the motif frequency. We next sought to test whether these trends held for our broader target panel.

We performed hierarchical clustering using all variants with >0.1% of sequencing reads from all the selections for our 96-target panel (Fig. 1f, Supplementary Data 4, see Methods for additional details). We used 300 clusters, which balanced merging unrelated variants against splitting related ones (Supplementary Fig. 12 and Methods). Across 53 targets, we observed 263 prominent clusters (>1% sequencing reads in the selection for a Target-Sampling Size pairing), 6 (+/-4 SD) prominent clusters per target for selections with 10^8^ sampling size (Fig. 2h, Supplementary Table 3, Supplementary Fig. 13). The number of unique sequences per cluster increased with sampling size (Fig. 2i and Supplementary Fig. 13), and 93 target-cluster pairs across 29 targets had six or more unique sequences; we performed convergence analysis to identify each binding motif and calculate its frequency (Supplementary Fig. 10) for each of these cluster-target pairs (Supplementary Table 3). On average, 7.5 positions were converged (+/-1.6) out of the 17 randomized positions (ranging from 3-11 positions; Supplementary Fig. 14). We again observed a strong correlation between estimated binder density from the sampling size required to identify binders and the calculated motif frequency (Fig. 2j, Supplementary Fig. 14, Pearson Correlation *r* = 0.61, *P* = 0.0001). Notably, the subset of clusters for which we identified a motif are those for which sampling depth greatly exceeded binder density; thus, these are characteristics of the simpler binding motifs within our broader set of target-binding motif pairs. In total, these results support our hypothesis that binding motif frequency determines binder density (Fig. 2a) and provides quantitative characteristics of *de novo* PPI fitness landscapes (see Supplementary Note 1 for implications for affinity maturation and library design).

## Target, not library, sets bindability

To assess how specific design features of the scaffold variation influence binder discovery, we constructed eight additional affibody designs based on three distinct design considerations: different randomized positions (Designs A–C); different amino acid usage via diverse codon degeneracy (Designs B, D–F); and altered hydrophobic cores to vary stability/expression (Designs G–I) (Fig. 3a, Supplementary Tables 4 and 5, Supplementary Fig. 15). Additionally, we constructed a fully randomized 30-mer NNY design (Design J; ∼2 × 10^35^ theoretical variants, Supplementary Table 4) to serve as a reference for scaffold-less binder density. A ∼10^8^ sampling of each design was screened across the 96-target panel (Fig. 3b, Supplementary Fig. 16, Supplementary Data 2, Supplementary Fig. 6), revealing substantial variation in hit counts. Targets with higher binder density with Design A more consistently yielded hits across alternative designs, suggesting these represent intrinsically “bindable” targets (Fig. 3c). Additionally, we compared the results with an alternative scaffold, affitin (10^8^ sampling size) which mirrored our binder density categories for the affibody library (Fig. 3b): 89% (16/18) high density targets, 67% (16/24) moderate density, 25% (1/4) low density, and 12% (3/25) for undetected density. This consistency suggests that the results observed with affibody are not specific to the affibody scaffold, but a general target characteristic.

**Fig. 3:**
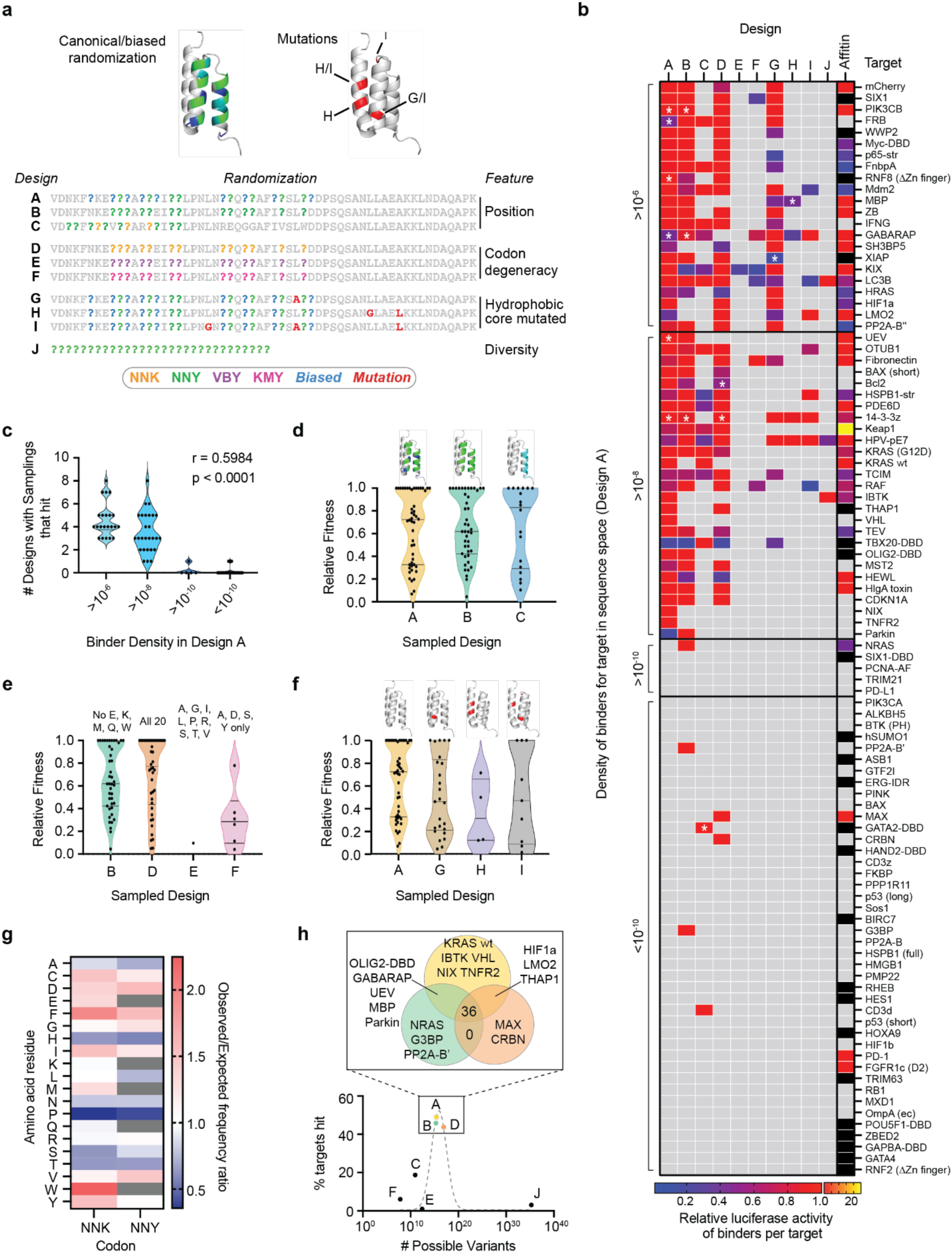
Elucidating the impact of randomization design on target-affibody binder density. **a**, Designs (Supplementary Table**s 4 and 5, Supplementary Fig. 15**) for nine affibodies (Designs A-I) and a tenth fully random NNY 30-mer (Design J) to assess the relative importance of randomization position, codon diversity, and scaffold stability. **b**, Heat map indicating the relative luciferase activity across all binders for a given target (based on the *E. coli* RNAP complementation luciferase assay) normalized to the highest-signal binder from Design A-J. Asterisks: selections where the variant stochastically was <3-fold as abundant as the top variant. The Affitin set is from a 10^8^ affitin library. Black boxes: selection with the affitin library was not performed on that target. If the affitin variant had a higher luciferase signal than the highest signal affibody, then the value can be >1 (see scale bar). **c**, Violin plot comparing binder density category and number of designs that hit with a 10^8^ sampling (two-tailed Pearson correlation, r = 0.5984, 95% C.I. 0.3972 to 0.7013, *P* <0.0001). **d–f**, Violin plots showing the relative fitness of all hits discovered in indicated sampled design, as measured via binding luciferase assay. **g**, Position-independent amino acid usage for degenerate codons NNK and NNY, calculated from all binder sequences across all designs containing NNK or NNY codons. Frequencies are expressed as an aggregated ratio between observed and expected frequencies across all positions containing an NNK or NNY codon. **h**, Scatterplot of sequence space diversity versus target hit rate. Dotted grey line shows a lognormal-like relationship between diversity and hit rate, included as a visual guide only due to limited (7) data points. Venn diagram of the targets hit by the top three-performing Designs A, B, and D, with targets where not all three designs hit listed.

Randomizing two helices (Design B) rather than one (Design C) increased the number of targets hit from 18 to 43 (Fig. 3b,d and Supplementary Figs. 17 and 18), whereas additional restricted randomization of surface positions (Design A) made little difference (Supplementary Fig. 17). Thus, while single-helix randomization reduces binder density, likely by restricting possible binding motifs, binders from Design C often show equal or better fitness. Therefore, two-helix randomization generally outperforms one-helix designs by accessing more possible binding motifs but at the cost of reducing the fitness of single-helix motif binders, likely via the additional non-converged positions in the second helix.

Next, we investigated how amino acid diversity at canonically randomized positions affects binder discovery. We restricted residue incorporation by changing the degeneracy in the codon randomization: Design D uses NNK (all 20 amino acids + stop), Design B uses NNY (no stop, Glu, Lys, Met, Gln, or Trp), Design E uses VBY (postulated primordial amino acids^33^), and Design F uses KMY (only Tyr, Ala, Ser, and Asp to test Tyr’s privileged role in PPIs^34^). We found that reduced amino acid diversity decreased the number of targets with binder densities >10^-8^ (Fig. 3b) and yielded lower-fitness binders when present (Fig. 3e). Samplings of Designs B and D (NNY and NNK codons) performed comparably (43 and 41 hits, 36 overlapping hits; Supplementary Fig. 17), with similar fitness (Fig. 3e), while the samplings of Designs E and F (VBY and KMY codons) performed poorly, producing only one and six low-fitness hits, respectively (Fig. 3e), each hit overlapping with hits for Design B (Supplementary Fig. 17). Nevertheless, when all amino acid categories are represented, additional chemical redundancy (e.g., Lys vs. Arg, Glu vs. Asp) offers only marginal gains, likely due to the conflicting balance between additional possible motifs and dilution of all motif frequencies.

Finally, in Designs with mutated hydrophobic cores, which alter the average expression level (a proxy of stability) of randomly isolated variants (Supplementary Fig. 19), we observed a precipitous loss of binder density and quality (Fig. 3f). Therefore, as expected, stability and structure play a major role in binder density^26,35^, and even a few core mutations can negate the effects of binder density increases imparted by a stable scaffold protein.

Every amino acid was used among hits from NNK- and NNY-randomized codons, with similar usage between the two (Fig. 3g and Supplementary Fig. 20). When comparing number of theoretically possible variants to hit rates from 10^8^ variant samplings across the 96-target panel (excluding core-mutated libraries), hit rates increased with more randomized positions and codon diversity but plateaued around 51%, with no gain beyond a diversity of ∼10^15^ before decreasing with increased diversity (Fig. 3h). These results indicate the existence of both an optimal diversity window and a practical ceiling for scaffold randomization design. We hypothesize this optimization is driven by the fact that increasing diversity at the position and codon levels increases the possible binding motifs in the design, while at the same time additional randomizations or redundancies in amino acid types dilute each individual binding motif and reduce the average stability of design variants. Therefore, while modifying randomization positions or codons is unlikely to enhance binder density, we anticipate that mutations that improve scaffold stability could lead to higher binder densities and improved fitness.

We next asked what target characteristics correlate with target “bindability” (binder density category as a proxy). Again, we relied on our binder density categories (high (>10^-6^), moderate (10^-6^ to 10^-8^), low (10^-8^ to 10^-10^), and undetectable (<10^-10^); Fig. 1f), modified to place “undetectable” targets for which one of our other Designs hit (Fig. 3b) into the “low” category (Supplementary Table 6). We grouped our 96-target panel by structural and functional features (Supplementary Table 7); however, median binder density did not differ significantly across groupings (Kruskal-Wallis *H* = 10.81, df = 7, *P* = 0.147) (Supplementary Fig. 21) suggesting that intrinsic target-specific properties outweigh group-level trends. We then tested a set of individual characteristics (Supplementary Table 8) for significant differences between binder density categories (Supplementary Fig. 22): expression level in PANCS (Pearson’s correlation, *r* = – 0.2775, *P* = 0.0241) and percent disorder (*r* = 0.2439, *P* = 0.0485) showed modest significance; however, all other features, including length, solvent-accessible surface area, secondary structure content, and AlphaFold pTM or ipTM scores, were not significantly correlated with binder density (Supplementary Fig. 22). Solvent-exposed residue distributions did not differ across binder density categories (Supplementary Fig. 23). A multiple linear regression incorporating all variables explained only 23% of the variance in binder difficulty rank (R² = 0.2301, *P* = 0.0270), with disorder (β = 0.285, *P* = 0.0087) and expression (β = –9.7×10^-6^, *P* = 0.0402) as the only significant contributors (Supplementary Table 9). Principal component analysis of all eight variables likewise failed to resolve clusters by binder density (Supplementary Fig. 24). Thus, while disorder and expression are mildly associated with target tractability, standard biophysical and structural parameters are not, pointing to the likely influence of additional, untested features.

## A model reveals interpretable features

The preceding sections show that binding is governed by motifs composed of a small number of amino acids (3-11 converged positions; Supplementary Fig. 14a) that are target specific (Fig. 3b). Each motif represents a distinct combination of sequence and structural features at the binding interface, and a binder that matches any one such combination can bind its target. Convergence analysis, however, is tied to a specific scaffold and is blind to fixed scaffold residues and to distributed structural features.

Motivated by these observations, and to disentangle the drivers of bindability, we constructed a model that scores binding as an interaction between contextual representations of binder and target residues. The interaction between each binder residue *i* and target residue *j* is *b_i_*^*T*^ *W t_j_*, a minimal pairwise energy that explicitly accounts for target identity, where *W* is a learned interaction matrix (Fig. 4a). Here, b_i_ and t_j_ are residue-level embeddings from ESM-2 650M^36^, a protein language model whose representations, learned purely from sequence, encode the local sequence and structural context of each position, including contributions from neighboring residues that are fixed in the library design. These pairwise scores are aggregated across all residue pairs via a statistical mechanical model, so that the overall binding score is dominated by the strongest local contacts rather than averaged over the full protein surface.

**Fig. 4:**
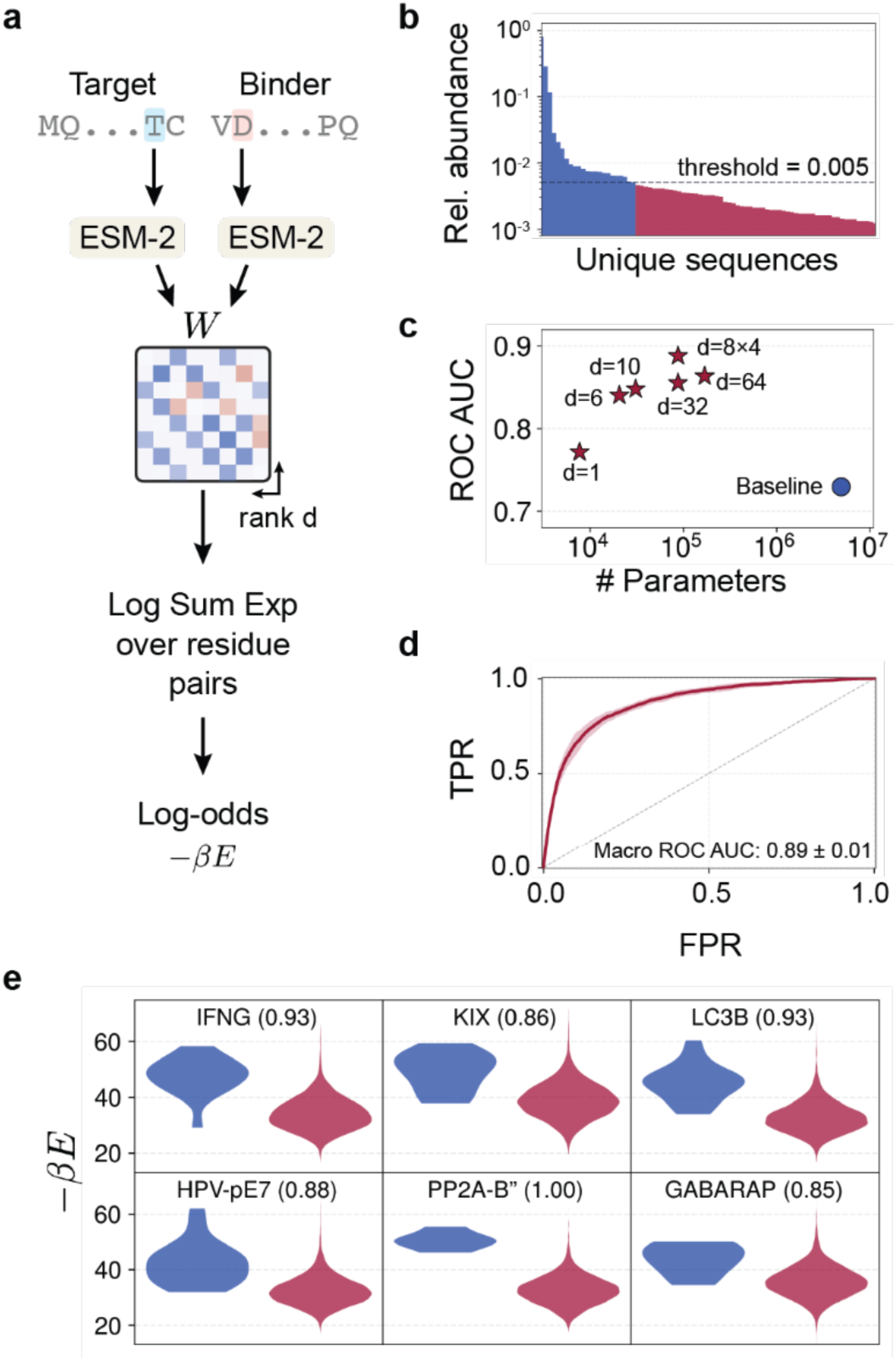
An ML model trained on PANCS selection data learns an interaction matrix that discriminates binders from non-binders across targets. **a**, ESM-2 embeddings of the target and binder are scored through a learned interaction matrix *W* of rank d, aggregated over all residue pairs via soft maximum to produce a log-odds, or a score−*βE* (**Methods**). **b**, Positive binders (blue) and non-binders (red) are defined by a relative abundance threshold of 0.005 in the selection data, corresponding to enrichment above background in NGS read counts. **c**, Macro ROC AUC as a function of model parameters at varying rank d (red stars) and MLP baseline (blue circle; **Methods**). d=8×4 denotes 4 matrices of rank 8. Error bars denote standard deviation across five-fold cross validation and are contained within the marker size. Metrics and parameter counts are given in Supplementary Table 10. **d**, ROC curve for the d=8×4 model (macro AUC = 0.89 ± 0.01). Shading denotes the standard deviation across five-fold cross-validation. **e**, Distributions of − *βE* for positive binders (blue) and non-binders (red) on the test set with per-target AUC in parentheses. Distribution for all targets are given in Supplementary Fig. 25.

We defined positive binders as sequences exceeding a relative abundance of 0.5% in the selection endpoint, a threshold that clearly separates binder enrichment from background (Fig. 4b; **Methods**). Since most sequences are non-binders, and even those differing by a single residue from a confirmed binder frequently fail to enrich, we trained the model using contrastive learning (**Methods**)^37^. This structured model outperforms an MLP baseline trained on the same data with ∼57-fold more parameters (Fig. 4c, Supplementary Table 10), indicating that the pairwise structure, combined with sufficiently expressive residue representations, captures the binding landscape efficiently. Despite its simplicity, the model achieved a macro-AUC of 0.89 (Fig. 4d), and the -*βE* score consistently separates binders from non-binders across individual targets (Fig. 4e, Supplementary Fig. 25).

The learned interaction matrix *W* captures the rules of binding but expresses them along the dense dimensions of ESM-2, in which individual biological concepts are encoded in superposition rather than in isolation^38^. To disentangle them, we re-expressed *W* through the InterPLM feature dictionary^39^, a sparse autoencoder that decomposes each ESM-2 embedding into 10,240 interpretable features corresponding to sequence and structural motifs (Fig. 5a; **Methods**). Since the dictionary is unannotated, we developed a concept-discovery pipeline that identifies the biological concept that each feature encodes. The pipeline discovers compositional motifs (conjunctions of residue identity, flanking context, and UniProt annotations^40^) through a greedy search over positional sequence contexts, with each annotation validated by held-out-data (**Methods**). These annotations are determined from Swiss-Prot, independently of the binding data, and are used for all analyses here. LLM-based annotations are also provided for comparison (Supplementary Data 5).

**Fig. 5:**
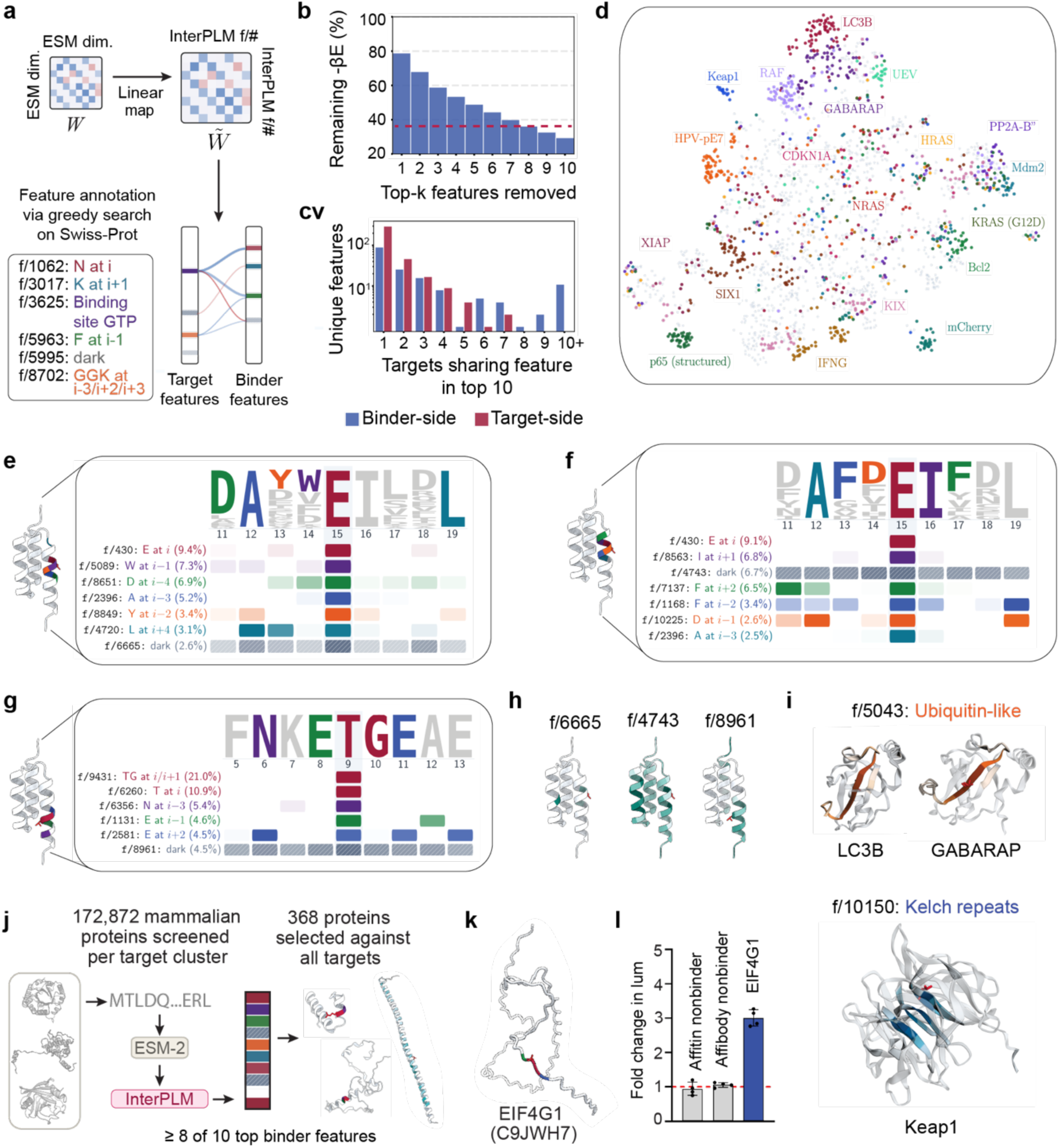
Decomposition of the interaction matrix into feature-feature couplings. **a**, The trained interaction matrix *W* (ESM-2 embedding space) is re-expressed in the InterPLM sparse autoencoder basis via a linear transform, yielding a feature-feature interaction matrix *W̃* that couples target-side features to binder-side features. Each feature is independently annotated against SwissProt concepts through a deterministic concept-discovery pipeline **(Methods)**. Features that resist annotation are classified as dark. Both deterministic and LLM-based annotations are provided in Supplementary Data 5. (b) By linearity, each feature’s contribution to the log-odds is additive and unique **(Methods)**. The top features account for the majority of the discriminative signal and removing them is sufficient to reduce the binding score below the non-binder mean (dashed line). **c**, Distribution of target-side (red) and binder-side (blue) feature reuse across targets. Each bar indicates how many unique features appear in the top 10 rankings of a given number of targets. Most target features appear in only a single target’s top 10, while binder-side features are extensively shared. **d**, UMAP projection of binder-side feature contributions to the score for all positive binder-target pairs, coloured by target identity. **e–g**, Top binder-side features for LC3B (e), GABARAP (f), and Keap1 (g) ranked by contribution to the predicted score. The sequence logo shows the converged motif at the indicated scaffold positions. Each row represents one InterPLM feature, with its annotation and relative importance (percentage of total discriminative signal) on the left. Features are classified as bright when they correspond to an annotatable concept, or dark (hatched). The activation grid shows that the top bright features all fire at the indicated position (aligned in centre), concentrating the discriminative signal at a single site. Their co-activation reconstructs the LIR motif (e), and the NxETGE motif (g). Shared features between LC3B and GABARAP (f/430, f/2396) reflect structural homology, while aromatic features diverge. The structure maps feature activations onto the affibody scaffold, with the stick residue marking the point of contact. **h**, Dark features from LC3B (f/6665), GABARAP (f/4743), and Keap1 (f/8961) activate broadly across the scaffold and detect structural determinants. **i**, Dominant target-side feature for LC3B and GABARAP (left) and Keap1 (right) all of which recover structural folds. **j**, Feature-based search for candidate natural interaction partners. The top binder-side features were used as a query against 172,872 human, mouse, and rat proteins in UniProt **(Methods)**, identifying proteins whose sequences activate the same feature constellation (Supplementary Data 7; Supplementary Data 8). **k**, For Keap1, the search recovers EIF4G1 coloured by binder-side feature activations matching the scheme in **g**. **l**, Experimental validation of the EIF4G1-Keap1 interaction by split-RNAP luciferase complementation assay. The fold-change in luciferase signal for the predicted interaction partner (blue) was divided by the average of two non-binding controls. Each bar plot shows the mean value (*n* = 4, error bars indicate SD).

Since the change of basis is exact for the specific model we used (a variant trained on SAE-reconstructed embeddings, macro-AUC 0.81 ± 0.02; Supplementary Table 10) and each feature’s contribution to a prediction is additive and unique (**Methods**), we next obtained a well-defined marginal importance for every feature in every prediction. As illustrated for a Keap1 binding-mode cluster (Fig. 5b), removing the top-ranked features is sufficient to reduce the binding score to the non-binder mean. These properties allowed us to rank features by how much the model’s decision depends on them and identify which binder- and target-side features binding prediction depends on for each target (Fig. 5c). On both sides the distribution is highly skewed, but with a striking asymmetry. On the binder side, only 165 unique features account for all top 10 rankings across targets, with a single feature appearing in the top 10 of 34 targets. On the target side, 379 unique features are required, the majority appearing in only a single target’s top 10 (Fig. 5c), reflecting the diversity of natural target surfaces against a shared binder scaffold.

To visualize how binder-side feature requirements vary across targets, we projected feature importances onto two dimensions using UMAP (Fig. 5d)^41^. Targets with related binding surfaces grouped together: LC3B, GABARAP, and UEV formed a loose grouping consistent with their shared autophagy-related binding surfaces^42^, while KRAS G12D, NRAS, and HRAS clustered along the right edge. IFNG resolved into two distinct modes, corresponding to the two binding motifs identified by convergence analysis (Fig. 2d). Other targets, including mCherry and p65, occupied isolated positions, while XIAP, SIX1, and KIX are distributed throughout the central region. The spatial organization in Fig. 5d is determined by which binder-side features contribute most to binding each target. To interpret these features, we examined the top-ranked features for targets where known structural biology enables validation. InterPLM features vary in their interpretability. Some correspond to clearly annotatable structural or sequence motifs (“bright” features), while others detect compositional patterns that resist simple biological annotation (“dark” features) (**Methods**). For LC3B, the top features are predominantly bright and reconstruct the extended LIR ^42^ motif at scaffold position 15: the core aromatic-acidic pair (Trp at *i-1*, Glu at *i*), flanked by upstream Asp and Ala residues (Fig. 5e). For GABARAP, two of the top-ranked features are shared with LC3B, and the aromatic features capture both orientations of the motif (Fig. 5f). For Keap1, the top features fire at position 9, reconstructing the ETGE motif, the canonical motif through which Nrf2 engages the Kelch-repeat domain of Keap1 (Fig. 5g)^43^. Notably, in both motifs, some of the top features detect scaffold (invariant) residues such as 16-Ile and 12-Ala in the LIR and 8-Glu in the ETGE as components of the binding interface. For all three targets, dark features appeared among the top-ranked contributors (Fig. 5h). These features tended to activate broadly across the scaffold rather than at contact positions, capturing local, structural and framework-level determinants of binding that positional convergence analysis cannot resolve.

The target-side features operated at a different level of resolution, reporting on structural folds and functional sites (Fig. 5i). For LC3B and GABARAP, the leading target feature was shared: *f/5043* detected the ubiquitin-like beta fold common to both proteins. For Keap1, the dominant target feature (*f/10150*) corresponded to Kelch repeats, a β-propeller fold unique to Keap1 among the target panel. These trends were general across the panel (Supplementary Data 6): the top target-side features recovered the Walker A P-loop for NRAS, the zinc-coordinating CxxC motif of the LMO2 LIM domain, and the bHLH dimerization interface for both Myc and OLIG2. On the binder side, the top features predominantly detected local sequence motifs at the binding interface, with some capturing broader structural context. A notable exception is HPV-pE7, where the target features converged on an acidic disordered region while the binder-side features reconstructed an arginine dipeptide with predominant contribution from dark features that capture structural determinants.

## Generalization to natural proteins

The binding determinants recovered here, such as the ETGE degron for Keap1 and the extended LIR motif ^42,43^ for LC3B and GABARAP, were each established through independent structural studies of natural protein-protein interactions, yet emerge independently from synthetic affibody selections. The combination of biological selection through PANCS and contrastive training of the model produces a set of binding rules that, for these targets, converge on the same motifs. The agreement is at the level of the motif, not its structural presentation: Nrf2 engages Keap1 through the ETGE degron in a beta-hairpin^43^, a fold element absent from the affibody scaffold. At the same time, the features are not simply motif detectors. The top feature (*f/9431*) fired with high precision on TG dipeptides, though the dipeptide alone is not a sufficient condition for activation. The remaining top features, including dark features, captured additional structural determinants.

The feature dictionary comes from ESM-2 via InterPLM, both trained overwhelmingly on natural proteins, among which affibodies are a vanishing fraction (Z-domain homologs, such as affibody, constitute tens out of the ∼65 million sequences ESM-2 was trained on)^36,39^. As such, these features are unlikely to encode scaffold-specific information. The model expressed binding rules on a basis shaped by natural protein statistics, which constrains the hypothesis space. This motivates a direct test: can the learned features identify interactions in natural proteins unrelated to the training set? We scanned roughly 170,000 mammalian proteins in UniProt for sequences activating the top binder features for multiple targets (Fig. 5j**; Methods;** Supplementary Data 7). The search was highly selective, yielding 368 proteins across 22 of the 34 targets searched (Supplementary Data 8). For Keap1, it returned 3 proteins. In contrast, a sequence search for ETGE returned 872 mammalian proteins, indicating that the features discriminate beyond the linear motif. Most identified interactions were previously unreported. A notable exception was HPV-pE7, where the pipeline recovered fragments of PIH1D1 and NLRX1, both known direct interactors^44,45^.

We tested selected predictions using the split-RNAP luciferase complementation assay (Supplementary Fig. 4, Supplementary Fig. 26). Among the candidates, EIF4G1, a translation initiation factor not previously reported as a Keap1 interactor, carries an ETGE motif in the region identified by the feature search (Fig. 5k). We assayed a 47-residue truncation of EIF4G1 containing the predicted binding feature and observed clear binding (Fig. 5l). We chose multiple predictions for two targets as well as individual predictions across seven targets based on the highest number of features matched in the prediction and shared expected localization within the cell. Across 18 total predictions against PIK3CB, GABARAP, KIX, HPV-pE7 and Bcl2, 10 showed binding using our sensitive in vivo reporter assays (Supplementary Fig. 26). Whether these features transfer to targets absent from the training set, and to scaffolds other than affibody, remains to be tested. However, the recovery of binding partners among natural proteins suggests that the learned features capture binding determinants that extend beyond the synthetic scaffold.

## Discussion

In natural evolution, proteins must navigate sequence space through sequential mutations (co-evolution) or via high diversity mutations (V(D)J recombination followed by somatic hypermutation in antibody development). Our results suggest that, for the evolution of a *de novo* PPI, certain proteins with simple binding motifs that specify few residues can, through a limited number of mutations, easily acquire additional binding partners. In other words, these proteins appear poised for evolution to form new interactions. This idea is supported by the observation that the “easy” targets in our classification, such as KIX and LC3B, are known to bind many proteins in nature. Conversely, some targets contain complicated motifs (8+ amino acids).

Our ML results complement these experimental observations. The feature decomposition by the model reveals that bindability is not a single-axis property. For Keap1, recognition is governed by Kelch repeats presenting a specific pocket that engages ETGE-like motifs in a defined geometry with scaffold contributions. For the Atg8 proteins, binding is mediated through the LIR docking site, engaging a distinct aromatic-hydrophobic motif. For HPV-pE7, recognition instead relies on electrostatic complementarity between an acidic disordered region on the target and an arginine-rich motif on the binder, with the structural presentation of the scaffold playing a dominant role. Across all targets, the model’s target-side features are overwhelmingly specific, and cannot be captured by simple summarized descriptions such as % disorder. At the same time, the recovery and experimental validation of natural interaction partners indicates that the binding rules resolved here may not be confined to the synthetic context in which they were discovered.

Experimentalists have long understood that the success of a binder-discovery effort depends more on the target than on the discovery method. Binder density turns that intuition into a measurement, and the motif-frequency framework explains it: a target is easy or hard according to how complex its minimal binding motif is. This also tells us where effort should go. The simplest randomization, full diversity at every position on one face of the three-helix bundle, was nearly optimal; scaffold stability, not library diversity, appears to be the most critical feature (Supplementary Note 1). Because of this, although the outcome for any single target can look random, both in whether it succeeds and in what binders emerge, measuring enough interactions across diverse targets reveals interpretable patterns. Simple target descriptors did not predict this bindability a priori, but once selection data were in hand, a model could learn the features that govern binding, and those features predicted natural interaction partners (10 of 18 tested bound).

Binder density also provides a baseline for evaluating *de novo* design strategies: if thousands or more of designed variants are required to be tested experimentally to find one that works^18^, then designs must be evaluated in comparison to a similar size of random variants. We argue for robust benchmarking by evaluating the difficulty of a target using binder density with traditional scaffolds as well as the evaluation of designs from the top and bottom design quartiles of *in silico* metrics as a necessary first order approximation of whether a design strategy is efficacious or whether a target is simply easy. As binder design moves to hard targets, the ability to audit what a model learns and to learn from where it fails might matter as much as predictive accuracy.

## Methods

### Cloning Plasmids

All plasmids (Supplementary Table 11) were generated with Gibson Assembly (GA) of PCR fragments made with Q5 DNA polymerase (NEB) following the manufacturer’s protocol. Primers and target inserts were ordered from Integrated DNA Technologies (Supplementary Table 12). To make plasmids, GA reactions were transformed into DH10b chemically competent *E. coli* with the following protocol: Incubation on ice for 30 min, heat shock at 42°C for 45s, 2 min recovery on ice, and addition of 2xYT media and outgrowth with agitation at 37°C for 1h prior to plating on LB agar supplemented with kanamycin (40µg/mL in water), chloramphenicol (33µg/mL in EtOH), and carbenicillin (100µg/mL in water) as appropriate, and overnight incubation at 37°C. Correct insertion was confirmed via Sanger sequencing.

### Generating Bacterial Strains

To create multi-transformed bacterial strains for PANCS (+/-AP) and the split T7 RNAP *E. coli* luciferase assay (2-22/C-term-target/N-term-binder), S1030 *E. coli* strains were made chemically competent by preparing a 1/100 subculture of cells at OD600 ∼0.3, and pelleting and washing with cold CaCl2-glycerol buffer (60mM CaCl2, 10mM HEPES pH 7.0, 15% glycerol) twice prior to resuspension in the same buffer. Plasmids were transformed one at a time and transformed into single- or double-transformant chemically competent S1030 *E. coli* cells.

### Plaque Assays

To perform activity-independent plaque assays to measure titer, 2µL of phage was added to 100µL of S1030-1059 *E. coli* cells at OD600 ∼0.3, and a 50X serial dilution of each phage stock was performed by adding 2µL of the previous phage-*E. coli* mixture to the next, for a total of four dilutions. The mixture was combined with 750µL of 50°C top agar (7 g/L agar, 25 g/L LB) and plated evenly onto a four-quadrant bottom agar (15g/L agar, 15g/L LB) plate. After overnight incubation at 37°C, plaques were counted in the quadrant containing 10-200 plaques to determine the phage titer. The phage titer was calculated with the following equation:

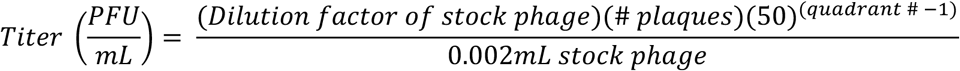

To perform activity-dependent plaque assays, the S1030-1059 strains were swapped for S1030 double-transformed +/- AP strains, consisting of the desired and a different target strain to confirm the presence or absence of on/off-target activity. In addition, to confirm the fidelity of the selections, plaque assays were performed with naïve S1030 strains to verify the absence of cheater phage in the absence of plaques.

### Production of Randomization Design Samplings

Affibody designs followed those of prior studies ^29,33,34,46,47^ and rational design. PCR conditions using Q5 DNA polymerase (NEB) were optimized for each design such that approximately 20-100µg product was formed at large scale, during which randomization schemes were introduced into an affibody template phage via primers containing degenerate codons (IDT) (**Supplementary Tables 11 and 12**). Following cleanup using the Wizard kit (Promega), restriction enzyme digestion with DpnI and NheI-NF (NEB), gel extraction and cleanup with the Zymo DNA clean and concentrate (Zymo DCC-5) kit, the digestion products were treated with T4 DNA ligase overnight. Ligated products were purified and concentrated with the Zymo DCC-5 kit prior to transformation via electroporation into S1030-1059 cells. The cells were added to 50mL SOC media warmed to 37°C and incubated with shaking for 2h, during which timepoints were collected to assess the titer and sampling size by plaque assay. At the end of the outgrowth, the cells were pelleted, and the supernatant was stored. To amplify the libraries, a 450mL S1030-1059 culture was grown to OD600 ∼0.6, and the phage supernatant was added. Following incubation for 6h, the cells were pelleted and the supernatant sterile-filtered and stored at 4°C. All libraries were subject to PANCS with naïve S1030 cells and a no-fusion negative control to ensure that libraries did not contain neither gIII nor N-term wild-type cheaters.

### Replication Assay

To determine the expression levels of new targets explored in this study in the PANCS system by measuring the phage replication rate, phage encoding the wild-type N-terminus of T7 RNAP were prepared to a diluted stock of 500 PFU/µL. 1000µL of phage were added to 1mL subcultures of S1030 transformed with each +AP in stationary phase, and incubated for 12h at 37°C with agitation. A no-culture (i.e. media only) control was included as a negative control. The cells were then pelleted, and the titer of supernatant was measured via plaque assay or qPCR. To calculate the replication rate of each target in the PANCS system, the titer of each output was divided by that of the media-only negative control.

### PANCS

PANCS was performed in deep 96-well plates in experiments where 10^8^ or fewer unique variants were tested. 1/10 subcultures of S1030 *E. coli* cells double-transformed with +/- APs were grown to OD600 ∼0.5, then seeded with the number of unique variants of interest (10^5^ -10^8^ PFU) in the first passage to obtain a final volume of 1mL per well and incubated with shaking for 12h. To continue the PANCS for a total of 6 passages, 5% of phage supernatant (50µL of 1mL) were passaged to the next round of selection following the same procedure.

To create our 10^10^ unique variant sampling of Design A, eight separate transformations (of ∼1-2*10^9^ unique variants each) were combined to generate a composite, 10^10^ sampling of Design A and used to inoculate 500mL cultures of +/-AP S1030 strains, as described before^14^. Subsequent passages were performed in progressively smaller volumes: 100mL (P2), 15mL (P3), and 5mL (P4+), while maintaining both a 1:1 phage-to-cell ratio and 5% passaging rate.

### qPCR

qPCR was utilized to determine the endpoint titer in a high-throughput manner following each PANCS experiment. The qPCR was optimized by testing various sets of primers that recognized different sites in the phage genome and determining the linear range and accuracy of the assay with the different primers and dilutions of phage. The primers VC-525 and VC-526 were selected to amplify a small section of the M13 phage gIII for assay generalizability to different libraries (Supplementary Table 12). 20µL qPCR reactions were assembled in the following way: 3µL water, 1µL each primer at 5µM, 10µL Power Up SYBR Mix (ThermoFisher, Cat#: A25777), and 5µL PANCS endpoint phage diluted 200X. The qPCR reactions were run on a QuantStudio6Pro with the following protocol: 95°C 10m denaturation, 40 cycles of 95°C 20s, 60°C 20s, and 72°C 20s, final elongation 72°C 20s, then a melt curve protocol of 95°C 10s, 65°C 1m, and 97°C 1s to ensure that only one amplicon exists. Each qPCR run included a standard curve of phage with known titers confirmed by plaque assay, later used to calculate the endpoint titers of each selection.

### Next Generation Sequencing (NGS)

NGS samples were prepared by appending adaptor and barcode sequences to PCR amplicons starting from the linker sequence and ending just downstream of the stop codon positioned following the affibody scaffold, using primers MS-1144-MS-1156 and MS-1346-1357 (Supplementary Table 12). The PCR protocol consisted of a 10 min initial denaturation step at 98°C, and 30 cycles of 15 s 98°C denaturation, 15 s 63 °C annealing, and 1 min 72 °C extension steps, and a 5 min final 72 °C extension step. The PCR products were pooled with other barcodes within the same NGS sample and purified with the Zymo DCC-5 kit prior to measuring their concentrations with the Qubit dsDNA High Sensitivity kit (ThermoFisher). Each NGS sample was diluted to 20ng/µL and 500ng total barcoded PCR products were submitted for analysis with the Amplicon EZ service at Genewiz (Azenta). Sequencing data was analysed by BB Merge to merge all paired end reads, then MATLAB was used to separate the results for each selection by barcode (Step1) and translate the sequences (Step2)^48,49^.

### Split-T7 RNAP *E. coli* Luciferase Assays

We followed the split-RNAP *E. coli* luciferase assay protocol detailed in our previous publication^14^. In summary, a double-transformant S1030 strain (2-22/C-term-target lux) was made chemically competent as described above and transformed with N-term binder and non-binder plasmids (**Supplementary Tables 11 and 12**). Colonies for each condition were picked and grown in 1mL of LB media with kanamycin, chloramphenicol, and carbenicillin overnight at 37°C with shaking. The next day, 1/20 subcultures for each colony were prepared in LB media supplemented with antibiotics and L-arabinose (2 mg/mL) and cultured for 3.5 h before measuring OD600 and luminescence signals using a BioTek Synergy Neo2 plate reader. To compare the activities of binders and non-binders, we computed the quotient of luminescence signal and OD600 and normalized all binder signals to those of the non-binder signals.

### Determination of putative hits for further validation

qPCR was performed following the final, sixth passage of PANCS to detect high titers, and selections with titers exceeding 1.0 × 10^7^ PFU/mL were deemed to contain putative binders. These selections were submitted for NGS analysis, and after data processing in MATLAB to assess the most enriched and relative abundances of the best variants. Of note, selections where a truncated variant (i.e. containing premature stop codon) was enriched was not considered for further analysis. Of the selections where an affibody variant was enriched, the top affibody variant or most abundant variant that could be identified, was subcloned into a N-term T7 RNAP plasmid for testing in our three-plasmid split T7 RNAP *E. coli* luciferase assay, to validate binding activity of the affibody with its target.

### Clustering and Fitness Analysis

For each collection of binders (data presented in **Fig. 2** and **Fig. 3** individually), sequences were one-hot encoded in the variable positions; PCA was performed on the one-hot encoded representation and the top 100 PCs were used to compute the 2-dimensional t-SNE embedding. We computed the associated Hamming distance matrix and performed hierarchical clustering with Ward linkages. For the 9-target panel (Fig. 2d-g) we used 30 clusters, and for the 96-target panel (Fig. 2h-j) we used 300 clusters (Supplementary Fig. 12). Furthermore, using the Hamming distance and the top 100 PCs resulted in identical clusters.

From the experimentally measured fold-change of top variants for each target in each sampling, we extrapolated the fitness of other variants identified in the NGS of that selection using the ratios of relative concentrations linearly:

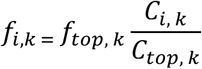

Where and *C_top_*, *_k_* are the relative concentrations and of variant *i* and the top variant; *f_i_*, *_k_* and *f_top_*, *_k_* are the extrapolated fitness for variant *i* and the measured fitness for the top variant. The total fitness is the sum of fitness for binders within a cluster and the normalized fitness is normalized over the number of binders.

### Amino acid distribution analysis

We followed a two-step approach to analyse the position-dependent amino acid usage at randomized positions in the affibodies using the Python libraries pandas, numpy, itertools, and Bio. First, the expected frequency of each amino acid for a given codon position was computed based on the design strategy. For each codon position, we considered all possible codons that could be generated from the nucleotides at each position, and the expected percent frequency of each amino acid was calculated by determining the fraction of codons that encode each amino acid. Observed percent frequencies were determined from the NGS binder sequences as reported in Step2 analysis, where we considered only sequences >1% abundance in their respective selections. The percent difference was calculated for each amino acid at every degenerate codon position using the formula:

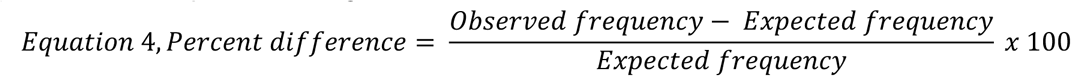

To assess the usage bias of each amino acid across the degenerate codons NNK and NNY, we identified positions in our designs that contained NNK or NNY and aggregated the percent differences of observed/expected amino acid frequencies across all binder sequences within a given design and across all designs.

### Statistical Analysis between Target Properties and Binder Density

The statistical analyses: Pearson’s and Spearman’s correlations, Kruskal-Wallis test for multi-group target comparisons, Dunn’s post-hoc pairwise tests, log-normal curve fitting, and multiple linear regression were all performed in GraphPad Prism 10 to assess meaningful relationships between target properties and binder density, analyse class-wide influences on target difficulty, and understand the relationship between sequence space diversity and hit rate, respectively. To process the split-T7 RNAP *E. coli* luciferase assay and qPCR data, Microsoft Excel was used to calculate fold changes in luminescence/OD600 over negative control signal and convert Cq values to titers based on a standard curve of phage with known PFU/mL. Target characteristics (Supplementary Table 8) were derived from AlphaFold 3 predictions^50^, NetSurfP-3.0^51^ and UniProt/MobiDB annotations.

### Data Preprocessing for Machine Learning

NGS data from PANCS provided affibody binder sequences for each target protein studied. All binder sequences returned from NGS were analysed, and their reads were collapsed by amino acid sequence to account for codon degeneracy. We removed sequences that were not exactly 58 amino acids in length or lacked the required affibody framework regions (N-terminal VDNK and C-terminal DPSQSANLLAEAKKLNDAQAPK). Relative abundance was defined as the number of reads for each unique protein sequence divided by the total sequence count for that selection. Across all targets and libraries in the multi-design screen (Designs A–J, Fig. 3b), sequences with relative abundance ≥0.5% were classified as binders. Negatives were defined as sequences appearing in NGS for other targets that do not satisfy the ≥0.5% threshold.

### Training Details for the ML Model

Each binder and target sequence was passed through a frozen ESM-2-650M model^36^ to obtain a contextualized embedding for each residue. We denote the binder and target embeddings as *b* ∈ ℝ*^N^*^b×1280^ and *t* ∈ ℝ*^N^*^t×1280^, where *N*_b_ = 58 for all affibody variants and *N*_t_ varies by target. We score the interaction between each binder residue and target residue through a bilinear form with *W* constrained to rank *d*. This is done by defining matrices *W*_B_ and *W*_T_ of dimension 1280 × *d*, and the interaction score between binder residue *i* and target residue *j* as:

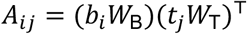

The log-odds of binding between a binder and target is defined as a soft maximum over all residue pairs:

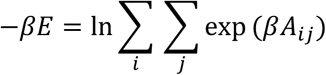

where *β* = 1/√*d*, a scaling factor that prevents the dot product from growing with *d*. Since positives make up a small fraction of the selection data, we train the model using an InfoNCE loss^37^, which scores each positive interaction between a target and its positive binder (with embedding *b*^+^) as:

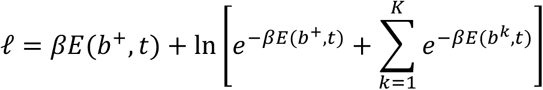

where *b^k^* runs over *K* = 100 negatives, redrawn every epoch. The loss reweights negatives so that those closest to the positive are penalized most. At optimality the score recovers the log density ratio between the binding and non-binding populations, up to an additive function of the target. The data is split into five cross-validation folds and stratified so that the same ratio of positive to negative binders is maintained across folds. For each fold, we trained models at *d* ∈ { 1,6,10,32,64,8 × 4}. The 8 × 4 configuration uses 4 independent interaction matrices of rank 8, each with its own *W*_B_ and *W*_T_, analogous to multi-head attention^52^. All models were trained with the Adam optimizer (lr = 10^−3^) for 500 epochs with a batch size of 240 positives, and the gradient norm was clipped at 1.0 to stabilize training. All models were trained on 4 A100 GPUs.

Evaluations use the model at the final checkpoint of each run, and are computed over every positive and negative in the test set. Model performance is evaluated by macro ROC-AUC, defined as the per-target ROC-AUC averaged across all targets. This metric measures how well the model discriminates binders from non-binders for each target independently, weighting all targets equally regardless of the number of positives. Macro ROC-AUC and parameter counts for each model configuration are reported in Supplementary Table 10 and plotted in Fig. 4c, with error bars showing the standard deviation across the five folds. The per-target AUC and score distributions are given in Supplementary Fig. 25.

Details of the baseline model are available in Ref. 16.

### Feature Decomposition of the Log-Odds of Binding

For a single binder residue embedding *b_i_*, a sparse autoencoder (SAE) encodes it into a sparse, higher-dimensional representation:

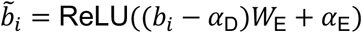

where *α*_D_ and *α*_E_ are biases, and *W*_E_ is the encoder weight matrix, which for InterPLM^39^ takes the dimension 1280 × 10240. The encoder expands the 1280-dimensional embedding by a factor of 8 into a non-negative representation *b*^j^*_i_* ∈ ℝ^10240^. Each element of *b*^j^*_i_* is referred to as a feature. SAEs are trained with a decoder that linearly maps the overcomplete representation back to the original embedding:

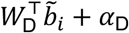

where *W*_D_ ∈ ℝ^10240×1280^ for InterPLM. SAEs are trained to reconstruct the original embedding from the overcomplete representation, with a penalty that enforces sparsity on the overcomplete basis *b̃_i_*. Under zero reconstruction error for binders and targets, 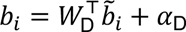 and 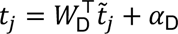. In this case, the contact score *A_ij_* can be written as:

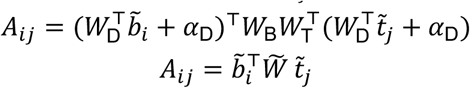

where *b̃_i_* and *t̃_j_* are augmented with a constant to absorb the decoder bias, and *W̃* ∈ ℝ^10241×10241^ is the feature-feature interaction matrix, with rank bounded by *d*.

The form above holds under zero reconstruction error. We train a variant of the model with *d* = 64, where the binder and target embeddings are given by the reconstructed embeddings. This model achieves a macro ROC-AUC of 0.81 ± 0.02 (Supplementary Table 10), with the gap attributable to reconstruction error, but makes the feature-level analysis exact.

To show this, we first note that the log-odds is a nonlinear function of the contact scores, so individual contributions cannot be read off directly. However, it can always be rewritten as a linear combination of contact scores under a distribution *p_ij_*, with the nonlinearity absorbed into an entropy term:

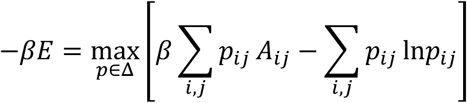

where Δ is the set of all values of *p_ij_* that define a valid probability distribution, i.e. *p_ij_* ≥ 0 and ∑*_i_*_,j_ *p_ij_* = 1. Here, we define *p_ij_* as a probability distribution over all binder-target residue pairs, representing how much each pair contributes to the overall binding score. The value of *p_ij_* that maximizes the equation is unique and given by the Boltzmann distribution over contact scores,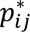 :

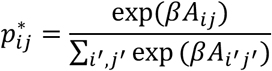

Hence, the log-odds separates into two terms:

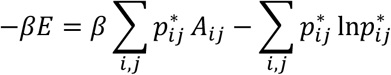

The first term is the average contact score over all residues weighted by *p*^∗^, and can be exactly decomposed into per-feature-pair contributions. We define *I_mn_* as the contribution of binder feature *m* paired with target feature *n* to the weighted contact score:

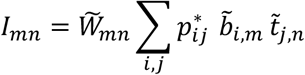

Here *W̃_mn_* is an element of the interaction matrix that tells us how strongly the model couples binder feature *m* with target feature *n*, *b̃_i,m_* is how active feature *m* is at binder residue *i*, *t̃_j,n_* is how active feature *n* is at target residue *j*, and 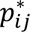 tells us how important the residue pair (*i*, *j*) is to binding. Marginalizing over target features gives the total contribution of a single binder feature, *I_m_* = ∑*_n_ I_mn_*.

The second term, 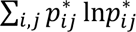, is a nonlinear function of all the contact scores, but is bounded between 0 and ln(*N*_b_ ⋅ *N*_t_). The two bounds reflect limiting cases: when the model’s prediction depends on a single residue pair, the term is 0, and when all residue pairs contribute equally, it reaches ln(*N*_b_ ⋅ *N*_t_). In practice, we find this term is near zero, so the weighted contact score, which *I_mn_* decomposes exactly, dominates the log-odds.

The marginal contributions *I_m_* rank which binder-side features the model relies on for each prediction. However, this ranking holds at fixed *p*^∗^: it assigns contributions for a given prediction, but removing a feature can change *p*^∗^ itself. To understand how the contributions relate to the effect of removal, we analyse the change in *p*^∗^ when individual features are set to zero. Formally, this can be understood as setting *b̃_i,m_* = 0 for all residues *i* and all *m* ∈ *M*. Defining −*βE′* as the log-odds after removal of these features, one can show that the change in log odds takes the form:

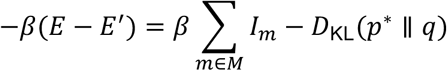

where *q* is the Boltzmann distribution after removal, and *D*_KL_ is the KL-divergence between the two distributions. Since KL-divergence is a non-negative quantity, we have −*β*(*E* − *E′*) ≤ *β* ∑*_m_*_∈*M*_ *I_m_*. In other words, the marginals overestimate the change in log odds after removal of features, and the equality holds when their removal does not reshuffle which residue pairs dominate the prediction. In our case, *p*^∗^ concentrates on a single residue pair, so the KL term collapses to −ln*q_i_*_∗j∗_, where (*i*^∗^, *j*^∗^) is the dominant residue pair under *p*^∗^. Hence, the bound is tight when removing the features does not modify the top contact position.

The dominant contact is a property of a single binder–target pair, and each target can have binders with different dominant residue-pairs. For all of our analyses, we first group binders that share the same dominant residue-pair contact, then rank features within each group. Per-cluster feature decompositions, cluster sizes, and contact pairs are provided in Supplementary Data 6.

### InterPLM Feature Annotation using Greedy Search

Sparse autoencoders are trained to reconstruct a PLM’s embeddings under a sparsity constraint. After training, the features need to be annotated to interpret what the dictionary corresponds to. Two approaches are typically used to annotate SAE features. In the first, each feature is assigned the single concept from a structured database (e.g. UniProt) whose presence best predicts feature activation^17^. In the second, a language model describes what each feature detects from its activation pattern, and the description is scored against held-out activations. Simon & Zou applied both methods to annotate the features of InterPLM trained for ESM-2-8M. For the SAE for ESM-2-650M that is used in this work.

For all analyses in the main text, we annotated all 10,240 features de novo against 515,725 reviewed Swiss-Prot proteins (164,913,378 total residues). For each feature, we find the combination of UniProt annotations (Domain, Active site, Binding site, Motif, Region, Modified residue) and local sequence context that best predicts feature activation.

The search proceeds in three stages. In the first, we replicate Simon & Zou’s single-concept method: each feature is matched to the one UniProt annotation that best predicts its activation. A match is scored by how often the feature fires within annotated regions (precision) and how often annotated regions contain a firing residue (recall). The threshold for firing is tuned on a validation set and all metrics are evaluated on separate held-out proteins. We consider two metrics. F1, the harmonic mean of precision and recall, measures how well a single concept accounts for the feature’s firing pattern. We also consider precision alone, which identifies concepts that are necessary but not sufficient for firing. A high precision tells us the concept is always present when the feature fires, but does not fully explain it, i.e. the feature requires additional conditions to activate.

We repeat the same matching: for each feature, we find the single best predictor of activation, now in terms of which amino acids appear at and around the active position. We find many features where a particular amino acid consistently appears at a fixed offset from the location where a feature fires, for example glutamate at two residues to the right. This is revealing in that it shows how PLMs capture context of a protein at a single residue. At the same time, recall and F1 are low: glutamate at position +2 appears in many places where the feature does not fire. The feature detects something more specific and requires several such conditions to hold at once.

The combination of amino acids at fixed positions defines a sequence motif, and the results above indicate that many features could be resolved by matching them to motifs. The space of possible motifs, however, grows combinatorially with the number of positions and amino acid types, making an exact search intractable. Here, we use a greedy search. Starting from the best single predictor of firing, we test every remaining amino acid–position pair and annotation as an additional requirement. If the best addition improves F1, we include it and repeat, up to a maximum of 12 steps. All validations are performed on the held-out set.

Across all stages, the search resolved 1,926 features at held-out test F1 ≥ 0.5, and identified 1,139 unique biological concepts. Simon & Zou reported 427 concepts across six layers of ESM-2-650M using single-concept matching alone.

The feature annotations are independent of any binding measurements, since the entire search is run on Swiss-Prot proteins with no overlap to the affibody binders. For the feature labels in **Fig. 5**, a feature was assigned its highest-scoring concept if per-residue precision ≥ 0.70 or held-out test F1 ≥ 0.30. Features meeting neither threshold are reported as dark features.

All annotations are included in Supplementary Data 5.

### LLM-based Annotations

As an independent check, we also ran the standard automated interpretability protocol^39^ on the same features. We reimplemented the protocol that Simon & Zou^39^ used for annotating ESM-2-8M, in which a language model describes what each feature detects from its activation pattern, and the description is scored against held-out activations.

To construct the input sets, each protein from Swiss-Prot is scored by its highest activation across all residues. These scores are normalized to [0, 1] and binned into ten equal-width bins. Two proteins are sampled per bin from 0.0 to 0.8, twelve from each of the two top bins, and ten zero-activation negatives, giving 38 per set. These examples are sorted by activation and assigned alternately to a description set and a held-out validation set. Features with fewer than 20 proteins above 0.7 are excluded.

During the description stage, an LLM sees the description-set proteins along with their Swiss-Prot annotations, maximum activation values, most-activated residue positions, and the amino acids at those positions. It is asked to describe what the feature detects and to summarise it in one sentence. During validation, a separate LLM receives the description and a set of proteins with their Swiss-Prot annotations, but no activation values. The model predicts how strongly the feature fires on each protein, and these predictions are compared to the measured values by Pearson correlation.

All annotations are produced with Claude Opus 5, using adaptive thinking at high effort and a maximum output of 100,000 tokens. Each response is checked before saving, and incorrectly formatted responses are discarded. About 22% were discarded on the first pass. We reran the setup for these failures, up to four times, to make sure all features were annotated.

The protocol was run on 1,680 of 1,696 features across the same protein set. Note that a description that can classify whether the feature fires or not, without any information about the magnitude of the response, can already reach r = 0.705.

These annotations are included in Supplementary Data 5, along with the deterministic ones.

### Feature-Based Proteome Search

Of the 54 targets in the training set, 45 had at least 10 positive binders. 34 of these targets produced binding-mode clusters with at least 6 positive binders, yielding 65 clusters in total. Feature importances I_m are computed from the reconstructed-embedding model (d=64, fold 1, checkpoint selected by best test AUC). For each cluster, we ask which SAE features are most consistently used for binding. Each positive binder nominates the 10 features with the largest importance I_m, features are ranked by how many binders nominate them, and the top 10 are retained. For each feature in this top-10 set, the activation threshold was set to the lowest non-zero activation of that feature across the cluster’s binders. The features and thresholds used for the search for each cluster are provided in Supplementary Data 7.

The search database comprised 172,872 human, mouse, and rat proteins (56,385,515 residues) from UniProt (downloaded 2026-03-30). Each protein was embedded with ESM-2 650M (layer 33) and decomposed through the InterPLM SAE. Each cluster’s top-10 features defined an independent search. For every residue in the database, we count how many of the 10 features have activation above their threshold. A residue is a hit if ≥8 of the 10 are active. Across all 65 clusters and 172,872 proteins, the search identified 383 hit residues across 368 distinct proteins. The median per-target hit rate across all 34 targets was 1 protein in 138,298; 12 targets returned no mammalian hits. Among the 22 targets with hits, the IQR ranged from 1 in 25,145 to 1 in 125,725. All hits are listed in Supplementary Data 8.

## Data availability

Raw NGS reads for all selections have been deposited in the NCBI Sequence Read Archive under BioProject [accession to be added]. Processed sequences, clustering results, feature annotations and proteome-search hits are provided as Supplementary Data 1-8. Source data are provided with this paper. Plasmids are available from Addgene [accession to be added].

## Code availability

Code for the interaction model, feature decomposition, concept-discovery pipeline and proteome search, together with trained model weights, is available at [GitHub/Zenodo DOI to be added].

## Acknowledgements

We thank the members of the Dickinson, Vaikuntanathan, and Dinner labs for providing constructive and insightful discussions about this study. We thank the University of Chicago Sanger Sequencing Core and Azenta’s Genewiz Amplicon-EZ service for sequencing support. We thank S. Ahmadiantehrani for assistance with manuscript preparation. A.N.S. acknowledges Anthropic for API credits through the AI for Science program. This work used resources provided through ACCESS allocation CIS261309, supported by the National Science Foundation.

## Author contributions

Conceptualization: SSL, MJS, CFG, ANS, AN, ARD, SV, BCD. Funding acquisition: SSL, MJS, CFG, ANS, AN, ARD, SV, BCD. Investigation: SSL, MJS, CFG, ANS, JY, EP, AN, JAP, ST. Methodology: SSL, MJS, CFG, AN, CB, JAP, ST. Conceived and developed the ML interaction model, feature decomposition, concept-discovery pipeline and the proteome search; performed all ML analysis; wrote the ML sections of the manuscript: A.N.S. Developed earlier ML model and data curation: CFG and AN. Supervision: ARD, SV, BCD. Writing (original draft): SSL, MJS, CFG, ANS, BCD. Writing (revision, review, & editing): SSL, MJS, CFG, ANS, AN, ARD, SV, BCD

## Additional information

Supplementary Information is available for this paper.

