## Supplementary Information for "Learning interpretable rules of protein binding from a trillion interactions"

### Table of Contents

**Supplementary Fig. 1** – Design/Construction of Affibody Library A
**Supplementary Table 1** – Details of Proteins in the Target Panel
**Supplementary Fig. 2** – Endpoint Titers from 9-Target Panel Selections
**Supplementary Fig. 3** – NGS Results from 9-Target Panel Selections
**Supplementary Fig. 4** – E. coli Binding Luciferase Assay for the 9-Target Panel Selections
**Supplementary Table 2** – Binder Density for the Proteins in the 9-Target Panel
**Supplementary Fig. 5** – Endpoint Titers from 96-Target Panel Selections
**Supplementary Fig. 6** – E. coli Binding Luciferase Assay for the 96-Target Panel Selections
**Supplementary Fig. 7** – Hierarchical Clustering of Binders for the 9-Target Panel Selections
**Supplementary Fig. 8** – Comparison of binder fitness across varied sampling size
**Supplementary Fig. 9** – Consistency in Fitness Landscape across Replicate Samplings
**Supplementary Fig. 10** – Identifying Binding Motifs from Prominent Clusters
**Supplementary Fig. 11** – Comparison of Motif Frequency to Motif Density for 9-Target Panel
**Supplementary Fig. 12** – Optimal Choice of Clustering Constraints
**Supplementary Fig. 13** – Characteristics of Prominent Clusters
**Supplementary Table 3** – Binding Motifs from Clusters
**Supplementary Fig. 14** – Unique Sequences per Cluster-Target Pair
**Supplementary Table 4** – Additional Library Construction – Libraries B-J
**Supplementary Table 5** – Possible Amino Acids in each Randomized Position in Each Library

**Supplementary Fig. 15** – Overlap in Library Design Sequence Space
**Supplementary Fig. 16** – Endpoint Titers from Varied Randomization Design Selections
**Supplementary Fig. 17** – Overlap in Hits between Randomization Designs
**Supplementary Fig. 18** – Comparing Hit Fitness between Randomization Designs
**Supplementary Fig. 19** – Expression Level of Random Library Variants
**Supplementary Fig. 20** – Additional Codon Use Comparisons
**Supplementary Table 6** – Targets sorted into Binder Density Categories
**Supplementary Table 7** – Grouping Target Panel by Functional/Structural Features
**Supplementary Fig. 21** – Comparing Binder Density Category across Target Groups
**Supplementary Table 8** – Individual Characteristics of each Target
**Supplementary Fig. 22** – Comparing Binder Density Category across Target Characteristics
**Supplementary Fig. 23** – Solvent-Exposed Amino Acid Distribution by Binder Density Category
**Supplementary Table 9** – Result of Multiple Linear Regression (Characteristics vs Bindability)
**Supplementary Fig. 24** – PCA of Characteristics for Binder Density Categories
**Supplementary Table 10** – Model Configurations, Parameter Counts, and Macro ROC-AUC
**Supplementary Fig. 25** – Per-Target Log Odds Distributions and Classification Performance
**Supplementary Fig. 26** – Assessment of Predicted Interactions
**Supplementary Table 11** - Plasmids used in this work
**Supplementary Table 12** - Primers used in this work
**Supplementary Note 1** – Clustering, convergence and library design
**References**

**Supplementary Data files**

**Supplementary Data 1** – NGS from 9-target panel selections
**Supplementary Data 2** – NGS from 96-target panel selections
**Supplementary Data 3** - Clustering for 9-target panel NGS
**Supplementary Data 4** - Clustering for 96-target panel NGS
**Supplementary Data 5** – Annotations of all SAE features referenced in this work (both deterministic and LLM-based)
**Supplementary Data 6** – Per-cluster feature decomposition
**Supplementary Data 7** – Activation threshold for the top-binder side features used for the proteome search
**Supplementary Data 8** – All hits from the proteome search

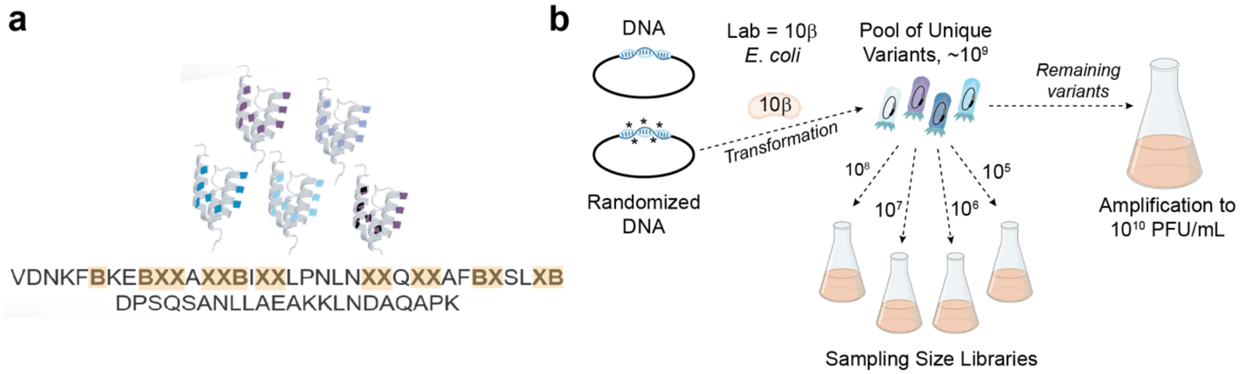

**Supplementary Fig. 1: Constructing independent libraries of varied sample size. a**, The affibody scaffold is randomized on one face of the three helix bundle along helix 1 and 2. The amino acid sequence of affibody is shown below with the randomized positions highlighted as B (biased mutations) and X (NNY, all types of amino acids, but no W, K, Q, E, or M) – this is the randomization design for affibody that we previously reported[1]. **b**, Varied, independent sampling of the library sequence space is produced by transforming plasmids containing the randomized DNA into *E. coli* to produce phage with those random variants. We assess the total number of variants produced by a single transformation, and then separate off different numbers of variants to amplify to produce an individual sampling of a defined number of unique variants. Independent transformations produce a unique set of variants such that each individual library of the same size has little to no overlap in which variants are in the sampling.

**Supplementary Table 1: Details of Target Panel Proteins.** Grey shading for 9-target panel.

| Target | AA Sequence | AlphaFold3 Structure | Other Notes |
|---|---|---|---|
| Very high (pLDDT > 90) | Confident (90 > pLDDT > 70) | Low (70 > pLDDT > 50) | Very low (pLDDT < 50) |
| **9-target panel** | | | |
| KIX | MGNIGSLSTIPTAAPPSSTGVRKGWH EHVTQDLRSHLVHKLVQAIFPTPDPA ALKDRRMENLVAYAKKVEGDMYESA NSRDEYYHLLAEKIYKIQKELEEKRRS RL | 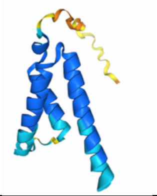 | Uniprot: Q92793. Human, nucleus, transcription factor, CREB Binding Protein (CBP) KIX domain (568-673) |
| PP2A-B" (622-1150) | MQILQETLTTSSQANLSVCRSPVGDK AKDTTSAVLIQQTPEVIKIQNKPEKKP GTPLPPPATSPSSPRPLSPVPHVNNV VNAPLSINIPRFYFPEGLPDTCSNHE QTLSRIETAFMDIEEQKADIYEMGKIA KVCGCPLYWKAPMFRAAGGEKTGF VTAQSFIAMWRKLLNNHHDDASKFIC LLAKPNCSSLEQEDFIPLLQDVVDTH PGLTFLKDAPEFHSRYITTVIQRIFYTV NRSWSGKITSTEIRKSNFLQTLALLEE EEDINQITDYFSYEHFYVIYCKFWELD TDHDLYISQADLSRYNDQASSSRIIER IFSGAVTRGKTIQKEGRMSYADFVWF LISEEDKRNPTSIEYWFRCMDVDGD GVLSMYELEYFYEEQCERMEAMGIE PLPFHDLLCQMLDLVKPAVDGKITLR DLKRCRMAHIFYDTFFNLEKYLDHEQ RDPFAVQKDVENDGPEPSDWDRFA AEEYETLVAEESAQAQFQEGFEDYE TDEPASPSEFGNKSNKILSASLPEKC GKLQSVDEE | 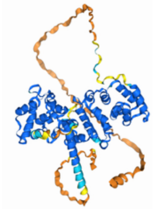 | Uniprot: Q06190. Human, cytoplasm, regulatory subunit of a phosphatase, primarily the structured domain. |
| CDKN1A | MSEPAGDVRQNPCGSKACRRLFGP VDSEQLSRDCDALMAGCIQEARERW NFDFVTETPLEGDFAWERVRGLGLP KLYLPTGPRRGRDELGGGRRPGTSP ALLQGTAEEDHVDLSLSCTLVPRSGE QAEGSPGGPGDSQGRKRRQTSMTD FYHSKRRLIFSKRKP | 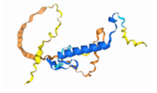 | Uniprot: P38936. Human, cytoplasm/nucleus, kinase inhibitor involved in cell cycle, intrinsically disordered, full length. |
| LC3B | MPSEKTFKQRRTFEQRVEDVRLIRE QHPTKIPVIIERYKGEKQLPVLDKTKF LVPDHVNMSELIKIIRRRLQLNANQAF FLLVNGHSMVSVSTPISEVYESEKDE DGFLYMVYASQETFGMKLSV | 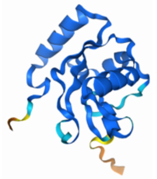 | Uniprot: A6NCE7. Human, cytoplasm/autophagosome membrane, autophagy cargo receptor, full length. |
| BTK (PH) | MAAVILESIFLKRSQQKKKTSPLNFKK RLFLLTVHKLSYYEYDFERGRRGSKK GSIDVEKITCVETVVPEKNPPPERQIP RRGEESSEMEQISIIERFPYPFQVVY DEGPLYVFSPTEELRKRWIHQLKNVI R | 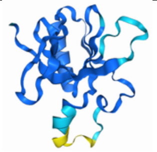 | Uniprot: Q06187. Human, cytoplasm, kinase, PH domain (1-133). |

| | | | |
|---|---|---|---|
| PP2A-B | MAGAGGGNDIQWCFSQVKGAVDDDVAEADIISTVEFNHSGELLATGDKGGRVVIFQQEQENKIQSHSRGEYNVYSTFQSHEPEFDYLKSLEIEEKINKIRWLPQKNAAQFLLSTNDKTIKLWKISERDKRPEGYNLKEEDGRYRDPTTVTTLRVPVFRPMDLMVEASPRRIFANAHTYHINSISINSDYETYLSADDLRINLWHLEITDRSFNIVDIKPANMEELTEVITAAEFHPNSCNTFVYSSSKGTIRLCDMRASALCDRHSKLFEEPEDPSNRSFFSEIISSISDVKFSHSGRYMMTRDYLSVKIWDLNMENRPVETYQVHEYLRSKLCSLYENDCIFDKFECCWNGSDSVVMTGSYNNFFRMFDRNTKRDITLEASRENNKPRTVLKPRKVCASGKRKKDEISVDSLDFNKKILHTAWHPKENIIAVATTNNLYIFQDKVN | 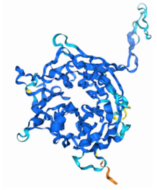 | Uniprot: P63151. Human, cytoplasm, regulatory subunit of a phosphatase, full length. |
| IFNG (24-156) | MQDPYVKEAENLKKYFNAGHSDVADNGTLFLGILKNWKEESDRKIMQSQIVSFYFKLFKNFKDDQSIQKSVETIKEDMNVKFFNSNKKKRDDFEKLTNYSVTDLNVQRKAIHELIQVMAELSPAAKTGKRKRSQ | 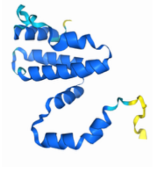 | Uniprot: P01579. Human, secreted, type II interferon, structured secreted domain only. |
| XIAP | MTFNSFEGSKTCVPADINKEEEFVEEFNRLKTFANFPSGSPVSASTLARAGFLYTGEGDTVRCFSCHAAVDRWQYGDSAVGRHRKVSPNCRFINGFYLENSATQSTNSGIQNGQYKVENYLGSRDHFALDRPSETHADYLLRTGQVVDISDTIYPRNPAMYSEEARLKSFQNWPDYAHLTPRELASAGLYYTGIGDQVQCFCCGGKLKNWEPCDRAWSEHRRHFPNCFFVLGRNLNIRSESDAVSSDRNFPNSTNLPRNPSMADYEARIFTFGTWIYSVNKEQLARAGFYALGEGDKVKCFHCGGGLTDWKPSEDPWEQHAKWYPGCKYLLEQKGQEYINNIHLTHSLEECLVRTTEKTPSLTRRIDDTIFQNPMVQEAIRMGFSFKDIKKIMEEKIQISGSNYKSLEVLVADLVNAQKDSMQDESSQTSLQKEISTEEQLRRLQEEKLCKICMDRNIAIVFVPCGHLVTCKQCAEAVDKCPMCYTVITFKQKIFMS | 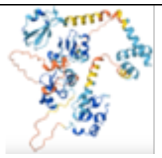 | Uniprot: P98170. Human, cytoplasm, nucleus, E3 ligase, full length |
| MAX (23-87) | MDKRAHHNALERKRRDHIKDSFHSLRDSVPSLQGEKASRAQILDKATEYIQYMRRKNHTHQQDIDDLKRQNALLEQQGEHP | 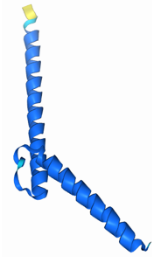 | Uniprot: P61244. Human, nucleus, transcription factor, intrinsically disordered, DNA binding domain only. |
| **Extended 96-target panel (includes 9-target panel above)** | | | |

| | | | |
|---|---|---|---|
| KRAS (G12D) | MTEYKLVVVGADGVGKSALTIQLIQNHFVDEYDPTIEDSYRKQVVIDGETCLLDILDTAGQEEYSAMRDQYMRTGEGFLCVFAINNTKSFEDIHHYREQIKRVKDSEDVPMVLVGNKCDLPSRTVDTKQAQDLARSYGIPFIETSAKTRQGVDDAFYTLVREIRKHKEKMSKDGKKKKKKSKTKCVIM | 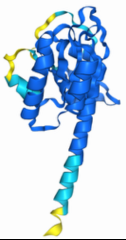 | Uniprot: P01116. Human, cytoplasm, GTPase, oncogene, full length isoform 4b G12D mutant. |
| ZB | MASEQLEKKLQALEKKLAQLEWKNQALEKKLAQ | 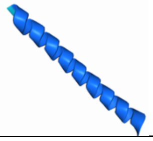 | Synthetic construct; PMID: 15631464 |
| p65 (17-293) | MSGPYVEIIEQPKQRGMRFRYKCEGRSAGSIPGERSTDTTKTHPTIKINGYTGPGTVRISLVTKDPPHRPHPHELVGKDCRDGFYEAELCPDRCIHSFQNLGIQCVKKRDLEQAISQRIQTNNNPFQVPIEEQRGDYDLNAVRLCFQVTVRDPSGRPLRLPPVLSHPIFDNRAPNTAELKICRVNRNSGSCLGGDEIFLLCDKVQKEDIEVYFTGPGWEARGSFSQADVHRQVAIVFRTPPYADPSLQAPVRVSMQLRRPSDRELSEPMEFQYLPDTD | 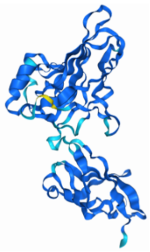 | Uniprot: Q04206. Human, aka NF-kb, nucleus, transcription factor, structured region |
| FGFR1c domain D2 (153-251) | MAPYWTNTEKMEKRLHAVPAANTVKFRCPAGGNPMPTMRWLKNGKEFKQEHRIGGYKVRNQHWSLIMESVVPSDKGNYTCVVENEYGSINHTYHLDVVER | 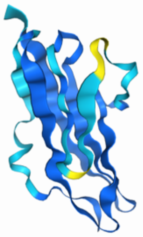 | Uniprot: P21802. Human, single-pass type 1 membrane protein, extracellular domain |
| TCIM | MKAKRSHQAVIMSTSLRVSPSIHGYHFDTASRKKAVGNIFENTDQESLERLFRNSGDKKAEERAKIIFAIDQDVEEKTRALMALKKRTKDKLFQFLKLRKYSIKVH | 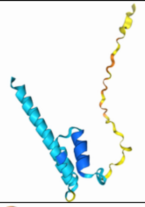 | Uniprot: Q9NR00. Human, nucleus/speckle/nucleolus, full length, disordered |
| TRIM21 (277-475) | MGLKKMLRTCAVHITLDPDTANPWLILSEDRRQVRLGDTQQSIPGNEERFDSYPMVLGAQHFHSGKHYWEVDVTGKEAWDLGVCRDSVRRKGHFLLSSKSGFWTIWLWNKQKYEAGTYPQTPLHLQVPPCQVGIFLDYEAGMVSFYNITDHGSLIYSFSECAFTGPLRPFFSPGFNDGGKNTAPLTLCPLNIGSQGSTDY | 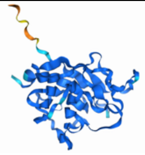 | Uniprot: P19474. Human, cytoplasm/nucleus, E3 ligase, structured domain, Fc binding region |
| PDE6D (1-150) | MSAKDERAREILRGFKLNWMNLRDAETGKILWQGTEDLSVPGVEHEARVPKKILKCKAVSRELNFSSTEQMEKFRLEQKVYFKGQCLEEWFFEFGFVIPNSTNTWQSLIEAAPESQMMPASVLTGNVIIETKFFDDDLLVSTSRVRLFYV | 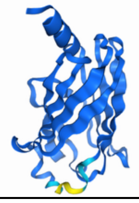 | Uniprot: O43924. Human, cytoplasm/membrane, binds/shuttles prenylated proteins including RAS family proteins; full length |

| | | | |
|---|---|---|---|
| RAF (52-131) | MSKTSNTIRVFLPNKQRTVVNVRNG<br>MSLHDCLMKALKVRGLQPECCAVFR<br>LLHEHKGKKARLDWNTDAASLIGEEL<br>QVDFL | 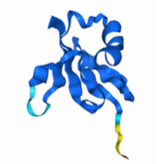 | Uniprot: P04049. Human, cytoplasm, proto oncogene, RAS binding domain |
| NRAS | MTEYKLVVVGAGGVGKSALTIQLIQN<br>HFVDEYDPTIEDSYRKQVVIDGETCL<br>LDILDTAGQEEYSAMRDQYMRTGEG<br>FLCVFAINNSKSFADINLYREQIKRVK<br>DSDDVPMVLVGNKCDLPTRTVDTKQ<br>AHELAKSYGIPFIETSAKTRQGVEDAF<br>YTLVREIRQYRMKKLNSSDDGTQGC<br>MGLPCVVM | 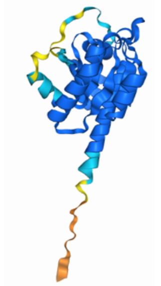 | Uniprot: P01111. Human, cytoplasm/cell membrane/Golgi apparatus membrane, proto oncogene, full length |
| Parkin | MIVFVRFNSSHGFPVEVDSDTSIFQL<br>KEVVAKRQGVPADQLRVIFAGKELRN<br>DWTVQNCDLDQQSIVHIVQRPWRKG<br>QEMNATGGDDPRNAAGGCEREPQS<br>LTRVDLSSSVLPGDSVGLAVILHTDS<br>RKDSPPAGSPAGRSIYNSFYVYCKG<br>PCQRVQPGKLRVQCSTCRQATLTLT<br>QGPSCWDDVLIPNRMSGECQSPHC<br>PGTSAEFFFKCGAHPTSDKETSVALH<br>LIATNSRNITCITCTDVRSPVLVFQCN<br>SRHVICLDCFHLYCVTRLNDRQFVHD<br>PQLGYSLPCVAGCPNSLIKELHHFRIL<br>GEEQYNRYQQYGAEECVLQMGGVL<br>CPRPGCGAGLLPEPDQRKVTCEGG<br>NGLGCGFAFCRECKEAYHEGECSAV<br>FEASGTTTQAYRVDERAAEQARWEA<br>ASKETIKKTTKPCPRCHVPVEKNGGC<br>MHMKCPQPQCRLEWCWNCGCEWN<br>RVCMGDHWFDV | 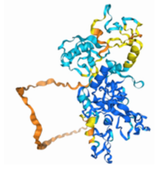 | Uniprot: O60260. Human, cytoplasm, E3 ligase, full length. |
| Mdm2 (1-188) | MCNTNMSVPTDGAVTTSQIPASEQE<br>TLVRPKPLLLKLLKSVGAQKDTYTMK<br>EVLFYLGQYIMTKRLYDEKQQHIVYC<br>SNDLLGDLFGVPSFSVKEHRKIYTMIY<br>RNLVVVNQQESSDSGTSVSENRCHL<br>EGGSDQKDLVQELQEEKPSSSHLVS<br>RPSTSSRRRAISETEENSDELSGERQ<br>RKRHKSDS | 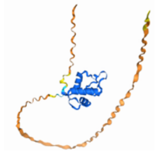 | Uniprot: Q00987. Human, cytoplasm/nucleus, E3 ligase and proto oncogene, region that binds p53. |
| HlgA (30-309) | MGHHHHHHAMENKIEDIGQGAEIIKR<br>TQDITSKRLAITQNIQFDFVKDKKYNK<br>DALVVKMQGFISSRTTYSDLKKYPYIK<br>RMIWPFQYNISLKTKDSNVDLINYLPK<br>NKIDSADVSQKLGYNIGGNFQSAPSI<br>GGSGSFNYSKTISYNQKNYVTEVES<br>QNSKGVKWGVKANSFVTPNGQVSA<br>YDQYLFAQDPTGPAARDYFVPDNQL<br>PPLIQSGFNPSFITTLSHERGKGDKS<br>EFEITYGRNMDATYAYVTRHRLAVDR<br>KHDAFKNRNVTVKYEVNWKTHEVKI<br>KSITPK | 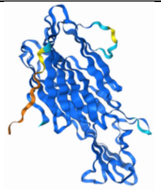 | Uniprot: P0A074. Staphylococcus aureus, secreted hemolytic toxin, folded domain (accidentally included His-tag). |

| | | | |
|---|---|---|---|
| Bcl2 | MAHAGRTGYDNREIVMKYIHYKLSQ<br>RGYEWDAGDVGAAPPGAAPAPGIFS<br>SQPGHTPHPAASRDPVARTSPLQTP<br>AAPGAAAGPALSPVPPVVHLTLRQA<br>GDDFSRRYRRDFAEMSSQLHLTPFT<br>ARGRFATVVEELFRDGVNWGRIVAF<br>FEFGGVMCVESVNREMSPLVDNIAL<br>WMTEYLNRHLHTWIQDNGGWDAFV<br>ELYGPSMRPLFDFSWLSLK | 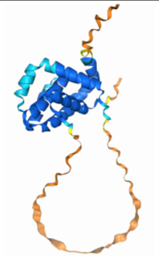 | Uniprot: P10415. Human, cytoplasm/mitochondrion outer membrane/nucleus membrane, suppresses apoptosis, full length. |
| HRAS | MTEYKLVVVGAGGVGKSALTIQLIQN<br>HFVDEYDPTIEDSYRKQVVIDGETCL<br>LDILDTAGQEEYSAMRDQYMRTGEG<br>FLCVFAINNTKSFEDIHQYREQIKRVK<br>DSDDVPMVLVGNKCDLAARTVESRQ<br>AQDLARSYGIPYIETSAKTRQGVEDA<br>FYTLVREIRQHKLRKLNPPDESGPGC<br>MSCKCVLS | 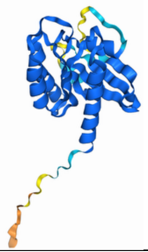 | Uniprot: P01112. Human, cytoplasm, GTPase, oncogene, full length. |
| KRAS wt | MTEYKLVVVGAGGVGKSALTIQLIQN<br>HFVDEYDPTIEDSYRKQVVIDGETCL<br>LDILDTAGQEEYSAMRDQYMRTGEG<br>FLCVFAINNTKSFEDIHHYREQIKRVK<br>DSEDVPMVLVGNKCDLPSRTVDTKQ<br>AQDLARSYGIPFIETSAKTRQGVDDA<br>FYTLVREIRKHKEKMSKDGKKKKKKS<br>KTKCVIM | 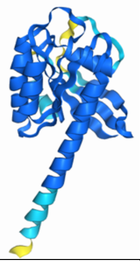 | Uniprot: P01116. Human, cytoplasm, GTPase, oncogene, full length isoform 4b. |
| G3BP (11-139) | MVGREFVRQYYTLLNQAPDMLHRFY<br>GKNSSYVHGGLDSNGKPADAVYGQ<br>KEIHRKVMSQNFTNCHTKIRHVDAHA<br>TLNDGVVVQVMGLLSNNNQALRRFM<br>QTFVLAPEGSVANKFYVHNDIFRYQD<br>EVFG | 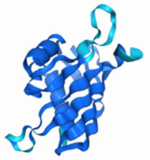 | Uniprot: Q13283. Human, cytoplasm, RAS GTPase activating protein, |
| NIX | MSSHLVEPPPPLHNNNNNCEENEQS<br>LPPPAGLNSSWVELPMNSSNGNDN<br>GNGKNGGLEHVPSSSSIHNGDMEKIL<br>LDAQHESGQSSSRGSSHCDSPSPQ<br>EDGQIMFDVEMHTSRDHSSQSEEEV<br>VEGEKEVEALKKSADWVSDWSSRPE<br>NIPPKEFHFRHPKRSVSLSMRKSGA<br>MKKGGIFSAEFLKVFIPSLFLSHVLAL<br>GLGIYIGKRLSTPSASTY | 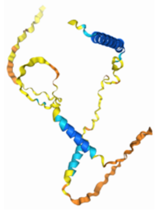 | Uniprot: O60238. Human, nucleus envelope/ER/mitochondrial outer membrane, single-pass membrane protein involved in apoptosis by suppressing BCl2, full length. |
| Sos1 (1136-1333) | MSSISLTKGTDEVPVPPPVPPRRRPE<br>SAPAESSPSKIMSKHLDSPPAIPPRQ<br>PTSKAYSPRYSISDRTSISDPPESPPL<br>LPPREPVRTPDVFSSSPLHLQPPPLG<br>KKSDHGNAFFPNSPSPFTPPPPQTP<br>SPHGTRRHLPSPPLTQEVDLHSIAGP<br>PVPPRQSTSQHIPKLPPKTYKREHTH<br>PSMHRDGPPLLENAHSS | 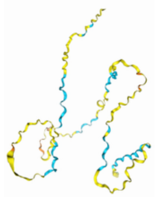 | Uniprot: Q07889. Human, cytoplasm, promotes RAS GTPase nucleotide exchange, disordered domain only. |
| PP2A-B' | MSSSSPPAGAASAAISASEKVDGFTR<br>KSVRKAQRQKRSQGSSQFRSQGSQ<br>AELHPLPQLKDATSNEQQELFCQKL<br>QQCCILFDFMDSVSDLKSKEIKRATL<br>NELVEYVSTNRGVIVESAYSDIVKMIS<br>ANIFRTLPPSDNPDFDPEEDEPTLEA | 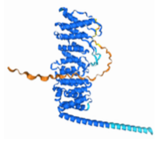 | Uniprot: Q15172. Human, cytoplasm, regulatory subunit of a phosphatase, full length. |

| | | | |
|---|---|---|---|
| | SWPHIQLVYEFFLRFLESPDFQPSIAK<br>RYIDQKFVQQLLELFDSEDPRERDFL<br>KTVLHRIYGKFLGLRAFIRKQINNIFLR<br>FIYETEHFNGVAELLEILGSIINGFALP<br>LKAEHKQFLMKVLIPMHTAKGLALFH<br>AQLAYCVVQFLEKDTTLTEPVIRGLLK<br>FWPKTCSQKEVMFLGEIEEILDVIEPT<br>QFKKIEEPLFKQISKCVSSSHFQVAE<br>RALYFWNNEYILSLIEENIDKILPIMFA<br>SLYKISKEHWNPTIVALVYNVLKTLME<br>MNGKLFDDLTSSYKAERQREKKKEL<br>EREELWKKLEELKLKKALEKQNSAYN<br>MHSILSNTSAE | | |
| mCherry | MVSKGEEDNMAIIKEFMRFKVHMEG<br>SVNGHEFEIEGEGEGRPYEGTQTSK<br>LKVTKGGPLPFAWDILSPQFMYGSKA<br>YVKHPADIPDYLKLSFPEGFKWERVM<br>NFEDGGVVTVTQDSSLQDGEFIYKVK<br>LRGTNFPSDGPVMQKKTMGWEASS<br>ERMYPEDGALKGEIKQRLKLKDGGH<br>YDAEVKTTYKAKKPVQLPGAYNVNIK<br>LDITSHNEDYTIVEQYERAEGRHSTG<br>GMDELYK | 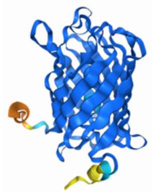 | Synthetic evolved protein from DsRed of Discosoma sea anemones. Red fluorescent protein full length. |
| Myc (DBD) | MNVKRRTHNVLERQRRNELKRSFFA<br>LRDQIPELENNEKAPKVVILKKATAYIL<br>SVQAEEQKLISEEDLLRKRREQLKHK<br>LEQL | 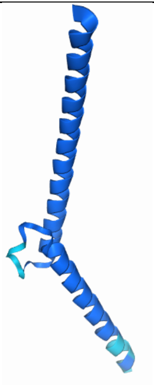 | Uniprot: P01106. Human, nucleus, proto oncogene transcription factor, DNA binding domain (368-451). |
| MBP (27-396) | MKIEEGKLVIWINGDKGYNGLAEVGK<br>KFEKDTGIKVTVEHPDKLEEKFPQVA<br>ATGDGPDIIFWAHDRFGGYAQSGLLA<br>EITPDKAFQDKLYPFTWDAVRYNGKL<br>IAYPIAVEALSLIYNKDLLPNPPKTWE<br>EIPALDKELKAKGKSALMFNLQEPYF<br>TWPLIAADGGYAFKYENGKYDIKDVG<br>VDNAGAKAGLTFLVDLIKNKHMNADT<br>DYSIAEAAFNKGETAMTINGPWAWS<br>NIDTSKVNYGVTVLPTFKGQPSKPFV<br>GVLSAGINAASPNKELAKEFLENYLLT<br>DEGLEAVNKDKPLGAVALKSYEEELA<br>KDPRIAATMENAQKGEIMPNIPQMSA<br>FWYAVRTAVINAASGRQTVDEALKD<br>AQTRITK | 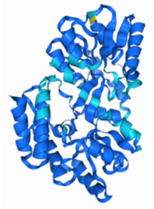 | Uniprot: P0AEX9. Escherichia coli, periplasm, maltose binding protein structured region. |

| | | | |
|---|---|---|---|
| Fibronectin (1080-1541) | MGVFTTLQPGSSIPPYNTEVTETTIVI<br>TWTPAPRIGFKLGVRPSQGGEAPRE<br>VTSDSGSIVVSGLTPGVEYVYTIQVLR<br>DGQERDAPIVNKVVTPLSPPTNLHLE<br>ANPDTGVLTVSWERSTTPDITGYRIT<br>TTPTNGQQGNSLEEVVHADQSSCTF<br>DNLSPGLEYNVSVYTVKDDKESVPIS<br>DTIIPEVPQLTDLSFVDITDSSIGLRWT<br>PLNSSTIIGYRITVVAAGEGIPIFEDFV<br>DSSVGYYTVTGLEPGIDYDISVITLING<br>GESAPTTLTQQTAVPPPTDLRFTNIG<br>PDTMRVTWAPPPSIDLTNFLVRYSPV<br>KNEEDVAELSISPSDNAVVLTNLLPG<br>TEYVVSVSSVYEQHESTPLRGRQKT<br>GLDSPTGIDFSDITANSFTVHWIAPRA<br>TITGYRIRHHPEHFSGRPREDRVPHS<br>RNSITLTNLTPGTEYVVSIVALNGREE<br>SPLLIGQQSTVSDV | 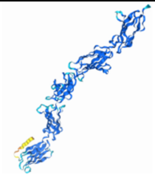 | Uniprot: P02751. Human, secreted, extracellular matrix, binds cell surfaces, various fibronectin type domains. |
| VHL (61-209) | MPVLRSVNSREPSQVIFCNRSPRVVL<br>PVWLNFDGEPQPYPTLPPGTGRRIH<br>SYRGHLWLFRDAGTHDGLLVNQTEL<br>FVPSLNVDGQPIFANITLPVYTLKERC<br>LQVVRSLVKPENYRRLDIVRSLYEDL<br>EDHPNVQKDLERLTQERIAHQ | 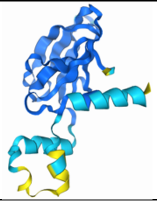 | Uniprot: P40337. Human, cytoplasm/ nucleus, E3 ligase, structured domain. |
| GABARAP | MKFVYKEEHPFEKRRSEGEKIRKKYP<br>DRVPVIVEKAPKARIGDLDKKKYLVPS<br>DLTVGQFYFLIRKRIHLRAEDALFFFV<br>NNVIPPTSATMGQLYQEHHEEDFFLY<br>IAYSDESVYGL | 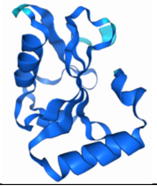 | Uniprot: O95166. Human, cytoplasm, involved in autophagy, full length. |
| GTF2I (1-106) | MAQVAMSTLPVEDEESSESRMVVTF<br>LMSALESMCKELAKSKAEVACIAVYE<br>TDVFVVGTERGRAFVNTRKDFQKDF<br>VKYCVEEEEKAAEMHKMKSTTQANR<br>MSVDA | 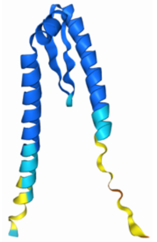 | Uniprot: P78347. Human, nucleus, transcription factor, DNA binding domain. |
| 14-3-3z Monomeric | MDKNELVQKAKQAEQAEDYDDMAA<br>CMKSVTEQGAELSNEERNLLSVAYK<br>NVVGARRSEWRVVSSIEQKTEGAEK<br>KQQMAREYREKIETELRDICNDVLSL<br>LEKFLIPNASQAESKVFYLKMKGDYY<br>RYLAEVAAGDDKKGIVDQSQQAYQE<br>AFEISKKEMQPTHPIRLGLALNFSVFY<br>YEILNSPEKACSLAKTAFDEAIAELDT<br>LSEESYKDSTLIMQLLRDNLTLWTSD<br>TQGDEAEAGEGGEN | 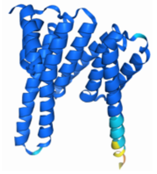 | Uniprot: P63104. Human, cytoplasm, interacts with phosphorylated proteins, full length. Modified to prevent dimerization (L12Q, R18E and S58E). |

| | | | |
|---|---|---|---|
| PCNA-AF | MVRTKADSVPGTYRKVVAARAPRKV LGSSTSATNSTSVSSRKAENKYAGG NPVCVRPTPKWQKGIGEFFRLSPKD SEKENQIPEEAGSSGLGKAKRKACPL QPDHTNDEKE | 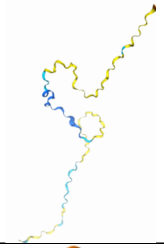 | Uniprot: Q15004. Human, nuclear, regulator of DNA replication, intrinsically disordered, full length. |
| TEV (2038-2274) | MGESLFKGPRDYNPISSTICHLTNES DGHTTSLYGIGFGPFIITNKHLFRRNN GTLLVQSLHGVFKVKNTTTLQQHLID GRDMIIIRMPKDFPPFPQKLKFREPQ REERICLVTTNFQTKSMSSMVSDTSC TFPSSDGIFWKHWIQTKDGQCGSPL VSTRDGFIVGIHSASNFTNTNNYFTS VPKNFMELLTNQEAQQWVSGWRLN ADSVLWGGHKVFMVKPEEPFQPVKE ATQLMN | 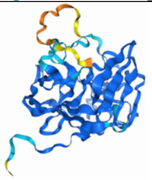 | Uniprot: Q88507. Tobacco etch virus, protease, proteolytic domain. |
| FnbpA (307-461) | MKYYTNLNGSIETFNKADNKFTHVAY VKPINGNKSESVSITGSLTQGSNVSG DSPIVKVYEYQGKETDLPKSVSVNLT DNSKFKDVTSDMQNKLTVQENGNYQ LNLEKLDKTYVIHYTGEYSKETDEVN FRTQVSAYPENSYRYYSYYNNHYTL TW | 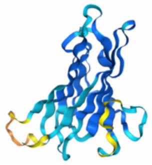 | Uniprot: P14738. Staphylococcus aureus, cell surface/secreted, fibronectin binding domain. |
| IBTK (1159-1339) | MLIDIISSKMIALTTKENNSGMNSMET VLFTPSKVPKPVNAWASSLHSVSSKS FRDFLLEEKKSVTSHSSGDHVKKVSF KGIENSQAPKIVRCSTHGTPGPEGNH ISDLPLLDSPNPWLSSSVTAPSMVAP VTFASIVEEELQQEAALIRSREKPLALI QIEEHAIQDLLVFYEAFGNPEEFVIVE RTPQGPLAVPMWNKHGC | 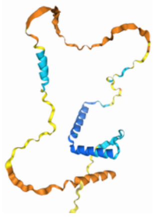 | Uniprot: Q9P2D0. Human, cytoplasm/nucleus, involved in B-cell development through BTK and NF-kb, disordered domain. |
| PIK3CB (3-115) | MFSFIMPPAMADILDIWAVDSQIASD GSIPVDFLLPTGIYIQLEVPREATISYIK QMLWKQVHNYPMFNLLMDIDSYMFA CVNQTAVYEELEDETRRLCDVRPFLP VLKLVTRS | 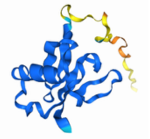 | Uniprot: P42338. Human, cytoplasm/nucleus, kinase, adaptor binding domain (ABD). |
| TNFR2 (33-205) | MAPEPGSTCRLREYYDQTAQMCCS KCSPGQHAKVFCTKTSDTVCDSCED STYTQLWNWVPECLSCGSRCSSDQ VETQACTREQNRICTCRPGWYCALS KQEGCRLCAPLRKCRPGFGVARPGT ETSDVVCKPCAPGTFSNTTSSTDICR PHQICNVVAIPGNASMDAVCTSTSP | 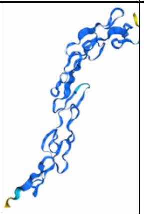 | Uniprot: P20333. Human, cell membrane, involved in regulating apoptosis, extracellular domain. |
| SH3BP5 | MLNHATQRVMEAEQTKTRSELVHKE TAARYNAAMGRMRQLEKKLKRAINK SKPYFELKAKYYVQLEQLKKTVDDLQ AKLTLAKGEYKMALKNLEMISDEIHER RRSSAMGPRGCGVGAEGSSTSVED L | 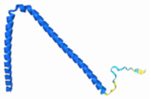 | Uniprot: C9JK30. Human, cytoplasm, guanine nucleotide exchange factor (GEF), full length. |

| | | | |
|---|---|---|---|
| OTUB1 (1-102) | MKPSWLSRTEFSKRLLCRTLWCQSG<br>WSSRSYTRSMLKMTTSINRRSRTST<br>KSTRTSARPGLTATVSIGLSDSPTWR<br>HCWMTARSCSGGEADLCRRPAGLL<br>Q | 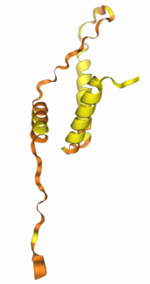 | Uniprot: Q96FW1. Human, cytoplasm, de-ubiquitinase, intrinsically disordered, full length. |
| PINK (107-353) | MGIPEFEEKQAESRRAVSACQEIQAI<br>FTQKSKPGPDPLDTRRLQGFRLEEYL<br>IGQSIGKGCSAAVYEATMPTLPQNLE<br>VTKSTGLLPGRGPGTSAPGEGQERA<br>PGAPAFPLAIKMMWNISAGSSSEAIL<br>NTMSQELVPASRVALAGEYGAVTYR<br>KSKRGPKQLAPHPNIIRVLRAFTSSVP<br>LLPGALVDYPDVLPSRLHPEGLGHG<br>RTLFLVMKNYPCTLRQYLCVNTPSPR<br>LAAMMLLQLLEGVDHL | 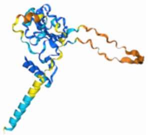 | Uniprot: Q9BXM7. Human, cytoplasm/mitochondrion membrane, kinase, kinase domain, partially disordered. |
| RB1 (1-103) | MPPKTPRKTAATAAAAAAEPPAPPPP<br>PPPEEDPEQDSGPEDLPLVRLEFEET<br>EEPDFTALCQKLKIPDHVRERAWLT<br>WEKVSSVDGVLVLRSPVFKPWLTMG<br>N | 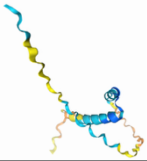 | Uniprot: P06400. Human, nucleus, regulates cell cycle, disordered domain (isoform 4). |
| FKBP (39-145) | MGVQVETISPGDGRTFPKRGQTCVV<br>HYTGMLEDGKKFDSSRDRNKPFKFM<br>LGKQEVIRGWEEGVAQMSVGQRAKL<br>TISPDYAYGATGHPGIIPPHATLVFDV<br>ELLKLE | 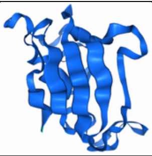 | Uniprot: Q0VDC6. Human, cytoplasm/sarcoplasmic reticulum membrane, cytoplasmic domain (excluding the membrane anchor). |
| HEWL (19-147) | MKVFGRCELAAAMKRHGLDNYRGY<br>SLGNWVCAAKFESNFNTQATNRNTD<br>GSTDYGILQINSRWWCNDGRTPGSR<br>NLCNIPCSALLSSDITASVNCAKKIVS<br>DGNGMNAWVAWRNRCKGTDVQAW<br>IRGCRL | 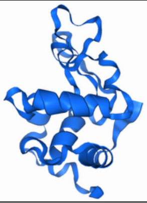 | Uniprot: P00698. Gallus gallus (chicken), secreted, lysozyme, structured domain (excluding signal domain). |
| HSPB1-str (90-171) | MHTADRWRVSLDVNHFAPDELTVKT<br>KDGVVEITGKHEERQDEHGYISRCFT<br>RKYTLPPGVDPTQVSSSLSPEGTLTV<br>EAPMPK | 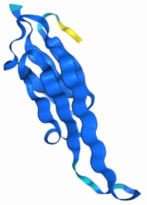 | Uniprot: P04792. Human, cytoplasm/nucleus, molecular chaperone, structured domain only. |
| LMO2 | MSSAIERKSLDPSEEPVDEVLQIPPSL<br>LTCGGCQQNIGDRYFLKAIDQYWHE<br>DCLSCDLCGCRLGEVGRRLYYKLGR<br>KLCRRDYLRLFGQDGLCASCDKRIRA<br>YEMTMRVKDKVYHLECFKCAACQKH<br>FCVGDRYLLINSDIVCEQDIYEWTKIN<br>GMI | 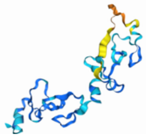 | Uniprot: P25791. Human, nucleus, transcription factor, mostly disordered, full length. |
| HIF1a (1-350) | MEGAGGANDKKKISSERRKEKSRDA<br>ARSRRSKESEVFYELAHQLPLPHNVS<br>SHLDKASVMRLTISYLRVRKLLDAGD<br>LDIEDDMKAQMNCFYLKALDGFVMV<br>LTDDGDMIYISDNVNKYMGLTQFELT | 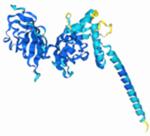 | Uniprot: Q16665. Human, cytoplasm/nucleus, transcriptional regulator, structured domain only. |

| | | | |
|---|---|---|---|
| | GHSVFDFTHPCDHEEMREMLTHRN<br>GLVKKGKEQNTQRSFFLRMKCTLTS<br>RGRTMNIKSATWKVLHCTGHIHVYDT<br>NSNQPQCGYKKPPMTCLVLICEPIPH<br>PSNIEIPLDSKTFLSRHSLDMKFSYCD<br>ERITELMGYEPEELLGRSIYEYYHALD<br>SDHLTKTHHDMFTKGQVTTGQYRML<br>AKRGGYVWVETQATVIYNTKNSQPQ<br>CIVCVNYVVSGIIQHDL | | |
| MXD1 | MAAAVRMNIQMLLEAADYLERRERE<br>AEHGYASMLPYNNKDRDALKRRNKS<br>KKNNSSSRSTHNEMEKNRRAHLRLC<br>LEKLKGLVPLGPESSRHTTLSLLTKAK<br>LHIKKLEDCDRKAVHQIDQLQREQRH<br>LKRQLEKLGIERIRMDSIGSTVSSERS<br>DSDREEIDVDVESTDYLTGDLDWSS<br>SSVSDSDERGSMQSLGSDEGYSSTS<br>IKRIKLQDSHKACLGL | 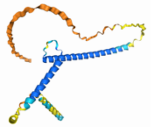 | Uniprot: Q05195. Human, nucleus, transcriptional repressor, intrinsically disordered, full length. |
| OmpA (ec, 84-224) | MADKQEAELRRQMEGTGVEVQRQG<br>DDIKLIMPGNITFATDSANIAPSFYAPL<br>NNLANSFKQYNQNTIEIVGYTDSTGS<br>RQHNMDLSQRRAQSVAGYLTAQGV<br>DGTRLSTRGMGPDQPIASNSTADGR<br>AQNRRVEVNLRPV | 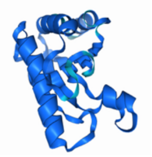 | Uniprot: Q9I5A7. Pseudomonas aeruginosa, outer membrane protein, extracellular domain. |
| CD3z | MKWKALFTAAILQAQLPITEAQSFGLL<br>DPKLCYLLDGILFIYGVILTALFLRVKF<br>SRSADAPAYQQGQNQLYNELNLGRR<br>EEYDVLDKRRGRDPEMGGKPQRRK<br>NPQEGLYNELQKDKMAEAYSEIGMK<br>GERRRGKGHDGLYQGLSTATKDTYD<br>ALHMQALPPR | 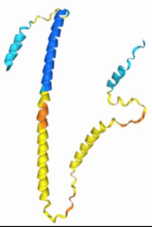 | Uniprot: P20963. Human, cell membrane, T-cell surface receptor, single pass-transmembrane protein, fully disordered, full length. |
| PIK3CA (1-105) | MPPRPSSGELWGIHLMPPRILVECLL<br>PNGMIVTLECLREATLITIKHELFKEAR<br>KYPLHQLLQDESSYIFVSVTQEAERE<br>EFFDETRRLCDLRLFQPFLKVIEPV | 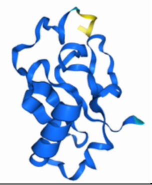 | Uniprot: P42336. Human, cytoplasm, kinase, adaptor binding domain (ABD). |
| UEV (1-145) | MAVSESQLKKMVSKYKYRDLTVRET<br>VNVITLYKDLKPVLDSYVFNDGSSRE<br>LMNLTGTIPVPYRGNTYNIPICLWLLD<br>TYPHNPPICFVKPTSSMTIKTGKHVD<br>ANGKIYLPYLHEWKHPQSDLLGLIQV<br>MIVVFGDEPPVFSRP | 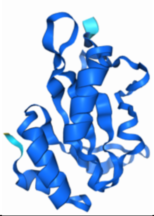 | Uniprot: Q99816. Human, cytoplasm/nucleus, binds ubiquitinated proteins, UEV domain. |
| FRB (2019-2114) | MVAILWHEMWHEGLEEASRLYFGER<br>NVKGMFEVLEPLHAMMERGPQTLKE<br>TSFNQAYGRDLMEAQEWCRKYMKS<br>GNVKDLTQAWDLYYHVFRRISKQ | 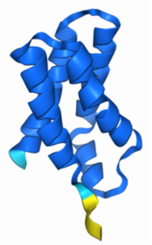 | Uniprot: P42345. Human, cytoplasm, kinase (mTOR), FKPB rapamycin associated domain (FRB. |

| | | | |
|---|---|---|---|
| PD-1 (25-160) | MLDSPDRPWNPPTFSPALLVVTEGD<br>NATFTCSFSNTSESFVLNWYRMSPS<br>NQTDKLAAFPEDRSQPGQDSRFRVT<br>QLPNGRDFHMSVVRARRNDSGTYLC<br>GAISLAPKAQIKESLRAELRVTERRAE<br>VPTAHPSPSP | 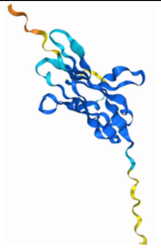 | Uniprot: Q15116. Human, cell membrane, inhibitor of T-cell activation, extracellular domain. |
| ALKBH5 | MAAASGYTDLREKLKSMTSRDNYKA<br>GSREAAAAAAAAVAAAAAAAAAAEP<br>YPVSGAKRKYQEDSDPERSDYEEQQ<br>LQKEEEARKVKSGIRQMRLFSQDEC<br>AKIEARIDEVVSRAEKGLYNEHTVDR<br>APLRNKYFFGEGYTYGAQLQKRGPG<br>QERLYPPGDVDEIPEWVHQLVIQKLV<br>EHRVIPEGFVNSAVINDYQPGGCIVS<br>HVDPIHIFERPIVSVSFFSDSALCFGC<br>KFQFKPIRVSEPVLSLPVRRGSVTVL<br>SGYAADEITHCIRPQDIKERRAVIILRK<br>TRLDAPRLETKSLSSSVLPPSYASDR<br>LSGNNRDPALKPKRSHRKADPDAAH<br>RPRILEMDKEENRRSVLLPTHRRRGS<br>FSSENYWRKSYESSEDCSEAAGSPA<br>RKVKMRRHSRGSRA | 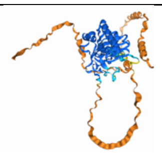 | Uniprot: Q6P6C2. Human, nucleus, RNA demethylase, partially disordered, full length. |
| HIF1b (1-474) | MAATTANPEMTSDVPSLGPAIASGNS<br>GPGIQGGGAIVQRAIKRRPGLDFDDD<br>GEGNSKFLRCDDDQMSNDKERFAR<br>SDDEQSSADKERLARENHSEIERRR<br>RNKMTAYITELSDMVPTCSALARKPD<br>KLTILRMAVSHMKSLRGTGNTSTDGS<br>YKPSFLTDQELKHLILEAADGFLFIVS<br>CETGRVVYVSDSVTPVLNQPQSEWF<br>GSTLYDQVHPDDVDKLREQLSTSEN<br>ALTGRILDLKTGTVKKEGQQSSMRM<br>CMGSRRSFICRMRCGSSSVDPVSVN<br>RLSFVRNRCRNGLGSVKDGEPHFVV<br>VHCTGYIKAWPPAGVSLPDDDPEAG<br>QGSKFCLVAIGRLQVTSSPNCTDMS<br>NVCQPTEFISRHNIEGIFTFVDHRCVA<br>TVGYQPQELLGKNIVEFCHPEDQQLL<br>RDSFQQVVKLKGQVLSVMFRFRSKN<br>QEWLWMRTSSFTFQNPYSDEIEYIIC<br>TNTNVKNSSQEPRPT | 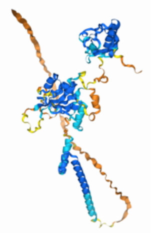 | Uniprot: P27540. Human, nucleus, heterodimerizes with ARNT to induce transcription, all structured regions, but only the N-terminal disordered domain. |
| HPV-pE7 | MHGDTPTLHEYMLDLQPETTDLYCY<br>EQLNDSSEEEDEIDGPAGQAEPDRA<br>HYNIVTFCCKCDSTLRLCVQSTHVDI<br>RTLEDLLMGTLGIVCPICSQKP | 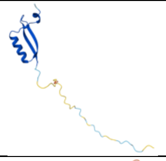 | Uniprot: P03129. Human papillomavirus, infects host cytoplasm and nucleus |
| MST2 (313-437) | MENSDEDELDSHTMVKTSVESVGTM<br>RATSTMSEGAQTMIEHNSTMLESDL<br>GTMVINSEDEEEEDGTMKRNATSPQ<br>VQRPSFMDYFDKQDFKNKSHENCN<br>QNMHEPFPMSKNVFPDNWKVPQDG<br>DF | 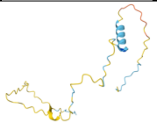 | Uniprot: Q13188. Cytoplasm, nucleus, cytoskeleton, Ser/Thr protein kinase, stress-activated and promotes apoptosis |

| | | | |
|---|---|---|---|
| p53 (long) (1-293) | MEEPQSDPSVEPPLSQETFSDLWKLLPENNVLSPLPSQAMDDLMLSPDDIEQWFTEDPGPDEAPRMPEAAPPVAPAPAAPTPAAPAPAPSWPLSSSVPSQKTYQGSYGFRLGFLHSGTAKSVTCTYSPALNKMFCQLAKTCPVQLWVDSTPPPGTRVRAMAIYKQSQHMTEVVRRCPHHERCSDSDGLAPPQHLIRVEGNLRVEYLDDRNTFRHSVVVPYEPPEVGSDCTTIHYNYMCNSSCMGGMNRRPILTIITLEDSSGNLLGRNSFEVRVCACPGRDRRTEEENLRKKG | 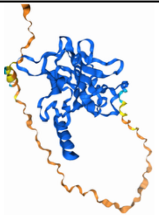 | Uniprot: P04637. Human, nucleus, transcription factor, N-terminal disordered domain and structured domain. |
| PD-L1 (18-239) | MAFTVTVPKDLYVVEYGSNMTIECKFPVEKQLDLAALIVYWEMEDKNIIQFVHGEEDLKVQHSSYRQRARLLKDQLSLGNAALQITDVKLQDAGVYRCMISYGGADYKRITVKVNAPYNKINQRILVVDPVTSEHELTCQAEGYPKAEVIWTSSDHQVLSGKTTTTNSKREEKLFNVTSTLRINTTTNEIFYCTFRRLDPEENHTAELVIPELPLAHPPNERT | 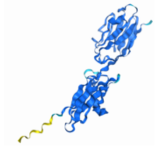 | Uniprot: Q9NZQ7. Human, cell membrane, important in T-cell activation, extracellular domain only. |
| BAX (short) (58-72) | MKKLSECLKRIGDELDS | 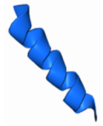 | Uniprot: Q07812. Human, cytoplasm, mitochondrion membrane, involved in mitochondrial apoptosis, BH3 domain. |
| p53 (short) (1-61) | MEEPQSDPSVEPPLSQETFSDLWKLLPENNVLSPLPSQAMDDLMLSPDDIEQWFTEDPGPD | 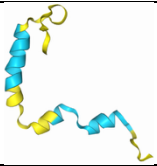 | Uniprot: P04637. Human, nucleus, transcription factor, N-terminal disordered activation domain. |
| CRBN (47-435) | MINFDTSLPTSHTYLGADMEEFHGRTLHDDDSCQVIPVLPQVMMILIPGQTLPLQLFHPQEVSMVRNLIQKDRTFAVLAYSNVQEREAQFGTTAEIYAYREEQDFGIEIVKVKAIGRQRFKVLELRTQSDGIQQAKVQILPECVLPSTMSAVQLESLNKCQIFPSKPVSREDQCSYKWWQKYQKRKFHCANLTSWPRWLYSLYDAETLMDRIKKQLREWDENLKDDSLPSNPIDFSYRVAACLPIDDVLRIQLLKIGSAIQRLRCELDIMNKCTSLCCKQCQETEITTKNEIFSLSLCGPMAAYVNPHGYVHETLTVYKACNLNLIGRPSTEHSWFPGYAWTVAQCKICASHIGWKFTATKKDMSPQKFWGLTRSALLPTIPDTEDEISP | 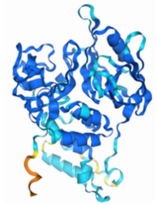 | Uniprot: Q96SW2. Human, cytoplasm/nucleus, E3 ligase, structured domain only. |
| PPP1R11 | MAEAGAGLSETVTETTVTVTTEPENRSLTIKLRKRKPEKKVEWTSDTVDNEHMGRRSSKCCCIYEKPRAFGESSTESDEEEEEGCGHTHCVRGHRKGRRRATLGPTPTTPPQPPDPSQPPPGPMQH | 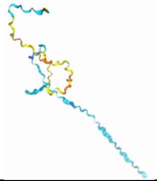 | Uniprot: O60927. Human, cytoplasm, nucleus, E3 ligase, intrinsically disordered, full-length. |

| | | | |
|---|---|---|---|
| PMP22 | MLLLLLSIIVLHVAVLVLLFVSTIVSQWIVGNGHATDLWQNCSTSSSGNVHHCFSSSPNEWLQSVQATMILSIIFSILSLFLFFCQLFTLTKGGRFYITGIFQILAGLCVMSAAAIYTVRHPEWHLNSDYSYGFAYILAWVAFPLALLSGVIYVILRKRE | 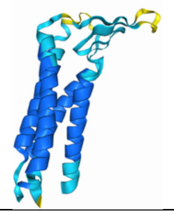 | Uniprot: Q01453. Human, cell membrane, involved in myelination, full length. |
| BAX (42-153) | MIAAVDTDSPREVFFRVAADMFSDGNFNWGRVVALFYFASKLVLKALCTKVPELIRTIMGWTLDFLRERLLGWIQDQGGWVRLLKPPHPHHRALTTAPAPPSLPPATPLGPW | 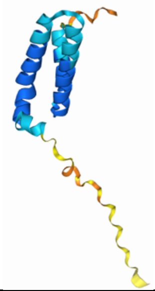 | Uniprot: Q07812. Human, cytoplasm, mitochondrion membrane, involved in mitochondrial apoptosis, isoform gamma. |
| Keap1 (321-611) | MAPKVGRLIYTAGGYFRQSLSYLEAYNPSDGTWLRLADLQVPRSGLAGCVVGGLLYAVGGRNNSPDGNTDSSALDCYNPMTNQWSPCAPMSVPRNRIGVGVIDGHIYAVGGSHGCIHHNSVERYEPERDEWHLVAPMLTRRIGVGVAVLNRLLYAVGGFDGTNRLNSAECYYPERNEWRMITAMNTIRSGAGVCVLHNCIYAAGGYDGQDQLNSVERYDVETETWTFVAPMKHRRSALGITVHQGRIYVLGGYDGHTFLDSVECYDPDTDTWSEVTRMTSGRSGVGVAVTME | 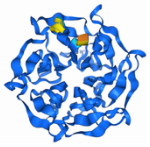 | Uniprot: Q14145. Human, cytoplasm/nucleus, E3 ligase, Kelch domains. |
| HSPB1 | MTERRVPFSLLRGPSWDPFRDWYPHSRLFDQAFGLPRLPEEWSQWLGGSSWPGYVRPLPPAAIESPAVAAPAYSRALSRQLSSGVSEIRHTADRWRVSLDVNHFAPDELTVKTKDGVVEITGKHEERQDEHGYISRCFTRKYTLPPGVDPTQVSSSLSPEGTLTVEAPMPKLATQSNEITIPVTFESRAQLGGPEAAKSDETAAK | 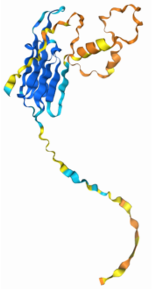 | Uniprot: P04792. Human, cytoplasm/nucleus, molecular chaperone, mostly disordered, full length. |
| HMGB1 | MGKGDPKKPRGKMSSYAFFVQTCREEHKKKHPDASVNFSEFSKKCSERWKTMSAKEKGKFEDMAKADKARYEREMKTYIPPKGETKKKFKDPNAPKRPPSAFFLFCSEYRPKIKGEHPGLSIGDVAKKLGEMWNNTAADDKQPYEKKAAKLKEKYEKDIAAYRAKGKPDAAKKGVVKAEKSKKKKEEEEDEEDEEDEEEEEDEEDEDEEEEDDDDE | 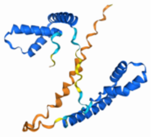 | Uniprot: P09429. Human, nucleus, transcription factor, partially disordered, full length. |
| hSUMO1 | MSDQEAKPSTEDLGDKKEGEYIKLKVIGQDSSEIHFKVKMTTHLKKLKESYCQRQGVPMNSLRFLFEGQRIADNHTPKELGMEEEDVIEVYQEQTGGHSTV | 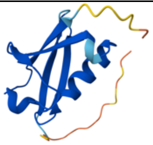 | Uniprot: P63165. Human, cytoplasm, nucleus, ubiquitin-like protein covalently attached to other proteins as a post-translational modification, full length |

| | | | |
|---|---|---|---|
| CD3d | MEHSTFLSGLVLATLLSQVSPFKIPIE<br>ELEDRVFVNCNTSITWVEGTVGTLLS<br>DITRLDLGKRILDPRGIYRCNGTDIYK<br>DKESTVQVHYRMCQSCVELDPATVA<br>GIIVTDVIATLLLALGVFCFAGHETGRL<br>SGAADTQALLRNDQVYQPLRDRDDA<br>QYSHLGGNWARNK | 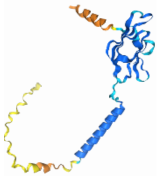 | Uniprot: P04234. Human, cell membrane, T-cell surface receptor, single pass-transmembrane protein, fully disordered, full length. |
| TRIM63 (160-353) | MSVFQGQKTELNNCISMLVAGNDRV<br>QTIITQLEDSRRVTKENSHQVKEELS<br>QKFDTLYAILDEKKSELLQRITQEQEK<br>KLSFIEALIQQYQEQLDKSTKLVETAI<br>QSLDEPGGATFLLTAKQLIKSIVEASK<br>GCQLGKTEQGFENMDFFTLDLEHIAD<br>ALRAIDFGTDEEEEEFIEEEDQEEEES<br>TEGKEEGHQ | 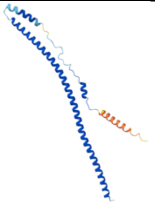 | Uniprot: Q969Q1. Human, cytoplasm, E3 ligase, truncation (160-353, excludes first RING-type Zn finger) |
| RHEB | MPQSKSRKIAILGYRSVGKSSLTIQFV<br>EGQFVDSYDPTIENTFTKLITVNGQE<br>YHLQLVDTAGQDEYSIFPQTYSIDING<br>YILVYSVTSIKSFEVIKVIHGKLLDMVG<br>KVQIPIMLVGNKKDLHMERVISYEEG<br>KALAESWNAAFLESSAKENQTAVDV<br>FRRIILEAEKMDGAASQGKSSCSVM | 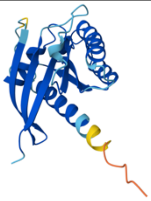 | Uniprot: Q15382. Human, endomembrane system, E3 ligase, full length |
| RNF2 (113-226, DZn finger) | MIYPSRDEYEAHQERVLARINKHNNQ<br>QALSHSIEEGLKIQAMNRLQRGKKQQ<br>IENGSGAEDNGDSSHCSNASTHSNQ<br>EAGPSNKRTKTSDDSGLELDNNNAA<br>MAIDPVMDGASEI | 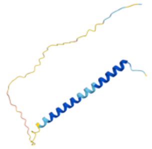 | Uniprot: X6RFN3. Human, nuclear body and nucleoplasm, E3 ligase, truncation (113-226, excludes first RING-type domain) |
| RNF8 (DZn finger) | MGEPGFFVTGDRAGGRSWCLRRVG<br>MSAGWLLLEDGCEVTVGRGFGVTY<br>QLVSKICPLMISRNHCVLKQNPEGQ<br>WTIMDNKSLNGVWLNRARLEPLRVY<br>SIHQGDYIQLGVPLENKENAEYEYEV<br>TEEDWETIYPCLSPKNDQMIEKNKEL<br>RTKRKFSLDELAGPGAEGPSNLKSKI<br>NKVSCESGQPVKSQGKGEVASTPSD<br>NLDPKLTALEPSKTTGAPIYPGFPKVT<br>EVHHEQKASNSSASQRSLQMFKVTM<br>SRILRLKIQMQEKHEAVMNVKKQTQK<br>GNSKKVVQMEQELQDLQSQLCAEQ<br>AQQQARVEQLEKTFQEEEQHLQGLE<br>IAQGEKDLKQQLAQALQEHWALMEE<br>LNRSKKDFEAIIQAKNKELEQTKEEKE<br>KMQAQKEEVLSHMNDVLENELQKDI<br>KSKTYSLVLDNCINKMVNNLSSEVKE<br>RRIVLIRERKAKRLF | 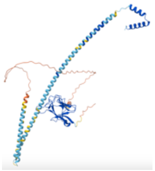 | Uniprot: O76064. Human, cytoplasm, nucleus, E3 ligase, excludes first RING-type domain |
| ASB1 (1-147) | MAEGGSPDGRAGPGSAGRNLKEWL<br>REQFCDHPLEHCEDTRLHDAAYVGD<br>LQTLRSLLQEESYRSRINEKSVWCCG<br>WLPCTPLRIAATAGHGSCVDFLIRKG<br>AEVDLVDVKGQTALYVAVVNGHLES<br>TQILLEAGADPNGSRHHRSTP | 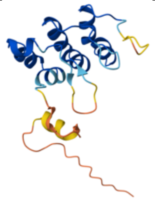 | Uniprot: Q9Y576. Human, nucleoplasm, E3 ligase, truncation (1-147) |

| | | | |
|---|---|---|---|
| BIRC7 | MGPKDSAKCLHRGPQPSHWAAGDG<br>PTQERCGPRSLGSPVLGLDTCRAWD<br>HVDGQILGQLRPLTEEEEEEGAGATL<br>SRGPAFPGMGSEELRLASFYDWPLT<br>AEVPPELLAAAGFFHTGHQDKVRCF<br>FCYGGLQSWKRGDDPWTEHAKWFP<br>SCQFLLRSKGRDFVHSVQETHSQLL<br>GSWDPWEEPEDAAPVAPSVPASGY<br>PELPTPRREVQSESAQEPGGVSPAE<br>AQRAWWVLEPPGARDVEAQLRRLQ<br>EERTCKVCLDRAVSIVFVPCGHLVCA<br>ECAPGLQLCPICRAPVRSRVRTFLS | 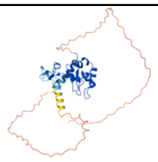 | Uniprot: Q96CA5. Human, cytoplasm, nucleus, E3 ligase, full length |
| SIX1-DBD (1-167) | MSMLPSFGFTQEQVACVCEVLQQG<br>GNLERLGRFLWSLPACDHLHKNESV<br>LKAKAVVAFHRGNFRELYKILESHQF<br>SPHNHPKLQQLWLKAHYVEAEKLRG<br>RPLGAVGKYRVRRKFPLPRTIWDGE<br>ETSYCFKEKSRGVLREWYAHNPYPS<br>PREKRELAEATGLTTTQ | 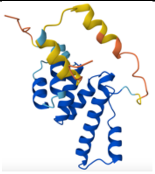 | Uniprot: Q3C2H5 (entry for chicken, but sequence identities are same). Human, cytoplasm, nucleus, transcription factor, DNA binding domain only |
| WWP2 | MASASSSRAGVALPFEKSQLTLKVVS<br>AKPKVHNRQPRINSYVEVAVDGLPSE<br>TKKTGKRIGSSELLWNEIIILNVTAQS<br>HLDLKVWSCHTLRNELLGTASVNLSN<br>VLKNNGGKMENMQLTLNLQTENKGS<br>VVSGGELTIFLDGPTVDLGNVPNGSA<br>LTDGSQLPSRDSSGTAVAPENRHQP<br>PSTNCFGGRSRTHRHSGASARTTPA<br>TGEQSPGARSRHRQPVKNSGHSGL<br>ANGTVNDEPTTATDPEEPSVVGVTS<br>PPAAPLSVTPNPNTTSLPAPATPAEG<br>EEPSTSGTQQLPAAAQAPDALPAGW<br>EQRELPNGRVYYVDHNTKTTTWERP<br>LPPGWEKRTDPRGRFYYVDHNTRTT<br>TWQRPTAEYVRNYEQWQSQRNQLQ<br>GAMQHFSQRFLYQSSSASTDHDPLG<br>PLPPGWEKRQDNGRVYYVNHNTRT<br>TQWEDPRTQGMIQEPALPPGWEMK<br>YTSEGVRYFVDHNTRTTTFKDPRPG<br>FESGTKQGSPGAYDRSFRWKYHQF<br>RFLCHSNALPSHVKISVSRQTLFEDS<br>FQQIMNMKPYDLRRRLYIIMRGEEGL<br>DYGGIAREWFFLLSHEVLNPMYCLFE<br>YAGKNNYCLQINPASSINPDHLTYFR<br>FIGRFIAMALYHGKFIDTGFTLPFYKR<br>MLNKRPTLKDLESIDPEFYNSIVWIKE<br>NNLEECGLELYFIQDMEILGKVTTHEL<br>KEGGESIRVTEENKEEYIMLLTDWRF<br>TRGVEEQTKAFLDGFNEVAPLEWLR<br>YFDEKELELMLCGMQEIDMSDWQKS<br>TIYRHYTKNSKQIQWFWQVVKEMDN<br>EKRIRLLQFVTGTCRLPVGGFAELIGS<br>NGPQKFCIDKVGKETWLPRSHTCFN<br>RLDLPPYKSYEQLREKLLYAIEETEGF<br>GQE | 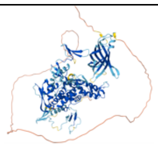 | Uniprot: O00308. Human, nucleus, E3 ligase, full length |

| | | | |
|---|---|---|---|
| SIX1 | MSMLPSFGFTQEQVACVCEVLQQG<br>GNLERLGRFLWSLPACDHLHKNESV<br>LKAKAVVAFHRGNFRELYKILESHQF<br>SPHNHPKLQQLWLKAHYVEAEKLRG<br>RPLGAVGKYRVRRKFPLPRTIWDGE<br>ETSYCFKEKSRGVLREWYAHNPYPS<br>PREKRELAEATGLTTTQVSNWFKNR<br>RQRDRAAEAKERENTENNNSSSNKQ<br>NQLSPLEGGKPLMSSSEEEFSPPQS<br>PDQNSVLLLQGNMGHARSSNYSLPG<br>LTASQPSHGLQTHQHLQDSLLGPL<br>TSSLVDLGS | 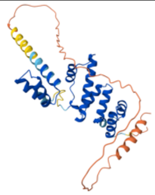 | Uniprot: Q3C2H5 (entry for chicken, but sequence identities are same). Human, cytoplasm, nucleus, transcription factor, full length |
| ERG-IDR (1-112) | MASTIKEALSVVSEDQSLFECAYGTP<br>HLAKTEMTASSSSDYGQTSKMSPRV<br>PQQDWLSQPPARVTIKMECNPSQVN<br>GSRNSPDECSVAKGGKMVGSPDTV<br>GMNYGSYMEEKH | 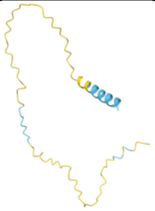 | Uniprot: P11308. Human, cytoplasm, nucleus, transcription factor, disordered domain only |
| HAND2-DBD (99-151) | MKRRGTANRKERRRTQSINSAFAEL<br>RECIPNVPADTKLSKIKTLRLATSYIAY<br>L | 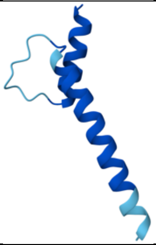 | Uniprot: P61296. Human, nucleus, transcription factor, DNA binding domain only |
| HES1 | MPADIMEKNSSSPVAATPASVNTTPD<br>KPKTASEHRKSSKPIMEKRRRARINE<br>SLSQLKTLILDALKKDSSRHSKLEKAD<br>ILEMTVKHLRNLQRAQMTAALSTDPS<br>VLGKYRAGFSECMNEVTRFLSTCEG<br>VNTEVRTRLLGHLANCMTQINAMTYP<br>GQPHPALQAPPPPPPGPGGPQHAPF<br>APPPPLVPIPGGAAPPPGGAPCKLGS<br>QAGEAAKVFGGFQVVPAPDGQFAFL<br>IPNGAFAHSGPVIPVYTSNSGTSVGP<br>NAVSPSSGPSLTADSMWRPWRN | 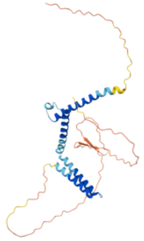 | Uniprot: Q14469. Human, nucleus, transcription factor, full length |
| HOXA9 | MATTGALGNYYVDSFLLGADAADEL<br>SVGRYAPGTLGQPPRQAATLAEHPD<br>FSPCSFQSKATVFGASWNPVHAAGA<br>NAVPAAVYHHHHHHPYVHPQAPVAA<br>AAPDGRYMRSWLEPTPGALSFAGLP<br>SSRPYGIKPEPLSARRGDCPTLDTHT<br>LSLTDYACGSPPVDREKQPSEGAFS<br>ENNAENESGGDKPPIDPNNPAANWL<br>HARSTRKKRCPYTKHQTLELEKEFLF<br>NMYLTRDRRYEVARLLNLTERQVKIW<br>FQNRRMKMKKINKDRAKDE | 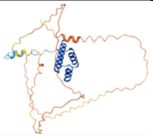 | Uniprot: P31269. Human, nucleus, cytoplasm, nucleus, transcription factor, full length |

| | | | |
|---|---|---|---|
| GATA2-DBD (295-373 | MCVNCGATATPLWRRDGTGHYLCN<br>ACGLYHKMNGQNRPLIKPKRRLSAA<br>RRAGTCCANCQTTTTTLWRRNANG<br>DPVCNAC | 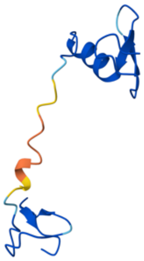 | Uniprot: P23769. Human, nucleus, transcription factor, DNA binding domain only |
| POU5F1-DBD (138-212) | DIKALQKELEQFAKLLKQKRITLGYTQ<br>ADVGLTLGVLFGKVFSQTTICRFEAL<br>QLSFKNMCKLRPLLQKWVEEAD | 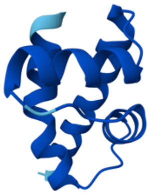 | Uniprot: D5K9R7. Human, nucleus, transcription factor, DNA binding domain only |
| THAP1 | MVQSCSAYGCKNRYDKDKPVSFHKF<br>PLTRPSLCKEWEAAVRRKNFKPTKY<br>SSICSEHFTPDCFKRECNNKLLKENA<br>VPTIFLCTEPHDKKEDLLEPQEQLPP<br>PPLPPPVSQVDAAIGLLMPPLQTPVN<br>LSVFCDHNYTVEDTMHQRKRIHQLE<br>QQVEKLRKKLKTAQQRCRRQERQLE<br>KLKEVVHFQKEKDDVSERGYVILPND<br>YFEIVEVPA | 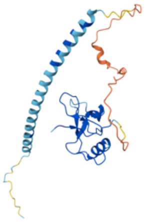 | Uniprot: Q9NVV9. Human, nucleus, transcription factor, full length |
| TBX20-DBD (109-288 | MLWDKFHELGTEMIITKSGRRMFPTI<br>RVSFSGVDPEAKYIVLMDIVPVDNKR<br>YRYAYHRSSWLVAGKADPPLPARLY<br>VHPDSPFTGEQLLKQMVSFEKVKLT<br>NNELDQHGHIILNSMHKYQPRVHIIKK<br>KDHTASLLNLKSEEFRTFIFPETVFTA<br>VTAYQNQLITKLKIDSNPFAKGFRD | 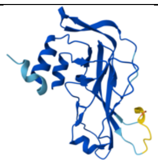 | Uniprot: Q9UMR3. Human, nucleus, transcription factor, DNA binding domain only |
| OLIG2-DBD (108-165) | MQLRLKINSRERKRMHDLNIAMDGLR<br>EVMPYAHGPSVRKLSKIATLLLARNYI<br>LML | 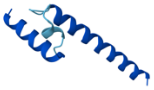 | Uniprot: Q13516. Human, cytoplasm, nucleus, transcription factor, DNA binding domain only |
| ZBED2 | MMRREDEEEEGTMMKAKGDLEMKE<br>EEEISETGELVGPFVSAMPTPMPHNK<br>GTRFSEAWEYFHLAPARAGHHPNQY<br>ATCRLCGRQVSRGPGVNVGTTALW<br>KHLKSMHREELEKSGHGQAGQRQD<br>PRPHGPQLPTGIEGNWGRLLEQVGT<br>MALWASQREKEVLRRERAVEWRER<br>AVEKRERALEEVERAILEMKWKVRAE<br>KEACQREKELPAAVHPFHFV | 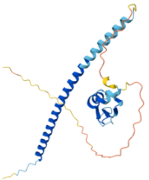 | Uniprot: Q9BTP6. Human, nucleus, transcription factor, full length |
| GAPBA-DBD (168-251) | AALEGYRKEQERLGIPYDPIQWSTDQ<br>VLHWVVWVMKEFSMTDIDLTTLNISG<br>RELCSLNQEDFFQRVPRGEILWSHLE<br>LLRKYV | 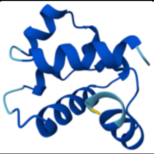 | Uniprot: A8IE48. Human, nucleus, transcription factor, DNA binding domain only |
| GATA4 | MYQSLAMAANHGPPPGAYEAGGPG<br>AFMHGAGAASSPVYVPTPRVPSSVL<br>GLSYLQGGGAGSASGGASGGSSGG<br>AASGAGPGTQQGSPGWSQAGADGA<br>AYTPPPVSPRFSFPGTTGSLAAAAAA<br>AAAREAAAYSSGGGAAGAGLAGRE | 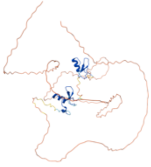 | Uniprot: A8IE48. Human, nucleus, transcription factor, full length |

| | |
|---|---|
| | QYGRAGFAGSYSSPYPAYMADVGA<br>SWAAAAAAASAGPFDSPVLHSLPGRA<br>NPAARHPNLDMFDDFSEGRECVNC<br>GAMSTPLWRRDGTGHYLCNACGLY<br>HKMNGINRPLIKPQRRLSASRRVGLS<br>CANCQTTTTTLWRRNAEGEPVCNAC<br>GLYMKLHGVPRPLAMRKEGIQTRKR<br>KPKNLNKSKTPAAPSGSESLPPASGA<br>SSNSSNATTSSSEEMRPIKTEPGLSS<br>HYGHSSSVSQTFSVSAMSGHGPSIH<br>PVLSALKLSPQGYASPVSQSPQTSSK<br>QDSWNSLVLADSHGDIITA |

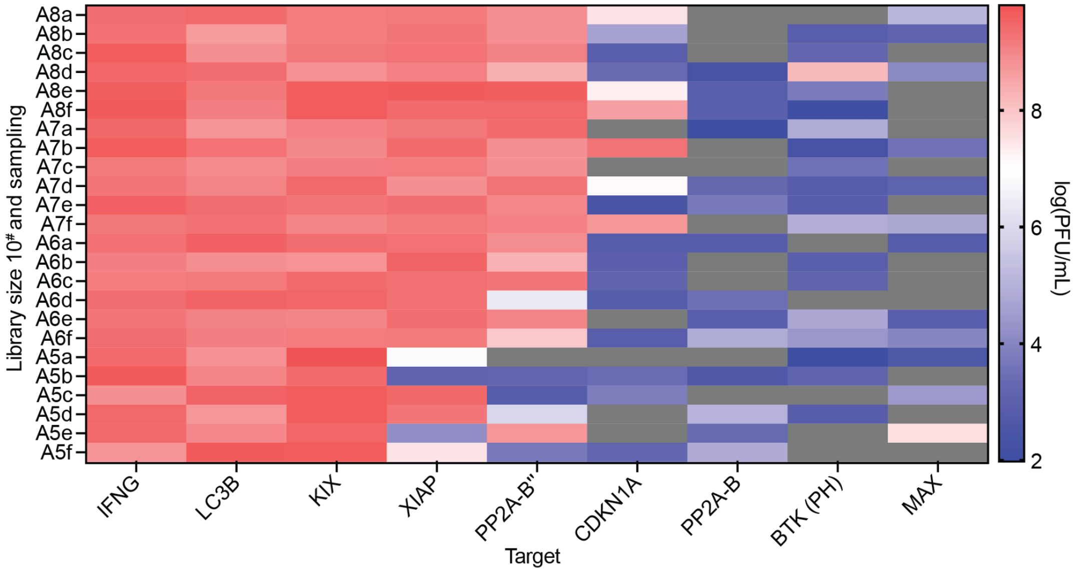

**Supplementary Fig. 2: Heat map of titers following PANCS with independent, varied size samplings.** PANCS was performed on the 9-target panel (Supplementary Table 1) with six independent replicates (a-f) of Library design A with varied sampling sizes ($10^{5-8}$): naming is design, log size, independent transformation as a triplet alphanumeric code. After completion of the sixth passage, the titer was determined by qPCR. Red indicates higher titers, blue lower titers, and gray indicates a titer lower than the threshold of detection (~$10^3$). All selections with a titer >1 × $10^7$ PFU/mL were subject to NGS analysis and validation in the split T7 RNAP *E. coli* luciferase assay. Each titer was determined as a single replicate.

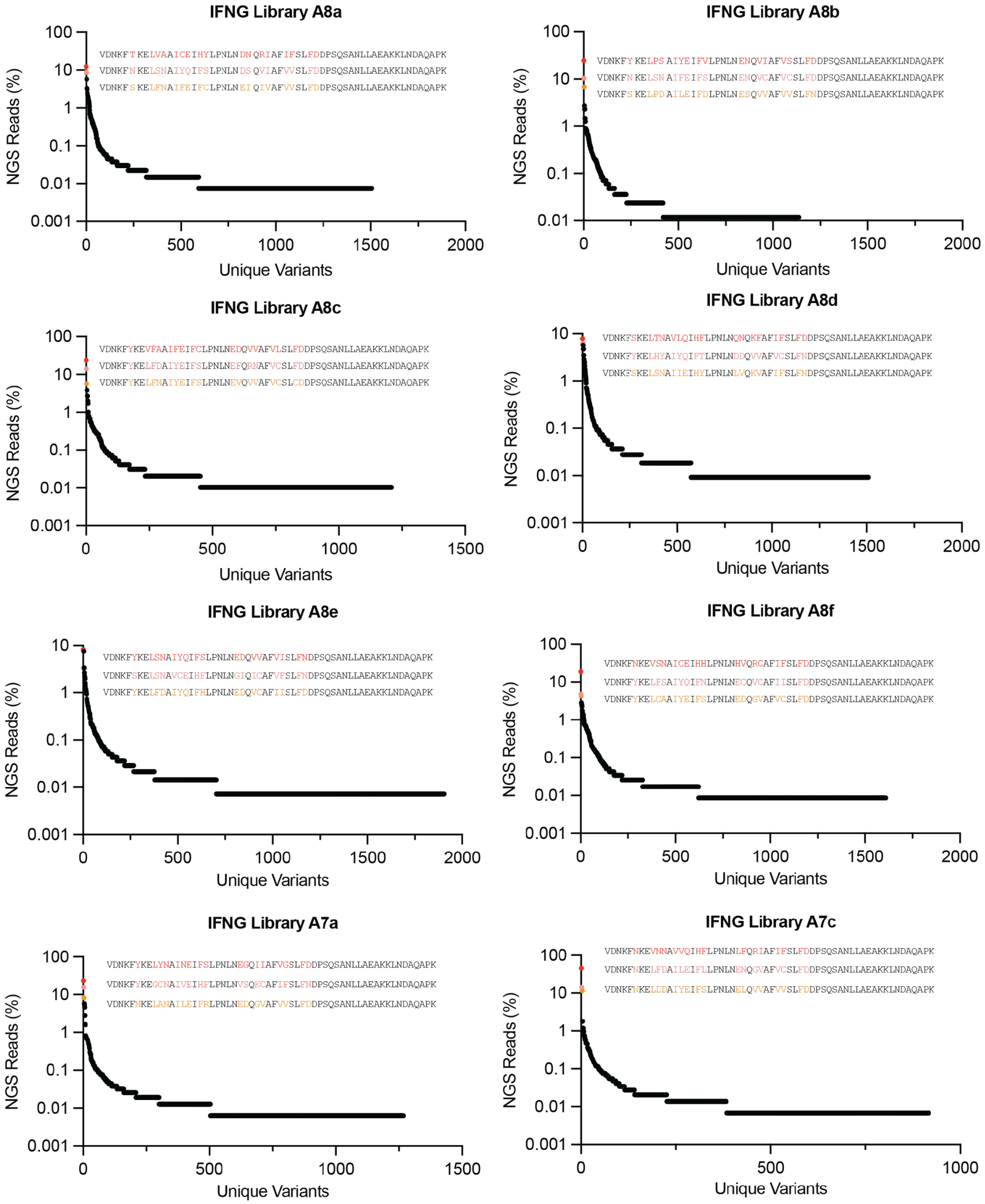
IFNG Library A8a
VDNKFTKELVAAICEIHYLPNLNDNQRIAFIFSLFDDPSQSANLLAEAKKLNDAQAPK
VDNKFNKELSNAIYQIFSLPNLNDSQVIAFVVSLFDDPSQSANLLAEAKKLNDAQAPK
VDNKFSKELFNAIFEIFCLPNLNEIQIVAFVVSLFDDPSQSANLLAEAKKLNDAQAPK
NGS Reads (%)
100
10
1
0.1
0.01
0.001
0
500
1000
1500
2000
Unique Variants
IFNG Library A8b
VDNKFYKELPSAIYEIFVLPNLNENQVIAFVSSLFDDPSQSANLLAEAKKLNDAQAPK
VDNKFNKELSNAIFEIFSLPNLNENQVCAFVCSLFDDPSQSANLLAEAKKLNDAQAPK
VDNKFSKELPDAILEIFDLPNLNESQVVAFVVSLFNDPSQSANLLAEAKKLNDAQAPK
NGS Reads (%)
100
10
1
0.1
0.01
0
500
1000
1500
2000
Unique Variants
IFNG Library A8c
VDNKFYKEVFAAIFEIFCLPNLNEDQVVAFVLSLFDDPSQSANLLAEAKKLNDAQAPK
VDNKFYKELFDAIYEIFSLPNLNEFQRNAFVCSLFDDPSQSANLLAEAKKLNDAQAPK
VDNKFYKELFNAIYEIFSLPNLNEVQVVAFVCSLCDDPSQSANLLAEAKKLNDAQAPK
NGS Reads (%)
100
10
1
0.1
0.01
0.001
0
500
1000
1500
Unique Variants
IFNG Library A8d
VDNKFSKELTNAVLQIHFLPNLNQNQKFAFIFSLFDDPSQSANLLAEAKKLNDAQAPK
VDNKFYKELHYAIYQIFTLPNLNDDQVVAFVCSLFNDPSQSANLLAEAKKLNDAQAPK
VDNKFSKELSNAIIEIHYLPNLNLVQKVAFIFSLFNDPSQSANLLAEAKKLNDAQAPK
NGS Reads (%)
10
1
0.1
0.01
0.001
0
500
1000
1500
2000
Unique Variants
IFNG Library A8e
VDNKFYKELSNAIYQIFSLPNLNEDQVVAFVISLFNDPSQSANLLAEAKKLNDAQAPK
VDNKFSKELSNAVCEIHFLPNLNGIQICAFVFSLFNDPSQSANLLAEAKKLNDAQAPK
VDNKFYKELFDAIYQIFHLPNLNEDQVCAFIISLFDDPSQSANLLAEAKKLNDAQAPK
NGS Reads (%)
10
1
0.1
0.01
0.001
0
500
1000
1500
2000
Unique Variants
IFNG Library A8f
VDNKFNKEVSNAICEIHHLPNLNHVQRCAFIFSLFDDPSQSANLLAEAKKLNDAQAPK
VDNKFYKELFSAIYQIFNLPNLNECQVCAFIISLFDDPSQSANLLAEAKKLNDAQAPK
VDNKFYKELCAAIYEIFSLPNLNEDQGVAFVCSLFDDPSQSANLLAEAKKLNDAQAPK
NGS Reads (%)
100
10
1
0.1
0.01
0.001
0
500
1000
1500
2000
Unique Variants
IFNG Library A7a
VDNKFYKELYNAINEIFSLPNLNEGQIIAFVGSLFDDPSQSANLLAEAKKLNDAQAPK
VDNKFYKEGCNAIVEIHFLPNLNVSQKCAFIFSLFNDPSQSANLLAEAKKLNDAQAPK
VDNKFNKELANAILEIFRLPNLNEDQGVAFVVSLFDDPSQSANLLAEAKKLNDAQAPK
NGS Reads (%)
100
10
1
0.1
0.01
0.001
0
500
1000
1500
Unique Variants
IFNG Library A7c
VDNKFNKEVNNAVVQIHFLPNLNLFQRIAFIFSLFDDPSQSANLLAEAKKLNDAQAPK
VDNKFNKELFDAILEIFLLPNLNENQGVAFVCSLFDDPSQSANLLAEAKKLNDAQAPK
VDNKFNKELDDAIYEIFSLPNLNELQVVAFVVSLFDDPSQSANLLAEAKKLNDAQAPK
NGS Reads (%)
100
10
1
0.1
0.01
0.001
0
500
1000
Unique Variants

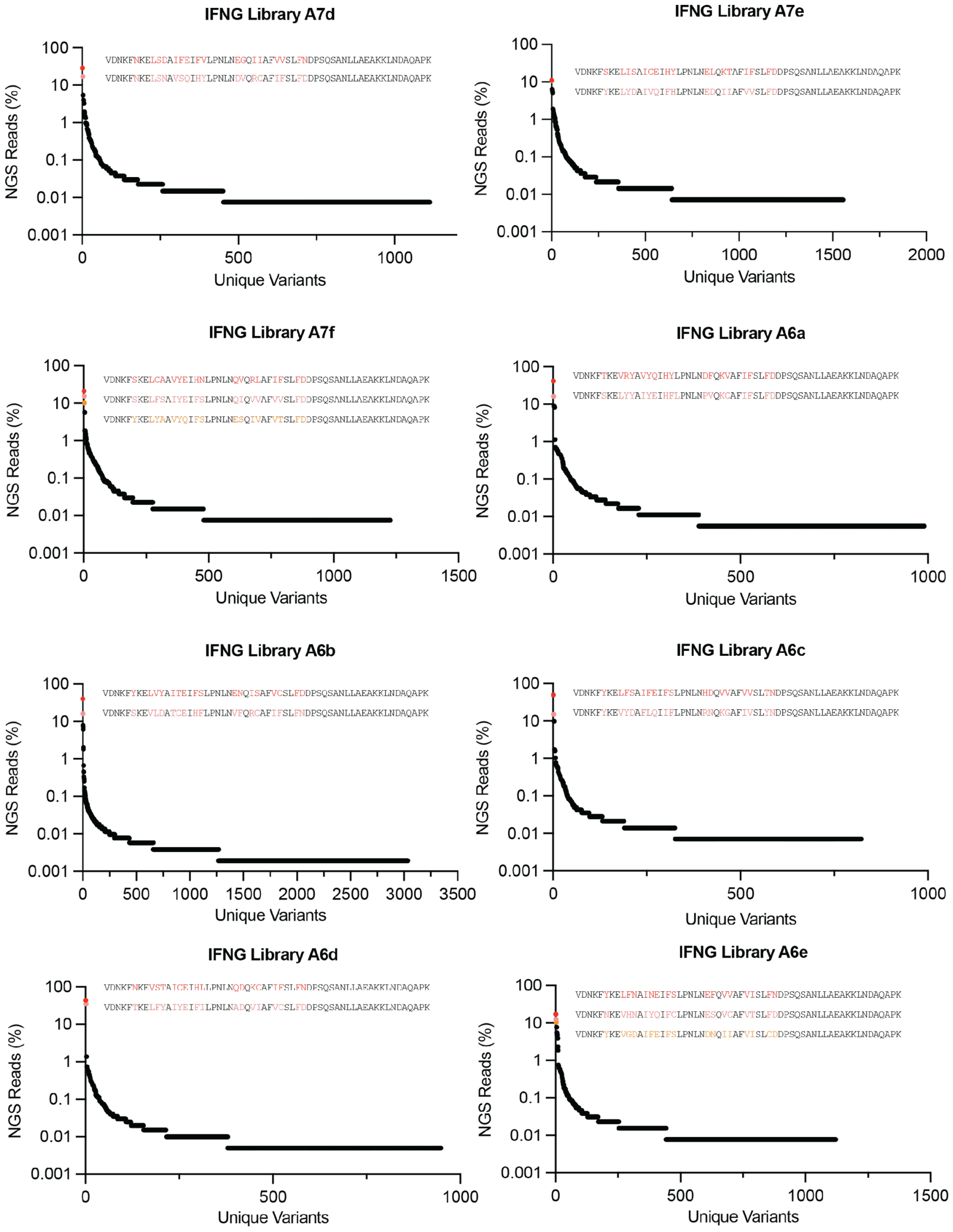
IFNG Library A7d
VDNKFNKELSDAIFEIFVLPNLNEGQIIAFVVSLFNDPSQSANLLAEAKKLNDAQAPK
VDNKFNKELSNAVSQIHYLPNLNDVQRCAFIFSLFDDPSQSANLLAEAKKLNDAQAPK
NGS Reads (%)
100
10
1
0.1
0.01
0.001
0
500
1000
Unique Variants
IFNG Library A7e
VDNKFSKELISAICEIHYLPNLNELQKTAFIFSLFDDPSQSANLLAEAKKLNDAQAPK
VDNKFYKELYDAIVQIFHLPNLNEDQIIAFVVSLFDDPSQSANLLAEAKKLNDAQAPK
NGS Reads (%)
100
10
1
0.1
0.01
0.001
0
500
1000
1500
2000
Unique Variants
IFNG Library A7f
VDNKFSKELCAAVYEIHNLPNLNQVQRLAFIFSLFDDPSQSANLLAEAKKLNDAQAPK
VDNKFSKELFSAIYEIFSLPNLNQIQVVAFVVSLFDDPSQSANLLAEAKKLNDAQAPK
VDNKFYKELYAAVYQIFSLPNLNESQIVAFVTSLFDDPSQSANLLAEAKKLNDAQAPK
NGS Reads (%)
100
10
1
0.1
0.01
0.001
0
500
1000
1500
Unique Variants
IFNG Library A6a
VDNKFTKEVRYAVYQIHYLPNLNDFQKVAFIFSLFDDPSQSANLLAEAKKLNDAQAPK
VDNKFSKELYYAIYEIHFLPNLNPVQKCAFIFSLFDDPSQSANLLAEAKKLNDAQAPK
NGS Reads (%)
100
10
1
0.1
0.01
0.001
0
500
1000
Unique Variants
IFNG Library A6b
VDNKFYKELVYAITEIFSLPNLNENQISAFVCSLFDDPSQSANLLAEAKKLNDAQAPK
VDNKFSKEVLDATCEIHFLPNLNVFQRCAFIFSLFNDPSQSANLLAEAKKLNDAQAPK
NGS Reads (%)
100
10
1
0.1
0.01
0.001
0
500
1000
1500
2000
2500
3000
3500
Unique Variants
IFNG Library A6c
VDNKFYKELFSAIFEIFSLPNLNHDQVVAFVVSLTNDPSQSANLLAEAKKLNDAQAPK
VDNKFYKEVYDAFLQIIFLPNLNRNQKGAFIVSLYNDPSQSANLLAEAKKLNDAQAPK
NGS Reads (%)
100
10
1
0.1
0.01
0.001
0
500
1000
Unique Variants
IFNG Library A6d
VDNKFNKEVSTAICEIHLLPNLNQDQKCAFIFSLFNDPSQSANLLAEAKKLNDAQAPK
VDNKFTKELFYAIYEIFILPNLNADQVIAFVCSLFDDPSQSANLLAEAKKLNDAQAPK
NGS Reads (%)
100
10
1
0.1
0.01
0.001
0
500
1000
Unique Variants
IFNG Library A6e
VDNKFYKELFNAINEIFSLPNLNEFQVVAFVISLFNDPSQSANLLAEAKKLNDAQAPK
VDNKFNKEVHNAIYQIFCLPNLNESQVCAFVTSLFDDPSQSANLLAEAKKLNDAQAPK
VDNKFYKEVGDAIFEIFSLPNLNDNQIIAFVISLCDDPSQSANLLAEAKKLNDAQAPK
NGS Reads (%)
100
10
1
0.1
0.01
0.001
0
500
1000
1500
Unique Variants

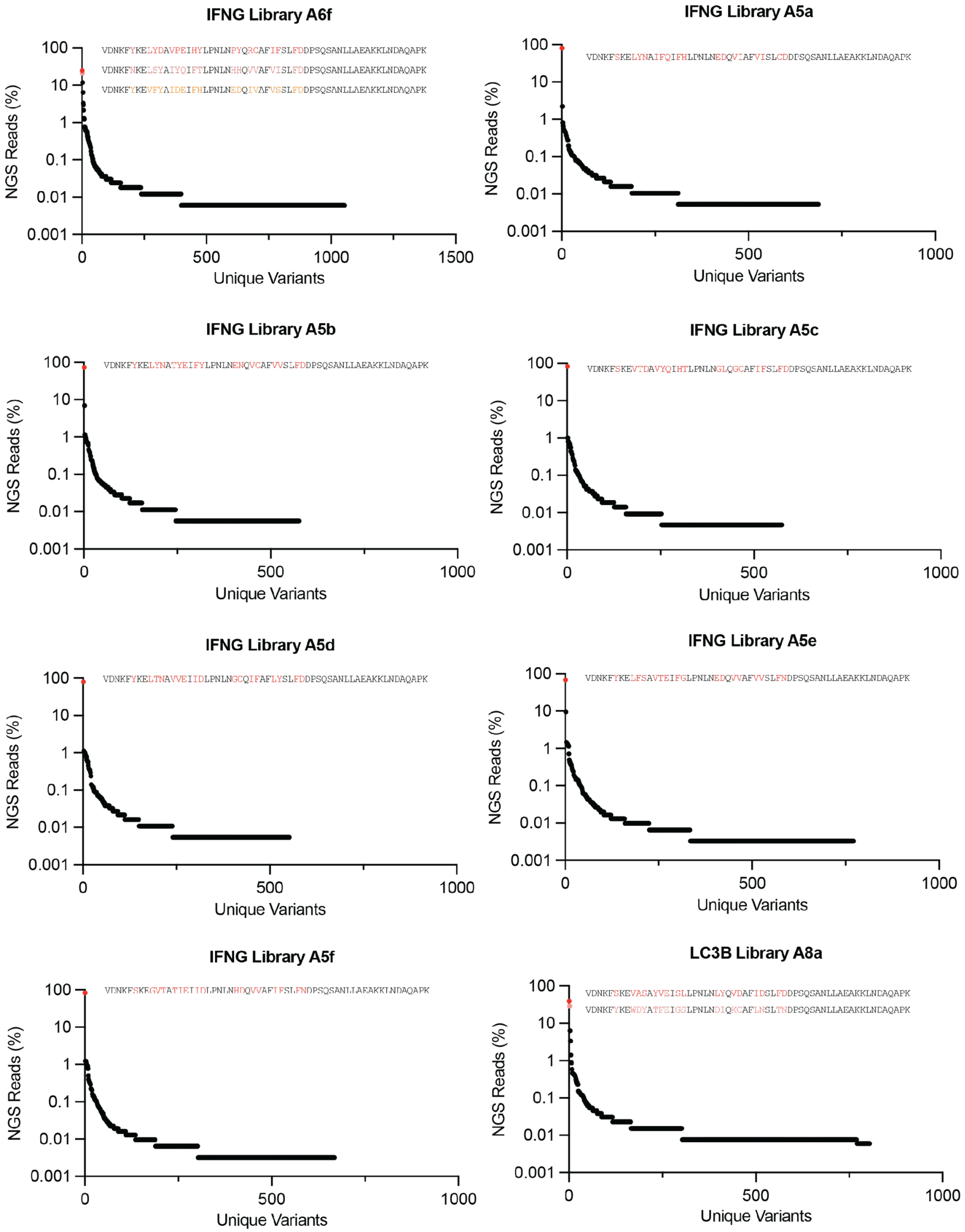
IFNG Library A6f
VDNKFYKELYDAVPEIHYLPNLNPYQRCAFIFSLFDDPSQSANLLAEAKKLNDAQAPK
VDNKFNKELSYAIYQIFTLPNLNHHQVVAFVISLFDDPSQSANLLAEAKKLNDAQAPK
VDNKFYKEVFYAIDEIFHLPNLNEDQIVAFVSSLFDDPSQSANLLAEAKKLNDAQAPK
NGS Reads (%)
100
10
1
0.1
0.01
0.001
0
500
1000
1500
Unique Variants
IFNG Library A5a
VDNKFSKELYNAIFQIFHLPNLNEDQVIAFVISLCDDPSQSANLLAEAKKLNDAQAPK
NGS Reads (%)
100
10
1
0.1
0.01
0.001
0
500
1000
Unique Variants
IFNG Library A5b
VDNKFYKELYNATYEIFYLPNLNENQVCAFVVSLFDDPSQSANLLAEAKKLNDAQAPK
NGS Reads (%)
100
10
1
0.1
0.01
0.001
0
500
1000
Unique Variants
IFNG Library A5c
VDNKFSKEVTDAVYQIHTLPNLNGLQGCAFIFSLFDDPSQSANLLAEAKKLNDAQAPK
NGS Reads (%)
100
10
1
0.1
0.01
0.001
0
500
1000
Unique Variants
IFNG Library A5d
VDNKFYKELTNAVVEIIDLPNLNGCQIFAFLYSLFDDPSQSANLLAEAKKLNDAQAPK
NGS Reads (%)
100
10
1
0.1
0.01
0.001
0
500
1000
Unique Variants
IFNG Library A5e
VDNKFYKELFSAVTEIFGLPNLNEDQVVAFVVSLFNDPSQSANLLAEAKKLNDAQAPK
NGS Reads (%)
100
10
1
0.1
0.01
0.001
0
500
1000
Unique Variants
IFNG Library A5f
VDNKFSKEGVTATIEIIDLPNLNHDQVVAFIFSLFNDPSQSANLLAEAKKLNDAQAPK
NGS Reads (%)
100
10
1
0.1
0.01
0.001
0
500
1000
Unique Variants
LC3B Library A8a
VDNKFSKEVASAYVEISLLPNLNLYQVDAFIDSLFDDPSQSANLLAEAKKLNDAQAPK
VDNKFYKEWDYATFEIGSLPNLNDIQKCAFLNSLTNDPSQSANLLAEAKKLNDAQAPK
NGS Reads (%)
100
10
1
0.1
0.01
0.001
0
500
1000
Unique Variants

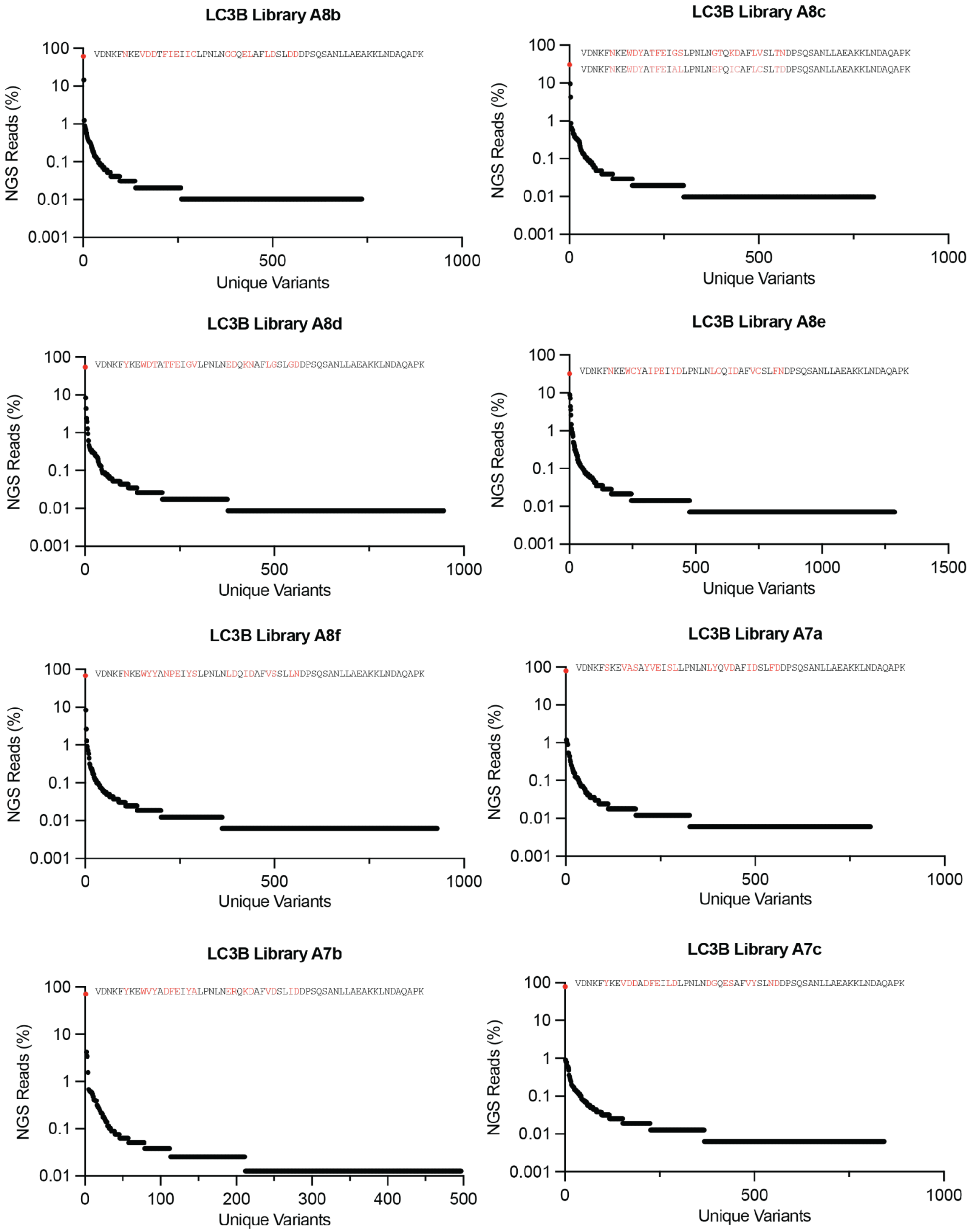
LC3B Library A8b
LC3B Library A8c
LC3B Library A8d
LC3B Library A8e
LC3B Library A8f
LC3B Library A7a
LC3B Library A7b
LC3B Library A7c
NGS Reads (%)
Unique Variants

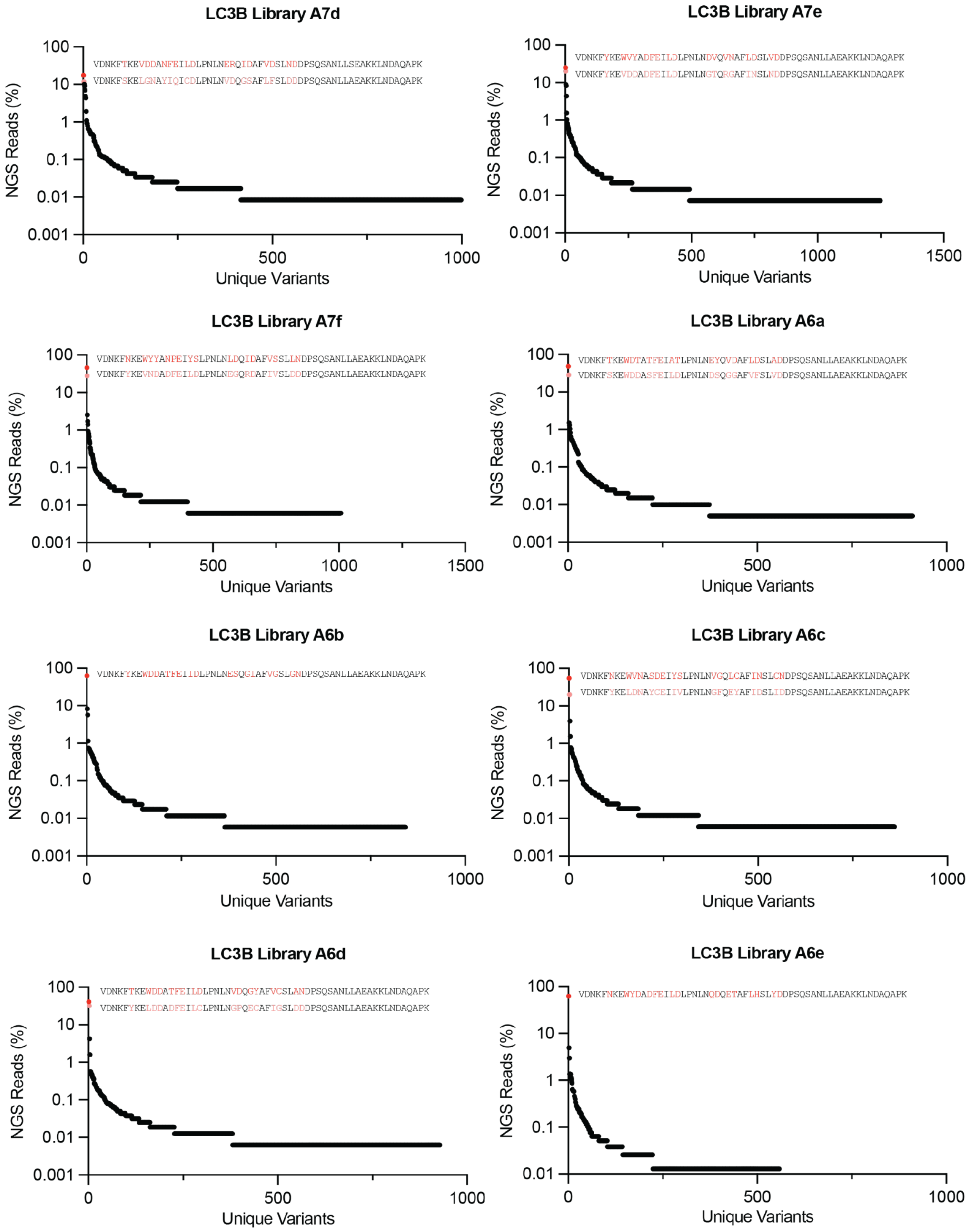

LC3B Library A7d
VDNKFTKEVDDANFEILDLPNLNERQIDAFVDSLNDDPSQSANLLSEAKKLNDAQAPK
VDNKFSKELGNAYIQICDLPNLNVDQGSAFLFSLDDDPSQSANLLAEAKKLNDAQAPK
NGS Reads (%)
100
10
1
0.1
0.01
0.001
0
500
1000
Unique Variants
LC3B Library A7e
VDNKFYKEWVYADFEILDLPNLNDVQVNAFLDSLVDDPSQSANLLAEAKKLNDAQAPK
VDNKFYKEVDDADFEILDLPNLNGTQRGAFINSLNDDPSQSANLLAEAKKLNDAQAPK
NGS Reads (%)
100
10
1
0.1
0.01
0.001
0
500
1000
1500
Unique Variants
LC3B Library A7f
VDNKFNKEWYYANPEIYSLPNLNLDQIDAFVSSLLNDPSQSANLLAEAKKLNDAQAPK
VDNKFYKEVNDADFEILDLPNLNEGQRDAFIVSLDDDPSQSANLLAEAKKLNDAQAPK
NGS Reads (%)
100
10
1
0.1
0.01
0.001
0
500
1000
1500
Unique Variants
LC3B Library A6a
VDNKFTKEWDTATFEIATLPNLNEYQVDAFLDSLADDPSQSANLLAEAKKLNDAQAPK
VDNKFSKEWDDASFEILDLPNLNDSQGGAFVFSLVDDPSQSANLLAEAKKLNDAQAPK
NGS Reads (%)
100
10
1
0.1
0.01
0.001
0
500
1000
Unique Variants
LC3B Library A6b
VDNKFYKEWDDATFEIIDLPNLNESQGIAFVGSLGNDPSQSANLLAEAKKLNDAQAPK
NGS Reads (%)
100
10
1
0.1
0.01
0.001
0
500
1000
Unique Variants
LC3B Library A6c
VDNKFNKEWVNASDEIYSLPNLNVGQLCAFINSLCNDPSQSANLLAEAKKLNDAQAPK
VDNKFYKELDNAYCEIIVLPNLNGFQEYAFIDSLIDDPSQSANLLAEAKKLNDAQAPK
NGS Reads (%)
100
10
1
0.1
0.01
0.001
0
500
1000
Unique Variants
LC3B Library A6d
VDNKFTKEWDDATFEILDLPNLNVDQGYAFVCSLANDPSQSANLLAEAKKLNDAQAPK
VDNKFYKELDDADFEILCLPNLNGPQECAFIGSLDDDPSQSANLLAEAKKLNDAQAPK
NGS Reads (%)
100
10
1
0.1
0.01
0.001
0
500
1000
Unique Variants
LC3B Library A6e
VDNKFNKEWYDADFEILDLPNLNQDQETAFLHSLYDDPSQSANLLAEAKKLNDAQAPK
NGS Reads (%)
100
10
1
0.1
0.01
0
500
1000
Unique Variants

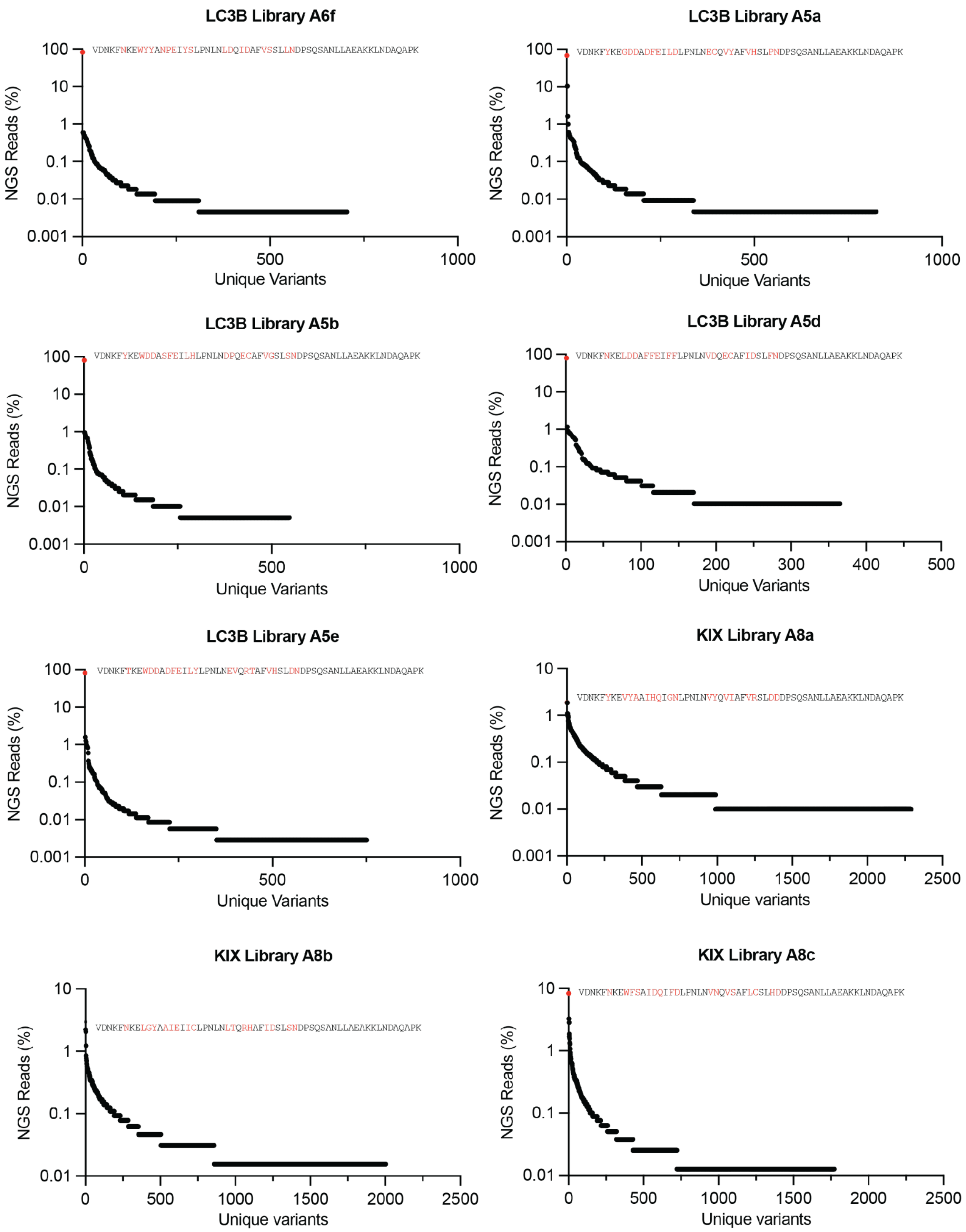

LC3B Library A6f
VDNKFNKEWYYANPEIYSLPNLNLDQIDAFVSSLLNDPSQSANLLAEAKKLNDAQAPK
NGS Reads (%)
100
10
1
0.1
0.01
0.001
0
500
1000
Unique Variants
LC3B Library A5a
VDNKFYKEGDDADFEILDLPNLNECQVYAFVHSLPNDPSQSANLLAEAKKLNDAQAPK
NGS Reads (%)
100
10
1
0.1
0.01
0.001
0
500
1000
Unique Variants
LC3B Library A5b
VDNKFYKEWDDASFEILHLPNLNDPQECAFVGSLSNDPSQSANLLAEAKKLNDAQAPK
NGS Reads (%)
100
10
1
0.1
0.01
0.001
0
500
1000
Unique Variants
LC3B Library A5d
VDNKFNKELDDAFFEIFFLPNLNVDQECAFIDSLFNDPSQSANLLAEAKKLNDAQAPK
NGS Reads (%)
100
10
1
0.1
0.01
0.001
0
100
200
300
400
500
Unique Variants
LC3B Library A5e
VDNKFTKEWDDADFEILYLPNLNEVQRTAFVHSLDNDPSQSANLLAEAKKLNDAQAPK
NGS Reads (%)
100
10
1
0.1
0.01
0.001
0
500
1000
Unique Variants
KIX Library A8a
VDNKFYKEVYAAIHQIGNLPNLNVYQVIAFVRSLDDDPSQSANLLAEAKKLNDAQAPK
NGS Reads (%)
10
1
0.1
0.01
0.001
0
500
1000
1500
2000
2500
Unique variants
KIX Library A8b
VDNKFNKELGYAAIEIICLPNLNLTQRHAFIDSLSNDPSQSANLLAEAKKLNDAQAPK
NGS Reads (%)
10
1
0.1
0.01
0
500
1000
1500
2000
2500
Unique variants
KIX Library A8c
VDNKFNKEWFSAIDQIFDLPNLNVNQVSAFLCSLHDDPSQSANLLAEAKKLNDAQAPK
NGS Reads (%)
10
1
0.1
0.01
0
500
1000
1500
2000
2500
Unique variants

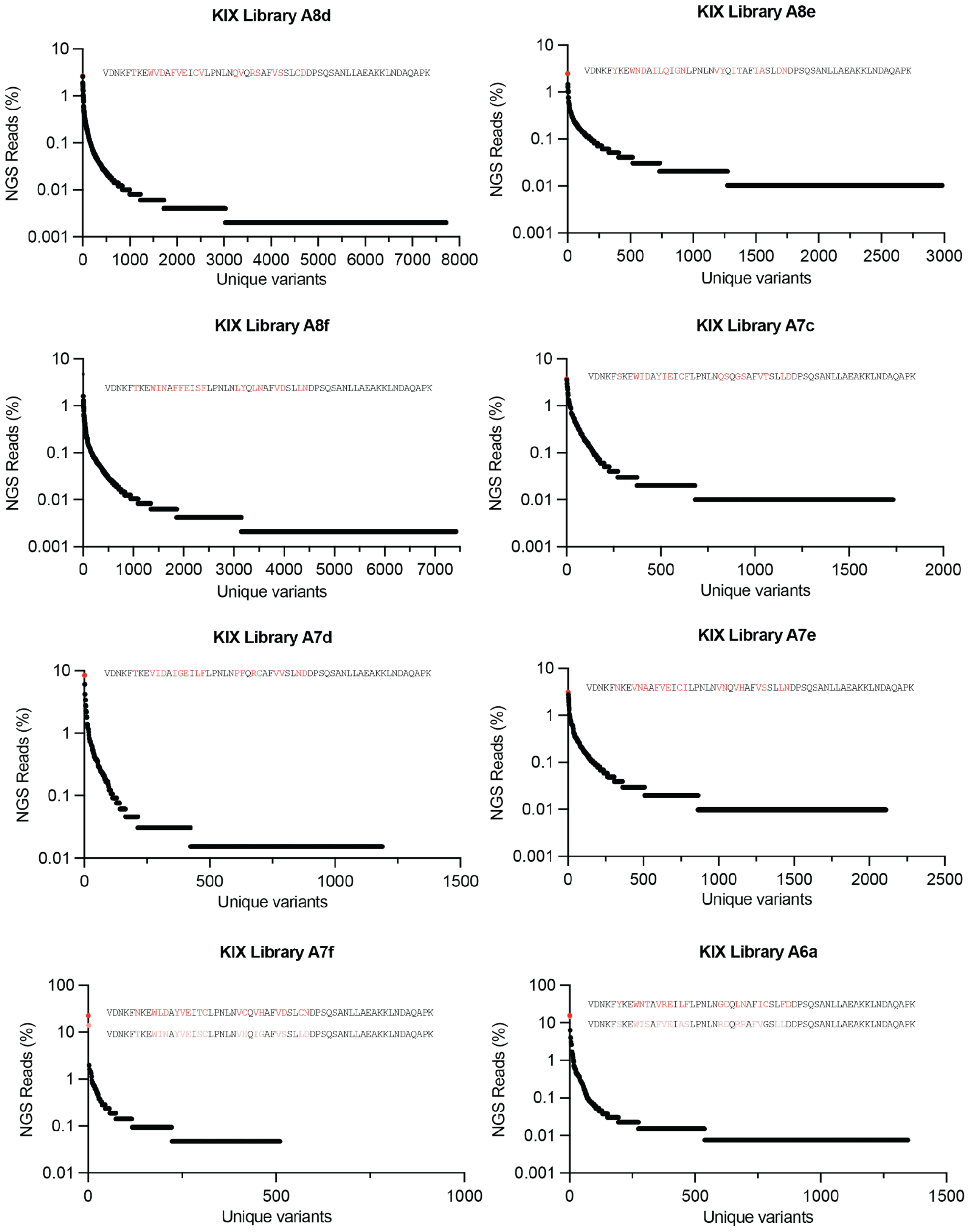
KIX Library A8d
VDNKFTKEWVDAFVEICVLPNLNQVQRSAFVSSLCDDPSQSANLLAEAKKLNDAQAPK
NGS Reads (%)
10
1
0.1
0.01
0.001
0
1000
2000
3000
4000
5000
6000
7000
8000
Unique variants
KIX Library A8e
VDNKFYKEWNDAILQIGNLPNLNVYQITAFIASLDNDPSQSANLLAEAKKLNDAQAPK
NGS Reads (%)
10
1
0.1
0.01
0.001
0
500
1000
1500
2000
2500
3000
Unique variants
KIX Library A8f
VDNKFTKEWINAFFEISFLPNLNLYQLNAFVDSLLNDPSQSANLLAEAKKLNDAQAPK
NGS Reads (%)
10
1
0.1
0.01
0.001
0
1000
2000
3000
4000
5000
6000
7000
Unique variants
KIX Library A7c
VDNKFSKEWIDAYIEICFLPNLNQSQGSAFVTSLLDDPSQSANLLAEAKKLNDAQAPK
NGS Reads (%)
10
1
0.1
0.01
0.001
0
500
1000
1500
2000
Unique variants
KIX Library A7d
VDNKFTKEVIDAIGEILFLPNLNPFQRCAFVVSLNDDPSQSANLLAEAKKLNDAQAPK
NGS Reads (%)
10
1
0.1
0.01
0
500
1000
1500
Unique variants
KIX Library A7e
VDNKFNKEVNAAFVEICILPNLNVNQVHAFVSSLLNDPSQSANLLAEAKKLNDAQAPK
NGS Reads (%)
10
1
0.1
0.01
0.001
0
500
1000
1500
2000
2500
Unique variants
KIX Library A7f
VDNKFNKEWLDAYVEITCLPNLNVCQVHAFVDSLCNDPSQSANLLAEAKKLNDAQAPK
VDNKFTKEWINAYVEISCLPNLNVHQIGAFVSSLLDDPSQSANLLAEAKKLNDAQAPK
NGS Reads (%)
100
10
1
0.1
0.01
0
500
1000
Unique variants
KIX Library A6a
VDNKFYKEWNTAVREILFLPNLNGCQLNAFICSLFDDPSQSANLLAEAKKLNDAQAPK
VDNKFSKEWISAFVEIASLPNLNRCQRRAFVGSLLDDPSQSANLLAEAKKLNDAQAPK
NGS Reads (%)
100
10
1
0.1
0.01
0.001
0
500
1000
1500
Unique variants

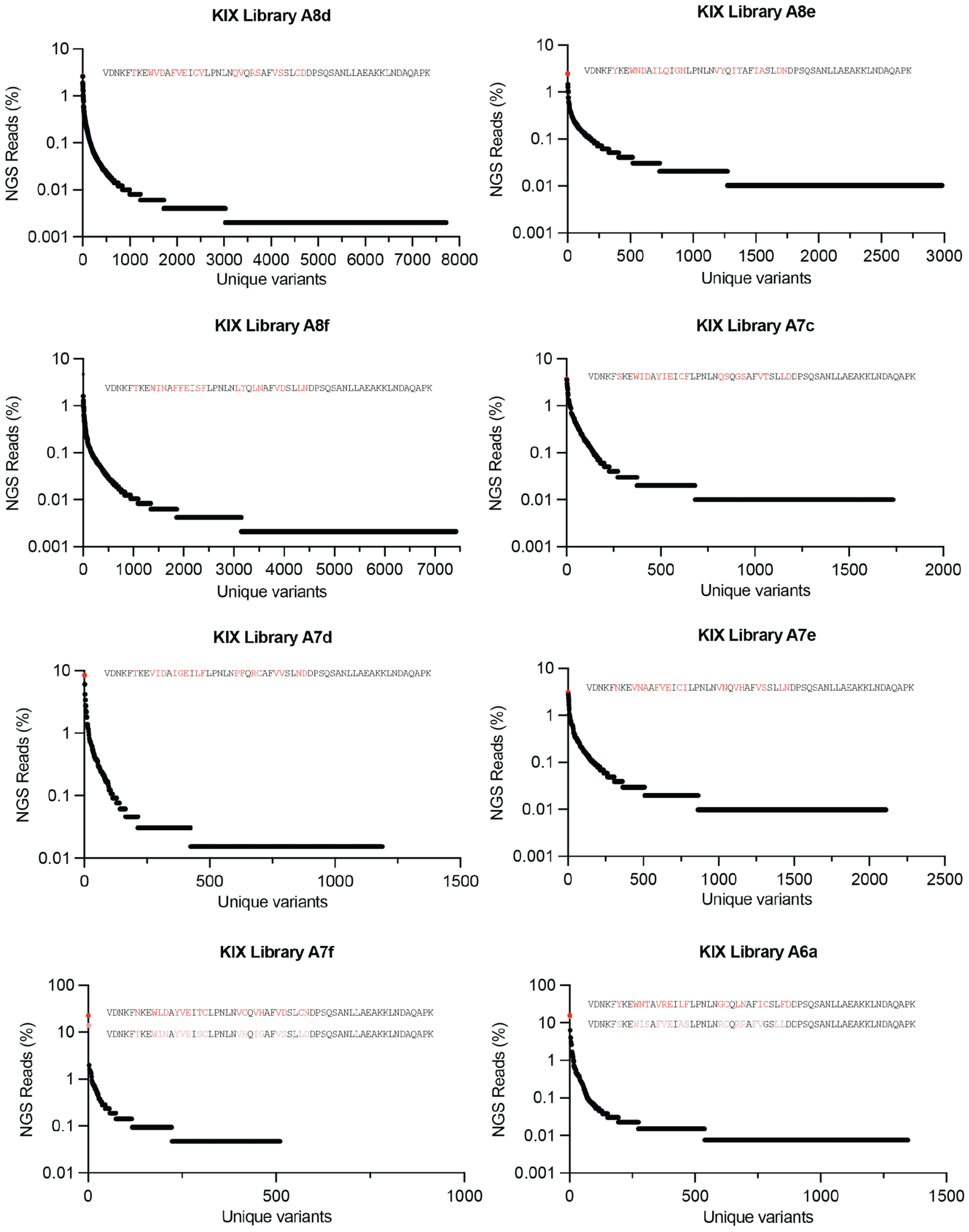
KIX Library A8d
VDNKFTKEWVDAFVEICVLPNLNQVQRSAFVSSLCDDPSQSANLLAEAKKLNDAQAPK
NGS Reads (%)
10
1
0.1
0.01
0.001
0
1000
2000
3000
4000
5000
6000
7000
8000
Unique variants
KIX Library A8e
VDNKFYKEWNDAILQIGNLPNLNVYQITAFIASLDNDPSQSANLLAEAKKLNDAQAPK
NGS Reads (%)
10
1
0.1
0.01
0.001
0
500
1000
1500
2000
2500
3000
Unique variants
KIX Library A8f
VDNKFTKEWINAFFEISFLPNLNLYQLNAFVDSLLNDPSQSANLLAEAKKLNDAQAPK
NGS Reads (%)
10
1
0.1
0.01
0.001
0
1000
2000
3000
4000
5000
6000
7000
Unique variants
KIX Library A7c
VDNKFSKEWIDAYIEICFLPNLNQSQGSAFVTSLLDDPSQSANLLAEAKKLNDAQAPK
NGS Reads (%)
10
1
0.1
0.01
0.001
0
500
1000
1500
2000
Unique variants
KIX Library A7d
VDNKFTKEVIDAIGEILFLPNLNPFQRCAFVVSLNDDPSQSANLLAEAKKLNDAQAPK
NGS Reads (%)
10
1
0.1
0.01
0
500
1000
1500
Unique variants
KIX Library A7e
VDNKFNKEVNAAFVEICILPNLNVNQVHAFVSSLLNDPSQSANLLAEAKKLNDAQAPK
NGS Reads (%)
10
1
0.1
0.01
0.001
0
500
1000
1500
2000
2500
Unique variants
KIX Library A7f
VDNKFNKEWLDAYVEITCLPNLNVCQVHAFVDSLCNDPSQSANLLAEAKKLNDAQAPK
VDNKFTKEWINAYVEISCLPNLNVHQIGAFVSSLLDDPSQSANLLAEAKKLNDAQAPK
NGS Reads (%)
100
10
1
0.1
0.01
0
500
1000
Unique variants
KIX Library A6a
VDNKFYKEWNTAVREILFLPNLNGCQLNAFICSLFDDPSQSANLLAEAKKLNDAQAPK
VDNKFSKEWISAFVEIASLPNLNRCQRRAFVGSLLDDPSQSANLLAEAKKLNDAQAPK
NGS Reads (%)
100
10
1
0.1
0.01
0.001
0
500
1000
1500
Unique variants

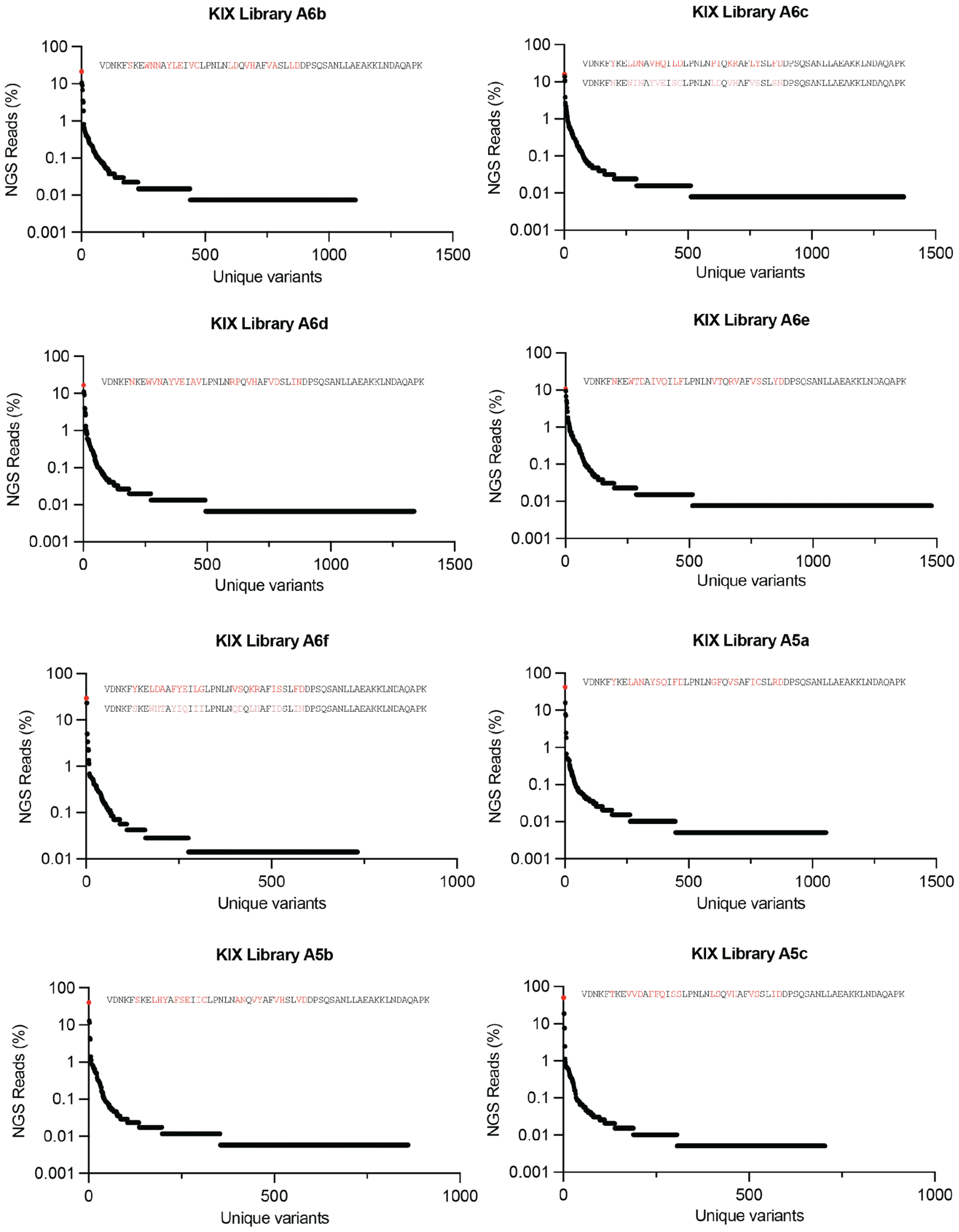

KIX Library A6b
VDNKFSKEWNNAYLEIVCLPNLNLDQVHAFVASLLDDPSQSANLLAEAKKLNDAQAPK
NGS Reads (%)
100
10
1
0.1
0.01
0.001
0
500
1000
1500
Unique variants
KIX Library A6c
VDNKFYKELDNAVHQILDLPNLNPIQKRAFLYSLFDDPSQSANLLAEAKKLNDAQAPK
VDNKFNKEWINAYVEISCLPNLNLDQVHAFVSSLSNDPSQSANLLAEAKKLNDAQAPK
NGS Reads (%)
100
10
1
0.1
0.01
0.001
0
500
1000
1500
Unique variants
KIX Library A6d
VDNKFNKEWVNAYVEIAVLPNLNRPQVHAFVDSLINDPSQSANLLAEAKKLNDAQAPK
NGS Reads (%)
100
10
1
0.1
0.01
0.001
0
500
1000
1500
Unique variants
KIX Library A6e
VDNKFNKEWTDAIVQILFLPNLNVTQRVAFVSSLYDDPSQSANLLAEAKKLNDAQAPK
NGS Reads (%)
100
10
1
0.1
0.01
0.001
0
500
1000
1500
Unique variants
KIX Library A6f
VDNKFYKELDAAFYEILGLPNLNVSQKRAFISSLFDDPSQSANLLAEAKKLNDAQAPK
VDNKFSKEWIITAYIQIIILPNLNQDQLIIAFIDSLINDPSQSANLLAEAKKLNDAQAPK
NGS Reads (%)
100
10
1
0.1
0.01
0
500
1000
Unique variants
KIX Library A5a
VDNKFYKELANAYSQIFDLPNLNGFQVSAFICSLRDDPSQSANLLAEAKKLNDAQAPK
NGS Reads (%)
100
10
1
0.1
0.01
0.001
0
500
1000
1500
Unique variants
KIX Library A5b
VDNKFSKELHYAFSEIICLPNLNANQVYAFVHSLVDDPSQSANLLAEAKKLNDAQAPK
NGS Reads (%)
100
10
1
0.1
0.01
0.001
0
500
1000
Unique variants
KIX Library A5c
VDNKFTKEVVDAFFQISSLPNLNLSQVIIAFVSSLIDDPSQSANLLAEAKKLNDAQAPK
NGS Reads (%)
100
10
1
0.1
0.01
0.001
0
500
1000
Unique variants

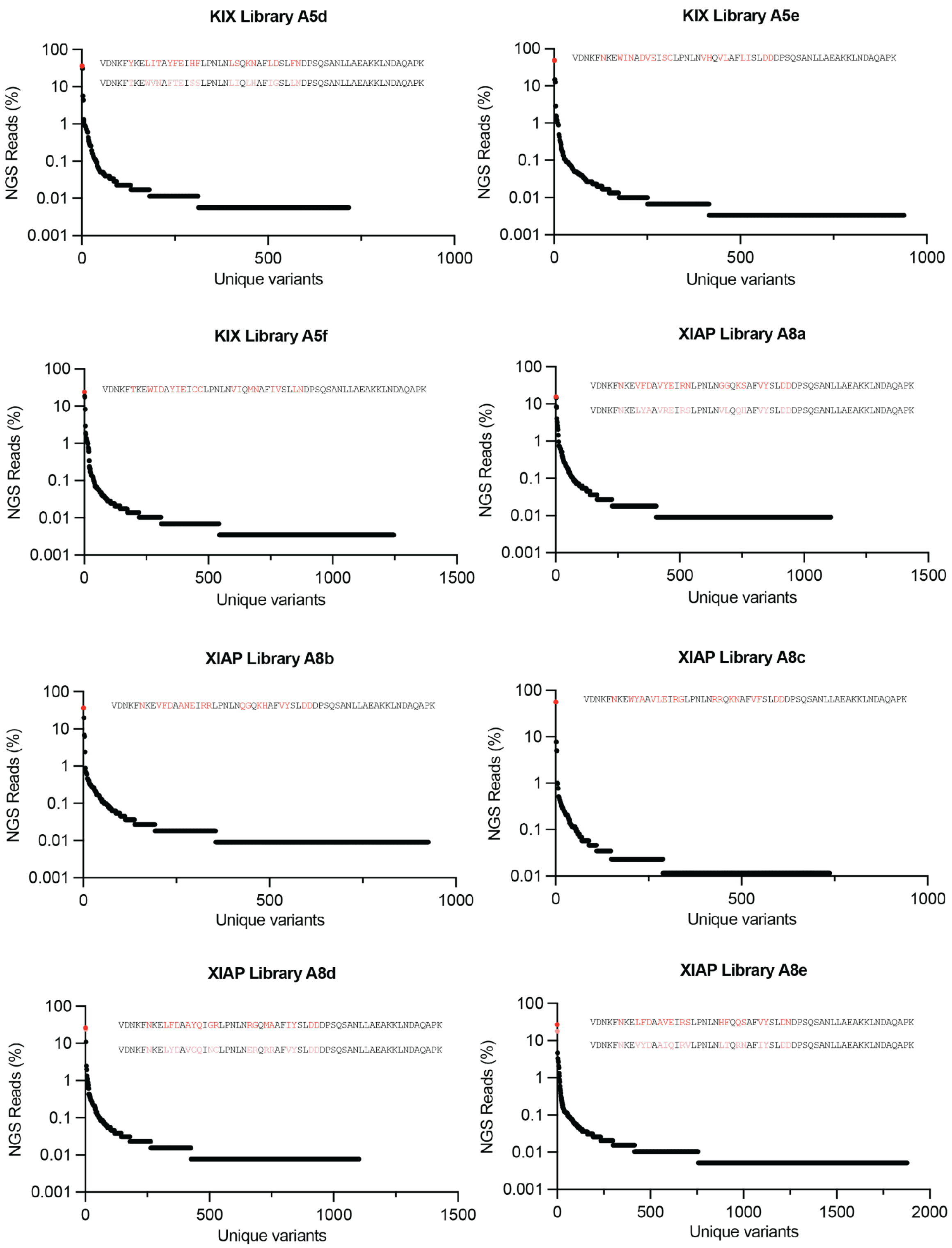
KIX Library A5d
VDNKFYKELITAYFEIHFLPNLNLSQKNAFLDSLFNDPSQSANLLAEAKKLNDAQAPK
VDNKFTKEWVNAFTEISSLPNLNLIQLHAFIGSLLNDPSQSANLLAEAKKLNDAQAPK
NGS Reads (%)
Unique variants
KIX Library A5e
VDNKFNKEWINADVEISCLPNLNVHQVLAFLISLDDDPSQSANLLAEAKKLNDAQAPK
NGS Reads (%)
Unique variants
KIX Library A5f
VDNKFTKEWIDAYIEICCLPNLNVIQMNAFIVSLLNDPSQSANLLAEAKKLNDAQAPK
NGS Reads (%)
Unique variants
XIAP Library A8a
VDNKFNKEVFDAVYEIRNLPNLNGGQKSAFVYSLDDDPSQSANLLAEAKKLNDAQAPK
VDNKFNKELYAAVREIRSLPNLNVLQQHAFVYSLDDDPSQSANLLAEAKKLNDAQAPK
NGS Reads (%)
Unique variants
XIAP Library A8b
VDNKFNKEVFDAANEIRRLPNLNQGQKHAFVYSLDDDPSQSANLLAEAKKLNDAQAPK
NGS Reads (%)
Unique variants
XIAP Library A8c
VDNKFNKEWYAAVLEIRGLPNLNRRQKNAFVFSLDDDPSQSANLLAEAKKLNDAQAPK
NGS Reads (%)
Unique variants
XIAP Library A8d
VDNKFNKELFDAAYQIGRLPNLNRGQMAAFIYSLDDDPSQSANLLAEAKKLNDAQAPK
VDNKFNKELYDAVCQINCLPNLNERQRRAFVYSLDDDPSQSANLLAEAKKLNDAQAPK
NGS Reads (%)
Unique variants
XIAP Library A8e
VDNKFNKELFDAAVEIRSLPNLNHFQQSAFVYSLDNDPSQSANLLAEAKKLNDAQAPK
VDNKFNKEVYDAAIQIRVLPNLNLTQRNAFIYSLDDDPSQSANLLAEAKKLNDAQAPK
NGS Reads (%)
Unique variants

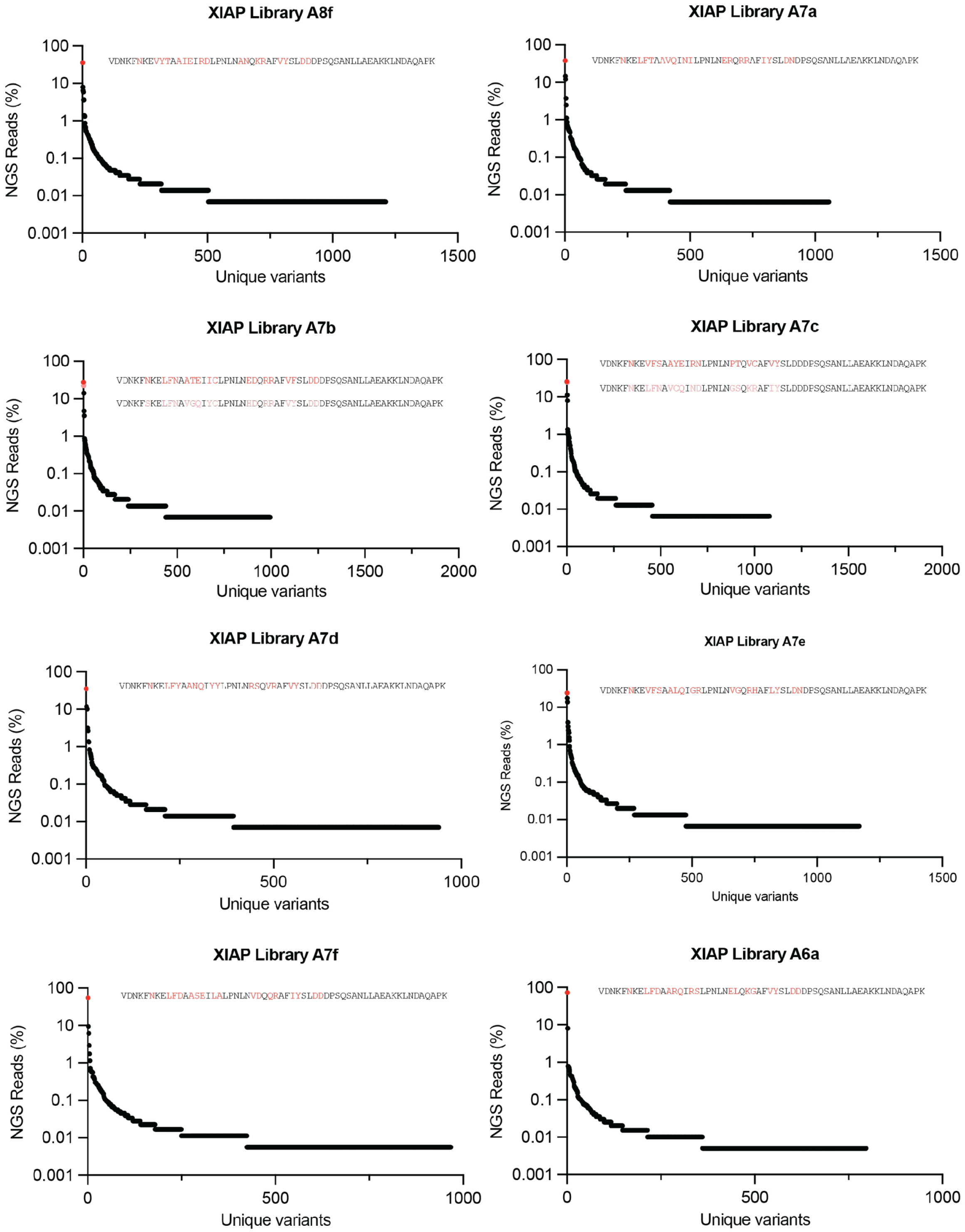

XIAP Library A8f
VDNKFNKEVYTAAIEIRDLPNLNANQKRAFVYSLDDDPSQSANLLAEAKKLNDAQAPK
NGS Reads (%)
Unique variants
XIAP Library A7a
VDNKFNKELFTAAVQINILPNLNERQRRAFIYSLDNDPSQSANLLAEAKKLNDAQAPK
NGS Reads (%)
Unique variants
XIAP Library A7b
VDNKFNKELFNAATEIICLPNLNEDQRRAFVFSLDDDPSQSANLLAEAKKLNDAQAPK
VDNKFSKELFNAVGQIYCLPNLNHDQRRAFVYSLDDDPSQSANLLAEAKKLNDAQAPK
NGS Reads (%)
Unique variants
XIAP Library A7c
VDNKFNKEVFSAAYEIRNLPNLNPTQVCAFVYSLDDDPSQSANLLAEAKKLNDAQAPK
VDNKFNKELFNAVCQINDLPNLNGSQKRAFIYSLDDDPSQSANLLAEAKKLNDAQAPK
NGS Reads (%)
Unique variants
XIAP Library A7d
VDNKFNKELFYAANQTYYLPNLNRSQVRAFVYSLDDDPSQSANLLAEAKKLNDAQAPK
NGS Reads (%)
Unique variants
XIAP Library A7e
VDNKFNKEVFSAALQIGRLPNLNVGQRHAFLYSLDNDPSQSANLLAEAKKLNDAQAPK
NGS Reads (%)
Unique variants
XIAP Library A7f
VDNKFNKELFDAASEILALPNLNVDQQRAFIYSLDDDPSQSANLLAEAKKLNDAQAPK
NGS Reads (%)
Unique variants
XIAP Library A6a
VDNKFNKELFDAARQIRSLPNLNELQKGAFVYSLDDDPSQSANLLAEAKKLNDAQAPK
NGS Reads (%)
Unique variants

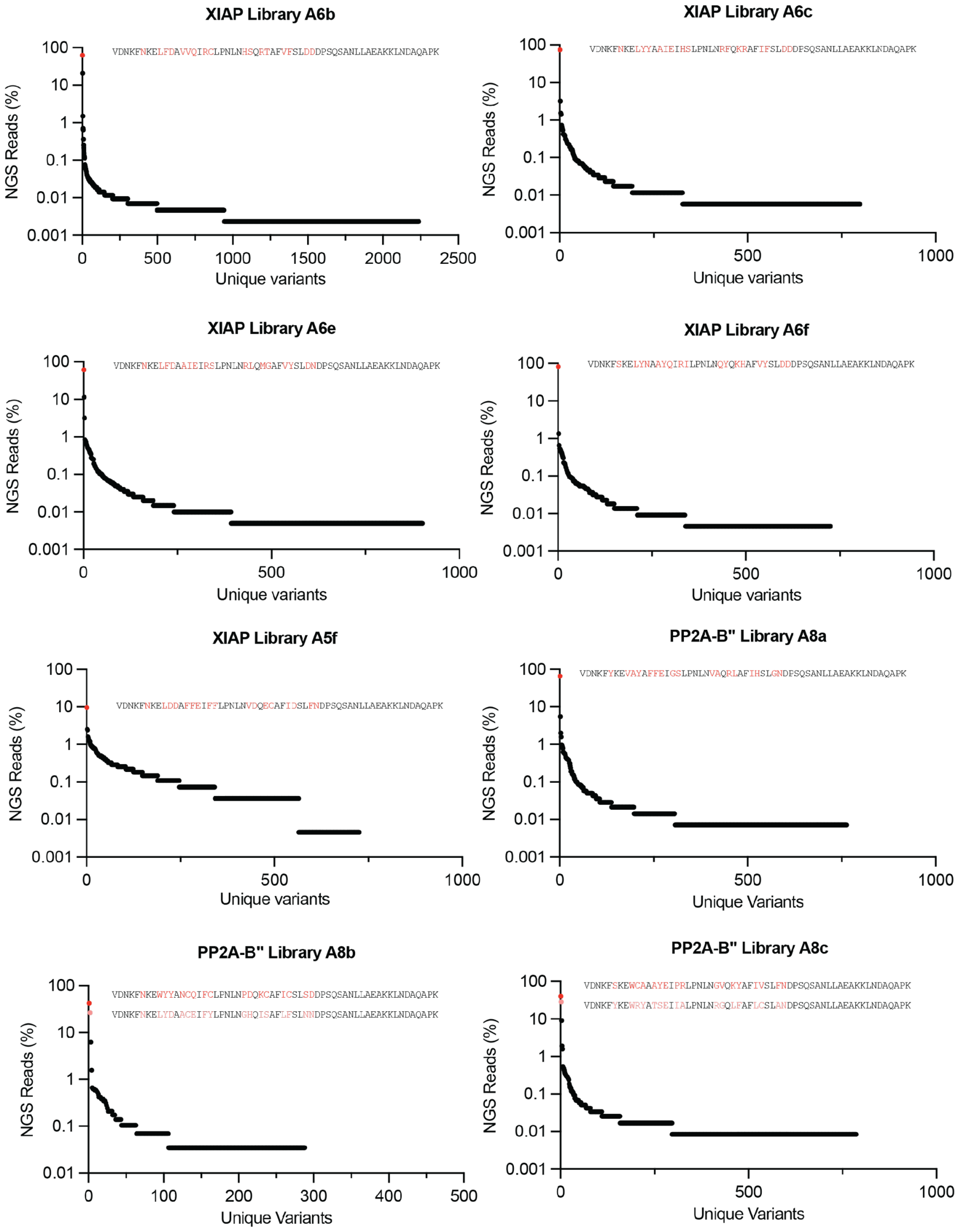
XIAP Library A6b
VDNKFNKELFDAVVQIRCLPNLNHSQRTAFVFSLDDDPSQSANLLAEAKKLNDAQAPK
NGS Reads (%)
Unique variants
XIAP Library A6c
VDNKFNKELYYAAIEIHSLPNLNRFQKRAFIFSLDDDPSQSANLLAEAKKLNDAQAPK
NGS Reads (%)
Unique variants
XIAP Library A6e
VDNKFNKELFDAAIEIRSLPNLNRLQMGAFVYSLDNDPSQSANLLAEAKKLNDAQAPK
NGS Reads (%)
Unique variants
XIAP Library A6f
VDNKFSKELYNAAYQIRILPNLNQYQKHAFVYSLDDDPSQSANLLAEAKKLNDAQAPK
NGS Reads (%)
Unique variants
XIAP Library A5f
VDNKFNKELDDAFFEIFFLPNLNVDQECAFIDSLFNDPSQSANLLAEAKKLNDAQAPK
NGS Reads (%)
Unique variants
PP2A-B" Library A8a
VDNKFYKEVAYAFFEIGSLPNLNVAQRLAFIHSLGNDPSQSANLLAEAKKLNDAQAPK
NGS Reads (%)
Unique Variants
PP2A-B" Library A8b
VDNKFNKEWYYANCQIFCLPNLNPDQKCAFICSLSDDPSQSANLLAEAKKLNDAQAPK
VDNKFNKELYDAACEIFYLPNLNGHQISAFLFSLNNDPSQSANLLAEAKKLNDAQAPK
NGS Reads (%)
Unique Variants
PP2A-B" Library A8c
VDNKFSKEWCAAAYEIPRLPNLNGVQKYAFIVSLFNDPSQSANLLAEAKKLNDAQAPK
VDNKFYKEWRYATSEIIALPNLNRGQLFAFLCSLANDPSQSANLLAEAKKLNDAQAPK
NGS Reads (%)
Unique Variants

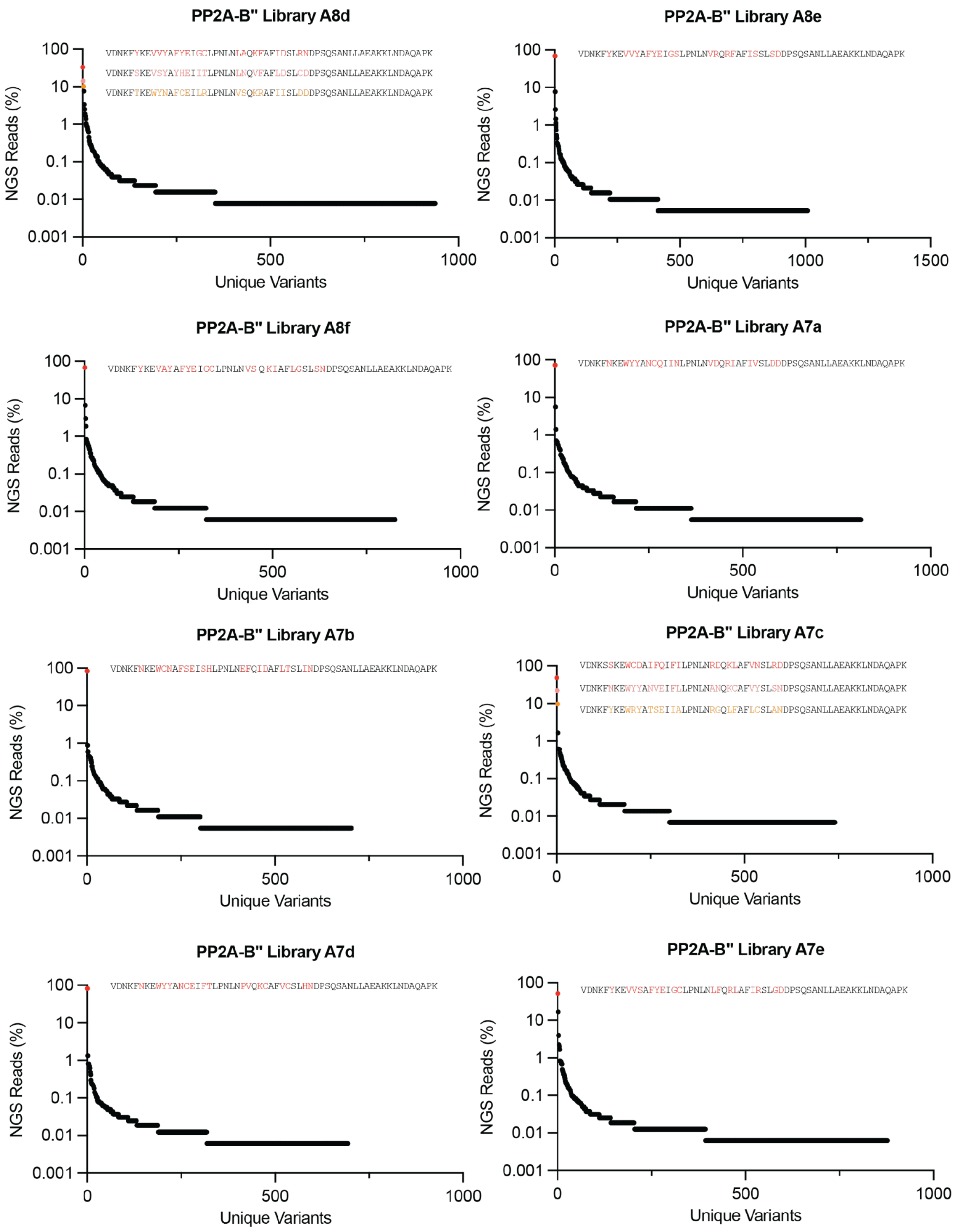
PP2A-B" Library A8d
VDNKFYKEVVYAFYEIGCLPNLNLAQKFAFTDSLRNDPSQSANLLAEAKKLNDAQAPK
VDNKFSKEVSYAYHEIITLPNLNLNQVFAFLDSLCDDPSQSANLLAEAKKLNDAQAPK
VDNKFTKEWYNAFCEILRLPNLNVSQKRAFIISLDDDPSQSANLLAEAKKLNDAQAPK
PP2A-B" Library A8e
VDNKFYKEVVYAFYEIGSLPNLNVRQRFAFISSLSDDPSQSANLLAEAKKLNDAQAPK
PP2A-B" Library A8f
VDNKFYKEVAYAFYEICCLPNLNVS Q KIAFLCSLSNDPSQSANLLAEAKKLNDAQAPK
PP2A-B" Library A7a
VDNKFNKEWYYANCQIINLPNLNVDQRIAFIVSLDDDPSQSANLLAEAKKLNDAQAPK
PP2A-B" Library A7b
VDNKFNKEWCNAFSEISHLPNLNEFQIDAFLTSLINDPSQSANLLAEAKKLNDAQAPK
PP2A-B" Library A7c
VDNKSSKEWCDAIFQIFILPNLNRDQKLAFVNSLRDDPSQSANLLAEAKKLNDAQAPK
VDNKFNKEWYYANVEIFLLPNLNANQKCAFVYSLSNDPSQSANLLAEAKKLNDAQAPK
VDNKFYKEWRYATSEIIALPNLNRGQLFAFLCSLANDPSQSANLLAEAKKLNDAQAPK
PP2A-B" Library A7d
VDNKFNKEWYYANCEIFTLPNLNPVQKCAFVCSLHNDPSQSANLLAEAKKLNDAQAPK
PP2A-B" Library A7e
VDNKFYKEVVSAFYEIGCLPNLNLFQRLAFIRSLGDDPSQSANLLAEAKKLNDAQAPK
NGS Reads (%)
100
10
1
0.1
0.01
0.001
Unique Variants
0
500
1000
1500

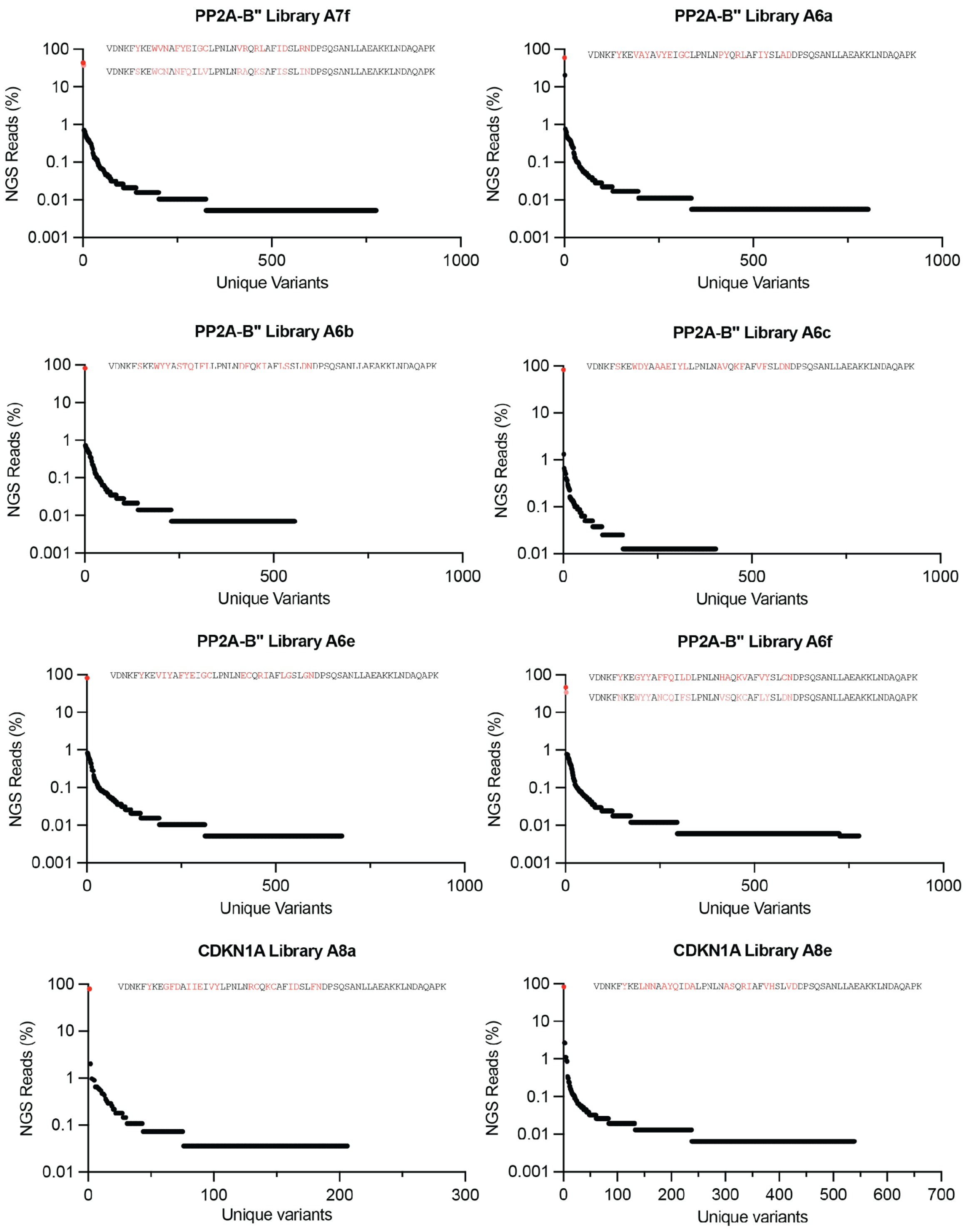
PP2A-B" Library A7f
VDNKFYKEWVNAFYEIGCLPNLNVRQRLAFIDSLRNDPSQSANLLAEAKKLNDAQAPK
VDNKFSKEWCNANFQILVLPNLNRAQKSAFISSLINDPSQSANLLAEAKKLNDAQAPK
NGS Reads (%)
100
10
1
0.1
0.01
0.001
0
500
1000
Unique Variants
PP2A-B" Library A6a
VDNKFYKEVAYAVYEIGCLPNLNPYQRLAFIYSLADDPSQSANLLAEAKKLNDAQAPK
NGS Reads (%)
Unique Variants
PP2A-B" Library A6b
VDNKFSKEWYYASTQIFLLPNLNDFQKTAFLSSLDNDPSQSANLLAEAKKLNDAQAPK
NGS Reads (%)
Unique Variants
PP2A-B" Library A6c
VDNKFSKEWDYAAAEIYLLPNLNAVQKFAFVFSLDNDPSQSANLLAEAKKLNDAQAPK
NGS Reads (%)
Unique Variants
PP2A-B" Library A6e
VDNKFYKEVIYAFYEIGCLPNLNECQRIAFLGSLGNDPSQSANLLAEAKKLNDAQAPK
NGS Reads (%)
Unique Variants
PP2A-B" Library A6f
VDNKFYKEGYYAFFQILDLPNLNHAQKVAFVYSLCNDPSQSANLLAEAKKLNDAQAPK
VDNKFNKEWYYANCQIFSLPNLNVSQKCAFLYSLDNDPSQSANLLAEAKKLNDAQAPK
NGS Reads (%)
Unique Variants
CDKN1A Library A8a
VDNKFYKEGFDAIIEIVYLPNLNRCQKCAFIDSLFNDPSQSANLLAEAKKLNDAQAPK
NGS Reads (%)
100
200
300
Unique variants
CDKN1A Library A8e
VDNKFYKELNNAAYQIDALPNLNASQRIAFVHSLVDDPSQSANLLAEAKKLNDAQAPK
NGS Reads (%)
400
600
700
Unique variants

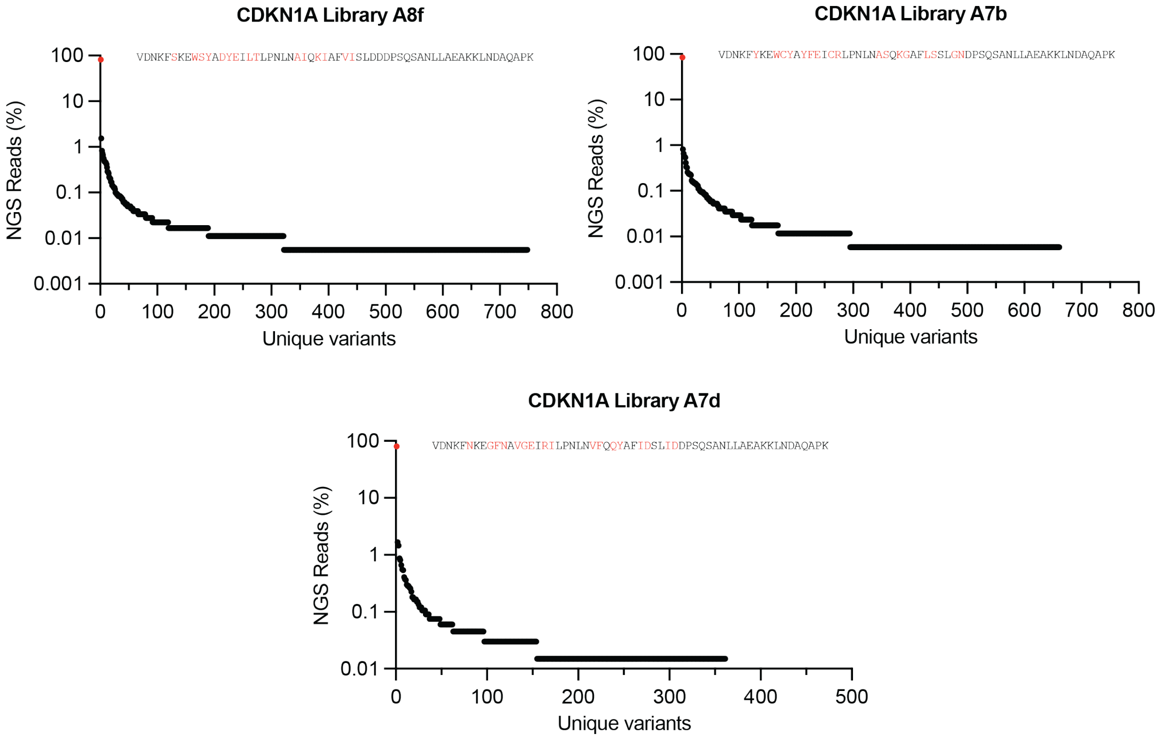

**Supplementary Fig. 3: NGS sequence prevalence plots for hits from 9-target panel.** After separating the reads by barcode and translating DNA sequences into amino acid sequences, the abundance of each unique sequence meeting the randomization design in the NGS results from each selection with >$10^7$ PFU/mL endpoint titer (**Supplementary Fig. 2**) was plotted as a percentage of total reads for that selection. The top variant (for nearly all) selections was subcloned from the endpoint phage for further testing in the split T7 RNAP *E. coli* luciferase assay (**Supplementary Fig. 4**).

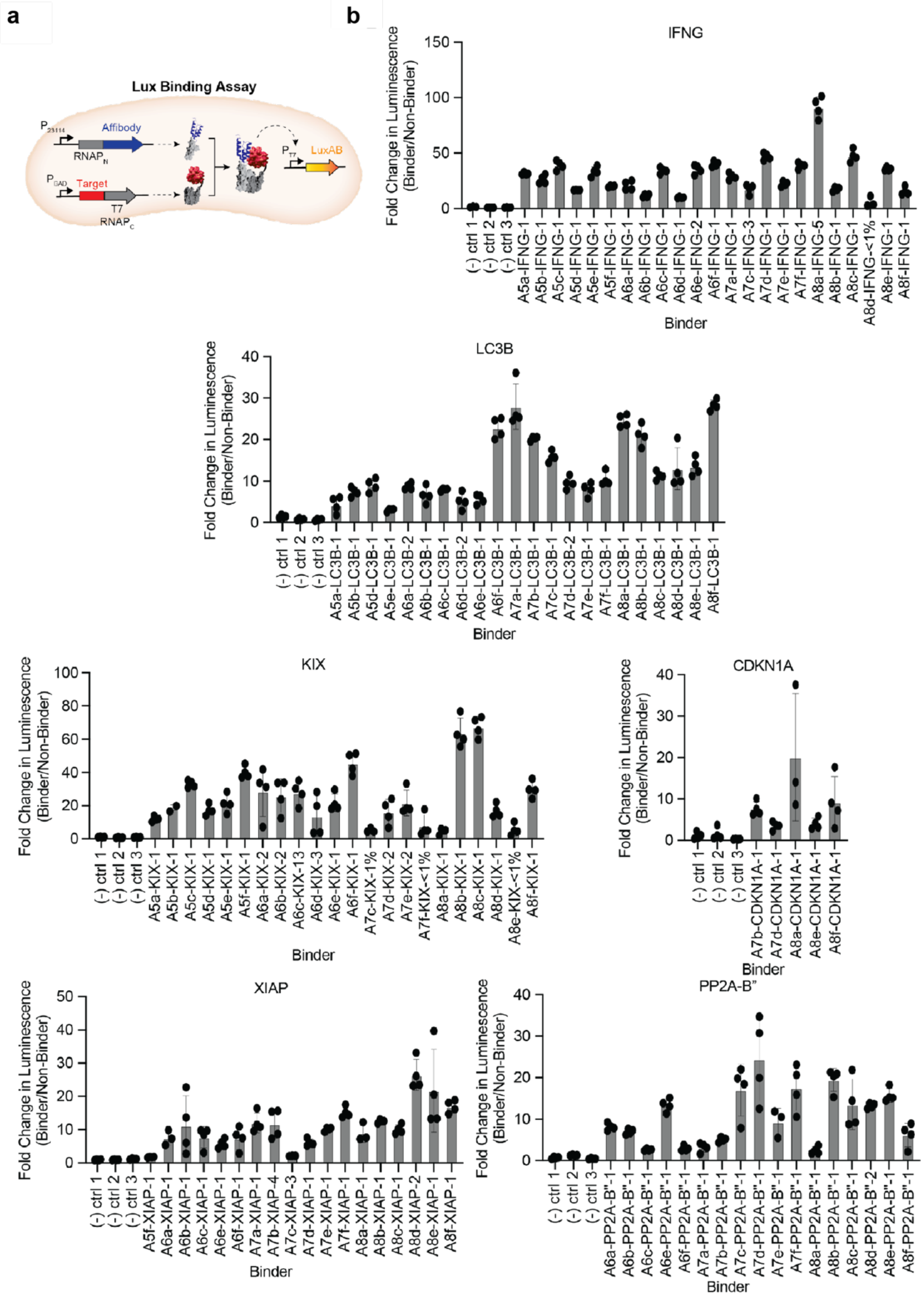

a
b
Lux Binding Assay
Affibody
RNAP
Target
T7
RNAP
LuxAB
IFNG
LC3B
KIX
CDKN1A
XIAP
PP2A-B”
Fold Change in Luminescence (Binder/Non-Binder)
Binder
(-) ctrl 1
(-) ctrl 2
(-) ctrl 3
A5a-IFNG-1
A5b-IFNG-1
A5c-IFNG-1
A5d-IFNG-1
A5e-IFNG-1
A5f-IFNG-1
A6a-IFNG-1
A6b-IFNG-1
A6c-IFNG-1
A6d-IFNG-1
A6e-IFNG-2
A6f-IFNG-1
A7a-IFNG-1
A7c-IFNG-3
A7d-IFNG-1
A7e-IFNG-1
A7f-IFNG-1
A8a-IFNG-5
A8b-IFNG-1
A8c-IFNG-1
A8d-IFNG-<1%
A8e-IFNG-1
A8f-IFNG-1
A5a-LC3B-1
A5b-LC3B-1
A5d-LC3B-1
A5e-LC3B-1
A6a-LC3B-2
A6b-LC3B-1
A6c-LC3B-1
A6d-LC3B-2
A6e-LC3B-1
A6f-LC3B-1
A7a-LC3B-1
A7b-LC3B-1
A7c-LC3B-1
A7d-LC3B-2
A7e-LC3B-1
A7f-LC3B-1
A8a-LC3B-1
A8b-LC3B-1
A8c-LC3B-1
A8d-LC3B-1
A8e-LC3B-1
A8f-LC3B-1
A5a-KIX-1
A5b-KIX-1
A5c-KIX-1
A5d-KIX-1
A5e-KIX-1
A5f-KIX-1
A6a-KIX-2
A6b-KIX-2
A6c-KIX-13
A6d-KIX-3
A6e-KIX-1
A6f-KIX-1
A7c-KIX-1%
A7d-KIX-2
A7e-KIX-2
A7f-KIX-<1%
A8a-KIX-1
A8b-KIX-1
A8c-KIX-1
A8d-KIX-1
A8e-KIX-<1%
A8f-KIX-1
A7b-CDKN1A-1
A7d-CDKN1A-1
A8a-CDKN1A-1
A8e-CDKN1A-1
A8f-CDKN1A-1
A5f-XIAP-1
A6a-XIAP-1
A6b-XIAP-1
A6c-XIAP-1
A6e-XIAP-1
A6f-XIAP-1
A7a-XIAP-1
A7b-XIAP-4
A7c-XIAP-3
A7d-XIAP-1
A7e-XIAP-1
A7f-XIAP-1
A8a-XIAP-1
A8b-XIAP-1
A8c-XIAP-1
A8d-XIAP-2
A8e-XIAP-1
A8f-XIAP-1
A6a-PP2A-B"-1
A6b-PP2A-B"-1
A6c-PP2A-B"-1
A6e-PP2A-B"-1
A6f-PP2A-B"-1
A7a-PP2A-B"-1
A7b-PP2A-B"-1
A7c-PP2A-B"-1
A7d-PP2A-B"-1
A7e-PP2A-B"-1
A7f-PP2A-B"-1
A8a-PP2A-B"-1
A8b-PP2A-B"-1
A8c-PP2A-B"-1
A8d-PP2A-B"-2
A8e-PP2A-B"-1
A8f-PP2A-B"-1

**c**

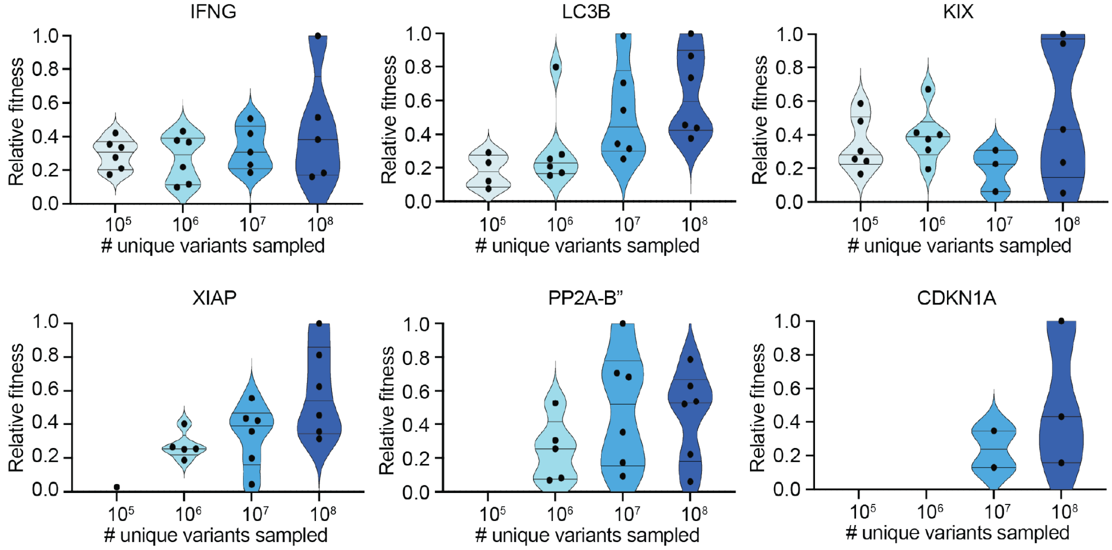

**Supplementary Fig. 4: Split T7 RNAP *E. coli* luciferase assay to assess binding to targets (fitness). a**, Schematic of Split T7 RNAP *E. coli* binding luciferase assay. The affibody variant from the phage is subcloned onto a plasmid as a fusion with $RNAP_N$, the target is cloned on an inducible plasmid fused to $RNAP_C$, and complementation of RNAP results in expression of LuxAB. **b**, Luciferase assay results for each target-binder pair with target-non-binder controls (affibody (PD-L1)[2], Monobody (hSUMO1)[3], and affitin (SasA)[4]). To calculate the fold-change of the binder over the non-targeting controls, the blank $OD_{600}$ signal (media only) was subtracted from the $OD_{600}$ of each sample, then the luminescence signal of each binder was divided by this modified $OD_{600}$. The average of all three negative, non-binding controls was calculated for each target, then the fold-change of each binder colony was divided by this non-binding control average for each target to produce the fold-change of the binder over the non-targeting controls. Each bar plot shows the mean, standard deviation, and individual values ($n$ = 4, error bars indicate SD). **c**, Comparison between the relative fitness of binders discovered from each sampling size. To calculate the relative fitness of binders, the fold-changes of all binders for a target were normalized to the highest luciferase signal binder for that target (across all sampling sizes).

**Supplementary Table 2: Affibody binder densities identified for the 9-target panel.**

| **Binder density** | **Targets in this category** |
|---|---|
| $>10^{-5}$ | IFNG, LC3B, KIX |
| $>10^{-6}$ | XIAP, PP2A-B” |
| $>10^{-7}$ | CDKN1A |
| $<10^{-8}$ | PP2A-B, BTK (PH), MAX |

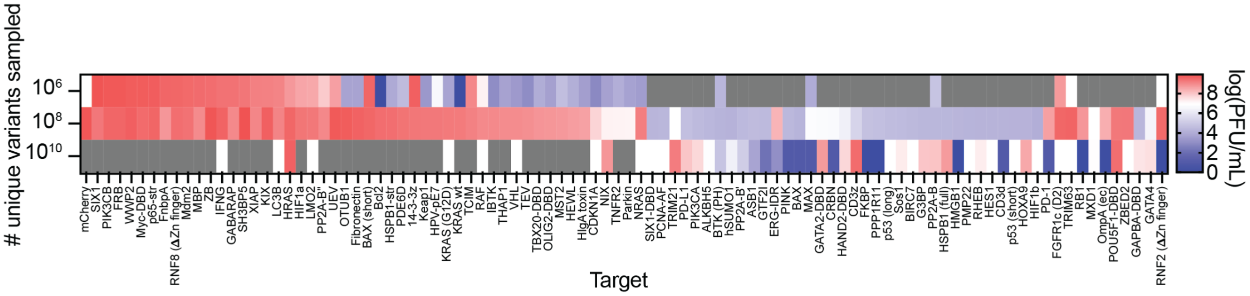

**Supplementary Fig. 5: Heat map of titers following PANCS of the 96-target panel with design A at $10^6$, $10^8$, $10^{10}$.** PANCS was performed on the 96-target panel (Supplementary Table 1) with a single independent replicates of Library design A with varied sampling sizes ($10^{6,\ 8,\ \text{or}\ 10}$). After completion of the sixth passage, the titer was determined by qPCR. Red indicates higher titers, blue lower titers, and gray indicates that a selection was not performed for a target at that sample size. All selections with a titer >1 × $10^7$ PFU/mL were subject to NGS analysis and validation in the split T7 RNAP *E. coli* luciferase assay. Each titer was determined as a single replicate.

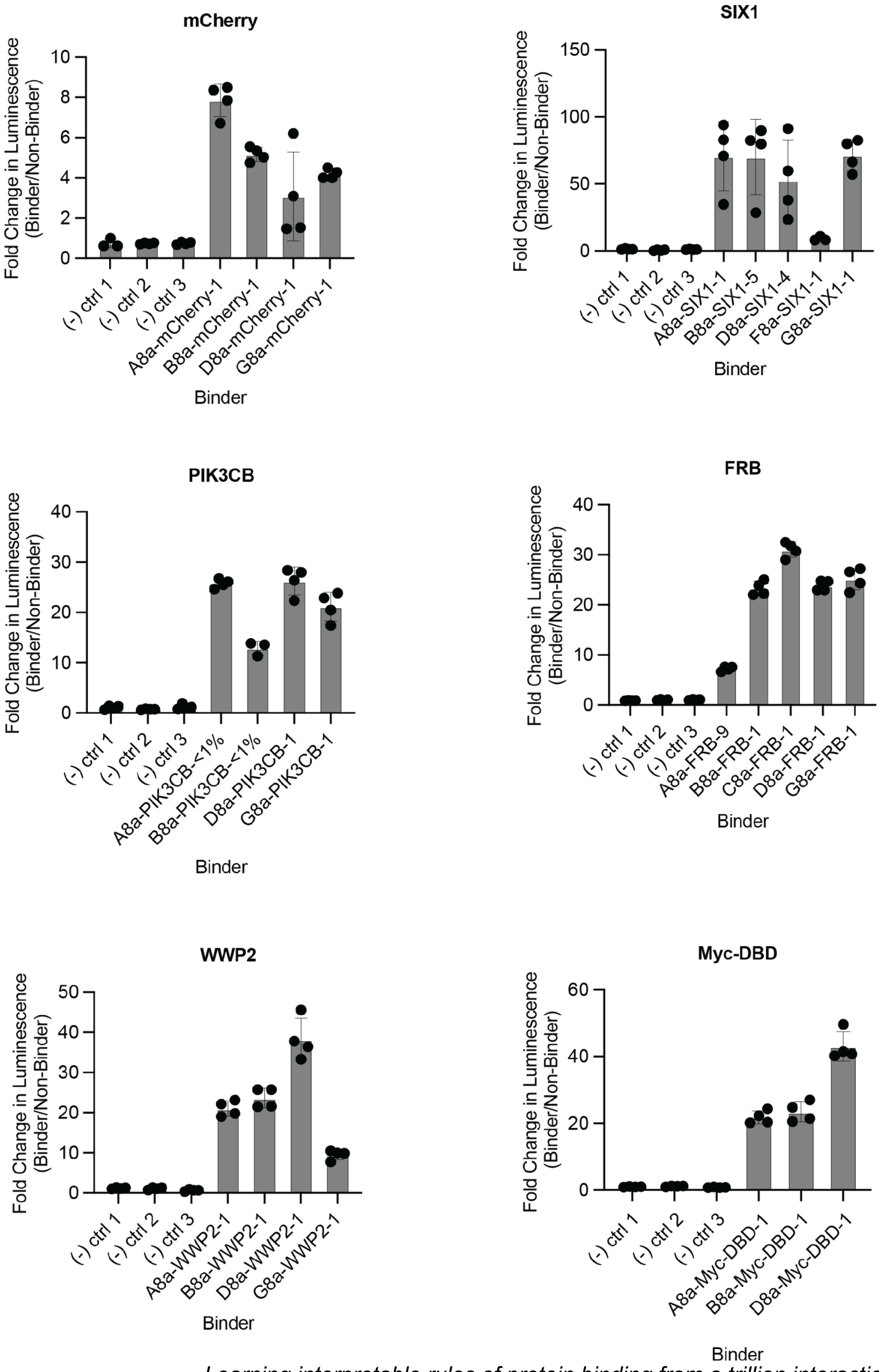
mCherry
Fold Change in Luminescence (Binder/Non-Binder)
0
2
4
6
8
10
(-) ctrl 1
(-) ctrl 2
(-) ctrl 3
A8a-mCherry-1
B8a-mCherry-1
D8a-mCherry-1
G8a-mCherry-1
Binder
SIX1
Fold Change in Luminescence (Binder/Non-Binder)
0
50
100
150
(-) ctrl 1
(-) ctrl 2
(-) ctrl 3
A8a-SIX1-1
B8a-SIX1-5
D8a-SIX1-4
F8a-SIX1-1
G8a-SIX1-1
Binder
PIK3CB
Fold Change in Luminescence (Binder/Non-Binder)
0
10
20
30
40
(-) ctrl 1
(-) ctrl 2
(-) ctrl 3
A8a-PIK3CB-<1%
B8a-PIK3CB-<1%
D8a-PIK3CB-1
G8a-PIK3CB-1
Binder
FRB
Fold Change in Luminescence (Binder/Non-Binder)
0
10
20
30
40
(-) ctrl 1
(-) ctrl 2
(-) ctrl 3
A8a-FRB-9
B8a-FRB-1
C8a-FRB-1
D8a-FRB-1
G8a-FRB-1
Binder
WWP2
Fold Change in Luminescence (Binder/Non-Binder)
0
10
20
30
40
50
(-) ctrl 1
(-) ctrl 2
(-) ctrl 3
A8a-WWP2-1
B8a-WWP2-1
D8a-WWP2-1
G8a-WWP2-1
Binder
Myc-DBD
Fold Change in Luminescence (Binder/Non-Binder)
0
20
40
60
(-) ctrl 1
(-) ctrl 2
(-) ctrl 3
A8a-Myc-DBD-1
B8a-Myc-DBD-1
D8a-Myc-DBD-1
Binder

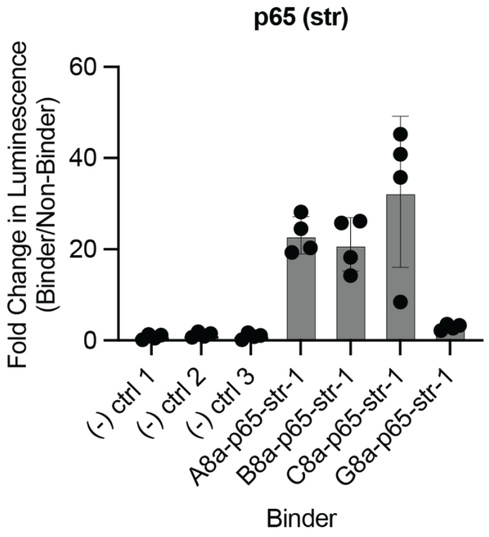
p65 (str)
Fold Change in Luminescence (Binder/Non-Binder)
0
20
40
60
(-) ctrl 1
(-) ctrl 2
(-) ctrl 3
A8a-p65-str-1
B8a-p65-str-1
C8a-p65-str-1
G8a-p65-str-1
Binder

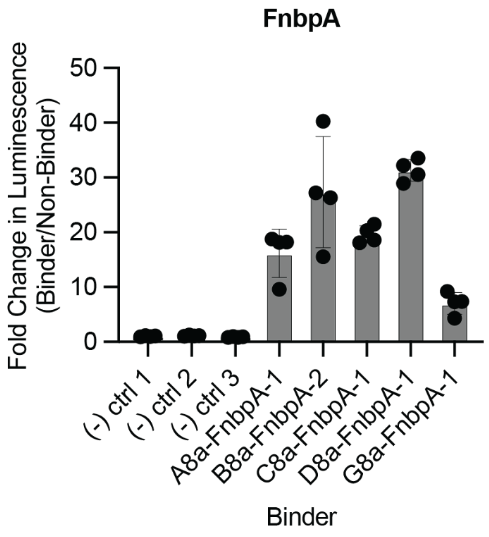
FnbpA
Fold Change in Luminescence (Binder/Non-Binder)
0
10
20
30
40
50
(-) ctrl 1
(-) ctrl 2
(-) ctrl 3
A8a-FnbpA-1
B8a-FnbpA-2
C8a-FnbpA-1
D8a-FnbpA-1
G8a-FnbpA-1
Binder

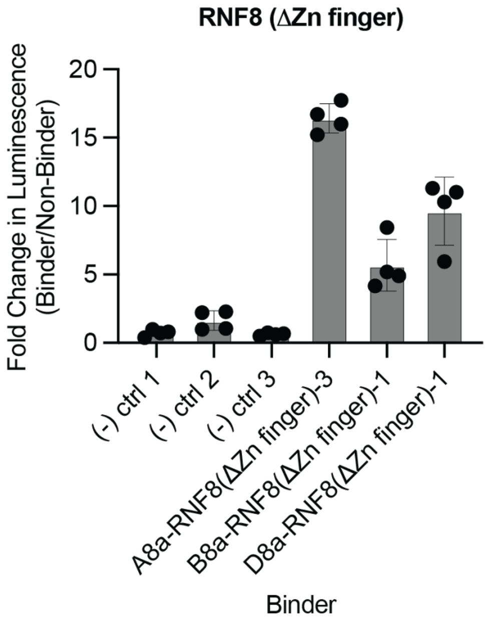
RNF8 (ΔZn finger)
Fold Change in Luminescence (Binder/Non-Binder)
0
5
10
15
20
(-) ctrl 1
(-) ctrl 2
(-) ctrl 3
A8a-RNF8(ΔZn finger)-3
B8a-RNF8(ΔZn finger)-1
D8a-RNF8(ΔZn finger)-1
Binder

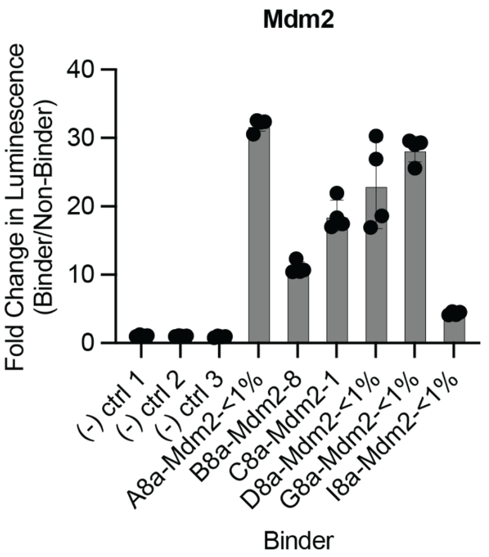
Mdm2
Fold Change in Luminescence (Binder/Non-Binder)
0
10
20
30
40
(-) ctrl 1
(-) ctrl 2
(-) ctrl 3
A8a-Mdm2-<1%
B8a-Mdm2-8
C8a-Mdm2-1
D8a-Mdm2-<1%
G8a-Mdm2-<1%
I8a-Mdm2-<1%
Binder

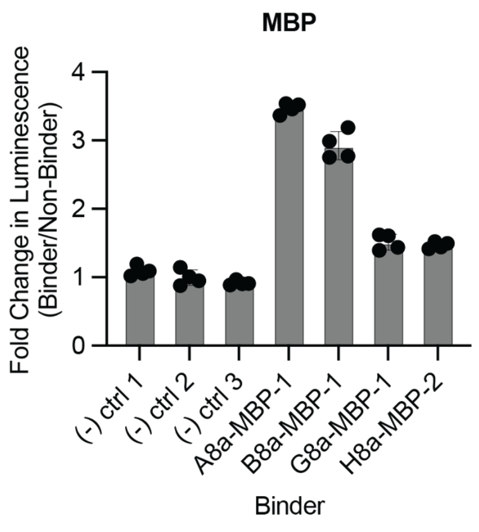
MBP
Fold Change in Luminescence (Binder/Non-Binder)
0
1
2
3
4
(-) ctrl 1
(-) ctrl 2
(-) ctrl 3
A8a-MBP-1
B8a-MBP-1
G8a-MBP-1
H8a-MBP-2
Binder

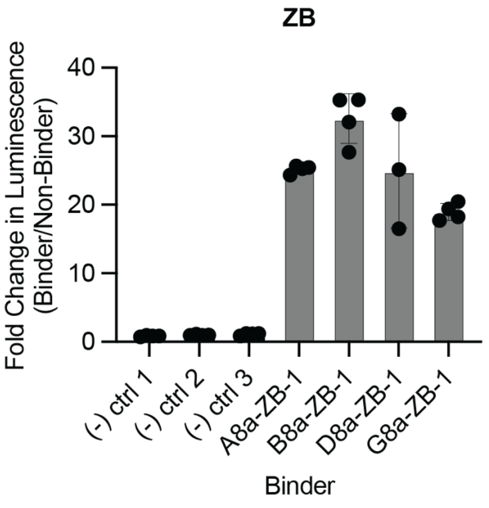
ZB
Fold Change in Luminescence (Binder/Non-Binder)
0
10
20
30
40
(-) ctrl 1
(-) ctrl 2
(-) ctrl 3
A8a-ZB-1
B8a-ZB-1
D8a-ZB-1
G8a-ZB-1
Binder

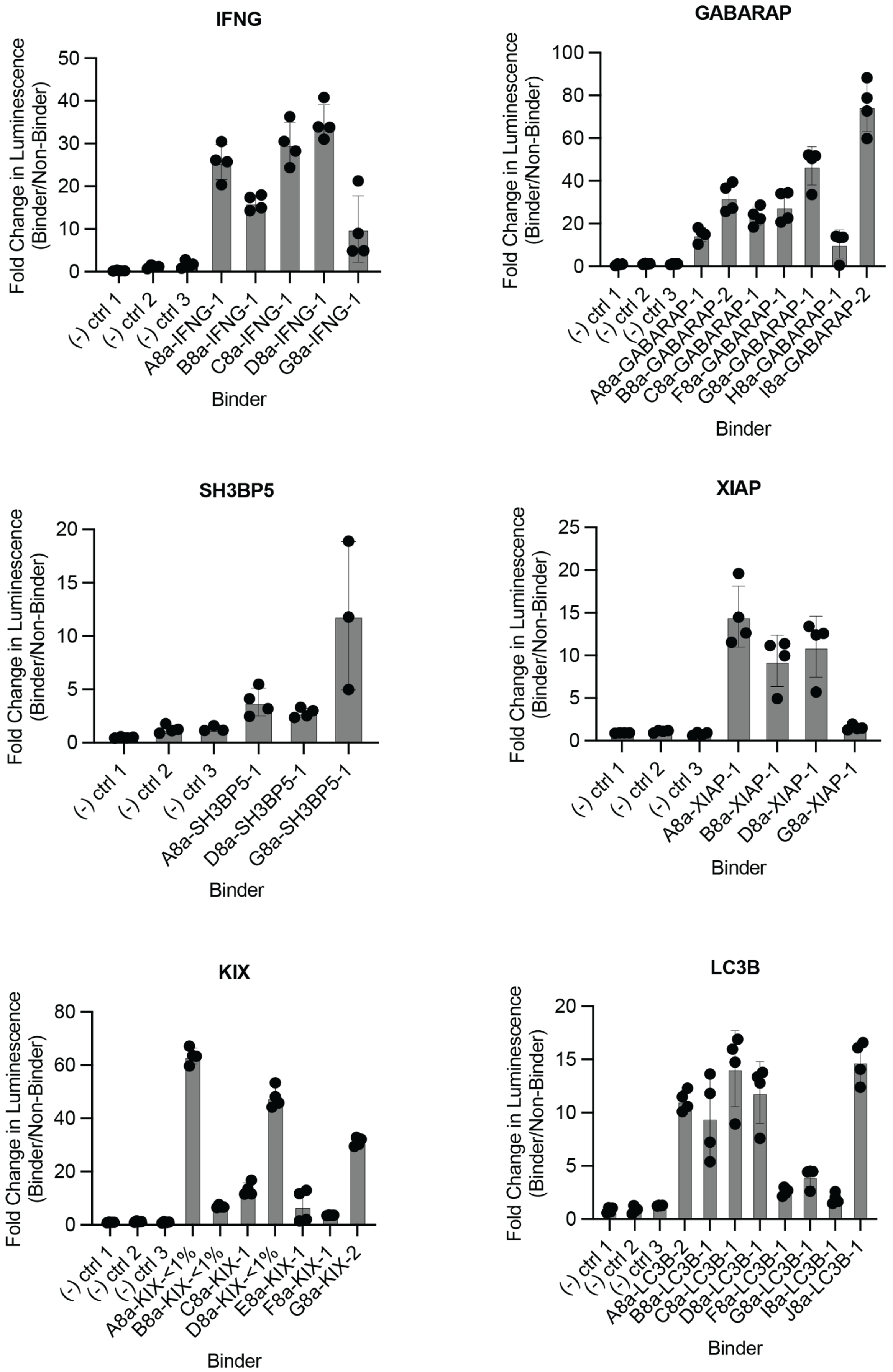
IFNG
Fold Change in Luminescence (Binder/Non-Binder)
0
10
20
30
40
50
(-) ctrl 1
(-) ctrl 2
(-) ctrl 3
A8a-IFNG-1
B8a-IFNG-1
C8a-IFNG-1
D8a-IFNG-1
G8a-IFNG-1
Binder
GABARAP
Fold Change in Luminescence (Binder/Non-Binder)
0
20
40
60
80
100
(-) ctrl 1
(-) ctrl 2
(-) ctrl 3
A8a-GABARAP-1
B8a-GABARAP-2
C8a-GABARAP-1
F8a-GABARAP-1
G8a-GABARAP-1
H8a-GABARAP-1
I8a-GABARAP-2
Binder
SH3BP5
Fold Change in Luminescence (Binder/Non-Binder)
0
5
10
15
20
(-) ctrl 1
(-) ctrl 2
(-) ctrl 3
A8a-SH3BP5-1
D8a-SH3BP5-1
G8a-SH3BP5-1
Binder
XIAP
Fold Change in Luminescence (Binder/Non-Binder)
0
5
10
15
20
25
(-) ctrl 1
(-) ctrl 2
(-) ctrl 3
A8a-XIAP-1
B8a-XIAP-1
D8a-XIAP-1
G8a-XIAP-1
Binder
KIX
Fold Change in Luminescence (Binder/Non-Binder)
0
20
40
60
80
(-) ctrl 1
(-) ctrl 2
(-) ctrl 3
A8a-KIX-<1%
B8a-KIX-<1%
C8a-KIX-1
D8a-KIX-<1%
E8a-KIX-1
F8a-KIX-1
G8a-KIX-2
Binder
LC3B
Fold Change in Luminescence (Binder/Non-Binder)
0
5
10
15
20
(-) ctrl 1
(-) ctrl 2
(-) ctrl 3
A8a-LC3B-2
B8a-LC3B-1
C8a-LC3B-1
D8a-LC3B-1
F8a-LC3B-1
G8a-LC3B-1
I8a-LC3B-1
J8a-LC3B-1
Binder

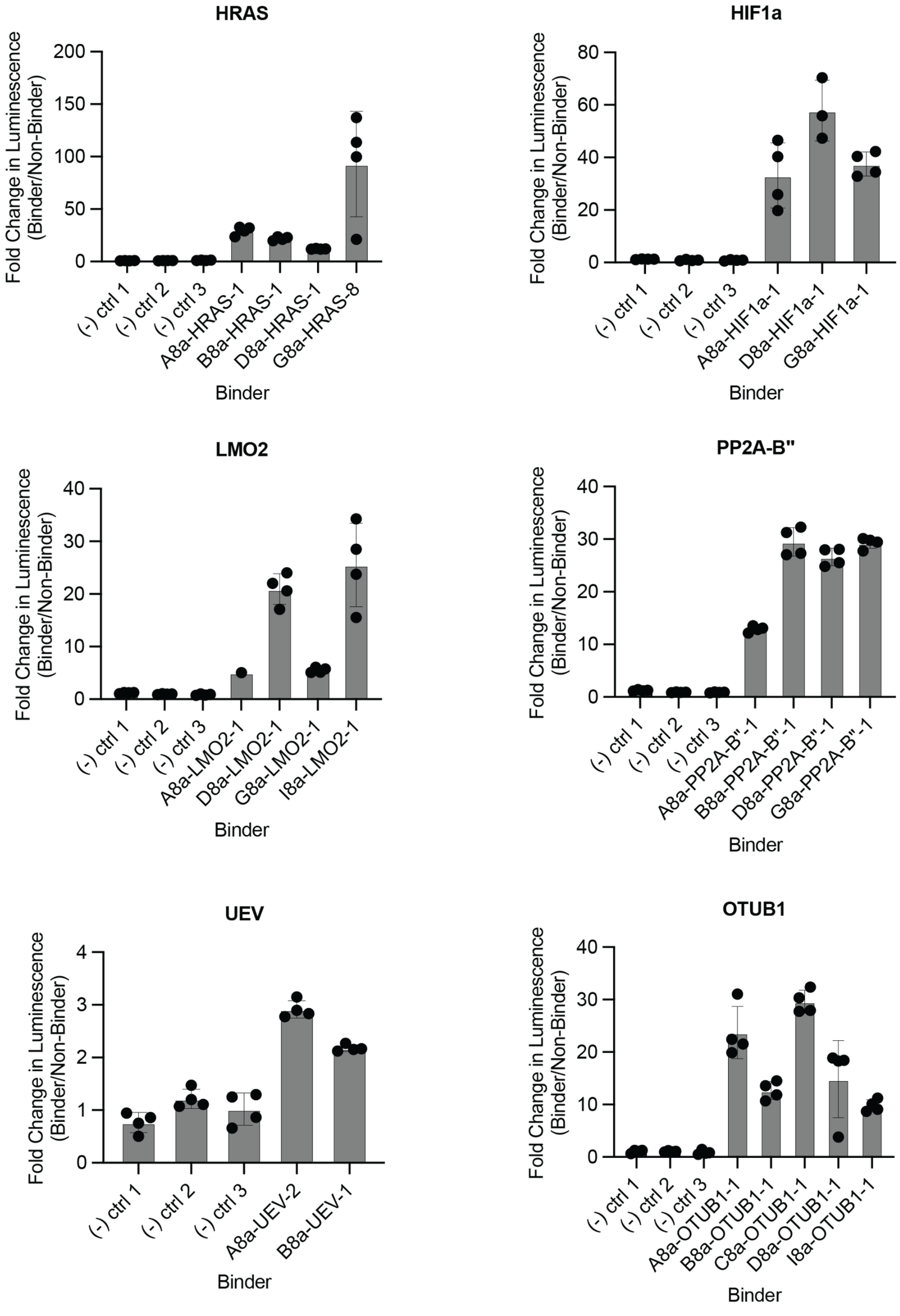
HRAS
Fold Change in Luminescence (Binder/Non-Binder)
0
50
100
150
200
(-) ctrl 1
(-) ctrl 2
(-) ctrl 3
A8a-HRAS-1
B8a-HRAS-1
D8a-HRAS-1
G8a-HRAS-8
Binder
HIF1a
Fold Change in Luminescence (Binder/Non-Binder)
0
20
40
60
80
(-) ctrl 1
(-) ctrl 2
(-) ctrl 3
A8a-HIF1a-1
D8a-HIF1a-1
G8a-HIF1a-1
Binder
LMO2
Fold Change in Luminescence (Binder/Non-Binder)
0
10
20
30
40
(-) ctrl 1
(-) ctrl 2
(-) ctrl 3
A8a-LMO2-1
D8a-LMO2-1
G8a-LMO2-1
I8a-LMO2-1
Binder
PP2A-B''
Fold Change in Luminescence (Binder/Non-Binder)
0
10
20
30
40
(-) ctrl 1
(-) ctrl 2
(-) ctrl 3
A8a-PP2A-B''-1
B8a-PP2A-B''-1
D8a-PP2A-B''-1
G8a-PP2A-B''-1
Binder
UEV
Fold Change in Luminescence (Binder/Non-Binder)
0
1
2
3
4
(-) ctrl 1
(-) ctrl 2
(-) ctrl 3
A8a-UEV-2
B8a-UEV-1
Binder
OTUB1
Fold Change in Luminescence (Binder/Non-Binder)
0
10
20
30
40
(-) ctrl 1
(-) ctrl 2
(-) ctrl 3
A8a-OTUB1-1
B8a-OTUB1-1
C8a-OTUB1-1
D8a-OTUB1-1
I8a-OTUB1-1
Binder

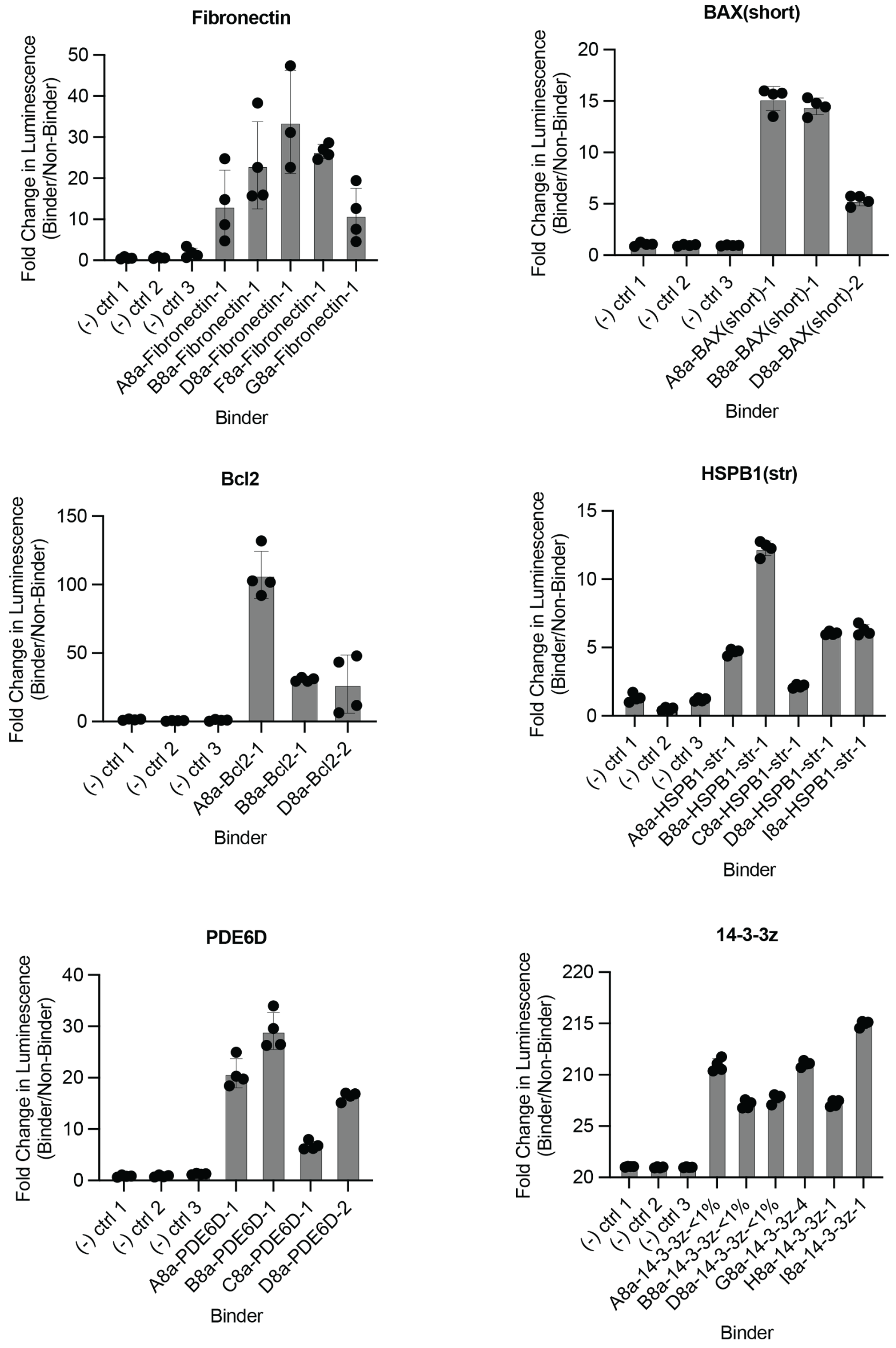
Fibronectin
Fold Change in Luminescence (Binder/Non-Binder)
50
40
30
20
10
0
(-) ctrl 1
(-) ctrl 2
(-) ctrl 3
A8a-Fibronectin-1
B8a-Fibronectin-1
D8a-Fibronectin-1
F8a-Fibronectin-1
G8a-Fibronectin-1
Binder
BAX(short)
Fold Change in Luminescence (Binder/Non-Binder)
20
15
10
5
0
(-) ctrl 1
(-) ctrl 2
(-) ctrl 3
A8a-BAX(short)-1
B8a-BAX(short)-1
D8a-BAX(short)-2
Binder
Bcl2
Fold Change in Luminescence (Binder/Non-Binder)
150
100
50
0
(-) ctrl 1
(-) ctrl 2
(-) ctrl 3
A8a-Bcl2-1
B8a-Bcl2-1
D8a-Bcl2-2
Binder
HSPB1(str)
Fold Change in Luminescence (Binder/Non-Binder)
15
10
5
0
(-) ctrl 1
(-) ctrl 2
(-) ctrl 3
A8a-HSPB1-str-1
B8a-HSPB1-str-1
C8a-HSPB1-str-1
D8a-HSPB1-str-1
I8a-HSPB1-str-1
Binder
PDE6D
Fold Change in Luminescence (Binder/Non-Binder)
40
30
20
10
0
(-) ctrl 1
(-) ctrl 2
(-) ctrl 3
A8a-PDE6D-1
B8a-PDE6D-1
C8a-PDE6D-1
D8a-PDE6D-2
Binder
14-3-3z
Fold Change in Luminescence (Binder/Non-Binder)
220
215
210
25
20
(-) ctrl 1
(-) ctrl 2
(-) ctrl 3
A8a-14-3-3z-<1%
B8a-14-3-3z-<1%
D8a-14-3-3z-<1%
G8a-14-3-3z-4
H8a-14-3-3z-1
I8a-14-3-3z-1
Binder

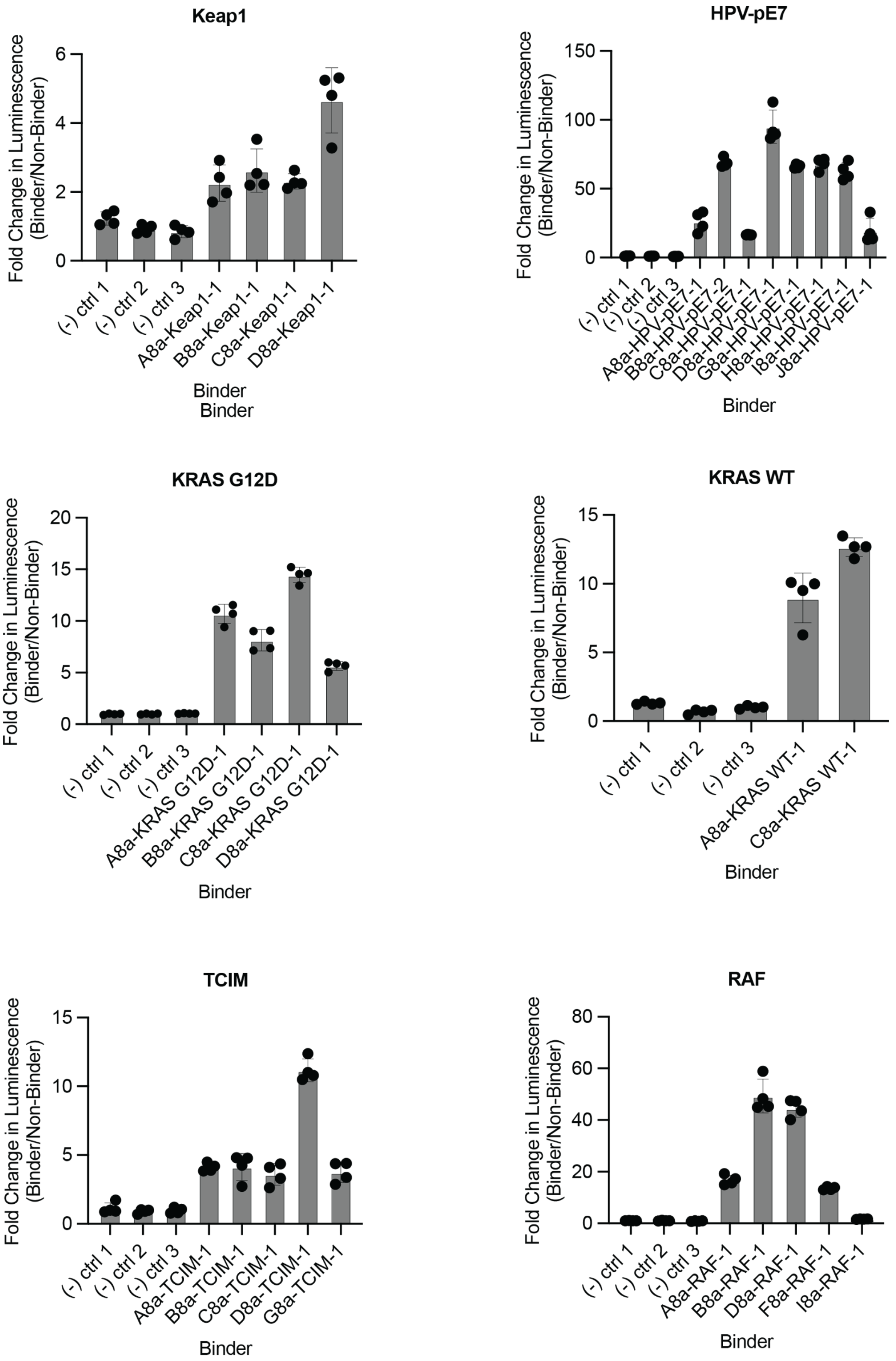
Keap1
Fold Change in Luminescence (Binder/Non-Binder)
0
2
4
6
(-) ctrl 1
(-) ctrl 2
(-) ctrl 3
A8a-Keap1-1
B8a-Keap1-1
C8a-Keap1-1
D8a-Keap1-1
Binder
Binder
HPV-pE7
0
50
100
150
(-) ctrl 1
(-) ctrl 2
(-) ctrl 3
A8a-HPV-pE7-1
B8a-HPV-pE7-2
C8a-HPV-pE7-1
D8a-HPV-pE7-1
G8a-HPV-pE7-1
H8a-HPV-pE7-1
I8a-HPV-pE7-1
J8a-HPV-pE7-1
KRAS G12D
0
5
10
15
20
A8a-KRAS G12D-1
B8a-KRAS G12D-1
C8a-KRAS G12D-1
D8a-KRAS G12D-1
KRAS WT
A8a-KRAS WT-1
C8a-KRAS WT-1
TCIM
A8a-TCIM-1
B8a-TCIM-1
C8a-TCIM-1
D8a-TCIM-1
G8a-TCIM-1
RAF
0
20
40
60
80
A8a-RAF-1
B8a-RAF-1
D8a-RAF-1
F8a-RAF-1
I8a-RAF-1

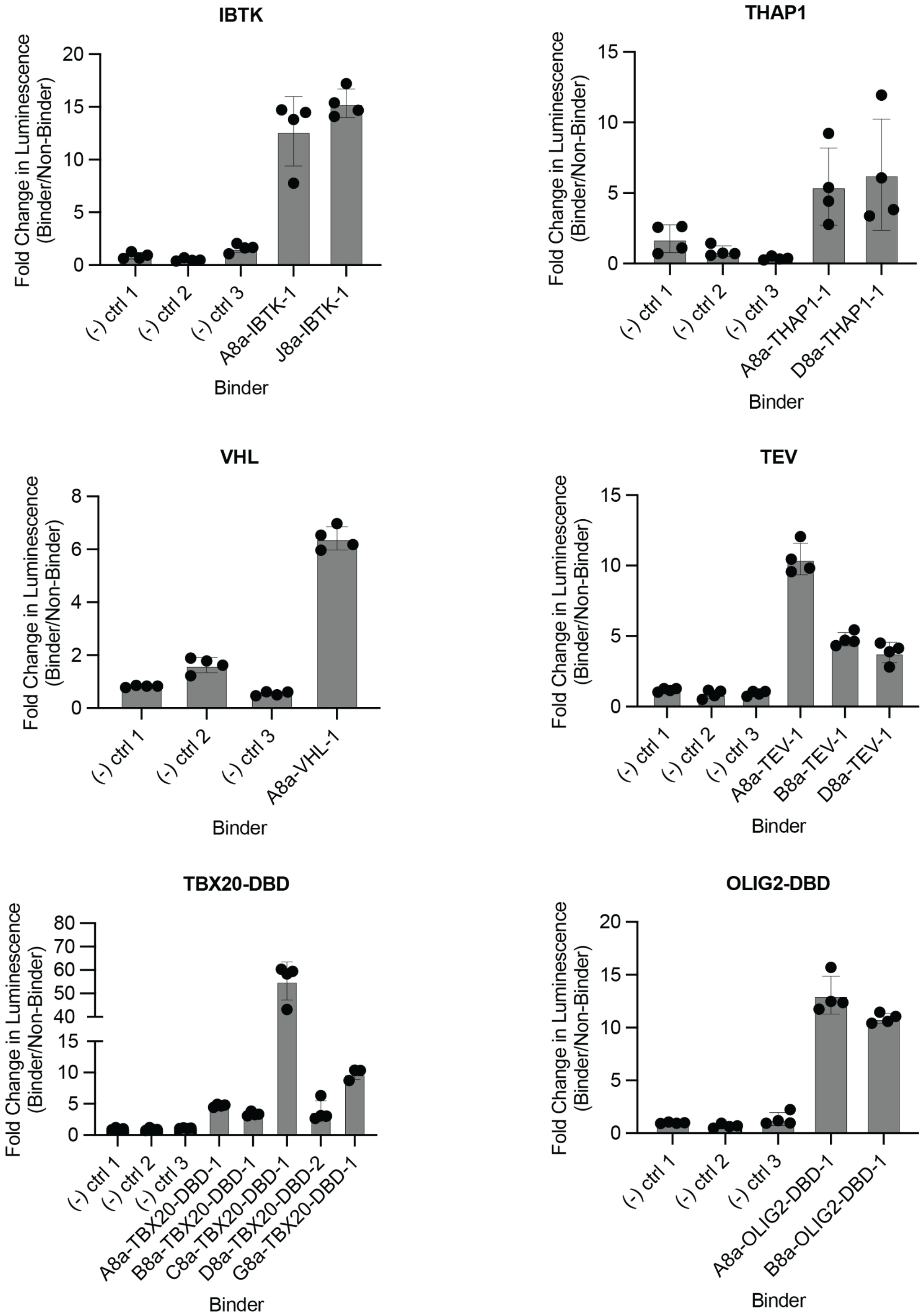
IBTK
Fold Change in Luminescence (Binder/Non-Binder)
0
5
10
15
20
(-) ctrl 1
(-) ctrl 2
(-) ctrl 3
A8a-IBTK-1
J8a-IBTK-1
Binder
THAP1
Fold Change in Luminescence (Binder/Non-Binder)
0
5
10
15
(-) ctrl 1
(-) ctrl 2
(-) ctrl 3
A8a-THAP1-1
D8a-THAP1-1
Binder
VHL
Fold Change in Luminescence (Binder/Non-Binder)
0
2
4
6
8
(-) ctrl 1
(-) ctrl 2
(-) ctrl 3
A8a-VHL-1
Binder
TEV
Fold Change in Luminescence (Binder/Non-Binder)
0
5
10
15
(-) ctrl 1
(-) ctrl 2
(-) ctrl 3
A8a-TEV-1
B8a-TEV-1
D8a-TEV-1
Binder
TBX20-DBD
Fold Change in Luminescence (Binder/Non-Binder)
0
5
10
15
40
50
60
70
80
(-) ctrl 1
(-) ctrl 2
(-) ctrl 3
A8a-TBX20-DBD-1
B8a-TBX20-DBD-1
C8a-TBX20-DBD-1
D8a-TBX20-DBD-2
G8a-TBX20-DBD-1
Binder
OLIG2-DBD
Fold Change in Luminescence (Binder/Non-Binder)
0
5
10
15
20
(-) ctrl 1
(-) ctrl 2
(-) ctrl 3
A8a-OLIG2-DBD-1
B8a-OLIG2-DBD-1
Binder

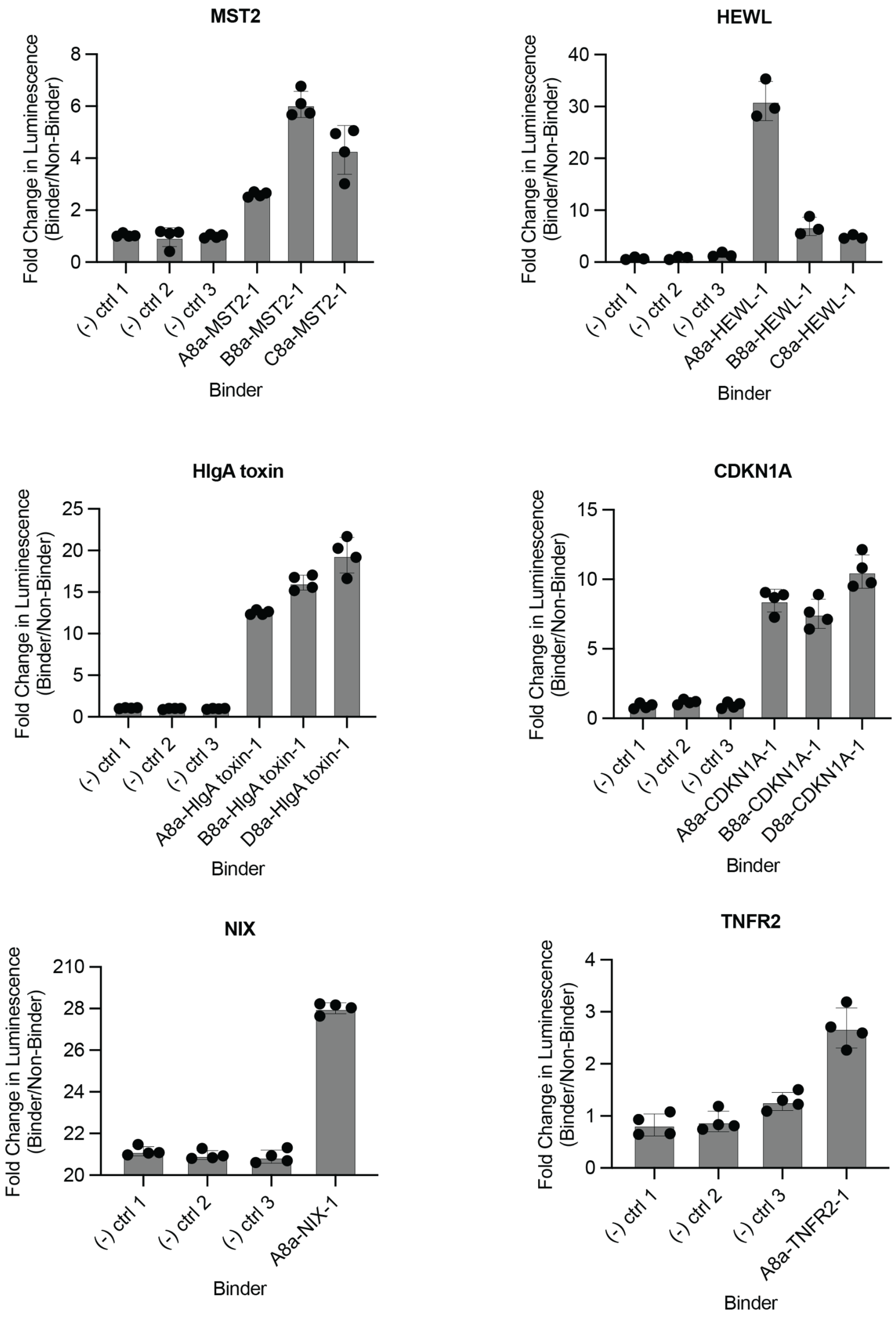
MST2
Fold Change in Luminescence (Binder/Non-Binder)
0
2
4
6
8
(-) ctrl 1
(-) ctrl 2
(-) ctrl 3
A8a-MST2-1
B8a-MST2-1
C8a-MST2-1
Binder
HEWL
Fold Change in Luminescence (Binder/Non-Binder)
0
10
20
30
40
(-) ctrl 1
(-) ctrl 2
(-) ctrl 3
A8a-HEWL-1
B8a-HEWL-1
C8a-HEWL-1
Binder
HlgA toxin
Fold Change in Luminescence (Binder/Non-Binder)
0
5
10
15
20
25
(-) ctrl 1
(-) ctrl 2
(-) ctrl 3
A8a-HlgA toxin-1
B8a-HlgA toxin-1
D8a-HlgA toxin-1
Binder
CDKN1A
Fold Change in Luminescence (Binder/Non-Binder)
0
5
10
15
(-) ctrl 1
(-) ctrl 2
(-) ctrl 3
A8a-CDKN1A-1
B8a-CDKN1A-1
D8a-CDKN1A-1
Binder
NIX
Fold Change in Luminescence (Binder/Non-Binder)
20
22
24
26
28
210
(-) ctrl 1
(-) ctrl 2
(-) ctrl 3
A8a-NIX-1
Binder
TNFR2
Fold Change in Luminescence (Binder/Non-Binder)
0
1
2
3
4
(-) ctrl 1
(-) ctrl 2
(-) ctrl 3
A8a-TNFR2-1
Binder

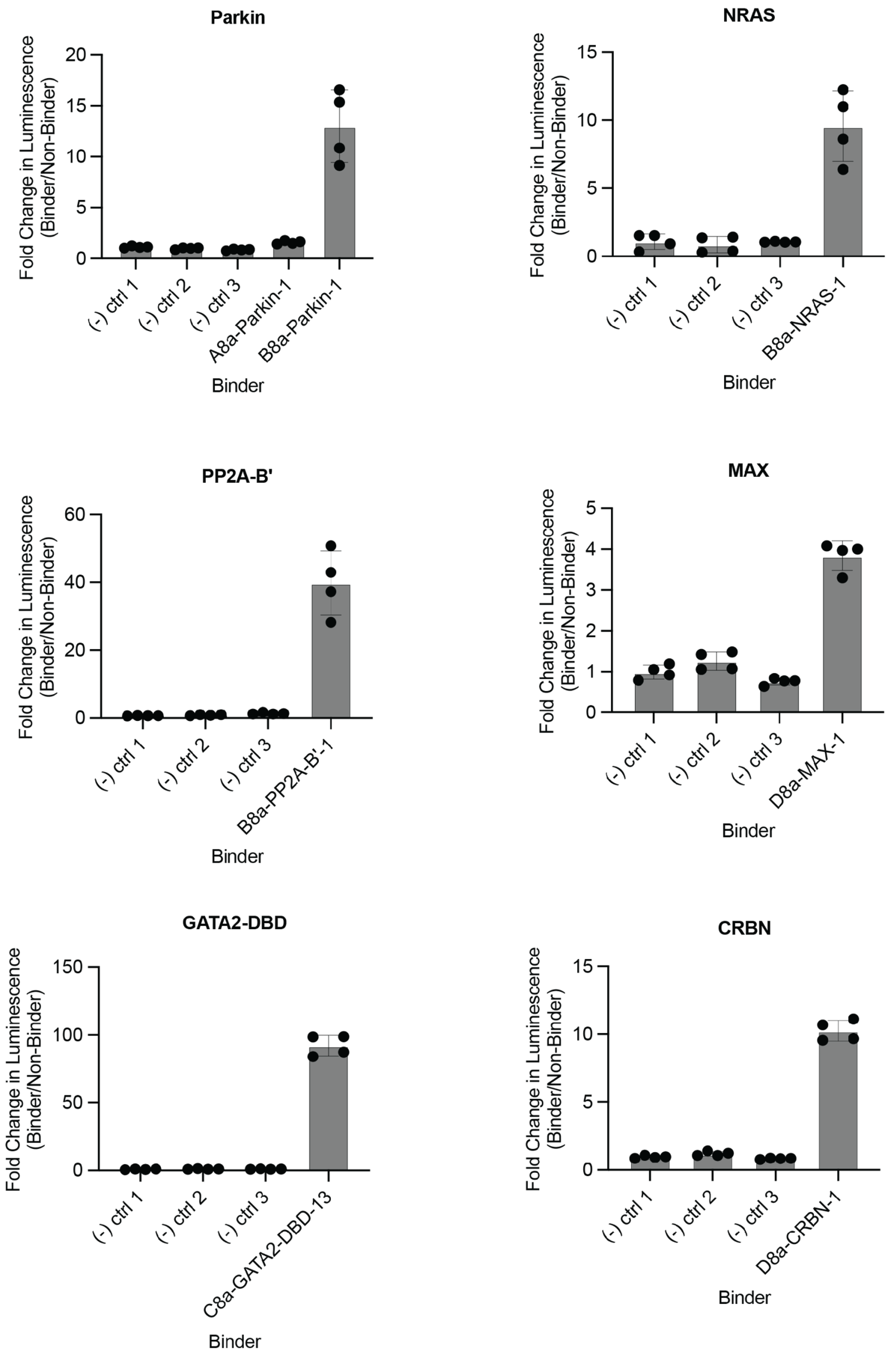
Parkin
Fold Change in Luminescence (Binder/Non-Binder)
0
5
10
15
20
(-) ctrl 1
(-) ctrl 2
(-) ctrl 3
A8a-Parkin-1
B8a-Parkin-1
Binder
NRAS
Fold Change in Luminescence (Binder/Non-Binder)
0
5
10
15
(-) ctrl 1
(-) ctrl 2
(-) ctrl 3
B8a-NRAS-1
Binder
PP2A-B'
Fold Change in Luminescence (Binder/Non-Binder)
0
20
40
60
(-) ctrl 1
(-) ctrl 2
(-) ctrl 3
B8a-PP2A-B'-1
Binder
MAX
Fold Change in Luminescence (Binder/Non-Binder)
0
1
2
3
4
5
(-) ctrl 1
(-) ctrl 2
(-) ctrl 3
D8a-MAX-1
Binder
GATA2-DBD
Fold Change in Luminescence (Binder/Non-Binder)
0
50
100
150
(-) ctrl 1
(-) ctrl 2
(-) ctrl 3
C8a-GATA2-DBD-13
Binder
CRBN
Fold Change in Luminescence (Binder/Non-Binder)
0
5
10
15
(-) ctrl 1
(-) ctrl 2
(-) ctrl 3
D8a-CRBN-1
Binder

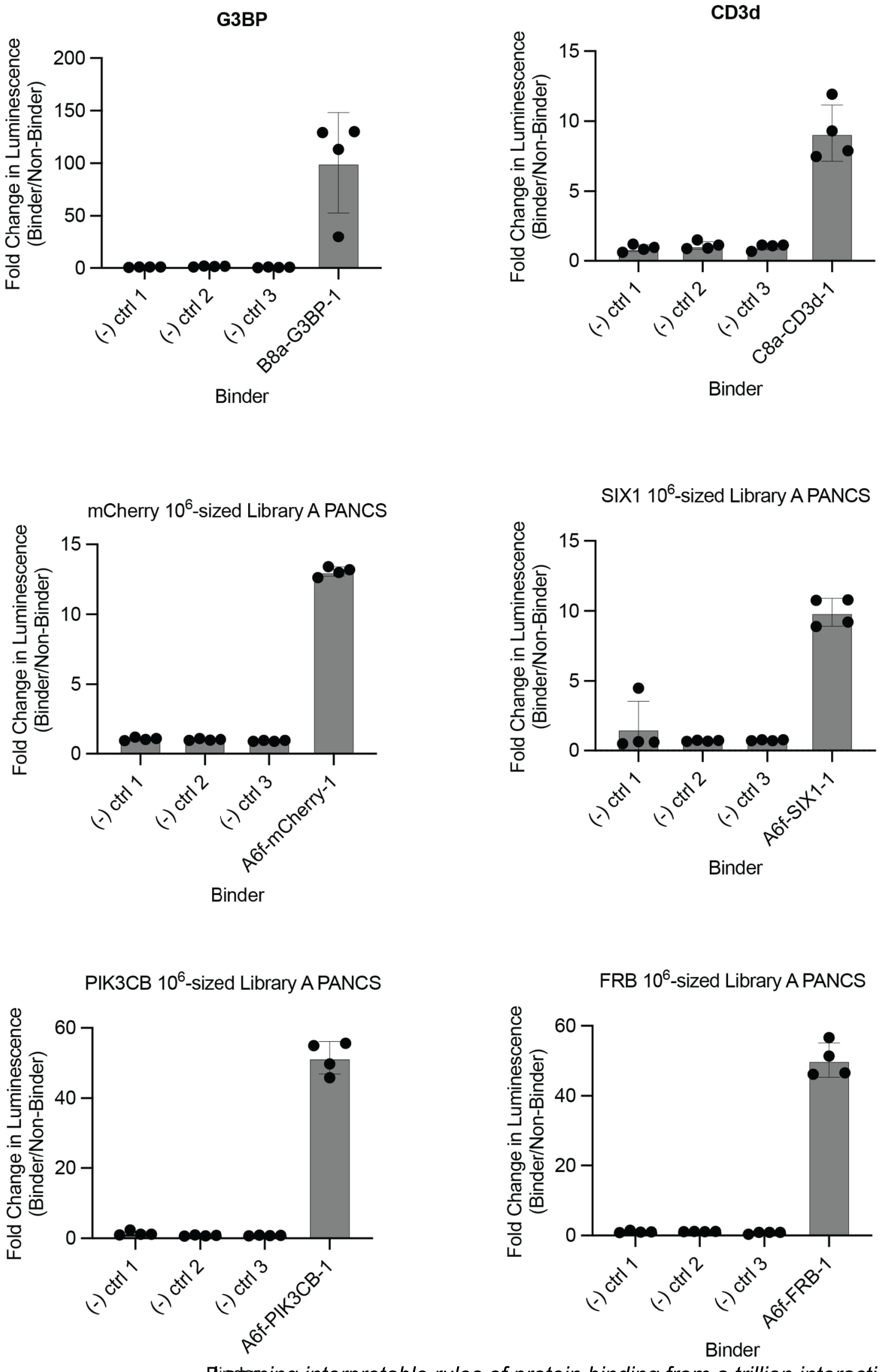
G3BP
Fold Change in Luminescence (Binder/Non-Binder)
0
50
100
150
200
(-) ctrl 1
(-) ctrl 2
(-) ctrl 3
B8a-G3BP-1
Binder
CD3d
Fold Change in Luminescence (Binder/Non-Binder)
0
5
10
15
(-) ctrl 1
(-) ctrl 2
(-) ctrl 3
C8a-CD3d-1
Binder
mCherry 10^6-sized Library A PANCS
Fold Change in Luminescence (Binder/Non-Binder)
0
5
10
15
(-) ctrl 1
(-) ctrl 2
(-) ctrl 3
A6f-mCherry-1
Binder
SIX1 10^6-sized Library A PANCS
Fold Change in Luminescence (Binder/Non-Binder)
0
5
10
15
(-) ctrl 1
(-) ctrl 2
(-) ctrl 3
A6f-SIX1-1
Binder
PIK3CB 10^6-sized Library A PANCS
Fold Change in Luminescence (Binder/Non-Binder)
0
20
40
60
(-) ctrl 1
(-) ctrl 2
(-) ctrl 3
A6f-PIK3CB-1
Binder
FRB 10^6-sized Library A PANCS
Fold Change in Luminescence (Binder/Non-Binder)
0
20
40
60
(-) ctrl 1
(-) ctrl 2
(-) ctrl 3
A6f-FRB-1
Binder

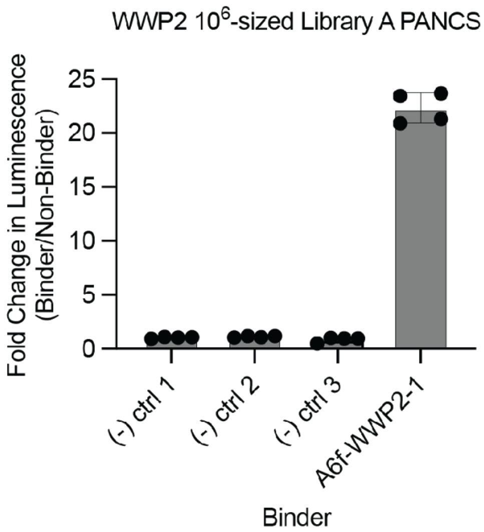
WWP2 $10^6$-sized Library A PANCS
Fold Change in Luminescence (Binder/Non-Binder)
0
5
10
15
20
25
(-) ctrl 1
(-) ctrl 2
(-) ctrl 3
A6f-WWP2-1
Binder

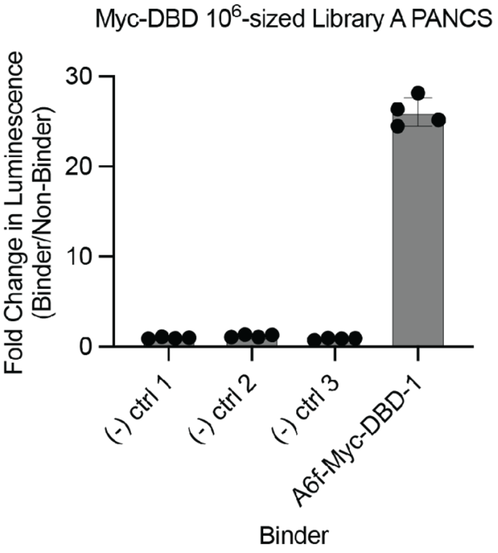
Myc-DBD $10^6$-sized Library A PANCS
Fold Change in Luminescence (Binder/Non-Binder)
0
10
20
30
(-) ctrl 1
(-) ctrl 2
(-) ctrl 3
A6f-Myc-DBD-1
Binder

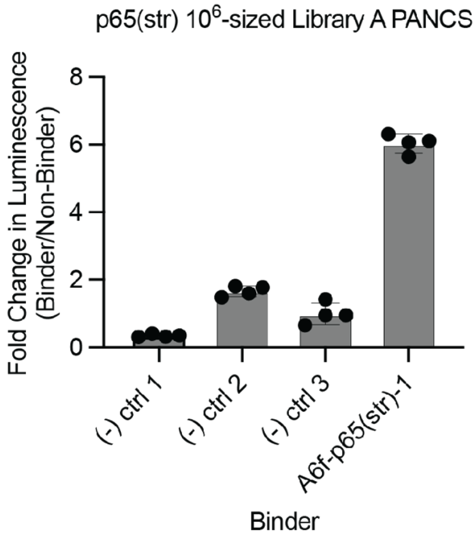
p65(str) $10^6$-sized Library A PANCS
Fold Change in Luminescence (Binder/Non-Binder)
0
2
4
6
8
(-) ctrl 1
(-) ctrl 2
(-) ctrl 3
A6f-p65(str)-1
Binder

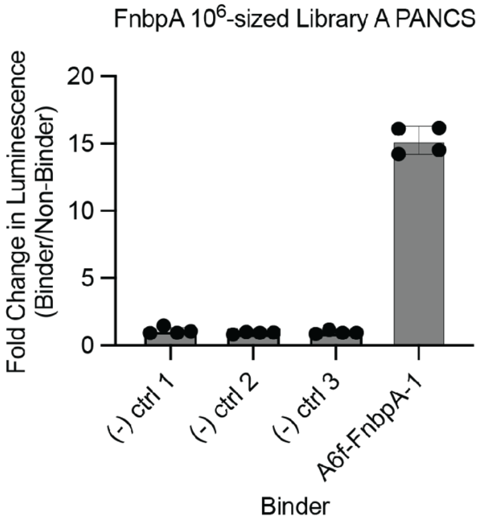
FnbpA $10^6$-sized Library A PANCS
Fold Change in Luminescence (Binder/Non-Binder)
0
5
10
15
20
(-) ctrl 1
(-) ctrl 2
(-) ctrl 3
A6f-FnbpA-1
Binder

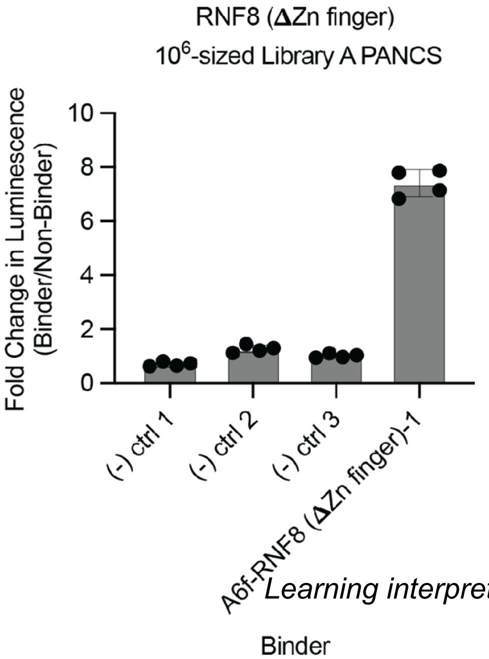
RNF8 (ΔZn finger)
$10^6$-sized Library A PANCS
Fold Change in Luminescence (Binder/Non-Binder)
0
2
4
6
8
10
(-) ctrl 1
(-) ctrl 2
(-) ctrl 3
A6f-RNF8 (ΔZn finger)-1
Binder

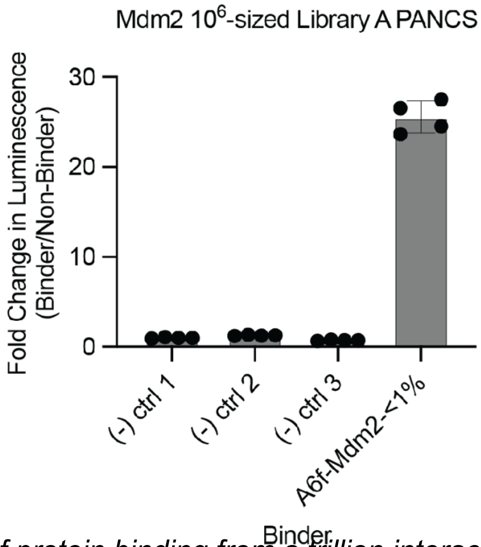
Mdm2 $10^6$-sized Library A PANCS
Fold Change in Luminescence (Binder/Non-Binder)
0
10
20
30
(-) ctrl 1
(-) ctrl 2
(-) ctrl 3
A6f-Mdm2-<1%
Binder

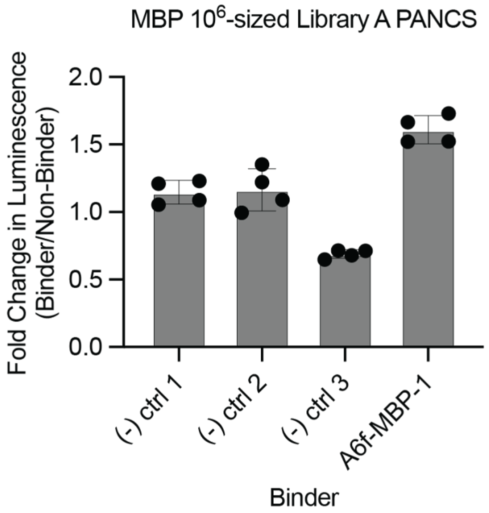
MBP 10^6-sized Library A PANCS
Fold Change in Luminescence (Binder/Non-Binder)
2.0
1.5
1.0
0.5
0.0
(-) ctrl 1
(-) ctrl 2
(-) ctrl 3
A6f-MBP-1
Binder

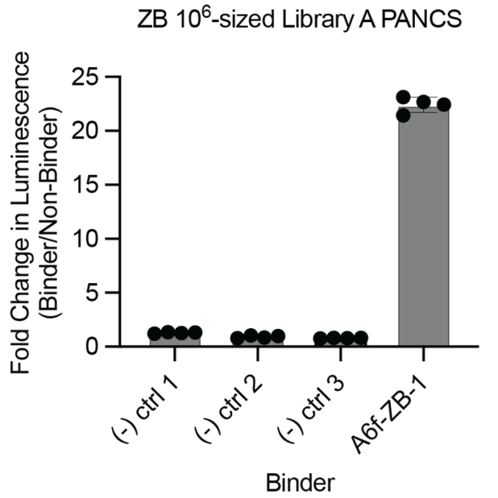
ZB 10^6-sized Library A PANCS
Fold Change in Luminescence (Binder/Non-Binder)
25
20
15
10
5
0
(-) ctrl 1
(-) ctrl 2
(-) ctrl 3
A6f-ZB-1
Binder

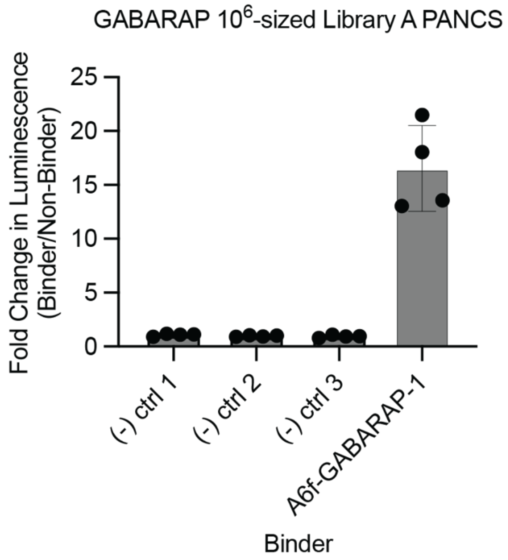
GABARAP 10^6-sized Library A PANCS
Fold Change in Luminescence (Binder/Non-Binder)
25
20
15
10
5
0
(-) ctrl 1
(-) ctrl 2
(-) ctrl 3
A6f-GABARAP-1
Binder

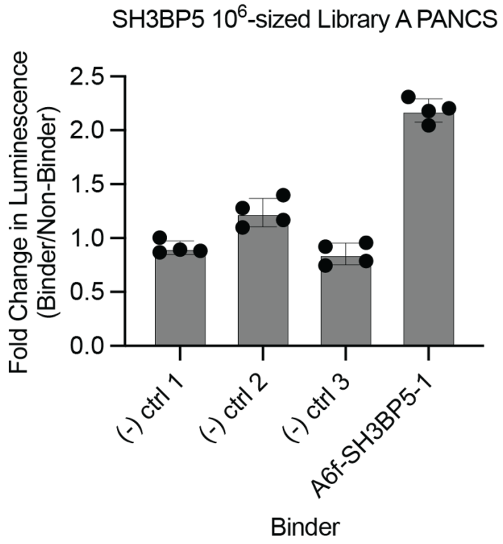
SH3BP5 10^6-sized Library A PANCS
Fold Change in Luminescence (Binder/Non-Binder)
2.5
2.0
1.5
1.0
0.5
0.0
(-) ctrl 1
(-) ctrl 2
(-) ctrl 3
A6f-SH3BP5-1
Binder

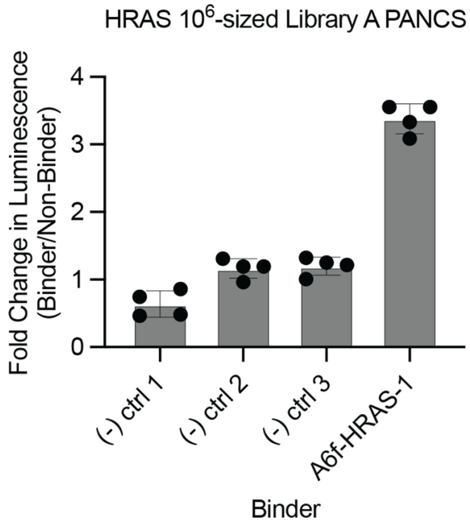
HRAS 10^6-sized Library A PANCS
Fold Change in Luminescence (Binder/Non-Binder)
4
3
2
1
0
(-) ctrl 1
(-) ctrl 2
(-) ctrl 3
A6f-HRAS-1
Binder

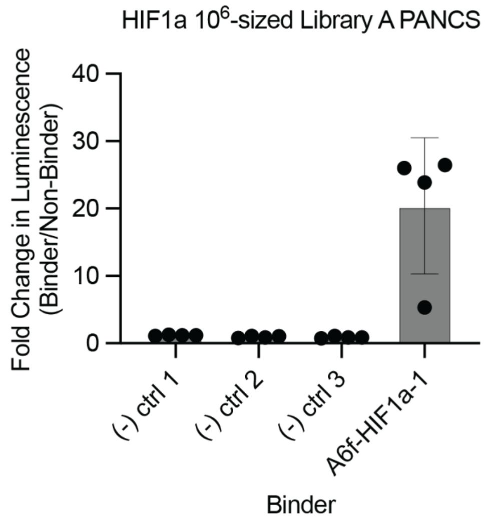
HIF1a 10^6-sized Library A PANCS
Fold Change in Luminescence (Binder/Non-Binder)
40
30
20
10
0
(-) ctrl 1
(-) ctrl 2
(-) ctrl 3
A6f-HIF1a-1
Binder

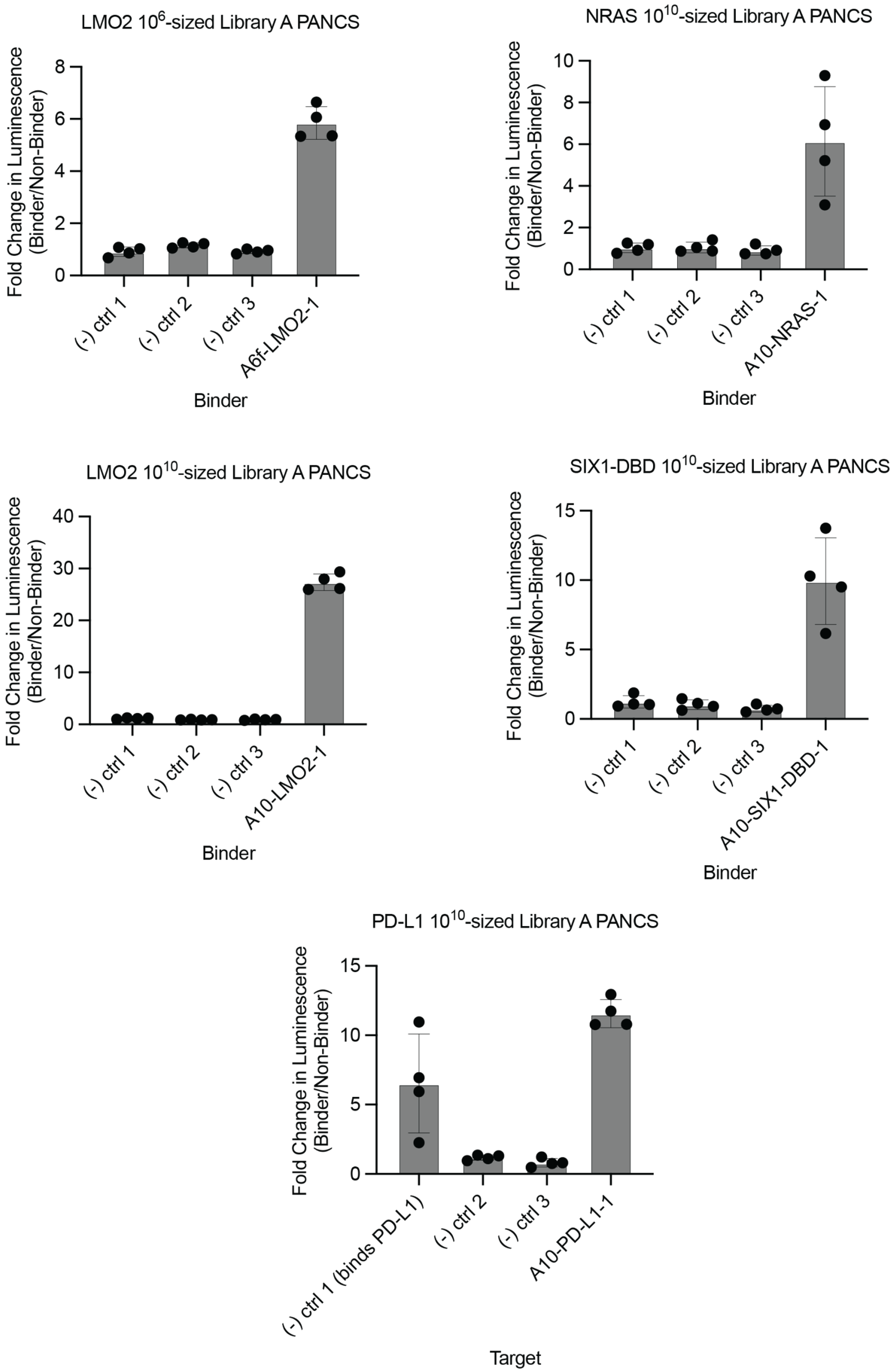
LMO2 10^6-sized Library A PANCS
Fold Change in Luminescence (Binder/Non-Binder)
0 2 4 6 8
(-) ctrl 1
(-) ctrl 2
(-) ctrl 3
A6f-LMO2-1
Binder
NRAS 10^10-sized Library A PANCS
Fold Change in Luminescence (Binder/Non-Binder)
0 2 4 6 8 10
(-) ctrl 1
(-) ctrl 2
(-) ctrl 3
A10-NRAS-1
Binder
LMO2 10^10-sized Library A PANCS
Fold Change in Luminescence (Binder/Non-Binder)
0 10 20 30 40
(-) ctrl 1
(-) ctrl 2
(-) ctrl 3
A10-LMO2-1
Binder
SIX1-DBD 10^10-sized Library A PANCS
Fold Change in Luminescence (Binder/Non-Binder)
0 5 10 15
(-) ctrl 1
(-) ctrl 2
(-) ctrl 3
A10-SIX1-DBD-1
Binder
PD-L1 10^10-sized Library A PANCS
Fold Change in Luminescence (Binder/Non-Binder)
0 5 10 15
(-) ctrl 1 (binds PD-L1)
(-) ctrl 2
(-) ctrl 3
A10-PD-L1-1
Target

**Supplementary Fig. 6: Split T7 RNAP *E. coli* luciferase binding assay for 96-target panel.** Luciferase assay results for each target-binder pair with target-non-binder controls (affibody (PD-L1)[2], Monobody (hSUMO1)[3], and affitin (SasA)[4]). To calculate the fold-change of the binder over the non-targeting controls, the blank $OD_{600}$ signal (media only) was subtracted from the $OD_{600}$ of each sample, then the luminescence signal of each binder was divided by this modified $OD_{600}$. The average of all three negative, non-binding controls was calculated for each target, then the fold-change of each binder colony was divided by this non-binding control average for each target to produce the fold-change of the binder over the non-targeting controls. Each bar plot shows the mean, standard deviation, and individual values ($n$ = 4, error bars indicate SD). All luciferase assay plots shown, unless otherwise labeled as $10^6$ or $10^{10}$, are representative of selections run with $10^8$ unique variants. Binder naming is design, log size, independent transformation as a triplet alphanumeric code.

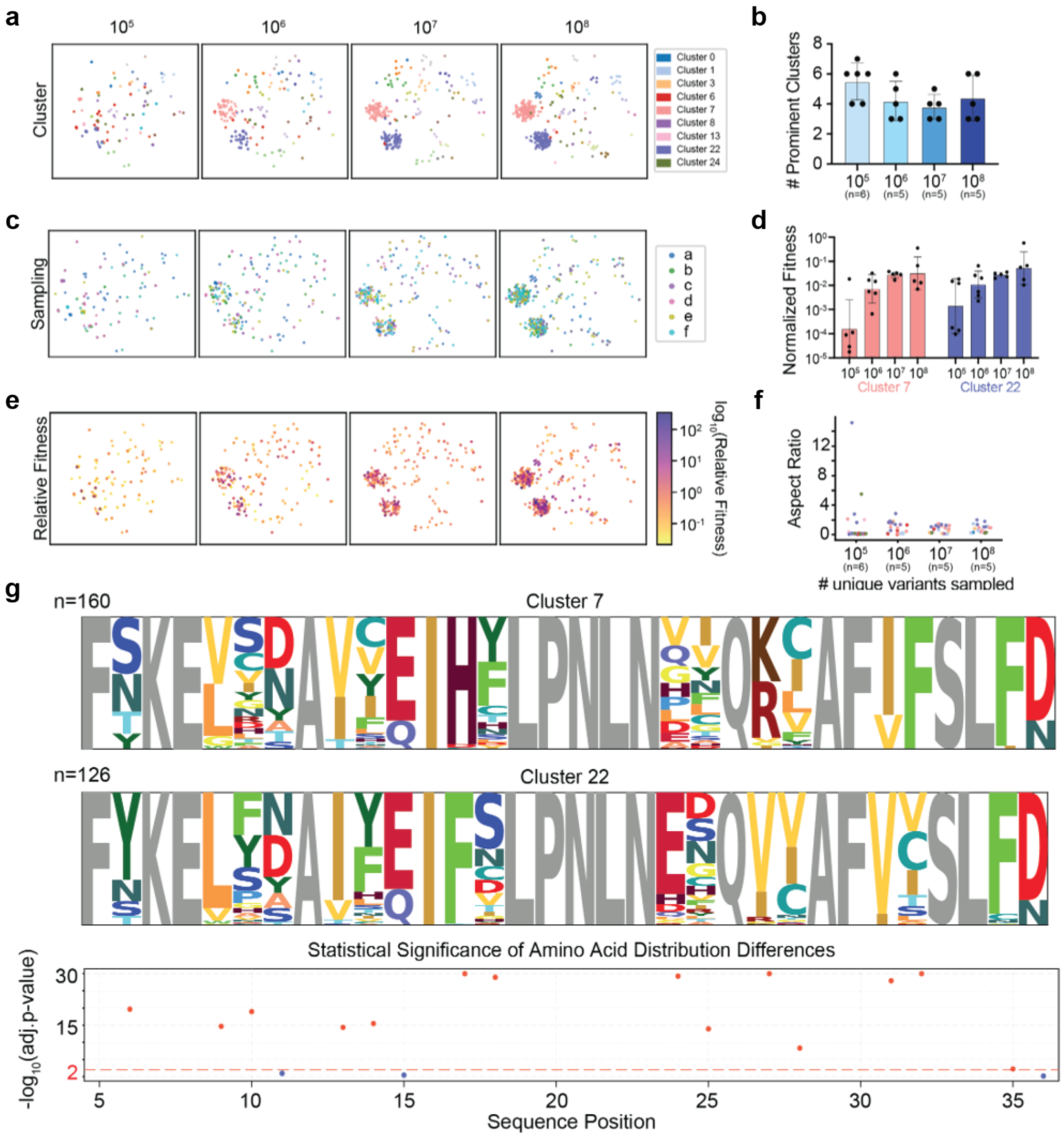

**Supplementary Fig. 7: Clustering details for IFNG.** Plot organization for each target: **a**, Binders for all samplings at sampling sizes from $10^5$ to $10^8$ unique variants for a target plotted on a t-SNE embedding colored by cluster associations. **b**, Number of prominent clusters (>1% of NGS reads) for the target for each sampling size. **c**, Same t-SNE plot as A, but colored by the sampling. **d**, Normalized fitness of each binder variant tested (top variant from the selection) sorted by the top two clusters (by the clusters' % of NGS reads in the $10^8$ selections). **e**, Same t-SNE as A, but coloring by relative fitness (see Methods for details on how the normalized fitness is used to compare fitness of all binders in the NGS across selections). **f**, The aspect ratio (relative fitness divided by the fraction of NGS reads for each cluster). **g**, Sequence logo plots illustrating amino acid distributions at every position for the clusters shown in **D** (frequency is a count of unique variants with that amino acid) – grey indicates not randomized; adjusted p-values at each protein position between the distributions of the two clusters, with the red dashed line corresponding to a significance level of 0.01. Four general landscape topologies were observed across the 9-target panel: IFNG displayed two prominent, well-separated clusters; XIAP displayed two prominent but highly similar clusters; KIX displayed both IFNG- and XIAP-like topologies; and LC3B, PP2A-B'' and CDKN1A had much sparser landscapes, consistent with the sampling size being nearer to the binder density for these three targets.

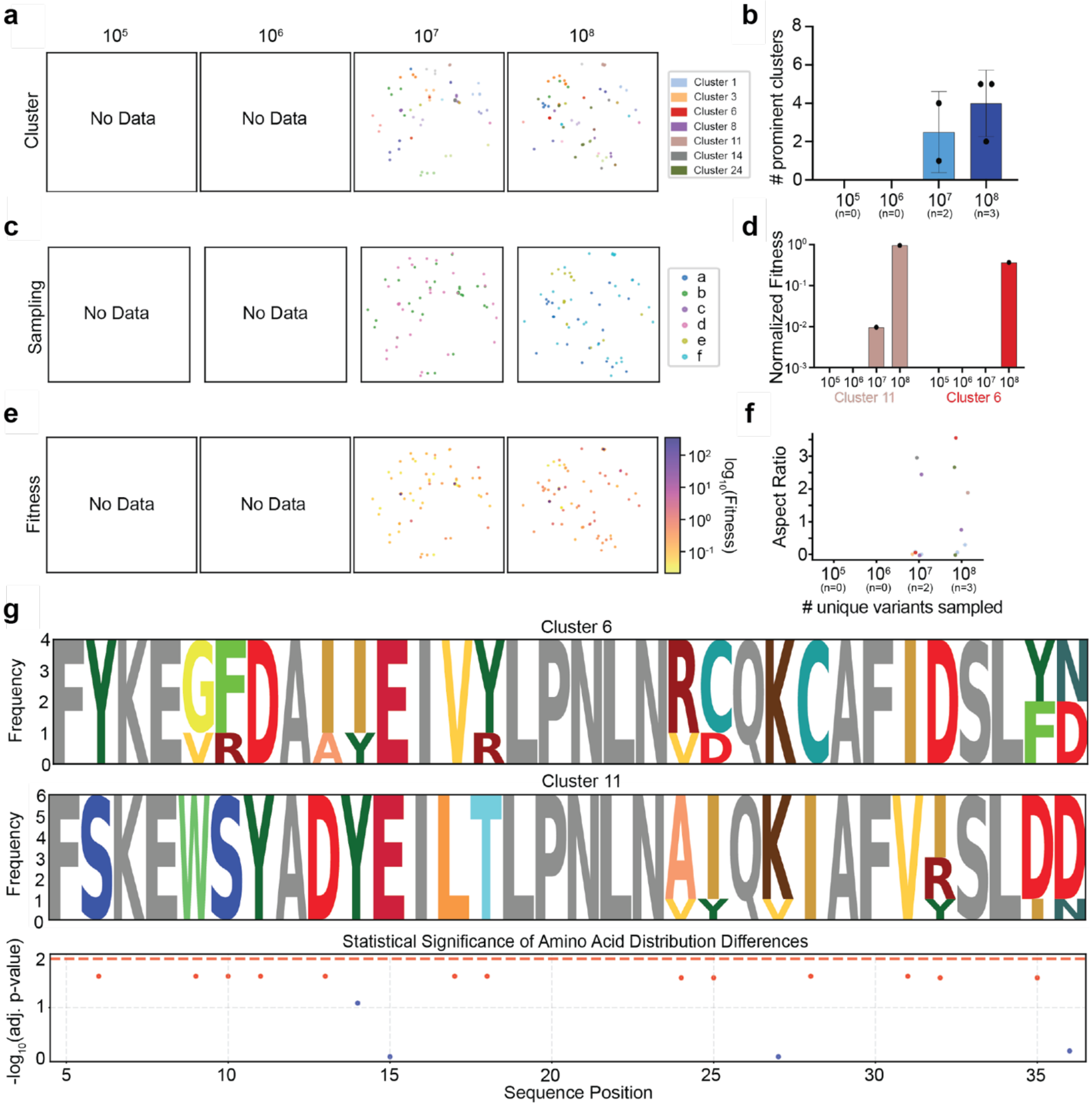

**Supplementary Fig. 7 (continued). Clustering details for CDKN1A.**

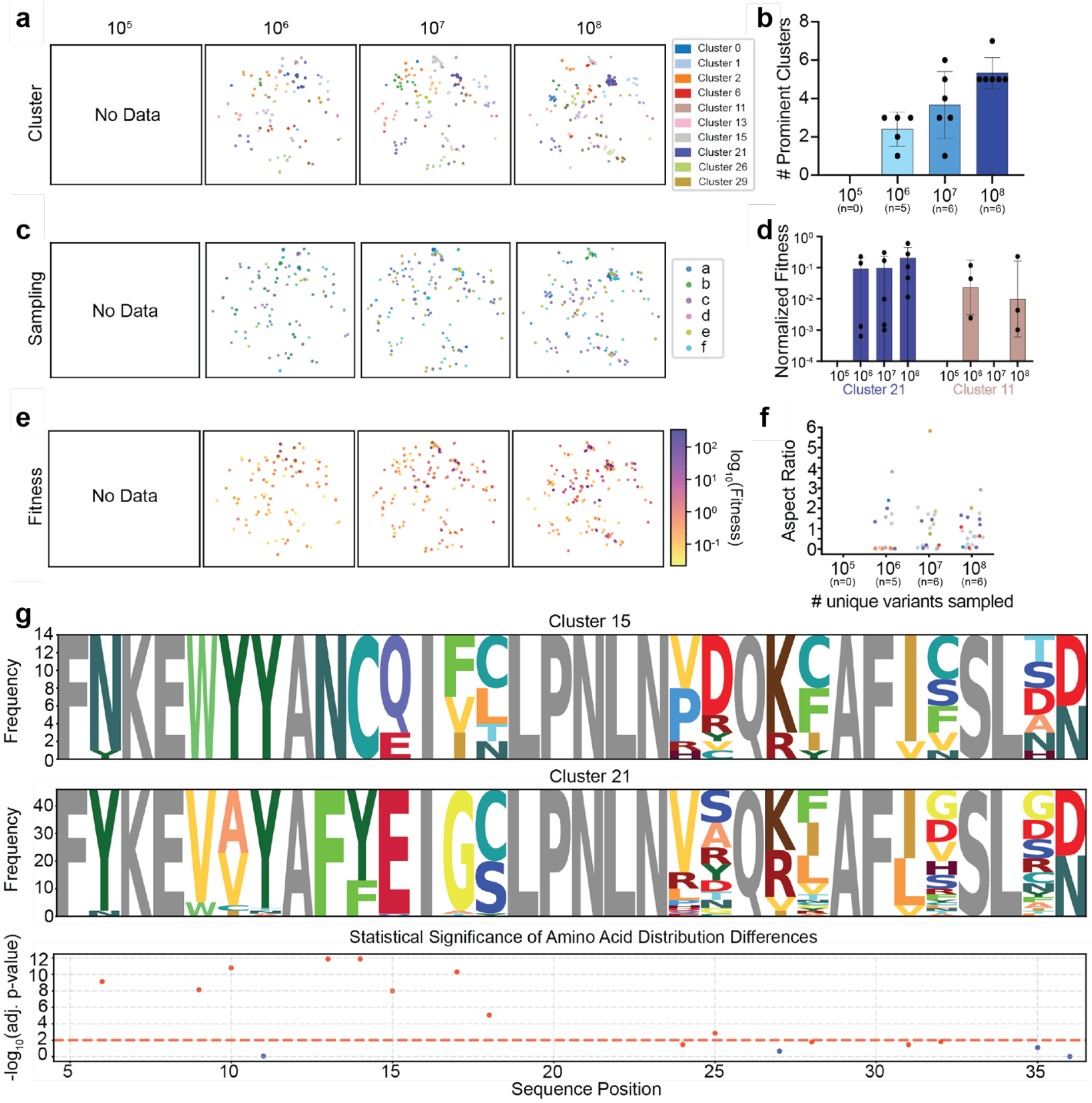

**Supplementary Fig. 7 (continued). Clustering details for PP2A-B''.**

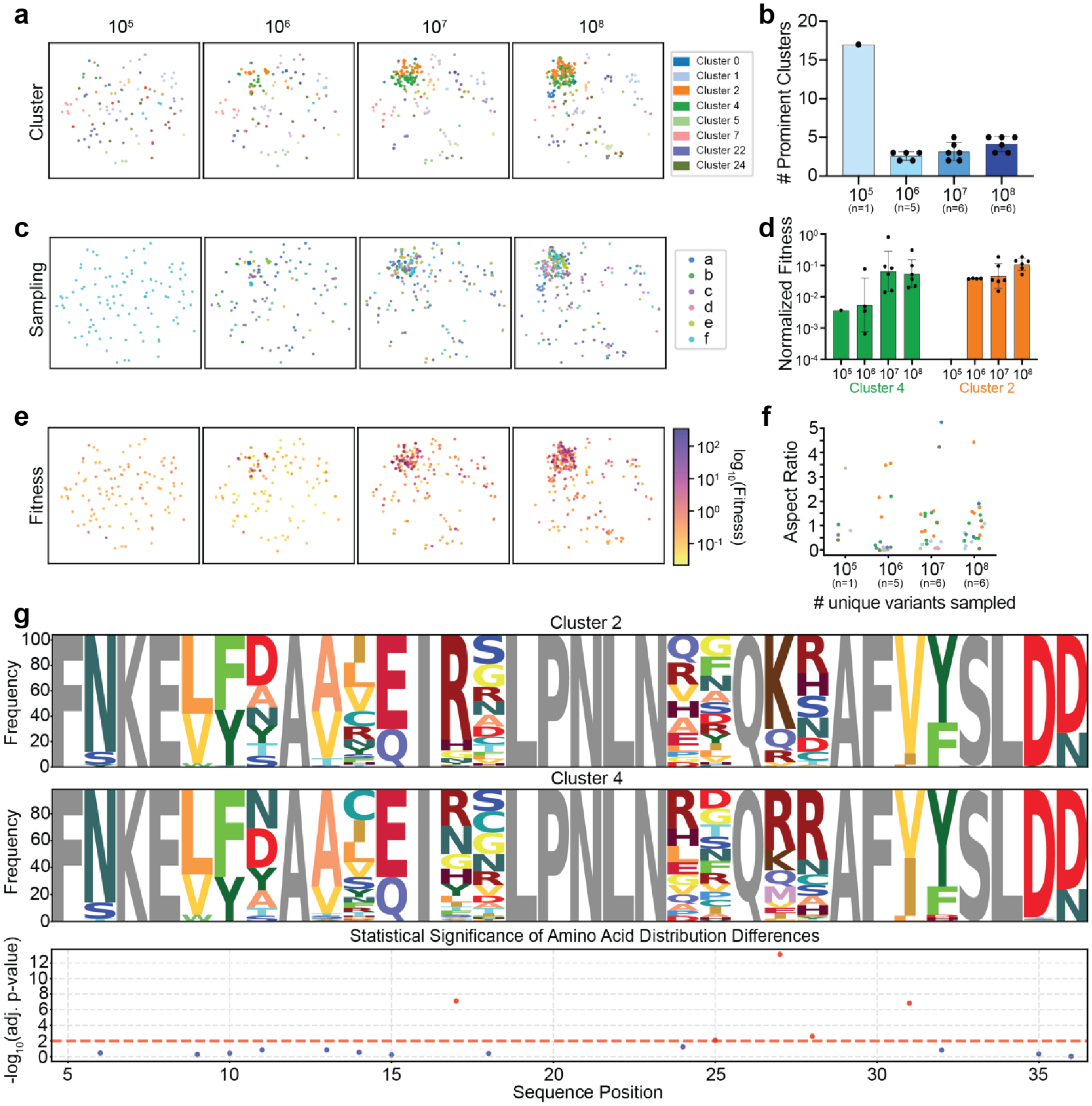

**Supplementary Fig. 7 (continued). Clustering details for XIAP.**

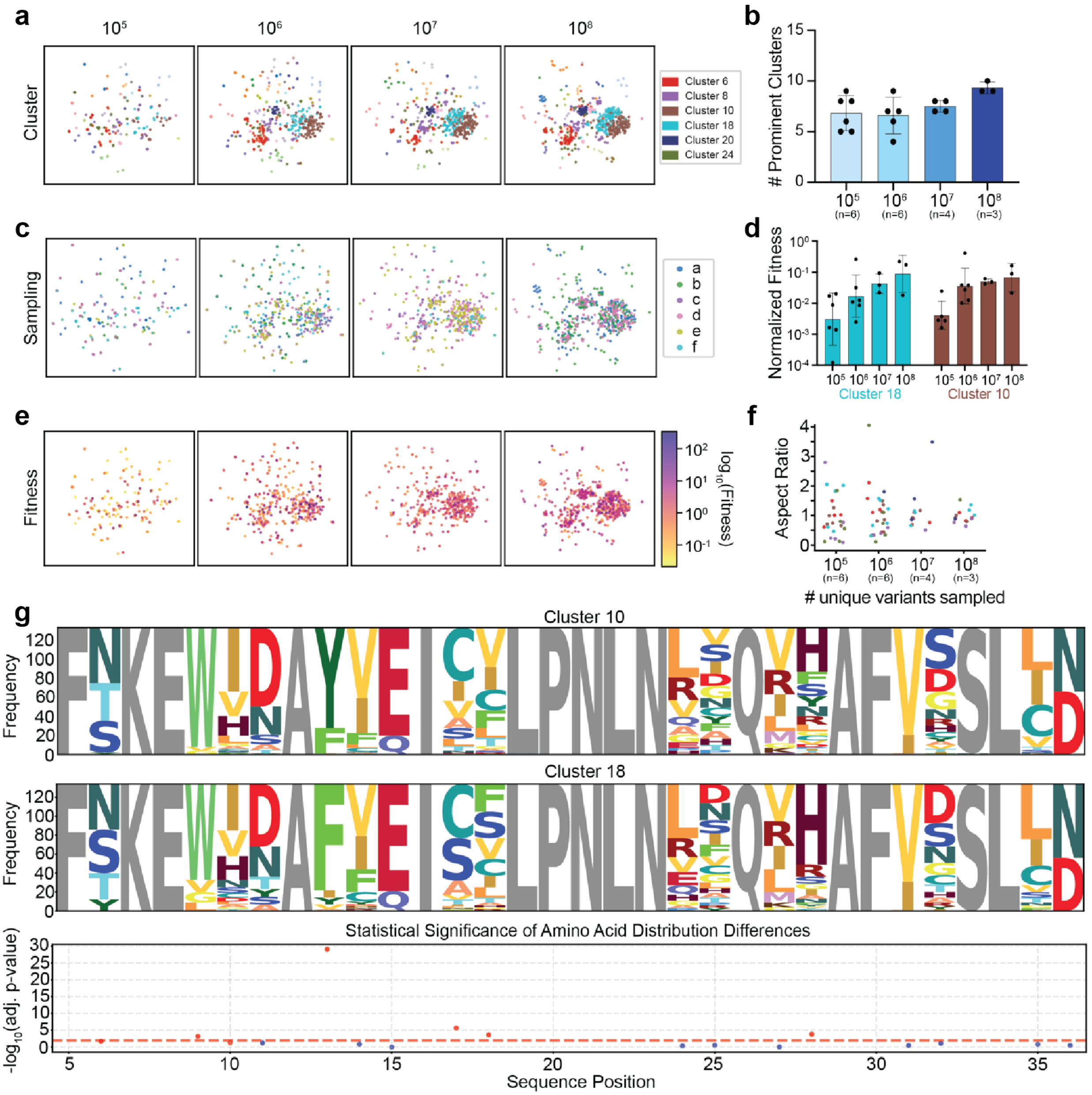

**Supplementary Fig. 7 (continued). Clustering details for KIX.**

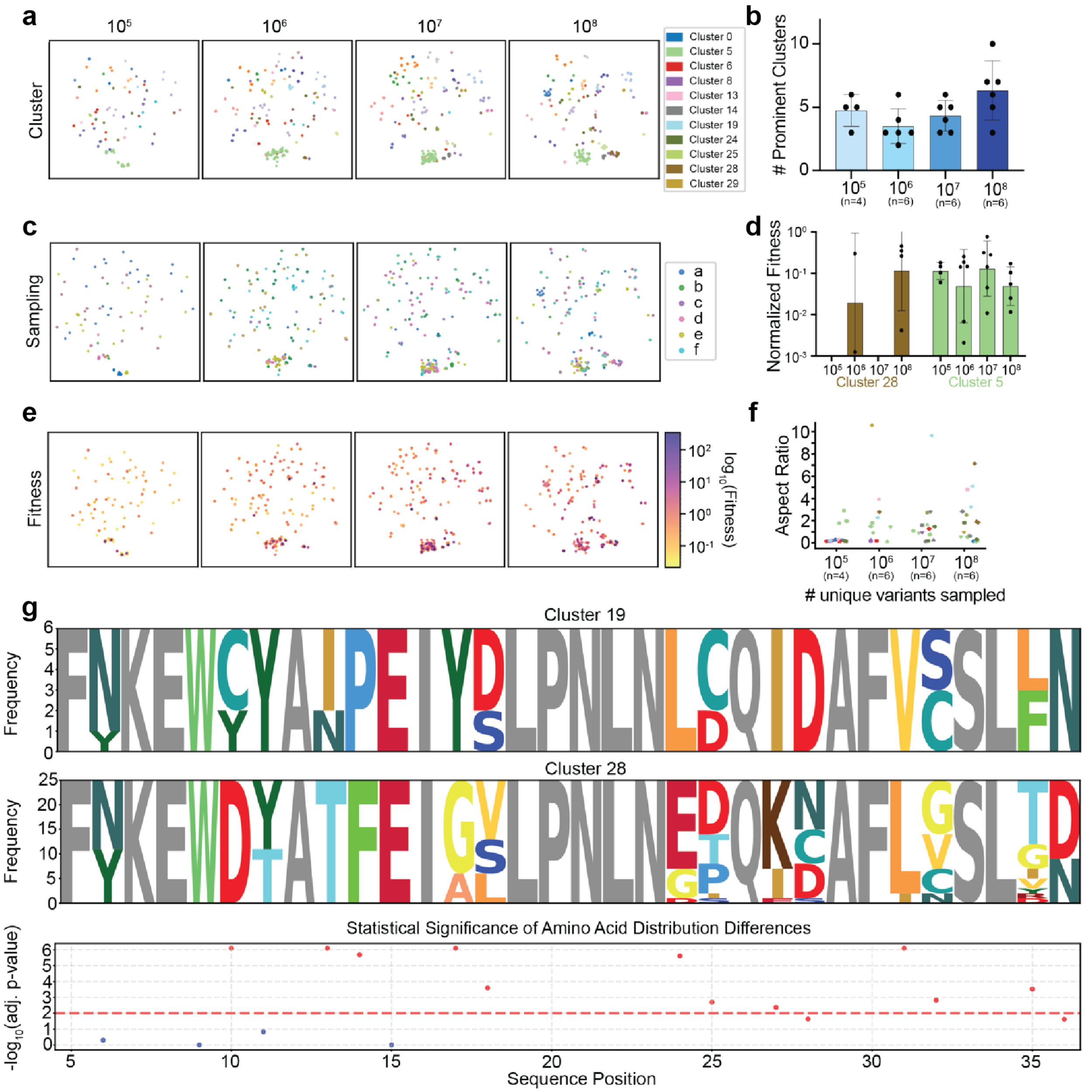

**Supplementary Fig. 7 (continued). Clustering details for LC3B.**

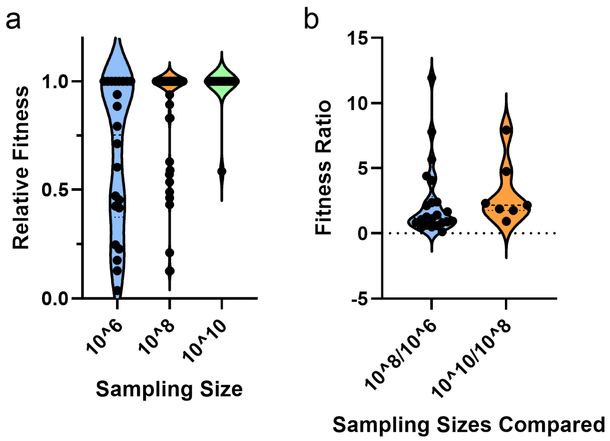

**Supplementary Fig. 8: Comparing binder fitness across sampling sizes. a**, The relative fitness value for each of the top binder variants from **Supplementary Fig. 6** is shown for each different sampling size. The average relative fitness varies from 0.66 (SD 0.35) for binders from the $10^6$ sampling size to 0.87 (SD 0.25) for binders from the $10^8$ sampling size to 0.97 (SD 0.12) for binders from the $10^{10}$ sampling size (note that these are skewed towards 1 in the $10^8$ set for any target that did not hit at $10^6$ and in the $10^{10}$ set for any target that did not hit at $10^8$. **b**, To correct for this, we also calculated the ratio of fitness for all targets for which we identified binders from the $10^6$ and $10^8$ or $10^8$ and $10^{10}$ sample size selections. On 4 out of 22 targets had substantially higher fitness in the $10^6$ sampling compared to the $10^8$ sampling (ratio of <0.8; 0/7 for $10^8$ vs $10^{10}$). The average ratios were 2.4 (SD 2.9) and 3.1 (SD 2.4), respectively. Consistent with the increasing affinity of variants from selections with different sampling sizes on the 9-target panel (**Supplementary Fig. 4**), the binding fitness of the dominant variant increases with larger sampling size.

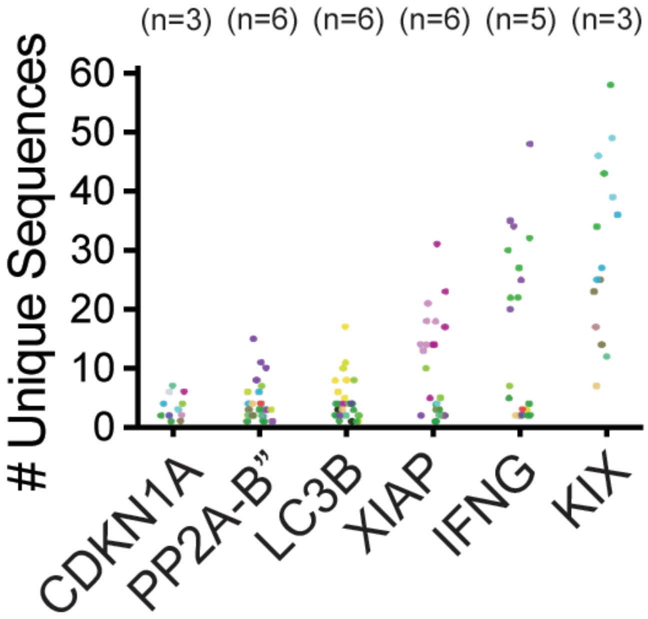

**Supplementary Fig. 9: Number of unique sequences per cluster ($10^8$ samplings only).** Same coloring as in **Supplementary Fig. 7.**

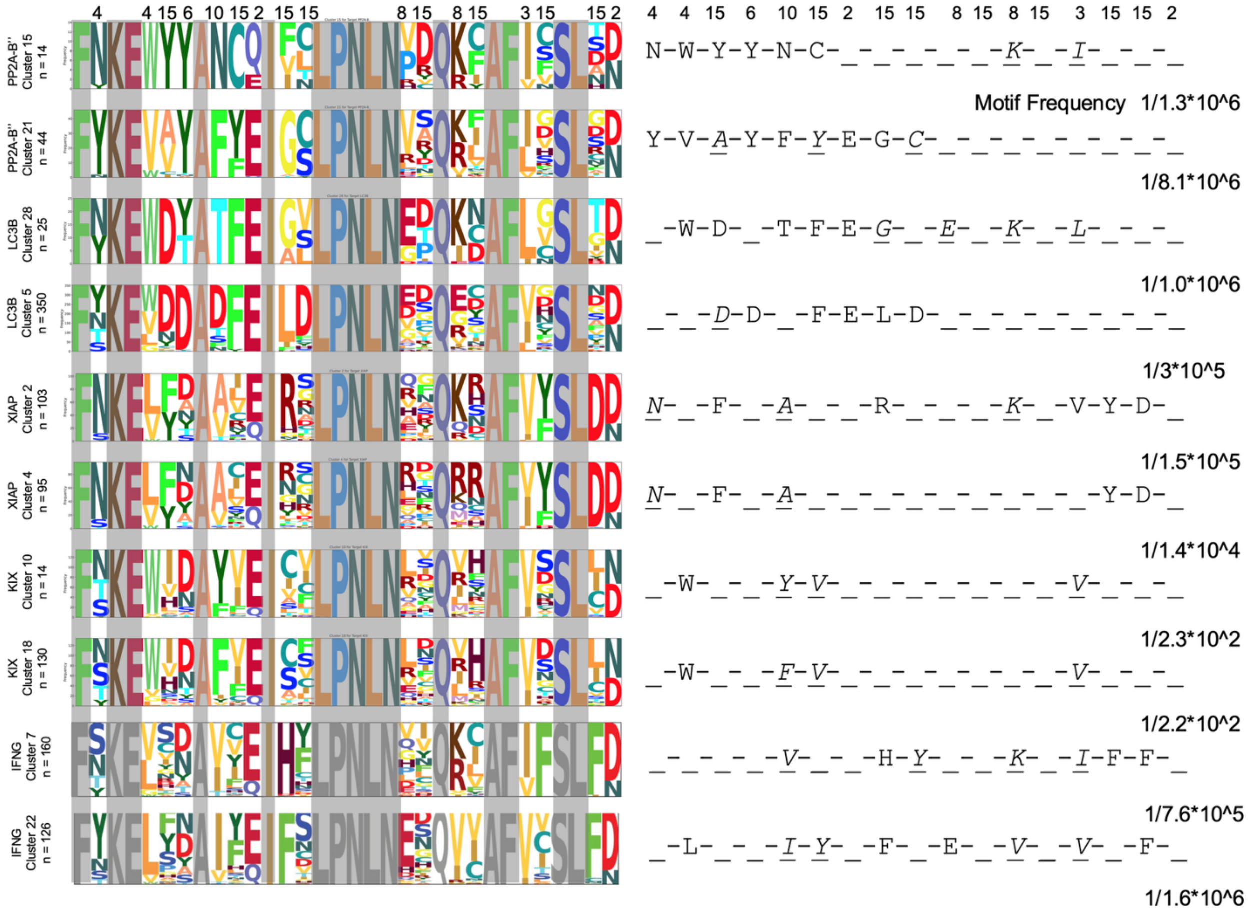

**Supplementary Fig. 10: Calculation of motif frequencies in randomization Design A**. Sequence logos (**Supplementary Fig. 7**) depicting the motif convergence of the two largest clusters for each target with at least 10 sequences, representing residues 4-36 of affibody only. To the right are the identified binding motifs with each position randomized listed as a single letter amino acid code or an underscore with dashes between each position. Highly conserved residues are identified as positions where a single amino acid has convergence equal to or greater than the expected percentage given the randomization (i.e. the first position can be four amino acids (listed above each position in the logo), so each as an expected percent of 25%) plus 45%; if two amino acids meet this threshold, then the greater of the two is indicated with italics and underlining at that position. Motif frequencies are calculated as multiplication of the individual identified positions (the number of amino acids possible is indicated on the top right). If a position is in italics/underlined (indicating two amino acids can be at that position), then the frequency is cut in half. For example, IFNG cluster 22 motif frequency is calculated as (1/4)*(1/5)*(1/7.5)*(1/15)*(1/8)*(1/4)*(1/1.5)*(1/15) = $6.2*10^{-7}$.

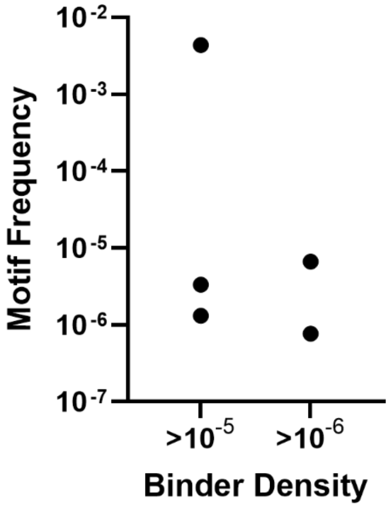

**Supplementary Fig. 11: Motif Frequency vs Binder Density for the 9-target panel**. Most frequent cluster shown for the 5 targets whose binding motif frequency was determined for the two most dominant clusters.

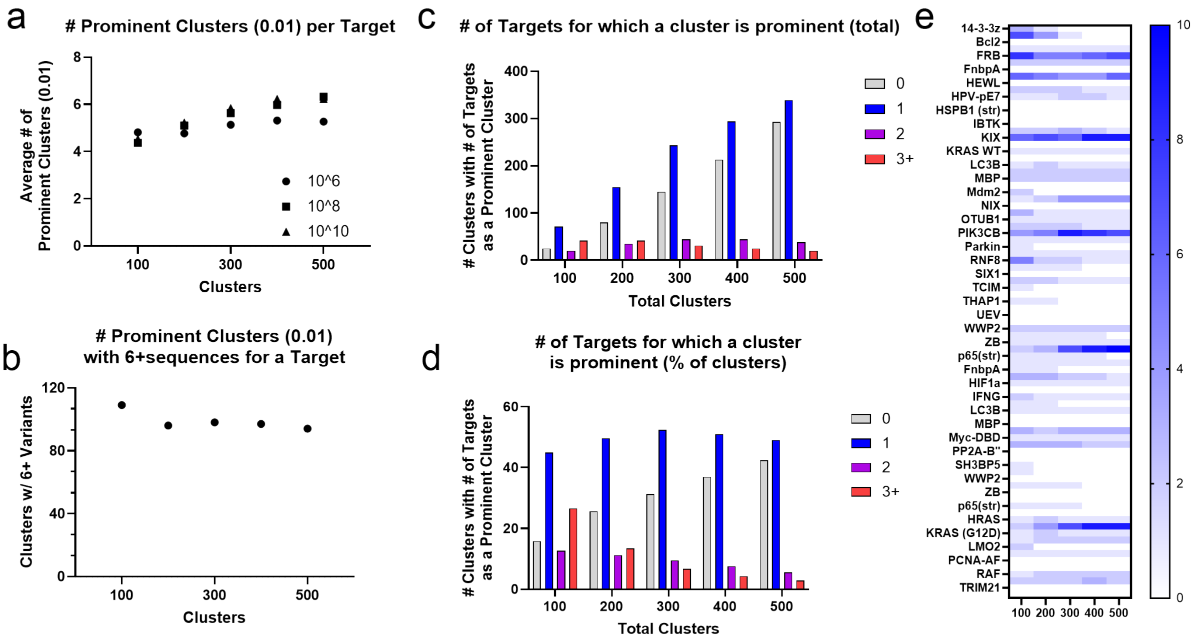

**Supplementary Fig. 12: How maximum number of clusters impacts hierarchical clustering results**. **a**, The number of prominent clusters per target for each selection size. **b**, The number of prominent clusters with 6+ sequences (total across all targets). **c**, Number of clusters for which 0, 1, 2, or 3+ targets have this cluster as prominent (>1% of NGS reads). **d**, Is the same as **C**, but instead of count, it's the % of clusters. **e**, The number of prominent clusters fore each target with varying maximum number of clusters. The maximum number of clusters (100–500) was varied to balance forcing unrelated variants together (with 100 clusters and 53 targets, there can be fewer than two prominent clusters unique to one target), putting each binder in its own cluster (>2,000 binders in this set), and the requirement for several (6+) unique sequences from one target in a cluster to identify its binding motif. We determined that 300 clusters provided an optimal balance, and the relevant parameters showed little variation between 200 and 500 clusters.

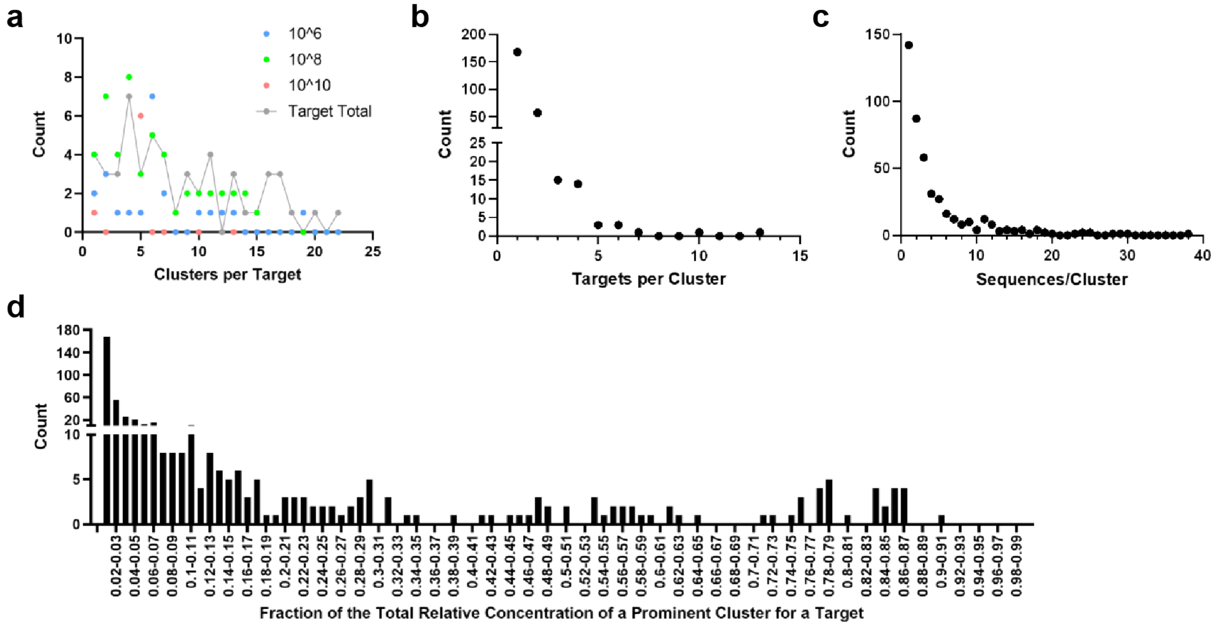

**Supplementary Fig. 13: Characteristics of Prominent Clusters. a**, Histogram of the number of prominent clusters per target for each different sampling size and the total for a target across sampling sizes. **b**, Histogram of the number of targets per prominent cluster. **c**, Histogram of the number of unique sequences in cluster-target pairs across sampling sizes. **d**, Fraction of NGS for a target for each prominent cluster. A total of 168 prominent clusters were specific to a single target, and 141 of these had >10% of NGS reads for that target (panels b, d). The mean number of unique sequences per cluster increased with sampling size (2.5 for $10^6$, 4.3 for $10^8$ and 4.2 for $10^{10}$; panel c), and only 93 target–cluster pairs across 29 targets had six or more unique sequences across the sampling sizes tested.

**Supplementary Table 3: Identified Binding Motifs.** X indicates not converged.

| Sampling Size[a] | | | | Randomization[b] | 4 | 4 | 15 | 6 | 9 | 15 | 2 | 15 | 15 | 8 | 15 | 8 | 15 | 3 | 15 | 15 | 2 | Motif |
|---|---|---|---|---|---|---|---|---|---|---|---|---|---|---|---|---|---|---|---|---|---|---|
| 10^6 | 10^8 | 10^10 | Target | Position[c] | 6 | 9 | 10 | 11 | 13 | 14 | 15 | 17 | 18 | 24 | 25 | 27 | 28 | 31 | 32 | 35 | 36 | Frequency[d] |
| 1 | 1 | | KIX | Cluster 0 | X | W | X | D | F/Y | I/V | X | C/S | X | X | X | X | X | V | X | X | X | 5.49E-05 |
| 1 | 1 | | KIX | Cluster 1 | X | W | X | X | Y/F | I/C | E | C | X | X | X | X | X | V | X | X | N | 4.12E-05 |
| 1 | | | Mdm2 | Cluster 5 | X | W | X | X | S/T | X | Q | L | X | X | X | E | X | V | X | X | X | 4.75E-05 |
| | 1 | | PIK3CB | Cluster 6 | Y | W | X | X | Y/F | X | E | X | X | X | X | K | X | X | X | X | X | 5.34E-04 |
| 1 | 1 | | p65 str | Cluster 7 | N | W | X | Y | F | F/Y | X | X | X | X | X | V | X | X | X | F | X | 7.91E-07 |
| | 1 | | KIX | Cluster 8 | S | W | X | X | I/F | I/L | X | V/L | F/C | X | X | X | X | I | S/A | X | N | 7.32E-07 |
| | 1 | | RNF8 | Cluster 9 | X | X | X | Y | I | F | Q | D | X | X | X | K | X | X | X | V | D | 1.06E-07 |
| | 1 | | TBX20-DBD | Cluster 12 | X | X | X | D | I/A | F | E | X | X | G | F | X | N | I | G | V/I | X | 7.50E-09 |
| 1 | 1 | | GABARAP | Cluster 13 | X | X | D | X | F | D | Q | X | D | X | X | V/L | X | X | X | X | X | 2.53E-06 |
| | 1 | | mCherry | Cluster 14 | X | X | X | X | D | X | X | X | X | X | X | K/R | I/V | X | R | I | X | 1.01E-05 |
| 1 | 1 | | PIK3CB | Cluster 15 | Y | G | X | X | I | F | X | S/A | X | X | X | R/K | X | X | X | X | X | 4.75E-06 |
| | 1 | | HIF1a | Cluster 18 | S | X | X | Y | X | X | X | X | X | Q | G | I | N | V | V | R | X | 5.63E-09 |
| | 1 | | HPV-pE7 | Cluster 19 | X | X | X | X | V | V/F | X | D | X | R | X | R | X | I | X | G/C | X | 3.38E-07 |
| | 1 | | HIF1a | Cluster 20 | X | X | F | X | X | Y/F | Q | V | S | X | X | I/V | S/T | X | X | R | X | 2.70E-08 |
| | 1 | | PDE6D | Cluster 20 | X | X | F | X | Y/F | X | X | R | R | X | X | X | X | X | D/G | D | D | 2.93E-07 |
| 1 | 1 | 1 | SIX1 | Cluster 21 | X | X | X | X | X | X | X | L | D | X | X | V | F | X | X | X | X | 2.28E-05 |
| 1 | 1 | | KIX | Cluster 23 | X | W | X | D | F/Y | V | X | X | X | X | X | X | X | V | X | X | X | 2.06E-04 |
| 1 | 1 | 1 | IFNG | Cluster 24 | Y | X | Y/F | X | I | Y/F | X | F | X | E | X | V | X | V | X | F | X | 5.63E-09 |
| 1 | 1 | | PIK3CB | Cluster 26 | X | W | X | X | I/V | Y/F | X | F | X | G | X | I/V | X | L | X | X | X | 2.53E-06 |
| 1 | 1 | 1 | IFNG | Cluster 27 | X | X | X | X | V | X | X | H | X | X | X | K/R | X | X | F | F | D | 2.53E-06 |
| | 1 | | OLIG2-DBD | Cluster 28 | X | W | V | D | F | X | E | F | X | X | X | X | D | L | X | X | D | 7.03E-08 |
| | 1 | | 14-3-3z | Cluster 31 | X | X | X | X | X | X | E | X | S/F | X | D | X | X | X | R | X | N | 1.48E-04 |
| | 1 | | MYC-DBD | Cluster 34 | S | V | L/F | X | F | D | X | A/V | X | X | X | R/K | X | X | D | X | X | 8.44E-08 |
| | 1 | | BAX | Cluster 37 | X | L | X | D | Y | F/Y | X | F | X | X | X | X | X | X | X | S/T | X | 5.49E-06 |
| | 1 | 1 | RAF | Cluster 38 | X | W | X | X | D | T | X | X | X | D | X | K/R | A/V | I | X | Y | X | 8.44E-08 |
| 1 | 1 | | MDM2 | Cluster 39 | Y | W | X | X | X | X | X | X | X | X | X | G/E | X | X | X | X | X | 9.62E-03 |
| 1 | 1 | | PIK3CB | Cluster 41 | Y | X | X | X | F/Y | F/Y | X | R | X | X | X | R | Y/F | X | X | V | X | 3.38E-07 |
| | 1 | | SH3BP5 | Cluster 43 | X | L | X | X | X | X | E | F | G/S | E | X | X | X | X | X | V/A | N | 7.41E-06 |
| | | 1 | SIX1 | Cluster 45 | X | X | X | X | F | D | X | Y/F | D | X | X | K/R | V | V | C | X | D | 3.75E-09 |
| 1 | | | XIAP | Cluster 48 | S | L | Y | N | A | Y | Q | R | I | X | X | X | X | X | X | X | X | 1.14E-08 |
| 1 | 1 | | MYC-DBD | Cluster 50 | Y | V | F | X | F | D | E | X | X | X | X | X | D | X | X | I | N | 3.43E-08 |
| 1 | 1 | | PIK3CB | Cluster 51 | X | W | X | X | Y | F | E | X | X | X | X | K/R | X | X | X | X | N | 7.12E-05 |
| 1 | 1 | | PIK3CB | Cluster 52 | X | L | X | X | F | F/Y | Q | R | S | X | X | X | X | X | X | V/I | X | 1.10E-06 |
| | 1 | | KIX | Cluster 53 | Y | W | X | X | I | X | E | C | X | X | X | E | X | X | X | F | D | 5.94E-07 |
| | 1 | | HPV-pE7 | Cluster 54 | X | X | X | D | V | X | X | X | P | X | X | R | R | X | F/Y | X | X | 8.44E-07 |
| | 1 | | ZB | Cluster 55 | T | L | X | X | I/V | X | X | X | F/Y | L/V | X | V/M | X | L | D | F | X | 8.44E-08 |
| | 1 | | HIF1a | Cluster 57 | X | X | X | X | X | V/I | X | I/L | X | X | X | L/I | X | I | R | R | N | 2.03E-06 |
| 1 | 1 | | mCherry | Cluster 60 | X | X | X | X | D | X | X | X | X | X | X | K/R | I | L | R | I/V | N | 1.69E-06 |
| 1 | 1 | | XIAP | Cluster 61 | N | L | F | X | V/A | X | E | X | X | X | X | K/R | X | V | Y | D | X | 1.06E-07 |
| 1 | 1 | | Myc-DBD | Cluster 64 | X | V | L/F | X | F | D | X | X | X | X | X | R | V/L | X | D | X | D | 8.44E-08 |
| 1 | 1 | | PIK3CB | Cluster 65 | Y | L | X | X | Y | X | X | R | X | X | X | X | C | X | G/S | V | D | 1.37E-07 |
| | 1 | | PDE6D | Cluster 66 | Y | L/V | V/I | X | X | R | X | R | X | X | X | K | X | X | D | D | D | 1.27E-08 |
| | 1 | | HPV-pE7 | Cluster 68 | X | X | X | X | V | X | E | X | X | R | X | R | R | V | F | X | X | 6.33E-07 |
| | 1 | | WWP2 | Cluster 74 | X | W/L | X | S/T | X | X | X | C | X | A | X | G | V/I | L | F | X | X | 2.53E-07 |
| | 1 | | CDKN1A | Cluster 75 | Y | W | Y | S | Y | N | E | A | D | X | X | K | X | X | X | X | X | 7.03E-10 |

[a] If "1" then that sampling size selection for that target had at least one unique sequence that clustered into that cluster.
[b] Number of possible amino acids in the degenerate codon in the randomization design.
[c] Position within the affibody scaffold.
[d] Calculated Motif frequency (same calculation as in Supplementary Fig. 10).

| Sampling Size [a] | | | | Randomization [b] | 4 | 4 | 15 | 6 | 9 | 15 | 2 | 15 | 15 | 8 | 15 | 8 | 15 | 3 | 15 | 15 | 2 | Motif |
|---|---|---|---|---|---|---|---|---|---|---|---|---|---|---|---|---|---|---|---|---|---|---|
| 10^6 | 10^8 | 10^10 | Target | Position [c] | 6 | 9 | 10 | 11 | 13 | 14 | 15 | 17 | 18 | 24 | 25 | 27 | 28 | 31 | 32 | 35 | 36 | Frequency [d] |
| 1 | 1 | | KIX | Cluster 76 | S | W | X | D | F/Y | I/F | E | X | X | X | X | X | H | X | X | I | X | 6.86E-07 |
| | 1 | | SH3BP5 | Cluster 80 | X | W/L | V/I | Y | V | X | X | F | X | E | X | G | X | L | R | X | X | 1.41E-08 |
| 1 | 1 | | GABARAP | Cluster 83 | N | W | X | X | F | D | Q | X | D | X | X | X | X | I | X | X | D | 2.57E-06 |
| 1 | 1 | | p65(str) | Cluster 86 | T | X | R | X | F | F/Y | E | X | X | X | X | V/L | X | L | X | F | D | 2.11E-07 |
| | 1 | | LMO2 | Cluster 95 | T | V | C | Y | Y | I | E | V | Y | X | X | X | X | X | X | X | X | 1.14E-08 |
| 1 | | | GABARAP | Cluster 97 | N | V | F | T | I | T | E | T | S | X | X | X | X | X | X | X | X | 1.14E-08 |
| 1 | 1 | | GABARAP | Cluster 98 | X | X | X | X | F | D | E | X | D | X | X | X | X | X | X | X | D | 1.23E-04 |
| | | 1 | NRAS | Cluster 99 | Y | V | F | Y | N | R | E | F | D | X | X | X | X | I | X | L | X | 2.54E-10 |
| | 1 | | FRB | Cluster 101 | X | X | C/I | X | A | S/A | X | F/Y | X | X | X | K/R | X | V | X | Y | X | 9.00E-07 |
| | 1 | | mCherry | Cluster 103 | X | X | X | X | X | X | X | F | F/Y | V | X | V | Y/F | X | X | D | X | 6.08E-07 |
| 1 | 1 | 1 | IFNG | Cluster 104 | X | L/V | X | X | V | X | X | H | X | X | X | R | X | I | F | F | D | 2.11E-07 |
| | 1 | | mCherry | Cluster 106 | X | X | X | X | D | X | Q | X | X | X | X | K | V/I | L | R | I/L | N | 8.44E-07 |
| 1 | 1 | | Mdm2 | Cluster 109 | Y | W | I/L | D | X | X | X | I/L | X | X | X | G | X | X | X | X | X | 1.14E-05 |
| | 1 | | HPV-pE7 | Cluster 110 | X | X | X | Y | V | X | Q | D | X | X | X | R/G | R | V | F/Y | X | D | 1.41E-07 |
| | 1 | | PIK3CB | Cluster 111 | X | X | Y | D | Y/F | F | E | S | X | X | X | R | I | I | X | X | X | 9.38E-09 |
| 1 | 1 | | PIK3CB | Cluster 112 | Y | V/L | X | X | Y | X | X | R | X | X | X | X | G/C | X | S | I/V | D | 5.49E-07 |
| | 1 | | Myc-DBD | Cluster 113 | X | V | X | X | F | X | X | X | X | L/V | X | K | D | X | X | I | X | 1.90E-06 |
| 1 | 1 | 1 | IFNG | Cluster 114 | Y | X | X | X | I/V | F/Y | X | F | X | E | X | V/I | X | X | X | F | X | 5.06E-07 |
| | 1 | 1 | IFNG | Cluster 115 | X | L/V | X | D | I/V | F/Y | X | F | X | E | X | I/V | X | V | V | F | X | 3.75E-09 |
| | | 1 | IFNG | Cluster 119 | X | L | X | X | I | F | X | F | X | E | X | V | V/I | V | I/C | F | D | 1.88E-10 |
| | 1 | | mCherry | Cluster 121 | X | X | X | X | D | X | X | X | X | X | X | K/R | I | X | R | X | N | 3.80E-05 |
| | 1 | 1 | IFNG | Cluster 123 | X | L | X | X | I | Y | X | F | X | E/D | X | I/V | V/C | V | V/C | F | D | 7.50E-10 |
| | 1 | | RAF | Cluster 129 | X | X | X | X | A | D | E | F | I/L | D | X | V | X | X | X | R | X | 1.69E-08 |
| | 1 | | THAP1 | Cluster 130 | N | X | D | Y | X | X | E | F | C | X | X | X | I/V | I | X | X | D | 1.37E-07 |
| 1 | 1 | | KIX | Cluster 131 | X | W | I | D | Y/F | V/I | E | C | X | X | X | X | X | V | X | X | D | 4.57E-07 |
| | 1 | | RNF8 | Cluster 134 | X | W | V/I | X | N | X | X | H | X | X | X | K/R | C/A | X | I/V | X | X | 6.75E-07 |
| 1 | 1 | 1 | IFNG | Cluster 136 | X | X | X | X | I | X | X | H | Y/F | X | X | R/K | X | X | F | F | X | 6.75E-07 |
| | 1 | | mCherry | Cluster 138 | X | X | X | N | D | X | X | X | X | X | X | K | I | X | R | X | X | 6.33E-06 |
| 1 | 1 | | KIX | Cluster 146 | Y | W/L | I | X | F/Y | X | E | X | F/L | X | X | X | X | X | D | X | X | 8.23E-06 |
| 1 | 1 | | FRB | Cluster 148 | N | X | F | X | X | C | X | X | G | X | X | R | X | X | X | X | X | 5.70E-06 |
| 1 | 1 | | PIK3CB | Cluster 149 | X | L | X | X | Y | X | X | R | X | X | X | R | X | I | S | I/V | X | 4.22E-07 |
| | 1 | | LMO2 | Cluster 168 | X | G | I | Y | I | S | Q | V | R | X | X | X | X | X | X | X | X | 4.57E-08 |
| | 1 | | LC3B | Cluster 198 | X | X | D | D | X | F | E | L | D | X | D | X | X | X | X | X | N | 6.10E-08 |
| | 1 | | LC3B | Cluster 202 | X | X | X | D | X | F | E | L | D | X | X | E/G | X | X | X | X | X | 3.04E-06 |
| 1 | 1 | | GABARAP | Cluster 204 | X | X | X | X | I | X | X | S/G | X | X | X | X | Y | L | X | Y/F | N | 2.19E-05 |
| | 1 | | FRB | Cluster 206 | N | X | F | X | X | C | X | X | G | X | X | R | X | X | X | X | N | 2.85E-06 |
| 1 | 1 | | GABARAP | Cluster 208 | X | X | Y | D | D | F | E | V | I/V | X | X | X | X | X | X | X | X | 3.66E-07 |
| | 1 | | OTUB1 | Cluster 216 | Y | W | D | X | D | Y | E | F | I | X | X | X | X | X | X | X | X | 6.86E-08 |
| 1 | 1 | | PIK3CB | Cluster 226 | Y | V | X | X | Y | X | X | R | X | X | X | R | X | X | X | V/A | X | 4.75E-06 |
| | 1 | | FRB | Cluster 232 | X | G | X | X | I/V | S | X | F/Y | X | D/E | X | K/R | X | V | I/V | F | X | 4.50E-08 |
| | | 1 | NRAS | Cluster 233 | N | L | S | S | A | S | Q | F | X | X | X | X | X | I | X | X | X | 3.52E-09 |
| 1 | | | PIK3CB | Cluster 262 | X | L | Y | D | X | F | E | X | A | X | X | R/K | Y/V | X | X | X | N | 6.33E-08 |
| | 1 | | WWP2 | Cluster 268 | Y | W | X | X | V/I | D | X | C/T | X | D | X | K/R | D | X | X | F | X | 8.44E-09 |
| | 1 | | GABARAP | Cluster 280 | X | X | X | D | D | F | E | I | D | X | X | X | X | X | X | X | X | 2.74E-06 |
| 1 | 1 | | PIK3CB | Cluster 290 | X | L | X | N | I/V | Y/F | X | F | I/C | G | X | V/I | X | I | X | X | X | 5.63E-08 |
| 1 | 1 | | mCherry | Cluster 292 | X | X | X | X | D | X | E | X | X | A/G | S/G | K | V | L | R | X | X | 1.69E-07 |
| 1 | 1 | | KIX | Cluster 294 | S | X | X | X | F | X | X | C/A | X | L | X | X | X | V | X | L/I | D | 8.23E-06 |
| | 1 | | FRB | Cluster 298 | X | X | X | X | X | X | X | D | X | V | G | K/R | I | I | X | X | N | 7.60E-07 |

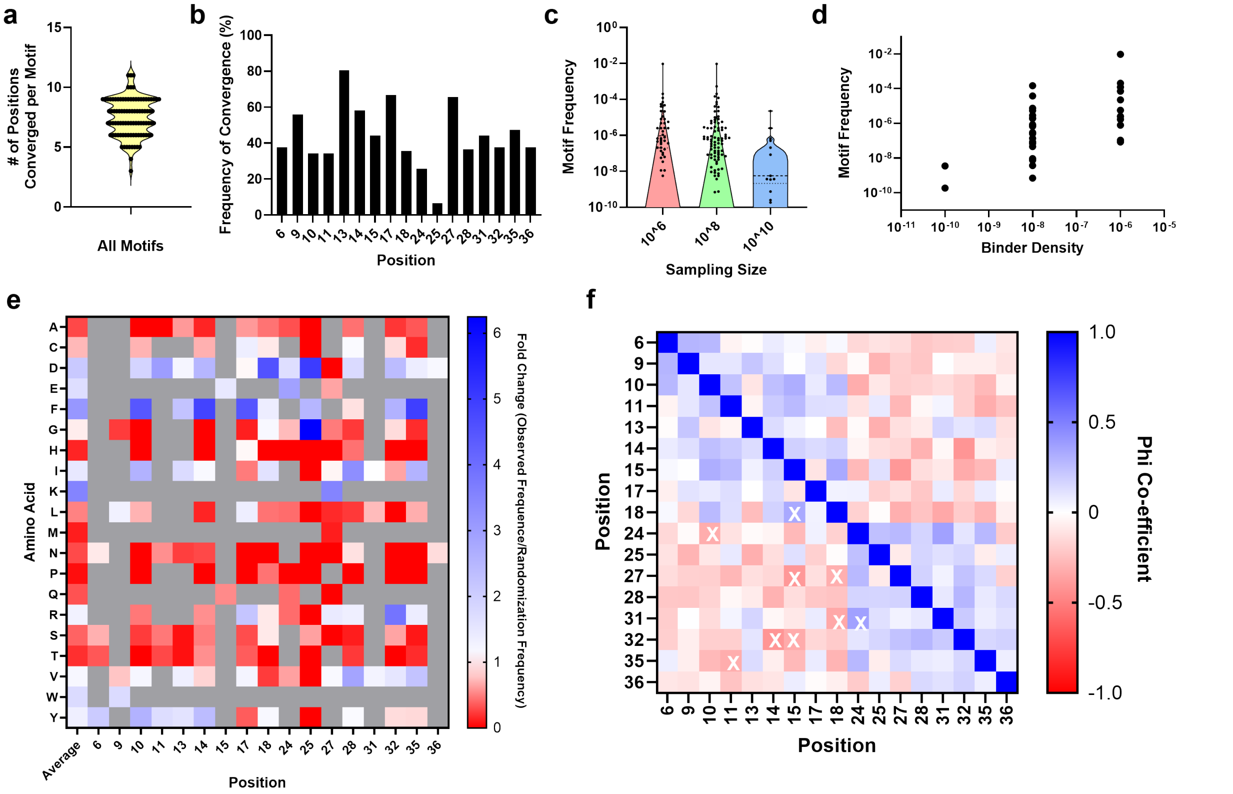

**Supplementary Fig. 14: Characterization of Binding Motifs. a**, The number of converged positions per binding motif (cluster). **b**, The frequency (%) that each position converged among the binding motifs (clusters). **c**, The motif frequency for each binding motif; as long as one variant from a selection clustered into that target-motif paring, then a point is shown for that selection under its sampling size. **d**, Binder Density vs Motif Frequency for the highest frequency motif from each target (separated by binder density). This was used for the Pearson's Correlation referred to in the main text. **e**, Fold change in the observed frequency over the randomization design frequency for each amino acid at each position (grey indicates that it wasn't not coded in the randomization frequency). **f**, Analysis of co-convergence between positions (Phi Co-efficient) where 1 indicates always co-occurs and -1 indicates mutual exclusivity; the white X's indicate pairings with strong statistical reliability (Fisher's exact test with a Benjamini-Hochberg FDR correction (motifs vary in how many positions converge) $q < 0.05$). Each position converged with a different frequency, from 6% to 81% of motifs (44% on average; SD 17%; panel b). The amino acid distribution within binding motifs varied strongly by position (panel e; for example, Gly is strongly de-enriched and Tyr strongly enriched at the position 10 NNY, while the inverse is observed for the NNY at position 25); aromatic and charged residues are highly enriched in binding motifs, while small and polar residues are strongly de-enriched. Because motifs were identified by convergence, these differences may reflect the greater interchangeability of small residues at binding interfaces rather than their frequency there. Significant intra-helix co-convergence was observed (panel f; positions 19 and 26 showed the highest co-convergence, and positions 9 and 27 the highest mutual exclusivity).

**Supplementary Table 4: Additional Library Construction.**

| Design | 40 min titer (PFU/mL) | 80 min titer (PFU/mL) | 120 min titer (PFU/mL) | Estimated number of variants | Amplified titer (PFU/mL) |
|---|---|---|---|---|---|
| **A** | 2.00E+08 | 3.25E+09 | 3.50E+09 | 1.73E+09 | 2.24E+10 |
| **B** | 1.25E+07 | 4.38E+07 | 7.50E+07 | 3.75E+09 | 2.75E+10 |
| **C** | 1.27E+07 | 2.70E+07 | 8.90E+07 | 4.45E+09 | 2.13E+11 |
| **D** | 1.75E+07 | 6.88E+07 | 2.50E+08 | 1.25E+10 | 2.21E+10 |
| **E** | 8.00E+05 | 6.20E+06 | 8.90E+08 | 4.45E+08 | 3.98E+11 |
| **F** | 5.00E+06 | 3.75E+07 | 7.50E+06 | 3.75E+09 | 6.81E+10 |
| **G** | 5.00E+05 | 1.30E+06 | 4.00E+06 | 2.00E+08 | 3.13E+11 |
| **H** | 4.70E+04 | 2.00E+05 | 2.00E+05 | 1.00E+07 | 6.88E+11 |
| **I** | 8.00E+05 | 1.80E+06 | 3.50E+06 | 1.75E+08 | 2.06E+12 |
| **J** | 1.50E+06 | 1.73E+06 | 2.93E+06 | 7.50E+07 | 1.75E+10 |

**Supplementary Table 5: Amino Acids possible at Each Position**

| Design | Description | Theoretical number unique variants |
|---|---|---|
| **A** | NNY; biased codons | 2.39E+15 |
| ***B*** | NNY only | 1.95E+15 |
| ***C*** | Single-helix NNY/NNK | 9.11E+10 |
| ***D*** | NNK only | 8.19E+16 |
| ***E*** | VBY only | 2.54E+12 |
| ***F*** | KMY only | 6.71E+07 |
| ***G*** | NNY; biased codons; L34A | 2.39E+15 |
| ***H*** | NNY; biased codons; L44G, A48L | 2.39E+15 |
| ***I*** | NNY; biased codons; L22G, L34A, A48G | 2.39E+15 |
| ***J*** | Full NNY random 30mer | 1.92E+35 |

| | **Amino acid positions 1-15** | | | | | | | | | | | | | | |
|---|---|---|---|---|---|---|---|---|---|---|---|---|---|---|---|
| **Design** | **1** | **2** | **3** | **4** | **5** | **6** | **7** | **8** | **9** | **10** | **11** | **12** | **13** | **14** | **15** |
| **A** | V | D | N | K | F | *WMY* | K | E | *KKG* | *NNY* | *DMY* | A | *DHY* | *NNY* | *SAR* |
| ***B*** | V | D | N | K | F | N | K | E | *NNY* | *NNY* | *NNY* | A | *NNY* | *NNY* | E |

| | | | | | | | | | | | | | | | |
|---|---|---|---|---|---|---|---|---|---|---|---|---|---|---|---|
| ***C*** | V | D | *NNY* | *NNY* | F | *NNY* | *NNK* | *NNY* | V | *NNY* | *NNK* | A | R | *NNK* | *NNY* |
| ***D*** | V | D | N | K | F | N | K | E | *NNK* | *NNK* | *NNK* | A | *NNK* | *NNK* | E |
| ***E*** | V | D | N | K | F | N | K | E | *VBY* | *VBY* | *VBY* | A | *VBY* | *VBY* | E |
| ***F*** | V | D | N | K | F | N | K | E | *KMY* | *KMY* | *KMY* | A | *KMY* | *KMY* | E |
| ***G*** | V | D | N | K | F | *WMY* | K | E | *KKG* | *NNY* | *DMY* | A | *DHY* | *NNY* | *SAR* |
| ***H*** | V | D | N | K | F | *WMY* | K | E | *KKG* | *NNY* | *DMY* | A | *DHY* | *NNY* | *SAR* |
| ***I*** | V | D | N | K | F | *WMY* | K | E | *KKG* | *NNY* | *DMY* | A | *DHY* | *NNY* | *SAR* |
| ***J*** | *NNY* | *NNY* | *NNY* | *NNY* | *NNY* | *NNY* | *NNY* | *NNY* | *NNY* | *NNY* | *NNY* | *NNY* | *NNY* | *NNY* | *NNY* |

| | **Amino acid positions 16-30** | | | | | | | | | | | | | | |
|---|---|---|---|---|---|---|---|---|---|---|---|---|---|---|---|
| **Design** | **16** | **17** | **18** | **19** | **20** | **21** | **22** | **23** | **24** | **25** | **26** | **27** | **28** | **29** | **30** |
| **A** | I | *NNY* | *NNY* | L | P | N | L | N | *SNS* | *NNY* | Q | *VDR* | *NNY* | A | F |
| ***B*** | I | *NNY* | *NNY* | L | P | N | L | N | *NNY* | *NNY* | Q | *NNY* | *NNY* | A | F |
| ***C*** | I | *NNY* | *NNY* | L | P | N | L | N | R | E | Q | G | G | A | F |
| ***D*** | I | *NNK* | *NNK* | L | P | N | L | N | *NNK* | *NNK* | Q | *NNK* | *NNK* | A | F |
| ***E*** | I | *VBY* | *VBY* | L | P | N | L | N | *VBY* | *VBY* | Q | *VBY* | *VBY* | A | F |
| ***F*** | I | *KMY* | *KMY* | L | P | N | L | N | *KMY* | *KMY* | Q | *KMY* | *KMY* | A | F |
| ***G*** | I | *NNY* | *NNY* | L | P | N | L | N | *SNS* | *NNY* | Q | *VDR* | *NNY* | A | F |
| ***H*** | I | *NNY* | *NNY* | L | P | N | L | N | *SNS* | *NNY* | Q | *VDR* | *NNY* | A | F |
| ***I*** | I | *NNY* | *NNY* | L | P | N | G | N | *SNS* | *NNY* | Q | *VDR* | *NNY* | A | F |
| ***J*** | *NNY* | *NNY* | *NNY* | *NNY* | *NNY* | *NNY* | *NNY* | *NNY* | *NNY* | *NNY* | *NNY* | *NNY* | *NNY* | *NNY* | *NNY* |

| | **Amino acid positions 31-45** | | | | | | | | | | | | | | |
|---|---|---|---|---|---|---|---|---|---|---|---|---|---|---|---|
| **Design** | **31** | **32** | **33** | **34** | **35** | **36** | **37** | **38** | **39** | **40** | **41** | **42** | **43** | **44** | **45** |
| **A** | *VTH* | *NNY* | S | L | *NNY* | *RAY* | D | P | S | Q | S | A | N | L | L |
| ***B*** | I | *NNY* | S | L | *NNY* | D | D | P | S | Q | S | A | N | L | L |
| ***C*** | I | V | S | L | W | D | D | P | S | Q | S | A | N | L | L |

| | | | | | | | | | | | | | | | |
|---|---|---|---|---|---|---|---|---|---|---|---|---|---|---|---|
| ***D*** | I | *NNK* | S | L | *NNK* | D | D | P | S | Q | S | A | N | L | L |
| ***E*** | I | *VBY* | S | L | *VBY* | D | D | P | S | Q | S | A | N | L | L |
| ***F*** | I | *KMY* | S | L | *KMY* | D | D | P | S | Q | S | A | N | L | L |
| ***G*** | *VTH* | *NNY* | S | A | *NNY* | *RAY* | D | P | S | Q | S | A | N | L | L |
| ***H*** | *VTH* | *NNY* | S | L | *NNY* | *RAY* | D | P | S | Q | S | A | N | G | L |
| ***I*** | *VTH* | *NNY* | S | A | *NNY* | *RAY* | D | P | S | Q | S | A | N | L | L |

| | **Amino acid position 46-58** | | | | | | | | | | | | |
|---|---|---|---|---|---|---|---|---|---|---|---|---|---|
| **Design** | **46** | **47** | **48** | **49** | **50** | **51** | **52** | **53** | **54** | **55** | **56** | **57** | **58** |
| **A** | A | E | A | K | K | L | N | D | A | Q | A | P | K |
| ***B*** | A | E | A | K | K | L | N | D | A | Q | A | P | K |
| ***C*** | A | E | A | K | K | L | N | D | A | Q | A | P | K |
| ***D*** | A | E | A | K | K | L | N | D | A | Q | A | P | K |
| ***E*** | A | E | A | K | K | L | N | D | A | Q | A | P | K |
| ***F*** | A | E | A | K | K | L | N | D | A | Q | A | P | K |
| ***G*** | A | E | A | K | K | L | N | D | A | Q | A | P | K |
| ***H*** | A | E | L | K | K | L | N | D | A | Q | A | P | K |
| ***I*** | A | E | G | K | K | L | N | D | A | Q | A | P | K |

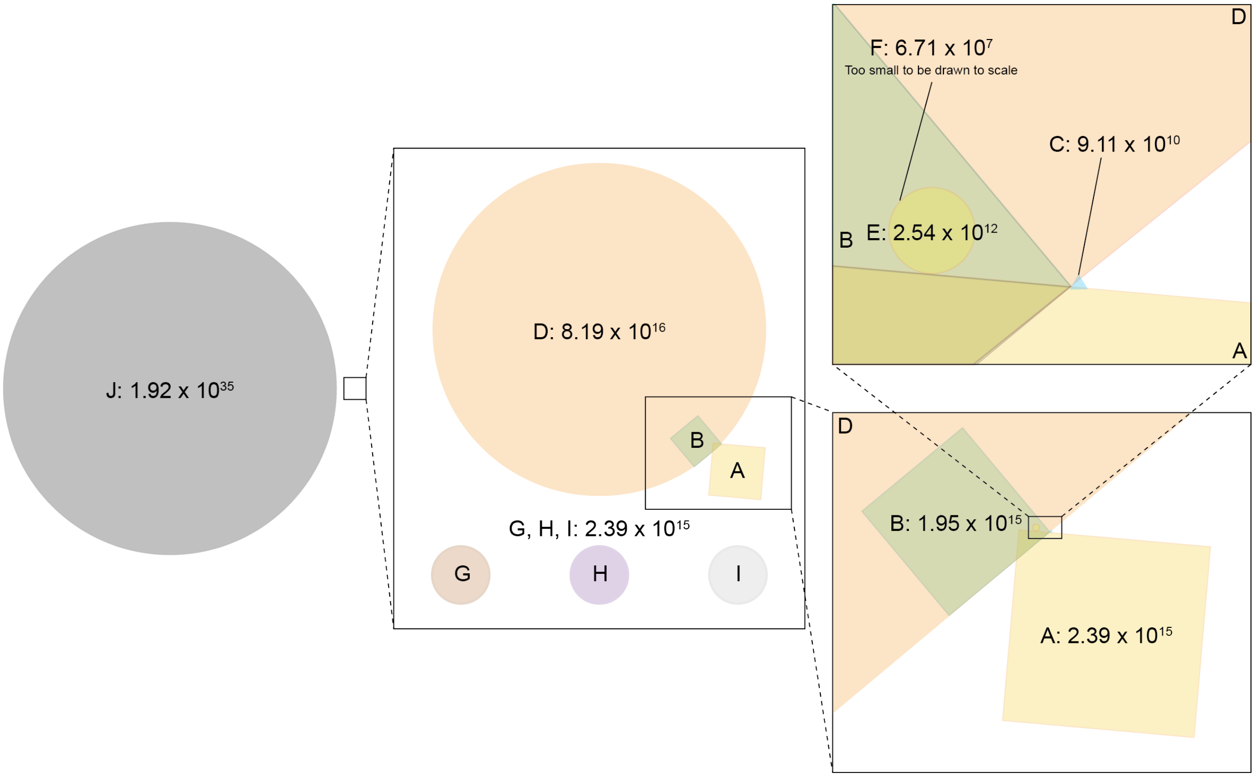

**Supplementary Fig. 15: Theoretical randomization design sequence space sizes and overlapping regions.** Calculations were performed by multiplying the number of common, possible amino acids at each position between designs of interest.

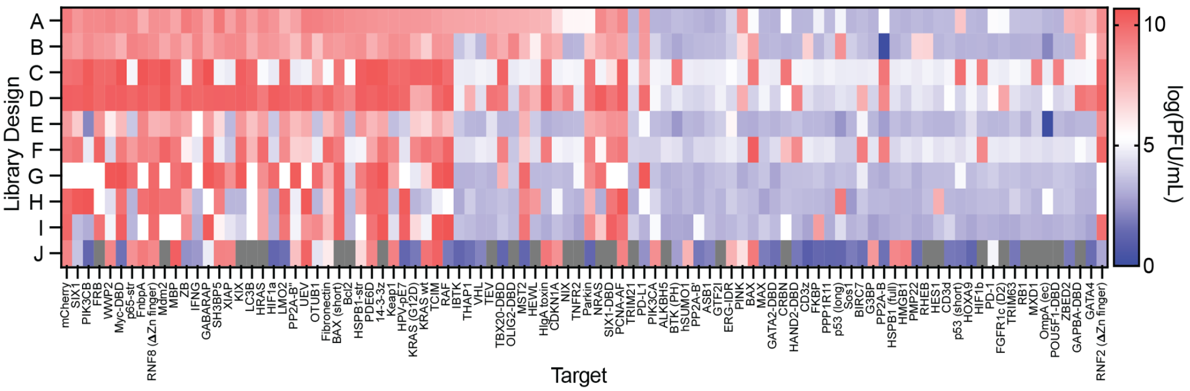

**Supplementary Fig. 16: Endpoint titer from all library selections at $10^8$ sampling size.** PANCS was performed on the 96-target panel (**Supplementary Table 1**) with a single independent replicate of each library with a sampling size of ~$10^8$. After completion of the sixth passage, the titer was determined by qPCR. Red indicates higher titers, blue lower titers, and gray indicates that a selection was not performed for a target at that sample size. All selections with a titer >1 × $10^7$ PFU/mL were subject to NGS analysis and validation in the split T7 RNAP *E. coli* luciferase assay. Each titer was determined as a single replicate.

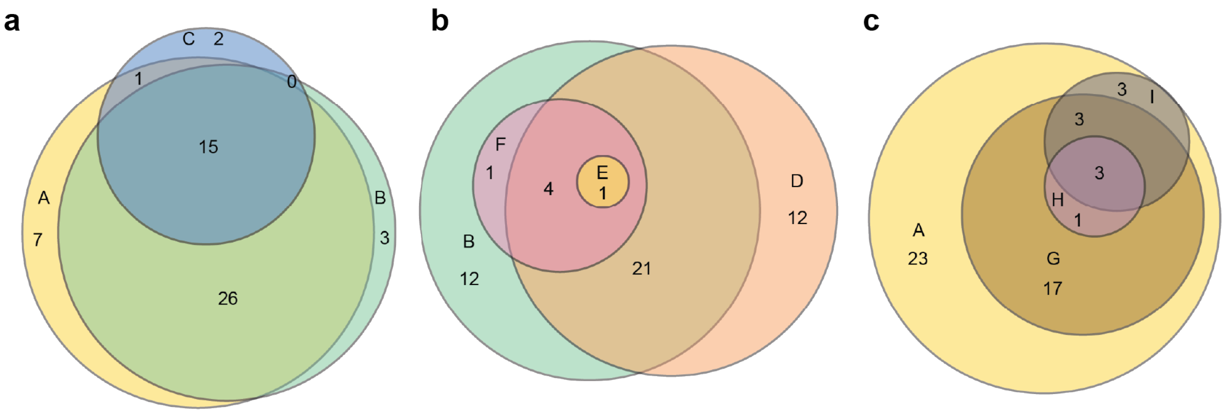

**Supplementary Fig. 17: Overlap in targets that hit between library designs. a**, Varied positions randomized. **b**, Varied amino acids available in the randomization via different codon degeneracies. **c**, Varied hydrophobic core stability via mutations to remove hydrophobic side chains in the affibody hydrophobic core.

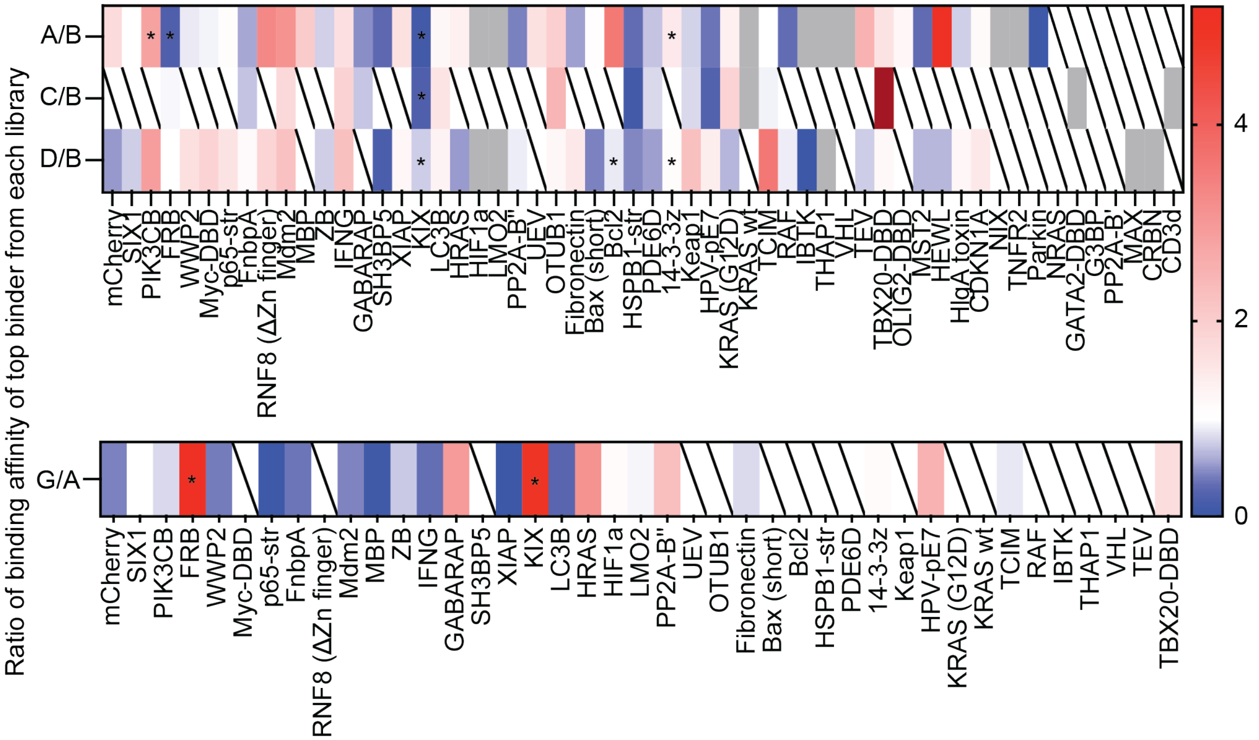

**Supplementary Fig. 18: Fold change in fitness (luciferase signal) between hits from different library designs. Top:** Red represents the binder from Designs A, C, or D outperforming that of B, and blue indicates that the design B binder was more fit. Boxes in gray represent targets that hit in one but not both designs compared, and boxes with a slash indicate no hits found in either design compared. White is set to 1.0, where the binding affinities of the two targets are comparable. **Bottom**: Same as top, but design G is compared to A rather than B.

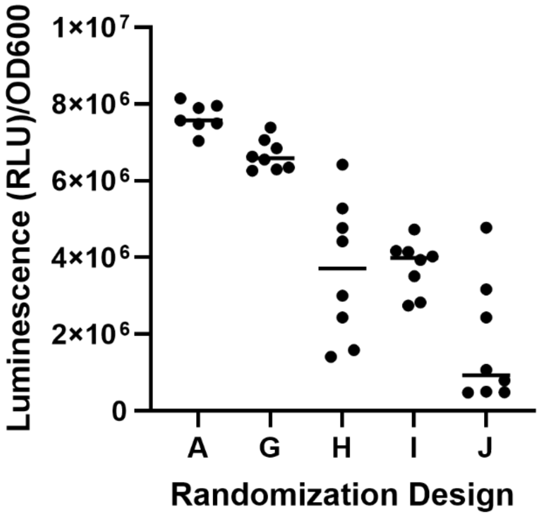

**Supplementary Fig. 19: Expression level of random variants from select randomization designs.** Individual randomization design variants were subcloned from the library phage into $RNAP_{N,WT}$ fusions for testing in the binding luciferase assay with ZB-$RNAP_C$ (the $RNAP_{N,WT}$ recombines with $RNAP_C$ independent of binding, thus this reports on expression). N =8, bar indicates mean.

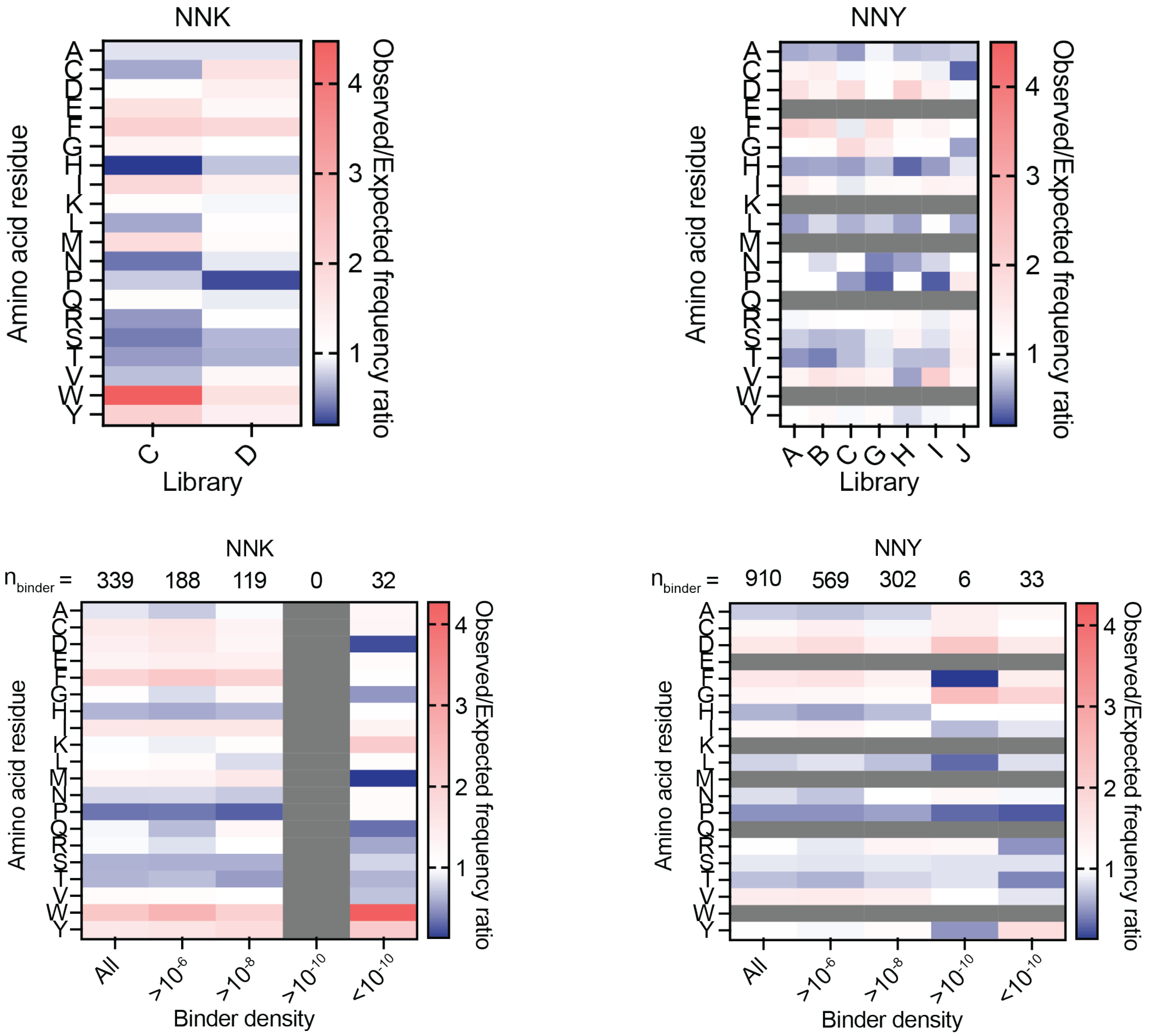

**Supplementary Fig. 20: Position-independent amino acid usage of binders discovered from different samplings.** To determine the position-dependent amino acid usages, the percent difference for position-dependent amino acid compositions were calculated, then an observed frequency to expected frequency ratio was calculated by taking the quotient of the aggregated frequencies for NNK and NNY codons, for each individual design.

**Supplementary Table 6: Targets sorted into binder density categories.**

| Rank | Target | Binder density in design A | Number of designs hit | Titer of hit in smallest sampling with design A |
|---|---|---|---|---|
| 1 | LC3B | 1.00E-06 | 8 | 1.26E+09 |
| 2 | GABARAP | 1.00E-06 | 7 | 2.37E+09 |
| 3 | KIX | 1.00E-06 | 7 | 1.39E+09 |
| 4 | Mdm2 | 1.00E-06 | 7 | 3.23E+09 |
| 5 | SIX1 | 1.00E-06 | 5 | 7.24E+09 |
| 6 | FRB | 1.00E-06 | 5 | 6.32E+09 |
| 7 | FnbpA | 1.00E-06 | 5 | 3.79E+09 |
| 8 | IFNG | 1.00E-06 | 5 | 2.37E+09 |
| 9 | mCherry | 1.00E-06 | 5 | 1.45E+10 |
| 10 | PIK3CB | 1.00E-06 | 4 | 6.39E+09 |
| 11 | WWP2 | 1.00E-06 | 4 | 5.50E+09 |
| 12 | p65-str | 1.00E-06 | 4 | 5.20E+09 |
| 13 | MBP | 1.00E-06 | 4 | 3.10E+09 |
| 14 | ZB | 1.00E-06 | 4 | 2.95E+09 |
| 15 | XIAP | 1.00E-06 | 4 | 1.46E+09 |
| 16 | HRAS | 1.00E-06 | 4 | 5.79E+08 |
| 17 | PP2A-B'' | 1.00E-06 | 4 | 8.46E+07 |
| 18 | Myc-DBD | 1.00E-06 | 3 | 5.43E+09 |
| 19 | RNF8 (ΔZn finger) | 1.00E-06 | 3 | 3.75E+09 |
| 20 | SH3BP5 | 1.00E-06 | 3 | 2.00E+09 |
| 21 | HIF1a | 1.00E-06 | 3 | 3.47E+08 |
| 22 | LMO2 | 1.00E-06 | 3 | 3.33E+08 |
| 23 | HPV-pE7 | 1.00E-08 | 8 | 1.91E+09 |
| 24 | 14-3-3z | 1.00E-08 | 6 | 1.98E+09 |
| 25 | Fibronectin | 1.00E-08 | 6 | 4.26E+09 |
| 26 | OTUB1 | 1.00E-08 | 5 | 5.53E+09 |
| 27 | HSPB1-str | 1.00E-08 | 5 | 2.74E+09 |
| 28 | TCIM | 1.00E-08 | 5 | 1.40E+09 |
| 29 | RAF | 1.00E-08 | 5 | 1.07E+09 |
| 30 | TBX20-DBD | 1.00E-08 | 5 | 6.69E+08 |
| 31 | PDE6D | 1.00E-08 | 5 | 2.28E+09 |
| 32 | Keap1 | 1.00E-08 | 4 | 1.91E+09 |
| 33 | KRAS (G12D) | 1.00E-08 | 4 | 1.82E+09 |
| 34 | BAX (short) | 1.00E-08 | 3 | 3.72E+09 |
| 35 | Bcl2 | 1.00E-08 | 3 | 2.98E+09 |
| 36 | TEV | 1.00E-08 | 3 | 8.30E+08 |
| 37 | MST2 | 1.00E-08 | 3 | 5.74E+08 |

| | | | | |
|---|---|---|---|---|
| 38 | HEWL | 1.00E-08 | 3 | 4.89E+08 |
| 39 | HlgA toxin | 1.00E-08 | 3 | 4.12E+08 |
| 40 | CDKN1A | 1.00E-08 | 3 | 5.51E+07 |
| 41 | UEV | 1.00E-08 | 3 | 6.07E+09 |
| 42 | KRAS WT | 1.00E-08 | 2 | 1.73E+09 |
| 43 | IBTK | 1.00E-08 | 2 | 1.03E+09 |
| 44 | THAP1 | 1.00E-08 | 2 | 9.48E+08 |
| 45 | OLIG2-DBD | 1.00E-08 | 2 | 6.29E+08 |
| 46 | Parkin | 1.00E-08 | 2 | 1.65E+07 |
| 47 | VHL | 1.00E-08 | 1 | 9.29E+08 |
| 48 | NIX | 1.00E-08 | 1 | 1.78E+07 |
| 49 | TNFR2 | 1.00E-08 | 1 | 1.73E+07 |
| 50 | NRAS | 1.00E-10 | 1 | 2.12E+10 |
| 51 | GATA2-DBD | 1.00E-10 | 1 | 2.56E+09 |
| 52 | G3BP | 1.00E-10 | 1 | 1.07E+09 |
| 53 | PP2A-B' | 1.00E-10 | 1 | 1.21E+07 |
| 54 | CRBN | <1.00E-10 | 1 | 0 |
| 55 | CD3d | <1.00E-10 | 1 | 0 |
| 56 | MAX | <1.00E-10 | 1 | 0 |
| 57 | SIX1-DBD | 1.00E-10 | 0 | 2.08E+10 |
| 58 | PCNA-AF | 1.00E-10 | 0 | 9.67E+09 |
| 59 | TRIM21 | 1.00E-10 | 0 | 1.63E+09 |
| 60 | PD-L1 | 1.00E-10 | 0 | 1.00E+08 |
| 61 | ALKBH5 | <1.00E-10 | 0 | 6.95E+06 |
| 62 | BTK (PH) | <1.00E-10 | 0 | 6.95E+06 |
| 63 | hSUMO1 | <1.00E-10 | 0 | 5.30E+06 |
| 64 | ASB1 | <1.00E-10 | 0 | 5.05E+05 |
| 65 | GTF2I | <1.00E-10 | 0 | 1.13E+05 |
| 66 | ERG-IDR | <1.00E-10 | 0 | 6.77E+04 |

The following 30 targets were not assigned a difficulty ranking because they all share the following in common - no hit discovered in any $10^8$-sized sampling, nor in a $10^{10}$-sized sampling. In some cases, the titer was too low for detection, or the putative hit was a false positive in luciferase assay - MXD1, FGFR1c (D2), TRIM63, ZBED2, PMP22, p53 (short), p53 (long), BIRC7, HES1, HIF1b, CD3z, POU5F1-DBD, HSPB1, HOXA9, PP2A-B, HAND2-DBD, PIK3CA, GATA4, GAPBA-DBD, Sos1, RHEB, PINK, BAX, FKBP, PPP1R11, HMGB1, PD-1, RB1, OmpA (ec), RNF2 (**Δ** Zn finger).

**Supplementary Table 7: Grouping target panel by broad functional/structural features.**

| Class | Kinases and phosphatases | E3 ligases | Transcription factors | EC domains of membrane proteins | Secreted | Disordered (>50%) | a-helical (>30%) | B-sheet (>30%) |
|---|---|---|---|---|---|---|---|---|
| Targets | KRAS (G12D), RAF, NRAS, HRAS, KRAS wt, PP2A-B, PP2A-B', PP2A-B", BTK (PH), PIK3CB, PIK3CA, FRB, MST2, RHEB | TRIM21, Parkin, Mdm2, VHL, CRBN, PPP1R11, TRIM63, RNF2, RNF8, ASB1, BIRC7, XIAP, WWP2 | p65-str, KIX, Myc-DBD, LMO2, HIF1a, MXD1, HIF1b, p53 (long), p53 (short), MAX, SIX1-DBD, SIX1, ERG-IDR, HAND2-DBD, HES1, HOXA9, GATA2-DBD, POU5F1-DBD, THAP1, TBX20-DBD, OLIG2-DBD, ZBED2, GAPBA-DBD, GATA4 | FGFR1c (D2), Bcl2, NIX, TNFR2, PINK, RB1, OmpA (ec), PD-1, PD-L1, PMP22 | HlgA toxin, Fibronectin, FnbpA, HEWL, IFNG | TCIM, NIX, Sos1, PCNA-AF, IBTK, CDKN1A, RB1, CD3z, HPV-pE7, MST2, PD-L1, BAX (short), p53 (short), PPP1R11, RNF2, SIX1-DBD, ERG-IDR, HES1, HOXA9, THAP1, ZBED2, GATA4 | PP2A-B', HES1, MXD1, Bcl2, SIX1, OmpA (ec), ZB, PP2A-B", HEWL, KRAS (G12D), RHEB, KRAS wt, MAX, IFNG, MBP, SH3BP5, GTF2I, FRB, RNF8, TCIM, HRAS, SIX1-DBD, HMGB1, BAX (short), GAPBA-DBD, POU5F1-DBD, Myc-DBD, 14-3-3z, ASB1, PMP22, TRIM63, NRAS, HAND2-DBD, OLIG2-DBD, ZBED2, BAX, KIX, XIAP | Fibronectin, HlgA toxin, p65-str, TEV, PDE6D, FKBP, TRIM21, FGFR1c (D2), TBX20-DBD, CRBN, HSPB1-str, PP2A-B, FnbpA, BTK (PH), PD-L1, G3BP, mCherry, PD-1, Keap1 |
| **n** | 14 | 13 | 24 | 10 | 5 | 22 | 38 | 19 |

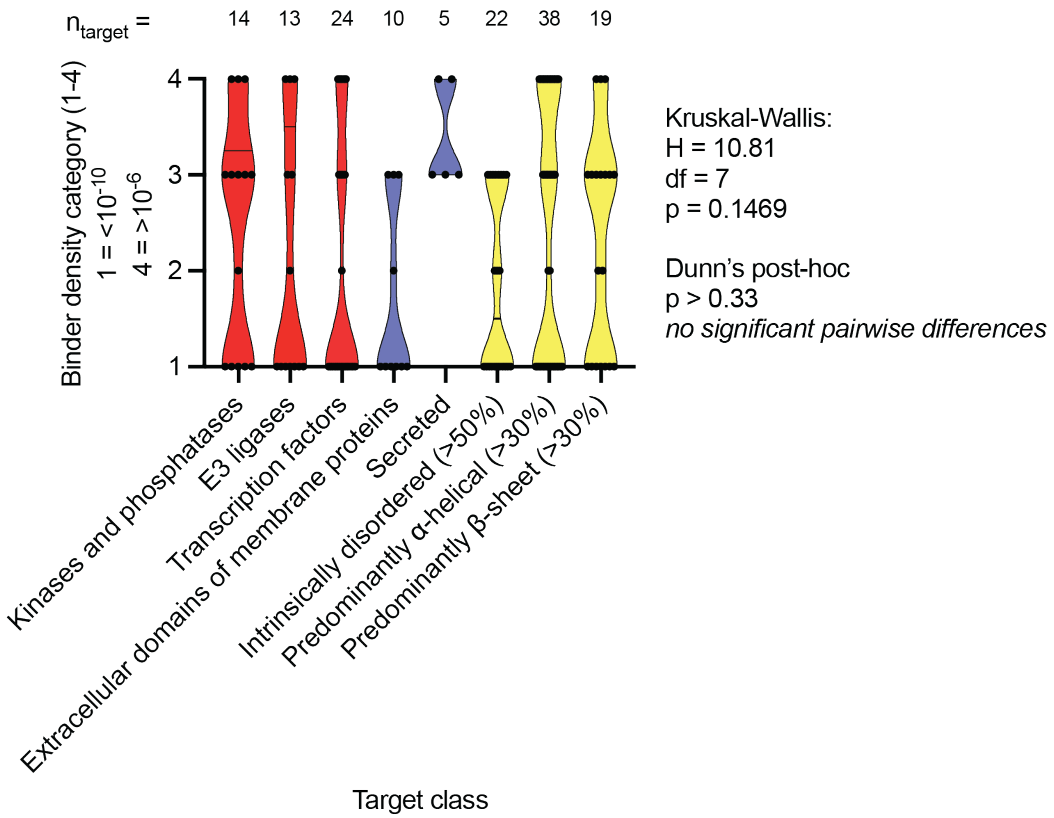

**Supplementary Fig. 21: Target class distributions of binder densities.** The 96-target panel was sorted into the above categories based on annotations available on Uniprot and MobiDB[7], predictions generated in AlphaFold3[5], and calculations made using NetSurfP-3.0[6]. The binder densities $>10^{-6}$, $>10^{-8}$, $>10^{-10}$, $<10^{-10}$ were converted to the values 4, 3, 2, and 1 respectively to allow plotting. The number of targets ($n_{targets}$) in each category is annotated above its corresponding violin.

**Supplementary Table 8: Target Characteristics.**

| Target | Expression level in PANCS *See plot below* | % disorder | Length (AA) | Solvent-accessible surface area (Å) | % Alpha Helix | % Beta Sheet | AlphaFold pTM of +AP | AlphaFold iPTM with top binder from A, B, D or best only hit in C |
|---|---|---|---|---|---|---|---|---|
| LC3B | 335000 | 0 | 125 | 9815.38092 | 25.6 | 22.4 | 0.81 | 0.65 |
| Mdm2 | 240000 | 25.5 | 188 | 18834.6704 | 20.7446809 | 3.19148936 | 0.45 | 0.16 |
| GABARAP | 2500000 | 0 | 117 | 8851.64606 | 29.9145299 | 23.9316239 | 0.9 | 0.21 |
| KIX | 100 | 27.36 | 106 | 9875.06945 | 59.047619 | 0 | 0.63 | 0.79 |
| SIX1 | 5.71923471 | 36.2 | 282 | 24752.6963 | 39.7887324 | 1.05633803 | 0.49 | 0.91 |
| FRB | 15000 | 0 | 96 | 7170.10885 | 79.1666667 | 0 | 0.86 | 0.57 |
| FnbpA | 150 | 0 | 154 | 10988.3595 | 1.92307692 | 37.1794872 | 0.79 | 0.28 |
| IFNG | 650000 | 8.27 | 133 | 10843.4333 | 63.4328358 | 0 | 0.76 | 0.17 |
| HRAS | 125000 | 10.6 | 189 | 12276.0705 | 34.3915344 | 19.5767196 | 0.86 | 0.84 |
| mCherry | 2000000 | 0 | 236 | 16429.1118 | 5.93220339 | 50 | 0.9 | 0.23 |
| PIK3CB | 1250 | 13.39 | 113 | 8326.07542 | 21.9298246 | 25.4385965 | 0.77 | 0.86 |
| WWP2 | 6.58416007 | 24.4 | 870 | 61709.94 | 23.7931034 | 18.1609195 | 0.63 | 0.38 |
| p65-str | 8800 | 49.6 | 277 | 18424.1857 | 7.55395683 | 38.8489209 | 0.8 | 0.22 |
| MBP | 4250 | 0 | 370 | 19184.818 | 47.4393531 | 18.328841 | 0.85 | 0.87 |
| ZB | 120000 | 0 | 33 | 4091.4816 | 81.8181818 | 0 | 0.54 | 0.34 |
| XIAP | 1.64475453 | 19.7 | 497 | 36825.0807 | 30.9859155 | 8.45070423 | 0.36 | 0.48 |
| PP2A-B" | 300 | 27.03 | 529 | 40020.5229 | 41.7769376 | 1.51228733 | 0.74 | 0.23 |
| Myc-DBD | 260000 | 0 | 84 | 8064.3284 | 84.3373494 | 0 | 0.36 | 0.28 |
| RNF8 (ΔZn finger) | 4.07577816 | 36.3 | 446 | 40188.3457 | 33.1838565 | 10.7623318 | 0.54 | 0.35 |
| SH3BP5 | 8250 | 21.1 | 128 | 12426.8112 | 78.125 | 0 | 0.83 | 0.22 |
| HIF1a | 1700 | 18.29 | 350 | 24555.8211 | 29.1428571 | 23.1428571 | 0.69 | 0.49 |
| LMO2 | 15500 | 8.2 | 158 | 12325.9729 | 14.556962 | 24.0506329 | 0.51 | 0.22 |

| | | | | | | | | |
|---|---|---|---|---|---|---|---|---|
| HPV-pE7 | 1550 | 52 | 98 | 8786.04469 | 9.18367347 | 15.3061224 | 0.92 | 0.18 |
| 14-3-3z | 1000000 | 0 | 245 | 17252.3282 | 77.9591837 | 0 | 0.9 | 0.59 |
| TBX20-DBD | 0.3484538 | 0 | 180 | 12263.4429 | 13.8121547 | 34.8066298 | 0.7 | 0.46 |
| Fibronectin | 625 | 0 | 460 | 31094.5299 | 0.64794816 | 48.1641469 | 0.22 | 0.89 |
| OTUB1 | 250 | 22.54 | 102 | 8260.75811 | 23.7623762 | 12.8712871 | 0.78 | 0.68 |
| HSPB1-str | 75000 | 1.22 | 82 | 6415.59171 | 7.22891566 | 50.6024096 | 0.87 | 0.31 |
| KRAS (G12D) | 90000 | 18 | 189 | 12882.8702 | 35.106383 | 19.1489362 | 0.43 | 0.3 |
| TCIM | 700 | 54.7 | 106 | 9277.11545 | 54.7169811 | 0 | 0.79 | 0.33 |
| RAF | 25000 | 0 | 90 | 6467.92089 | 21.25 | 25 | 0.88 | 0.74 |
| PDE6D | 4200 | 0 | 150 | 10823.7544 | 12 | 48.6666667 | 0.95 | 0.24 |
| Keap1 | 10500 | 2.06 | 291 | 15290.4563 | 0 | 48.630137 | 0.03 | 0.37 |
| BAX (short) | 2500000 | 100 | 15 | 1871.20416 | 76.4705882 | 0 | 0.67 | 0.47 |
| Bcl2 | 50 | 31.8 | 239 | 16696.0783 | 48.6238532 | 0 | 0.87 | 0.86 |
| KRAS WT | 335000 | 18 | 189 | 12773.0069 | 35.106383 | 19.6808511 | 0.89 | 0.86 |
| TEV | 1250 | 0 | 238 | 15599.662 | 9.74576271 | 36.8644068 | 0.2 | 0.27 |
| MST2 | 1000 | 100 | 125 | 14339.9345 | 0 | 0 | 0.93 | 0.24 |
| HEWL | 12950 | 0 | 129 | 7487.45308 | 44.1860465 | 3.87596899 | 0.9 | 0.22 |
| HIgA toxin | 150 | 0 | 280 | 19148.6616 | 5.86206897 | 47.5862069 | 0.4 | 0.87 |
| CDKN1A | 200 | 54.3 | 164 | 18254.5717 | 14.0243902 | 2.43902439 | 0.91 | 0.47 |
| UEV | 160000 | 1.38 | 145 | 10180.4631 | 27.5862069 | 29.6551724 | 0.27 | 0.17 |
| IBTK | 1250 | 80.42 | 181 | 19489.9885 | 19.7044335 | 4.43349754 | 0.42 | 0.47 |
| THAP1 | 2.43839714 | 61 | 213 | 21599.1711 | 27.6995305 | 8.92018779 | 0.72 | 0.82 |
| OLIG2-DBD | 0.57622328 | 0 | 58 | 4675.0372 | 76.7857143 | 0 | 0.66 | 0.64 |
| Parkin | 11500 | 18.2 | 465 | 32053.5665 | 18.0645161 | 23.8709677 | 0.72 | 0.17 |
| VHL | 500000 | 3.36 | 149 | 10793.3489 | 27.5167785 | 29.5302013 | 0.2 | 0.39 |
| NIX | 196.25 | 100 | 219 | 23157.4097 | 12.3287671 | 0 | 0.71 | 0.24 |
| TNFR2 | 15000 | 4.62 | 172 | 15533.0219 | 0 | 21.9653179 | 0.87 | 0.34 |

| | | | | | | | | |
|---|---|---|---|---|---|---|---|---|
| NRAS | 445000 | 15.8 | 189 | 12350.5343 | 34.9206349 | 19.5767196 | 0.47 | 0.15 |
| GATA2-DBD | 0.27824576 | 0 | 23 | 6278.36231 | 12.5 | 15 | 0.87 | 0.4 |
| G3BP | 7100 | 0 | 129 | 8752.38068 | 24.0310078 | 45.7364341 | 0.75 | 0.41 |
| PP2A-B' | 375 | 18.3 | 486 | 36414.9377 | 65.2263374 | 0 | 0.89 | 0.27 |
| CRBN | 250000 | 0 | 388 | 25401.2352 | 22.8791774 | 30.0771208 | 0.46 | 0.32 |
| CD3d | 15000 | 38.6 | 171 | 14449.2012 | 11.1111111 | 21.6374269 | 0.73 | 0.11 |
| MAX | 1900 | 15.38 | 65 | 7398.70295 | 85 | 0 | 0.66 | 0.26 |
| SIX1-DBD | 0.46171414 | 100 | 167 | 12719.3695 | 58.6826347 | 2.39520958 | 0.23 | No hit |
| PCNA-AF | 750 | 100 | 111 | 13979.3439 | 3.6036036 | 0 | 0.92 | No hit |
| TRIM21 | 1350 | 0 | 199 | 12248.7366 | 4.5 | 34 | 0.77 | No hit |
| PD-L1 | 4250 | 100 | 222 | 16653.5281 | 5.82959641 | 49.7757848 | 0.59 | No hit |
| ALKBH5 | 682.5 | 45.5 | 394 | 32834.4653 | 13 | 14 | 0.85 | No hit |
| BTK (PH) | 375 | 0 | 133 | 11125.8294 | 12.0300752 | 30.075188 | 0.68 | No hit |
| hSUMO1 | 0 | 29.7 | 101 | 9084.31409 | 17.8217822 | 26.7326733 | 0.69 | No hit |
| ASB1 | 0.13215216 | 19.73 | 147 | 10133.6271 | 41.4965986 | 0 | 0.61 | No hit |
| GTF2I | 750 | 38.68 | 106 | 9499.94055 | 47.1698113 | 11.3207547 | 0.14 | No hit |
| ERG-IDR | 3.37094791 | 99.55 | 112 | 10855.645 | 0 | 2.67857143 | 0.82 | No hit |
| FGFR1c (D2) | 12450 | 16.16 | 99 | 7411.17854 | 4.04040404 | 52.5252525 | 0.64 | No hit |
| TRIM63 | 2.01442144 | 26.29 | 193 | 20041.6367 | 66.4948454 | 0 | 0.34 | No hit |
| ZBED2 | 3.0127846 | 51.4 | 218 | 22831.8525 | 39.4495413 | 4.12844037 | 0.83 | No hit |
| PMP22 | 247.5 | 0 | 160 | 8715.92983 | 62.5 | 10 | 0.22 | No hit |
| p53 (short) | 1500 | 100 | 61 | 6817.67574 | 11.4754098 | 0 | 0.7 | No hit |
| p53 (long) | 5 | 34.81 | 293 | 23392.9032 | 6.48464164 | 28.3276451 | 0.41 | No hit |
| BIRC7 | 0.50552561 | 49.7 | 298 | 21960.1438 | 22.147651 | 6.7114094 | 0.38 | No hit |
| MXD1 | 375 | 47.5 | 221 | 24637.9915 | 38.0090498 | 0 | 0.34 | No hit |
| HES1 | 0.15255698 | 55 | 280 | 24726.2558 | 32.8571429 | 3.57142857 | 0.43 | No hit |
| HIF1b | 4.55 | 35.65 | 474 | 35650.1622 | 23.2067511 | 20.0421941 | 0.25 | No hit |

| | | | | | | | | |
|---|---|---|---|---|---|---|---|---|
| CD3z | 25000 | 86.5 | 164 | 17382.3237 | 14.6341463 | 0 | 0.84 | No hit |
| POU5F1-DBD | 1.06227406 | 0 | 75 | 5824.02851 | 74.6666667 | 0 | 0.45 | No hit |
| HSPB1 | 5700 | 24.3 | 205 | 19572.0933 | 2.92682927 | 25.3658537 | 0.29 | No hit |
| HOXA9 | 111.198321 | 80.5 | 272 | 30878.8141 | 14.7058824 | 0 | 0.89 | No hit |
| PP2A-B | 1100 | 2.5 | 447 | 30304.4442 | 0.67114094 | 32.4384787 | 0.82 | No hit |
| HAND2-DBD | 1.61878119 | 33.96 | 53 | 4893.14477 | 61.1111111 | 0 | 0.7 | No hit |
| PIK3CA | 1550 | 0 | 105 | 7957.16988 | 20.952381 | 26.6666667 | 0.87 | No hit |
| GATA4 | 0.77721193 | 76 | 442 | 48523.0084 | 4.75113122 | 2.48868778 | 0.19 | No hit |
| GAPBA-DBD | 0.69797803 | 0 | 84 | 6580.10507 | 65.4761905 | 0 | 0.85 | No hit |
| Sos1 | 150 | 100 | 198 | 23371.7986 | 0 | 0 | 0.11 | No hit |
| RHEB | 2.51653994 | 7.6 | 184 | 11970.378 | 35.326087 | 20.6521739 | 0.88 | No hit |
| PINK | 50000 | 10.53 | 247 | 17756.7427 | 24.291498 | 14.9797571 | 0.72 | No hit |
| BAX | 65000 | 35.53 | 112 | 9040.44548 | 54.4642857 | 0 | 0.6 | No hit |
| FKBP | 6650 | 0 | 107 | 7260.99139 | 12.1495327 | 38.317757 | 0.92 | No hit |
| PPP1R11 | 7850 | 100 | 126 | 15246.6175 | 0 | 0 | 0.19 | No hit |
| HMGB1 | 4850 | 38.1 | 215 | 22168.9657 | 49.7674419 | 0 | 0.37 | No hit |
| PD-1 | 250 | 16.18 | 136 | 11083.9263 | 2.18978102 | 43.0656934 | 0.75 | No hit |
| RB1 | 0.025 | 59.2 | 103 | 10605.5423 | 22.3300971 | 2.91262136 | 0.34 | No hit |
| OmpA (ec) | 1500000 | 0 | 141 | 9279.30182 | 41.4285714 | 20 | 0.92 | No hit |
| RNF2 (**Δ**Zn finger) | 74.2498266 | 54.39 | 113 | 12406.3015 | 26.0869565 | 0 | 0.27 | No hit |

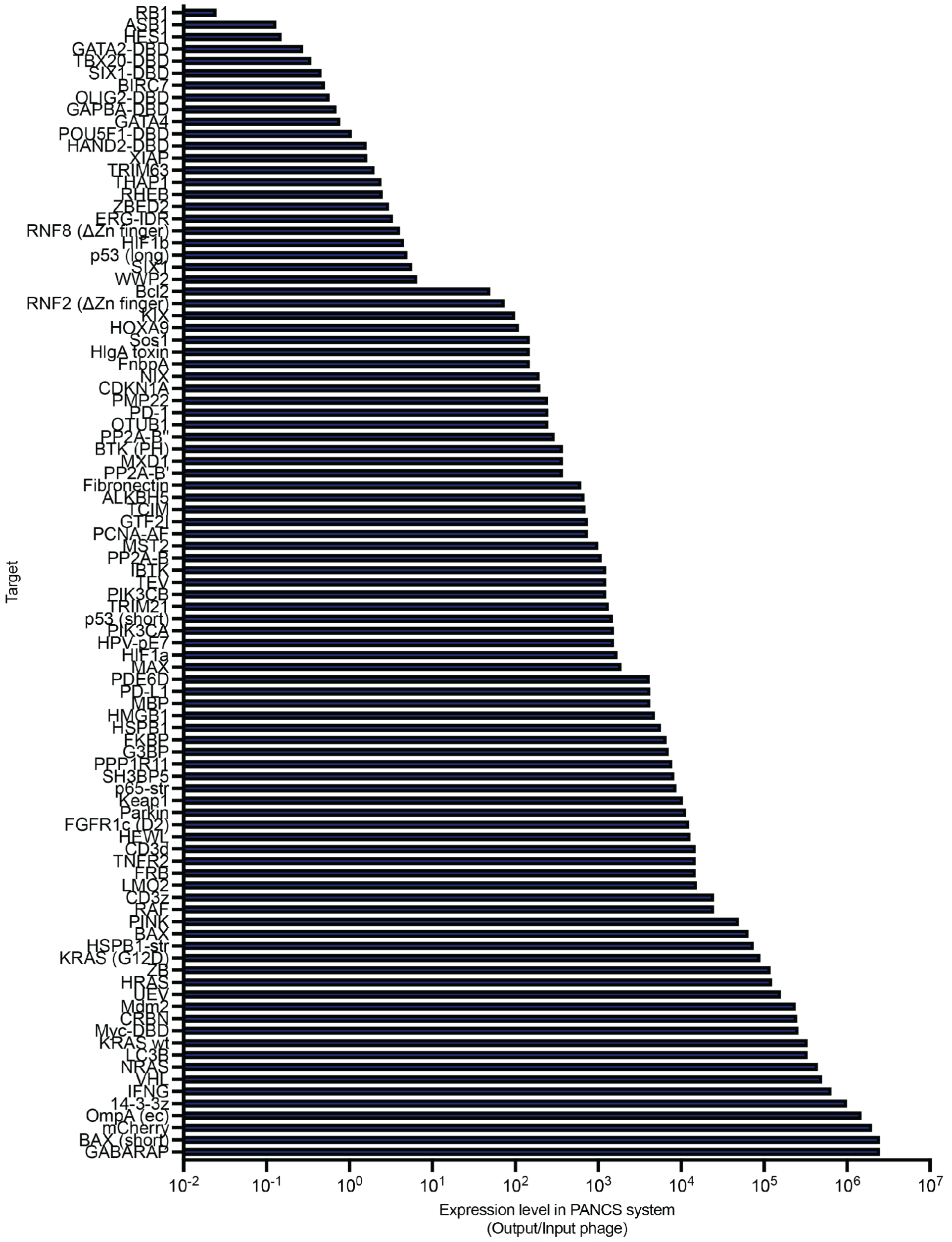

**Bar plot depicting the expression level of each target in the PANCS system.** The expression level of each target was measured by infecting S1030 E. coli transformed with the +AP only with 1000 PFU of phage encoding Nwt T7 RNAP N-term variants. After 10h incubation, the titer was measured and normalized to the titer of the input phage, generating an output/input phage value.

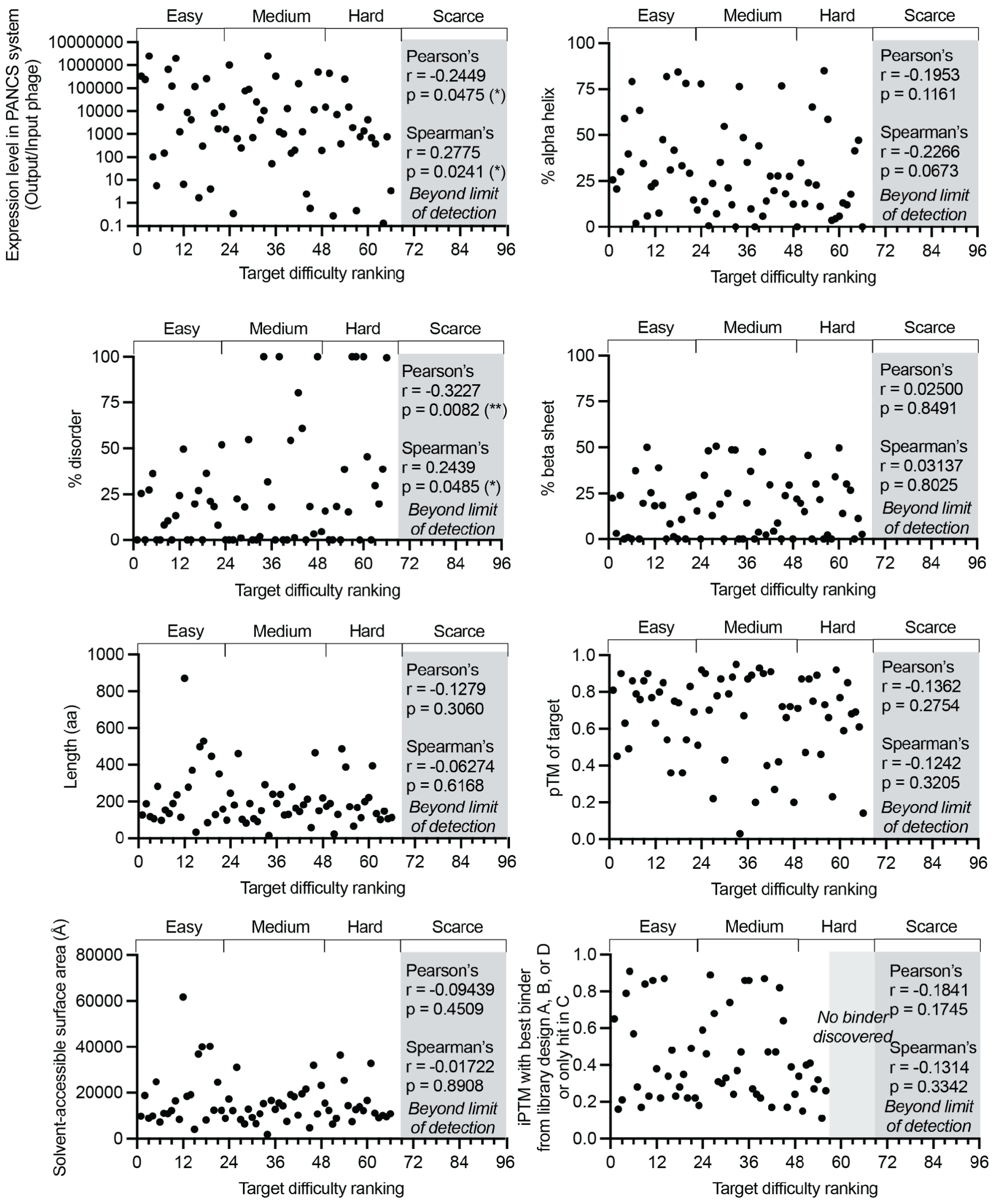

**Supplementary Fig. 22: Scatterplots of target difficulty ranking versus intrinsic target characteristic, with Pearson's and Spearman's correlations.** All data to make scatterplots is available in **Supplementary Table 8**, and represents the intrinsic target characteristics of expression level in PANCS system, % disorder, length (amino acids), solvent-accessible surface area, % alpha helix, % beta sheet, AlphaFold 3 pTM, and iPTM with best binder from designs A, B, D, or only hit in C.

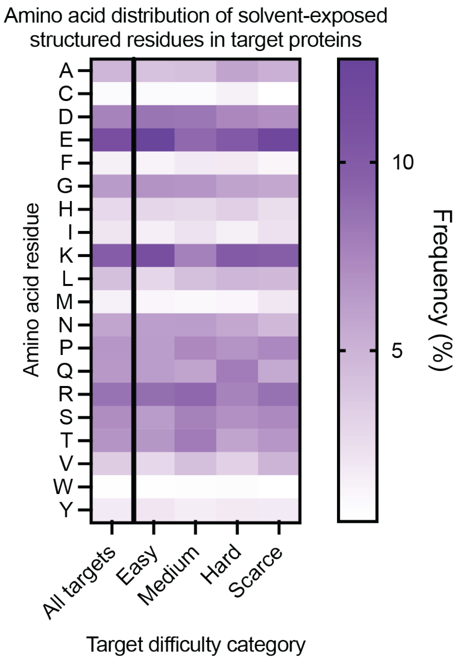

**Supplementary Fig. 23: Heat map depicting the frequency (%) of amino acid usage in solvent-accessible residues in target proteins, categorized by target difficulty.** Target difficulty was first re-defined in the two most difficult categories by designating all targets with hits discovered in either a $10^{10}$-sized sampling with design A or a $10^{8}$-sized other design (B-J) as “hard”, and anything where no hits were discovered, as “scarce”. After using NetSurfP-3.0 and AlphaFold 3 to determine if a residue is relatively solvent exposed (RSA value >0.25, AF3 pLDDT value >70), the frequency of each amino acid was calculated as a percentage of all residues identified across all solvent-accessible residues in our 96-target panel.

**Supplementary Table 9: Results of the Multiple Linear Regression of Target Characteristics vs Binder Density Category.**

| Variable | Sum of Squares | Degrees of freedom | P-value | Estimate (ß coefficient) | VIF |
|---|---|---|---|---|---|
| Regression | 5512 | 7 | 0.0270 | | |
| Residual | 18841 | 58 | | | |
| Total | 23953 | 65 | | | |
| Expression in PANCS system | 1400 | 1 | 0.0402 | -9.735E-06 | 1.134 |
| % disorder | 2344 | 1 | 0.0087 | 0.2845 | 2.263 |
| Length (AA) | 10.32 | 1 | 0.8576 | -0.01499 | 30.62 |
| Solvent accessible surface are a ($Å^2$) | 3.819 | 1 | 0.9131 | -0.0001293 | 31.12 |
| % alpha helix | 62.62 | 1 | 0.6588 | -0.07003 | 3.131 |
| % beta sheet | 23.20 | 1 | 0.7880 | 0.07497 | 4.328 |
| AlphaFold 3 pTM | 230.2 | 1 | 0.3983 | 15.57 | 3.630 |

Where n = 66, $R^2 = 0.2301$

| Normality of Residuals | Statistics | P-value | Passed normality test? |
|---|---|---|---|
| D'Agostino-Pearson omnibus (K2) | 3.688 | 0.1582 | Yes |
| Anderson-Darling (A2*) | 0.5731 | 0.1316 | Yes |
| Shapiro-Wilk (W) | 0.9668 | 0.0743 | Yes |
| Kolmogorov-Smirnov (distance) | 0.08786 | >0.1000 | Yes |

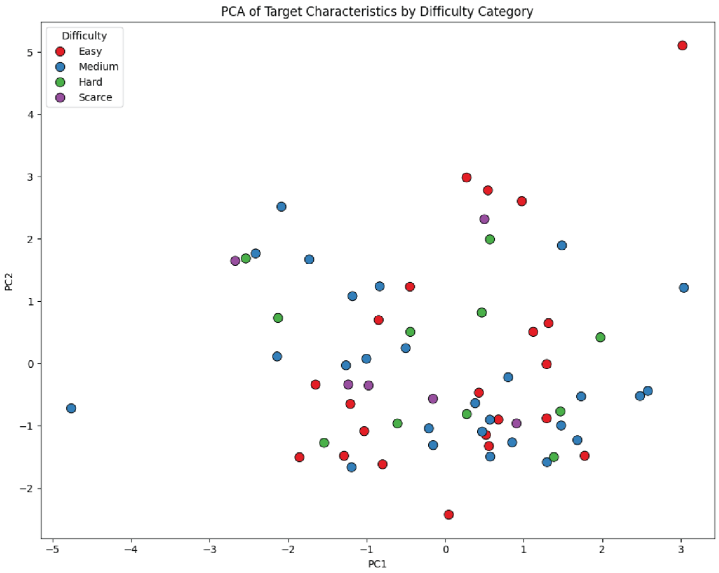

**Supplementary Fig. 24: Principal component analysis (PCA) of 8 myriad intrinsic target features.** The data from Supplementary Table 8 was used as an input matrix for attempting to perform PCA to determine if target features can cause targets to cluster by target difficulty categories. The PC loadings are included as well to demonstrate the impact and direction of each parameter on each axis.

| Model | Parameters | Macro AUC |
|---|---|---|
| d=1, 1 head | 7,680 | 0.771 ± 0.018 |
| d=6, 1 head | 20,480 | 0.840 ± 0.016 |
| d=10, 1 head | 30,720 | 0.848 ± 0.014 |
| d=32, 1 head | 87,040 | 0.855 ± 0.015 |
| d=8, 4 heads | 87,040 | 0.888 ± 0.011 |
| d=64, 1 head | 168,960 | 0.863 ± 0.008 |
| d=64, 1 head * | 163,840 | 0.807 ± 0.023 |
| Cosine-triplet MLP[16] | 4,931,841 | 0.727 ± 0.014 |

**Supplementary Table 10: Model configurations, parameter counts, and macro ROC-AUC.** Values are mean ± s.d. across the five cross-validation folds. *d* is the rank of the interaction matrix. *Variant operating on SAE-reconstructed embeddings without LayerNorm, used for the feature decomposition in **Fig. 5**.

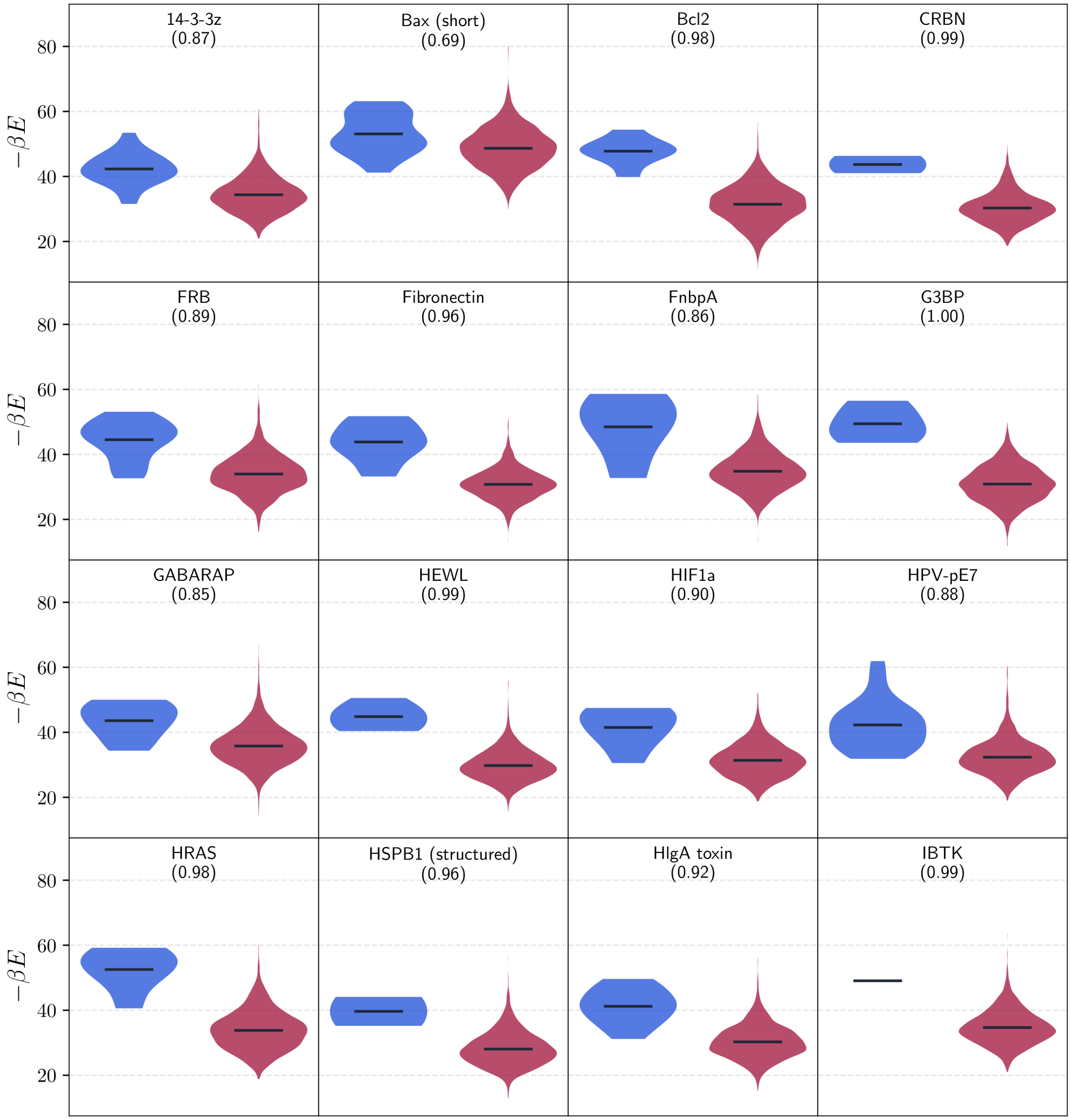

**Supplementary Fig. 25: Per-target log odds distribution and classification performance.** Violin plots of the log odds (−βE) for binders (blue) and non-binders (red) for each of the 52 targets, shown for the $d$ = 8, 4-head model (fold 1). The horizontal line marks the median, and the per-target ROC-AUC is given in parentheses below each target name.

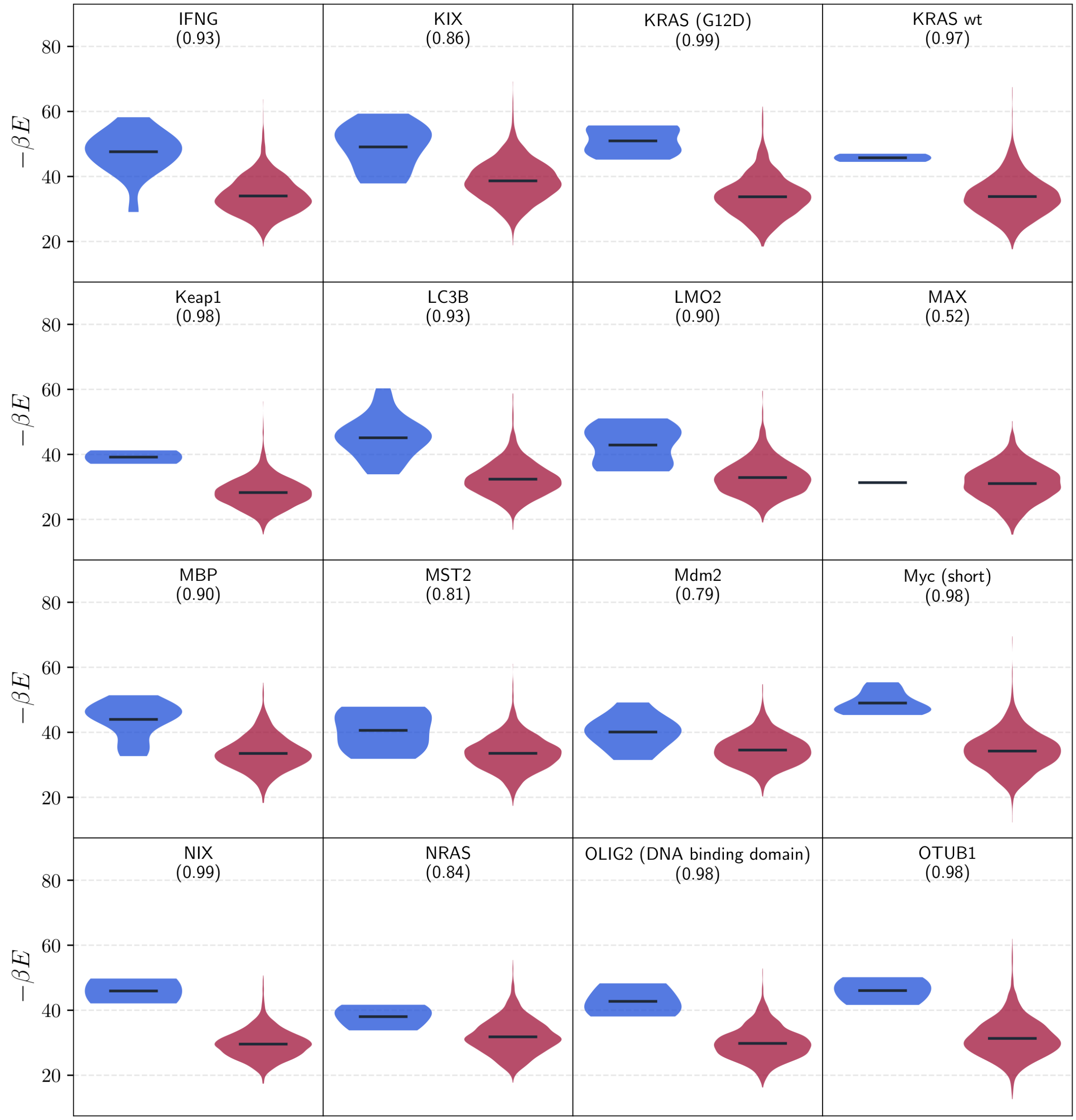

**Supplementary Fig. 25 (continued).**

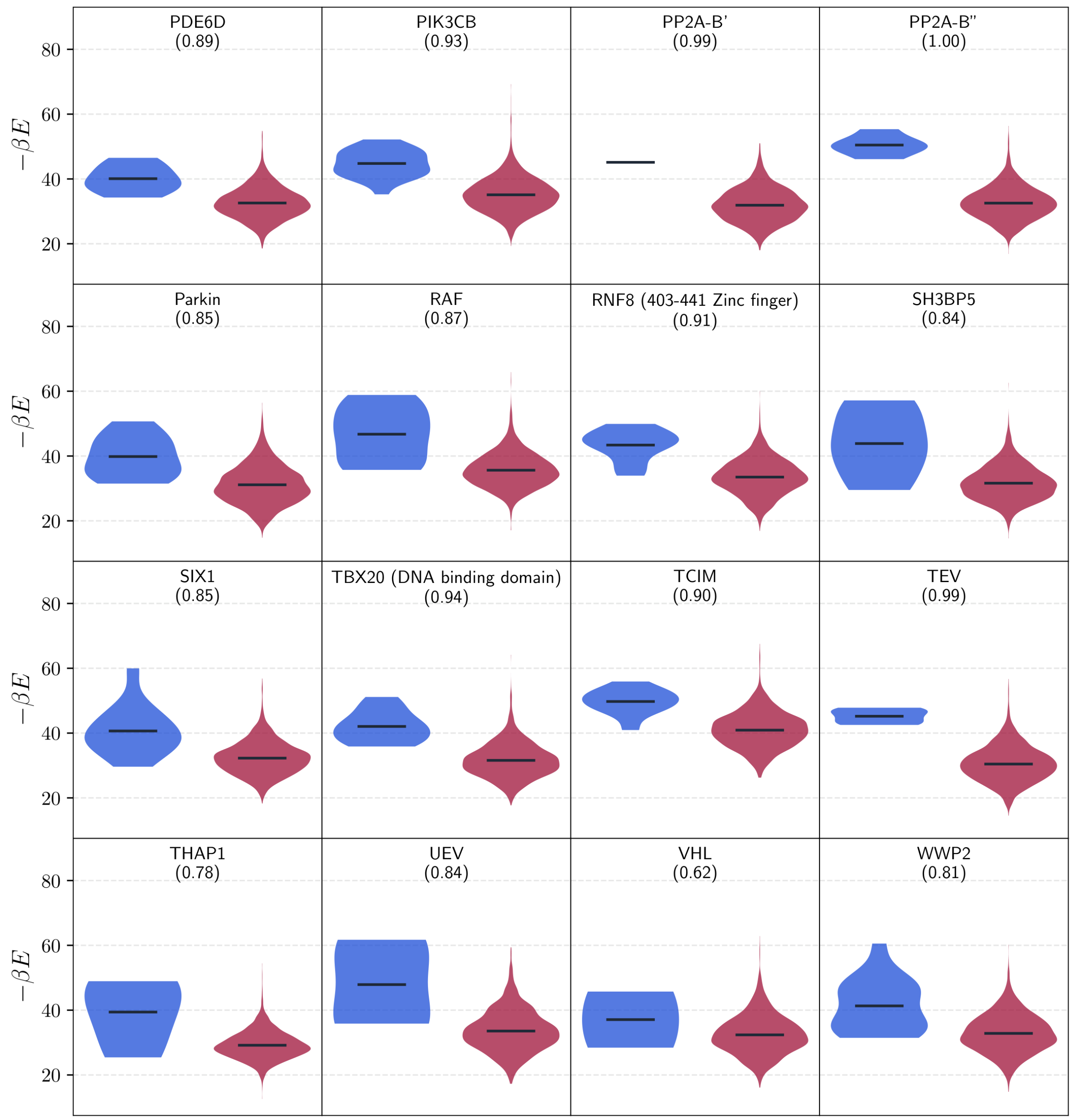

**Supplementary Fig. 25 (continued).**

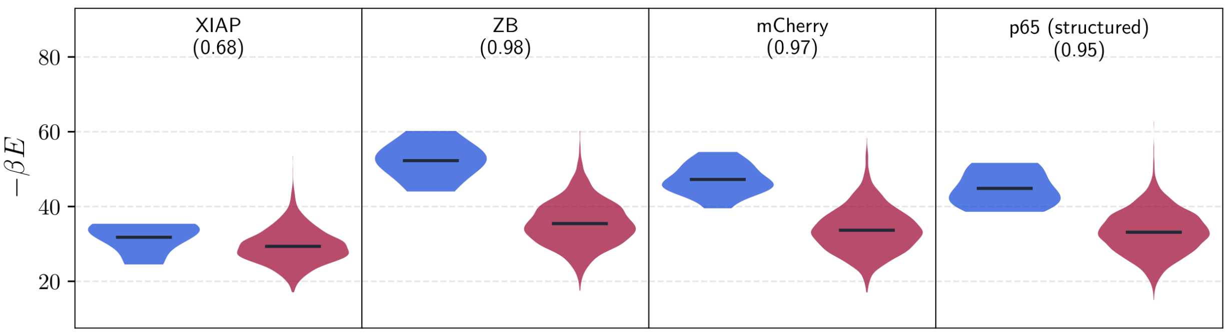

**Supplementary Fig. 25 (continued).**

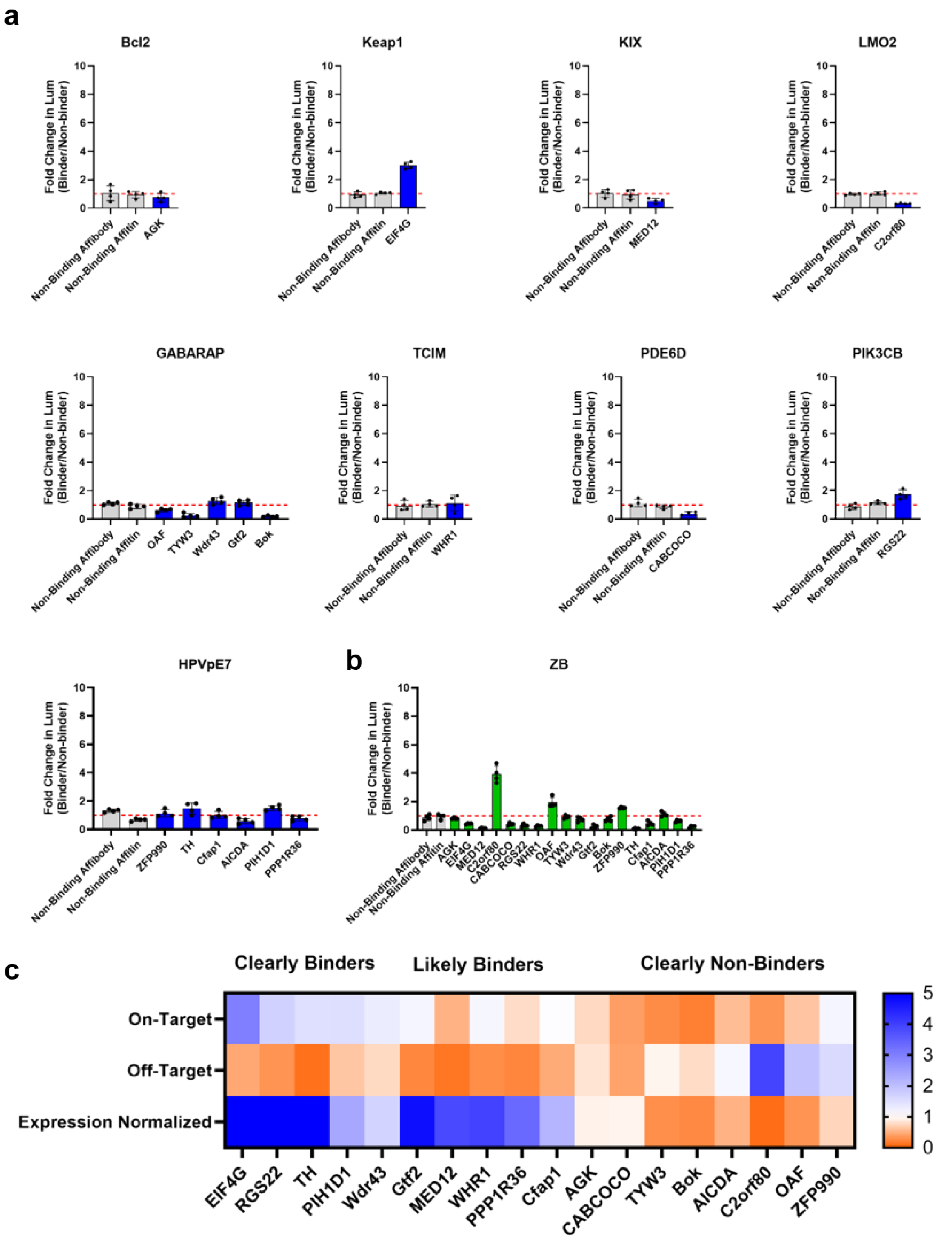

**Supplementary Fig. 26: Assessment of predicted target-mammalian protein interactions.** **a**, Each target (listed at graph title) with two non-binding controls (grey bars) and each predicted binder (blue bars) for all tested predictions using our split RNA polymerase luciferase assay (**Supplementary Fig. 4**). Each bar plot shows the mean, standard deviation, and individual values (*n* = 4, error bars indicate SD). **b**, Each predicted binder assessed for binding to a ZB, as a non-interacting target, to assess differences in expression level of each predicted binder. In selection outputs, scaffolds are expected to express similarly to the nonbinding controls (an assumption that holds when tested, **Supplementary Fig. 19**). Each bar plot shows the mean, standard deviation, and individual values ($n$ = 4, error bars indicate SD). **c**, Summary (averages) of the direct on-target (top row) binding observed (normalized to background of two high expressing non-binding scaffolds), the off-target (middle row) binding as a measure of expression level for each predicted binder, and the expression normalized binding signal (on-target/off-target; bottom row) for each predicted binder-target pair.

**Supplementary Table 11: List of plasmids used in this study.**

| ID | Description | Plasmid Map |
|---|---|---|
| Library cloning template | | |
| 76-31 | Template for cloning affibody libraries and full random 30mer library | https://benchling.com/s/seq-02e90z0Dmx8QenFnZSty?m=slm-OkP6P9eDWs34LeLC4wRx |
| -AP | | |
| 73-79 | $ZB_{neg}$, 60 AA linker, sd8/sd8 | https://benchling.com/s/seq-qptnks4V9vCAkgQsGtmf?m=slm-0SHHTsCgDhaxvivYq6yY |
| +APs | | |
| 75-154 | p15a, SD8/SD8; 60 AA linker | https://benchling.com/s/seq-OAkkUSRN3TJhdoLilhRD?m=slm-bEfoe3bp41FrsbQWjGA9 |
| 73-21 | KRAS (G12D), p15a, SD8/SD8; 60 AA linker | https://benchling.com/s/seq-Zr3IVLYCsltuBwWaWxJr?m=slm-wRhiEYUzN1wIXpbqdbih |
| 73-36 | ZB, p15a, SD8/SD8; 60 AA linker | https://benchling.com/s/seq-B3VpJwTHBJNtDB96TRU3?m=slm-iQ706YAxBZMMwGKFHJVF |
| 73-150 | p65 (17-293), p15a, SD8/SD8; 60 AA linker | https://benchling.com/s/seq-cdSliWqWg0tdHYgBTdOP?m=slm-pEDkDzXE0Z4F8vYUSiit |
| 73-155 | FGFR1c domain D2, p15a, SD8/SD8; 60 AA linker | https://benchling.com/s/seq-Uzm5jeJGfrrqaKG4tbn6?m=slm-OanxydYCDzyfpqszed47 |
| 73-156 | TCIM, p15a, SD8/SD8; 60 AA linker | https://benchling.com/s/seq-6IddRZ1HDqTN3MCPSaK8?m=slm-Ls84NSQmf0GHZLtK9WM8 |
| 73-157 | TRIM21 (277-475) , p15a, SD8/SD8; 60 AA linker | https://benchling.com/s/seq-oEa6R5xeroyTLQdBFxcy?m=slm-qT4xnij6laqu2uwpCMdM |
| 73-158 | PDE6D, p15a, SD8/SD8; 60 AA linker | https://benchling.com/s/seq-DNJTqsoqchb6Ifc8LmkH?m=slm-S7qdc6giMN9TxTm03COK |
| 75-122 | RAF, p15a, SD8/SD8; 60 AA linker | https://benchling.com/s/seq-653ffOYStxIyWy0crwUG?m=slm-popOBq01ASaK3UElPBLr |
| 75-124 | NRAS, p15a, SD8/SD8; 60 AA linker | https://benchling.com/s/seq-rBJD0ntLGgEuAuq1kSED?m=slm-yshqst61vCjixpRdgNbY |
| 75-125 | Parkin, p15a, SD8/SD8; 60 AA linker | https://benchling.com/s/seq-0sHCMSn1WiM7RUEPmNr1?m=slm-DqrmF7gFJy493PGy4yNk |
| 75-128 | Mdm2, p15a, SD8/SD8; 60 AA linker | https://benchling.com/s/seq-PxdMGjVyvDqAJDmdvnLQ?m=slm-QGrbcwpi2958fCky1PHM |
| 75-130 | HlgA toxin, p15a, SD8/SD8; 60 AA linker | https://benchling.com/s/seq-Et7y9HSeppcxczv2YEmK?m=slm-clebjjpTK98RJnKI19EK |
| 75-131 | Bcl2, p15a, SD8/SD8; 60 AA linker | https://benchling.com/s/seq-RMXbBBMr4cQa8yuBErNu?m=slm-phe0A6NggSSQHM8pYvx9 |

| | | |
|---|---|---|
| 75-133 | KIX, p15a, SD8/SD8; 60 AA linker | https://benchling.com/s/seq-75hSB7CK7Ss5mdiXwroT?m=slm-dFHu3SzeYs6wCnenE2hQ |
| 75-135 | HRAS, p15a, SD8/SD8; 60 AA linker | https://benchling.com/s/seq-GOe4zjTl1Mdf4PRb31OS?m=slm-aVc4rYGqhQtzmXMU7JX5 |
| 75-136 | KRAS wt, p15a, SD8/SD8; 60 AA linker | https://benchling.com/s/seq-ZeZ4aYcPMa4jt6kE0JA8?m=slm-urQbtWwArEqCnNkcetG1 |
| 75-137 | G3BP, p15a, SD8/SD8; 60 AA linker | https://benchling.com/s/seq-RmHYPArmkMpqTnNpr0OP?m=slm-2kBpsjPaVT6hPhPNxFxx |
| 75-143 | NIX, p15a, SD8/SD8; 60 AA linker | https://benchling.com/s/seq-r3wjG4NZ4yLcCRmEG7Yd?m=slm-P4Nzro5JL8BBBd0ORv5r |
| 75-140 | PP2A-B, p15a, SD8/SD8; 60 AA linker | https://benchling.com/s/seq-18IkmH1oviCvDSJqO8dh?m=slm-I67s5YoyoAAaUOnnArDm |
| 75-146 | Sos1, p15a, SD8/SD8; 60 AA linker | https://benchling.com/s/seq-w80QMJY61wsyX3exlMz4?m=slm-kRREHX73IefmCI7iWAO6 |
| 75-147 | PP2A-B', p15a, SD8/SD8; 60 AA linker | https://benchling.com/s/seq-TwI5xEyskYV7Sw9GsjRh?m=slm-TzAhfO0Rng5ajuOAwFgi |
| 75-148 | mCherry, p15a, SD8/SD8; 60 AA linker | https://benchling.com/s/seq-So0ZUDzTf8WRPuUMf9t0?m=slm-bnWoNtGNBoNtU3YjwZPc |
| 75-149 | Myc-DBD, p15a, SD8/SD8; 60 AA linker | https://benchling.com/s/seq-EnqwDf6RRDLp9d1xJV4T?m=slm-V2L8jbdYZITnjwyPReYe |
| 75-152 | PP2A-B'', p15a, SD8/SD8; 60 AA linker | https://benchling.com/s/seq-4gOERZifGohryh49vugB?m=slm-zclfZd87C5VyT8h1OZ68 |
| 75-153 | MBP, p15a, SD8/SD8; 60 AA linker | https://benchling.com/s/seq-CTYBnSJpKhNMPCGe49xt?m=slm-Yy5Ydwaik5tUJ7PRdDW7 |
| 75-155 | Fibronectin, p15a, SD8/SD8; 60 AA linker | https://benchling.com/s/seq-zdsltF3MW4Ugw2tCggjl?m=slm-pY7ugHj1aXcbiLaCPTIR |
| 75-156 | VHL, p15a, SD8/SD8; 60 AA linker | https://benchling.com/s/seq-qb7tgAd3oKFG9Z6EqSdo?m=slm-jGDEhoC2D3a3kO81tTL6 |
| 75-157 | BTK (PH), p15a, SD8/SD8; 60 AA linker | https://benchling.com/s/seq-5ALCwW2eLTMX3flVJet3?m=slm-sBY39ws0zDeS7oLoG5SH |
| 75-161 | GABARAP, p15a, SD8/SD8; 60 AA linker | https://benchling.com/s/seq-CU3BT33LpDQcqNz52AYW?m=slm-pM8ur5mMQ5m8gMYjfW4x |
| 75-164 | GTF2I, p15a, SD8/SD8; 60 AA linker | https://benchling.com/s/seq-D6ZmbBc9m2P7cdeZfjkI?m=slm-wbxKJ1T7nHnx8u0dFvP3 |
| 75-166 | 14-3-3z Monomeric (L12Q, R18E and S58E), p15a, SD8/SD8; 60 AA linker | https://benchling.com/s/seq-IWbQqjRmQzcjUYtigywe?m=slm-yt4r5Yag09nYxefCzHFJ |

| | | |
|---|---|---|
| 75-167 | PCNA-AF, p15a, SD8/SD8; 60 AA linker | https://benchling.com/s/seq-XJUWLK9eWZX7HAWauHcu?m=slm-oDX0s1RnucKQJcBeHREu |
| 75-169 | TEV, p15a, SD8/SD8; 60 AA linker | https://benchling.com/s/seq-81ffIJaX0kRAI6VLInsY?m=slm-galMvGSadT65JKoFySYK |
| 75-170 | FnbpA, p15a, SD8/SD8; 60 AA linker | https://benchling.com/s/seq-GjkFFNf2siUBskbdkmDc?m=slm-ASm8m8Enw4i3jjTBtBmf |
| 75-171 | IBTK, p15a, SD8/SD8; 60 AA linker | https://benchling.com/s/seq-Fy1aT64LESd9RoHJb4yy?m=slm-nfQUjcyqVNACboxbxfRP |
| 75-173 | PIK3CB (3-115), p15a, SD8/SD8; 60 AA linker | https://benchling.com/s/seq-7A7cpLNdIIJM7KkPgZaS?m=slm-Y9jqMOLRNBhwlpGµLJOH |
| 75-178 | TNFR2 (33-205), p15a, SD8/SD8; 60 AA linker | https://benchling.com/s/seq-mAyNSHYPq2EBro3MaKM7?m=slm-x8FVA5aJGLiVHCDET5Li |
| 75-179 | CDKN1A, p15a, SD8/SD8; 60 AA linker | https://benchling.com/s/seq-PQ5irZBkcDjh3R4UHBKN?m=slm-cFdFMaHd2Gl1r6awo41C |
| 75-180 | SH3BP5, p15a, SD8/SD8; 60 AA linker | https://benchling.com/s/seq-Avc55NRDP6F4mjVqeGlV?m=slm-o4BfkBdFLzii1YkDeJOP |
| 75-181 | OTUB1 (1-102), p15a, SD8/SD8; 60 AA linker | https://benchling.com/s/seq-O82hpMpccs8FoUNuFrfY?m=slm-DGRFyYIG6QHybTY31P3b |
| 75-150 | PINK, p15a, SD8/SD8; 60 AA linker | https://benchling.com/s/seq-qPq6x39UehzUtZRCla0I?m=slm-yHzeA88dr7KzF6ILd18b |
| 75-185 | RB1, p15a, SD8/SD8; 60 AA linker | https://benchling.com/s/seq-22kbw64bDo5f2vL4T6J3?m=slm-IGZoUdtzriYriAAKfviD |
| 75-188 | FKBP, p15a, SD8/SD8; 60 AA linker | https://benchling.com/s/seq-x40cA8lpEv7xirBM9uxs?m=slm-asEJgRJgltfptOrbcjjj |
| 76-50 | HEWL, p15a, SD8/SD8; 60 AA linker | https://benchling.com/s/seq-CxlSVSbklEqF3Syw042I?m=slm-H3KNC82Pha4FHZB37MbH |
| 75-193 | HSPB1-str (90-171), p15a, SD8/SD8; 60 AA linker | https://benchling.com/s/seq-PJZGJ4mun3IKcOJErxe0?m=slm-Xs5wlOr4SfvbdgRUM6Ed |
| 76-51 | LMO2 (28-150) , p15a, SD8/SD8; 60 AA linker | https://benchling.com/s/seq-NXc2ysSolIG8dC1PXNgL?m=slm-YfpX1I0eijcrk5tYWpZ8 |
| 76-52 | HIF1a, p15a, SD8/SD8; 60 AA linker | https://benchling.com/s/seq-LrW4a6u72I2krof0lNiK?m=slm-Fo74FDHKJbx8z2PrQe0z |
| 76-23 | MXD1, p15a, SD8/SD8; 60 AA linker | https://benchling.com/s/seq-bBeNxjomjgPQRSFiWGOn?m=slm-QyDn8Obf66PvZrYt6iKI |
| 75-195 | OmpA (ec, 84-224) , p15a, SD8/SD8; 60 AA linker | https://benchling.com/s/seq-qGyYe5Xj4xBsdfY7kAAg?m=slm-3Pw3eZnnYOG1s1rfDqRN |

| | | |
|---|---|---|
| 76-40 | CD3z, p15a, SD8/SD8; 60 AA linker | https://benchling.com/s/seq-fF2AIC8XFYµLI5dEVeJk?m=slm-qUnT9qGpKKdfn3OOS1p1 |
| 76-41 | PIK3CA (1-105) , p15a, SD8/SD8; 60 AA linker | https://benchling.com/s/seq-Aned8XSfCRP1MoDFpYwR?m=slm-9Q9yuEB1ZIsZuzYopY4X |
| 76-44 | UEV, p15a, SD8/SD8; 60 AA linker | https://benchling.com/s/seq-WTDkd5RV746qVTqvZFsw?m=slm-I1U9FTb2LkTO29rWI5Wp |
| 76-46 | FRB, p15a, SD8/SD8; 60 AA linker | https://benchling.com/s/seq-mtFkW0QpGYU17MfoCaHn?m=slm-P9kNCCZeZbWASii3O9Bx |
| 76-47 | PD-1, p15a, SD8/SD8; 60 AA linker | https://benchling.com/s/seq-bKsWCHYUYtEDZhayfEFE?m=slm-kkprEYggLqVT9u5bZgwT |
| 75-126 | LC3B, p15a, SD8/SD8; 60 AA linker | https://benchling.com/s/seq-BVnxoGkEIF8WqzyciCqM?m=slm-zsHlH0xb2vn5ws0dScqE |
| 75-127 | ALKBH5, p15a, SD8/SD8; 60 AA linker | https://benchling.com/s/seq-4J5UYICjPsrfCcBQexOw?m=slm-62da3X8otCUcHupJT2yw |
| 75-134 | HIF1b (1-474), p15a, SD8/SD8; 60 AA linker | https://benchling.com/s/seq-ao4jpxAORmJ8nMHYWkdF?m=slm-TRnAWFqxP1nEEQ8iCra6 |
| 75-139 | HPV-pE7, p15a, SD8/SD8; 60 AA linker | https://benchling.com/s/seq-UYuY1jGvfm2m3kCj41aw?m=slm-akfdGrZMUyWMOVSfZH7z |
| 75-144 | MST2 (313-437), p15a, SD8/SD8; 60 AA linker | https://benchling.com/s/seq-jtOStbjKLwrI2hWHuPye?m=slm-z79ntiBq1r4xrktdn7Qj |
| 75-145 | p53 (long), p15a, SD8/SD8; 60 AA linker | https://benchling.com/s/seq-B4Nf7Hq5PWnYghWasWdA?m=slm-Xdi24mTPOXd1sMbq0sHi |
| 76-48 | PD-L1, p15a, SD8/SD8; 60 AA linker | https://benchling.com/s/seq-B74sRi7ocrSyHvdQiQVZ?m=slm-xLrtaQpvUHFhWyCJ84mv |
| 76-1 | BAX (short), p15a, SD8/SD8; 60 AA linker | https://benchling.com/s/seq-8J7TAMAqN4ZLESQkEQ4G?m=slm-SFiX8Vx8PpbEJ9s39bMJ |
| 75-151 | p53 (short), p15a, SD8/SD8; 60 AA linker | https://benchling.com/s/seq-2OLFYp087OjPugBjKupY?m=slm-YIQqIGdSkzx0rHiTQfkZ |
| 75-158 | IFNG (24-156), p15a, SD8/SD8; 60 AA linker | https://benchling.com/s/seq-KsZIMXJQ7LRRHC7GSqti?m=slm-yIqgGLjFLyxIbwwJ2he8 |
| 75-162 | CRBN, p15a, SD8/SD8; 60 AA linker | https://benchling.com/s/seq-7mvwe76RtMPWca9NQqpC?m=slm-oLSx4DVQMj0Mh58fhIgZ |
| 75-163 | MAX, p15a, SD8/SD8; 60 AA linker | https://benchling.com/s/seq-rtp66xstwB8HFXFEQVje?m=slm-duRSyOvzl1gABrDn4rus |
| 75-168 | PPP1R11, p15a, SD8/SD8; 60 AA linker | https://benchling.com/s/seq-MbPMAihvvKbWdKD6Jtg0?m=slm-D92oVzXfQµLUeLExAAve |

| | | |
|---|---|---|
| 75-183 | PMP22, p15a, SD8/SD8; 60 AA linker | https://benchling.com/s/seq-sPUgwnqHHCrprRrHXZk7?m=slm-JomYsxi1wgt1rgWU8Fai |
| 75-189 | BAX, p15a, SD8/SD8; 60 AA linker | https://benchling.com/s/seq-dv8svMLWo7jRgBNANgao?m=slm-0XX5aejmRwEcRVHgMrIl |
| 76-42 | Keap1, p15a, SD8/SD8; 60 AA linker | https://benchling.com/s/seq-EdPHe3rCN1vOvTwxeQCZ?m=slm-KCWjIy09bMTDnKUY92o0 |
| 76-43 | HSPB1, p15a, SD8/SD8; 60 AA linker | https://benchling.com/s/seq-IfKi5eLDKox1xmAmAeah?m=slm-CwP7gFgtQY7tnrVkHMt5 |
| 76-45 | HMGB1, p15a, SD8/SD8; 60 AA linker | https://benchling.com/s/seq-5hGuCN4IoBaLzdNTzHz2?m=slm-ZdlLMfeR7jQGkp5RVitC |
| 76-92 | hSUMO1, p15a, SD8/SD8; 60 AA linker | https://benchling.com/s/seq-EjX9pw5JrR1wN4YXQ5V7?m=slm-hWO7GEglMHG65iIHwYqG |
| 76-93 | CD3d, p15a, SD8/SD8; 60 AA linker | https://benchling.com/s/seq-4hR2jNVµLOXJPDLGbX8B?m=slm-P33HEy8NnH2bAUbX7wbM |
| 77-7 | TRIM63 (160-353), , p15a, SD8/SD8; 60 AA linker | https://benchling.com/s/seq-dAM1NFZ5jRr0NZWb4yve?m=slm-OEXh6iKDgKs1dJ0QNmY9 |
| 77-9 | RHEB, p15a, SD8/SD8; 60 AA linker | https://benchling.com/s/seq-aspxSfXqTdhCJofvWwCe?m=slm-Bd8U3yV2aFL3HU0xLfUd |
| 77-17 | RNF2 (113-226, DZn finger), p15a, SD8/SD8; 60 AA linker | https://benchling.com/s/seq-bIEaHDwwkxUtCx6nNytB?m=slm-DNLMJnB0EdvWfyTui8fE |
| 77-19 | RNF8 (DZn finger), p15a, SD8/SD8; 60 AA linker | https://benchling.com/s/seq-jhCGtufQ9gtJtRWGbhAi?m=slm-1rjUTv4hilZ2u0k9h4Ne |
| 77-23 | ASB1 (1-147), p15a, SD8/SD8; 60 AA linker | https://benchling.com/s/seq-od0PeMXVesxKaaMMiDnQ?m=slm-Wcx8Z0qJDLlvSzADrKAd |
| 77-26 | BIRC7, p15a, SD8/SD8; 60 AA linker | https://benchling.com/s/seq-YdgbIvDMGaXml7knAqn4?m=slm-8BOwAEM6Py0217iCd40k |
| 77-37 | XIAP, p15a, SD8/SD8; 60 AA linker | https://benchling.com/s/seq-e0Om1642QH9XnVLx5qW8?m=slm-TcZdBzJ3UnC31UWANuOW |
| 77-160 | SIX1-DBD (1-167), p15a, SD8/SD8; 60 AA linker | https://benchling.com/s/seq-M4kq9TfnCrhV0xUyXhDL?m=slm-KouftFUTfxhbVwcyKCH6 |
| 77-45 | WWP2, p15a, SD8/SD8; 60 AA linker | https://benchling.com/s/seq-7pwiUrKFYfdjuJ9xTndX?m=slm-WQaUXaLIBR6f6KybeuYN |
| 77-121 | SIX1, p15a, SD8/SD8; 60 AA linker | https://benchling.com/s/seq-ox0KtlsyVaXReEr3By6s?m=slm-bI62gkglZVVvKCsbT1Ec |
| 77-126 | ERG-IDR (1-112), p15a, SD8/SD8; 60 AA linker | https://benchling.com/s/seq-JWRPkmFhTK9dMw8lV6If?m=slm-RPaucK16pOP1Xrv811I5 |

| | | |
|---|---|---|
| 77-129 | HAND2-DBD (99-151), p15a, SD8/SD8; 60 AA linker | https://benchling.com/s/seq-Z2Y8vncq5k5G1aqR5eWn?m=slm-eY4hhw4o0Wr6KGKfcOWH |
| 77-133 | HES1, p15a, SD8/SD8; 60 AA linker | https://benchling.com/s/seq-2jDU62wW983US6hLjtYz?m=slm-Iv68zjIUn5c0AFsolFPf |
| 77-161 | HOXA9, p15a, SD8/SD8; 60 AA linker | https://benchling.com/s/seq-R535LuMGxYlNeQe9zC5l?m=slm-H4fEprgAi5JQAIQisUEt |
| 77-182 | GATA2-DBD (295-373), p15a, SD8/SD8; 60 AA linker | https://benchling.com/s/seq-S2zqgQuEPdTkOggyPzQU?m=slm-7UHl6NM8h5tvwpGgZUF3 |
| 77-183 | POU5F1-DBD (138-212), p15a, SD8/SD8; 60 AA linker | https://benchling.com/s/seq-arQtTyDT2zIoMYJhaZgw?m=slm-FGGrKL3Fx2d3Gg2wPyn1 |
| 77-184 | THAP1, p15a, SD8/SD8; 60 AA linker | https://benchling.com/s/seq-axg3xcbrZ9dteUiKKX1l?m=slm-Q1mSs0BjwMIca7T8HfvK |
| 77-186 | TBX20-DBD (109-288), p15a, SD8/SD8; 60 AA linker | https://benchling.com/s/seq-Ih2T3TLSjt8iSgyQTNNb?m=slm-RbzplWOmKom4NyT2LMA9 |
| 77-188 | OLIG2-DBD (108-165), p15a, SD8/SD8; 60 AA linker | https://benchling.com/s/seq-FQyiMTP9UpNIYwS1aOjU?m=slm-z0ph3bh8GzelVa3YJ5Ys |
| 77-190 | ZBED2, p15a, SD8/SD8; 60 AA linker | https://benchling.com/s/seq-XdSq64wmTwEVcMKKZQd6?m=slm-52dopiggz6oSV1YvbYUN |
| 77-193 | GAPBA-DBD (168-251), p15a, SD8/SD8; 60 AA linker | https://benchling.com/s/seq-9R9g90KCBGaoZGrAu54S?m=slm-gxzDhVcQFBFpkyL2PJAT |
| 77-196 | GATA4, p15a, SD8/SD8; 60 AA linker | https://benchling.com/s/seq-6o9iID39NVgTpHjLKM0U?m=slm-a5NDROJEUCi18raYmjUq |
| Luciferase Assay | | |
| 2-22 | pSC101, LuxAB under T7 RNAP promoter | https://benchling.com/s/seq-Udyz6dIESk9lOHBzqfsO?m=slm-pPVnaTxvld7HcIZ8InVO |
| 77-153 | Lux-N Affibody (PDL1) | https://benchling.com/s/seq-4Hat7A1iLPOjVB5SfuyL?m=slm-kG73luQP4lmO9k1o5Xk0 |
| 77-144 | Lux-N Affitin (SasA) | https://benchling.com/s/seq-MzeKf9F5Wl4KL6VcGZvu?m=slm-EWzvy4QEH0p4MplFX396 |
| 77-147 | Lux-N Monobody (hSUMO1) | https://benchling.com/s/seq-DIRqB3iHFxxrwaJ30gE7?m=slm-ZXarjzck1DUgI2snLzHR |
| 65-50 | Lux-N RNAP$_{N(WT)}$- IFNG (not PPI dependent) | https://benchling.com/s/seq-z6k7WcWzlkehV2RIfEsz?m=slm-nF8asOivpVyPfh7mAXFB |
| 73-21 | KRAS (G12D) | https://benchling.com/s/seq-bHUdLMQyJLyYaNnCzWYo?m=slm-kqR0h1614DqdUkezQdiP |
| 73-22 | ZB | https://benchling.com/s/seq-cUHUKYXIIfvrpiwQffeK?m=slm-f8l0TBya1R6o360d4148 |

| | | |
|---|---|---|
| 76-152 | p65 (17-293) | https://benchling.com/s/seq-MFerrPNUPyioan1e5CPY?m=slm-7aRHx4Zp6Q72hCtJL1od |
| 76-105 | FGFR1c domain D2 | https://benchling.com/s/seq-9Ex1W5lAuuNbWmXbwFtM?m=slm-EVPbFTSOCZimmw58YBLK |
| 73-167 | TCIM | https://benchling.com/s/seq-nTnoMw1XCU0jQnn1ZXIE?m=slm-BYn2HKhFztBOk5msXYea |
| 73-173 | TRIM21 (277-475) | https://benchling.com/s/seq-qCMpyJNRnolctgFACU7q?m=slm-pOqLvGfB2rU0lwy0zrm3 |
| 76-88 | PDE6D | https://benchling.com/s/seq-8pMSWqjiCrQuri2DX3Kl?m=slm-3knkHTbWEwJnUEY4iboI |
| 73-23 | RAF | https://benchling.com/s/seq-O0XHCpWbTjdtbr6hzREO?m=slm-6HSgWO74Jc1t0JNGMmUi |
| 76-97 | NRAS | https://benchling.com/s/seq-MNEcNbqEA5bJjd5OxWp4?m=slm-kltXGR5rmeQg46qZz5G8 |
| 76-170 | Parkin | https://benchling.com/s/seq-Y1b8m3ZTzutnly70SRFH?m=slm-yEQvZsabomStJwvI3Ox1 |
| 76-74 | Mdm2 (1-188) | https://benchling.com/s/seq-lTuETCqeoOd8g8H8z2Xm?m=slm-dYuzgB8XONA4AEyrfbsV |
| 76-145 | HlgA toxin | https://benchling.com/s/seq-4rYld71VjIo3hrfWFvjX?m=slm-DNEtvtTLscs8Bld0O0JV |
| 76-99 | Bcl2 | https://benchling.com/s/seq-XXBGkJPjCBIubOQuCogn?m=slm-DRPjkgVNKU0Dowo7PDjX |
| 76-80 | KIX | https://benchling.com/s/seq-mkPkHaYFfPfnW81wRv1f?m=slm-FalsloRZmTpa36t0T2Ly |
| 76-98 | HRAS | https://benchling.com/s/seq-pgTuUgg65DUpR1fn20uo?m=slm-zeyRxnr9T8TrgjagcSCj |
| 76-96 | KRAS wt | https://benchling.com/s/seq-YNF06s3yMNWhoWHllQZm?m=slm-pOpYJTkPYLfYd25kMJqP |
| 76-106 | G3BP | https://benchling.com/s/seq-p6tyxpHvpRKNiw5c2wfR?m=slm-YTOOFgotSluGRC3VgMAY |
| 76-154 | NIX | https://benchling.com/s/seq-xefyGFfkOtoGCYsSLXtH?m=slm-pKPjhLMHVIswd1rsarhV |
| 76-175 | PP2A-B | https://benchling.com/s/seq-NUTSz9eO3JOJlfiwyrE5?m=slm-yAu6yfORB4nt6Pzc8qrU |
| 76-169 | Sos1 | https://benchling.com/s/seq-BetRKoyZuNBd1nUuMrp6?m=slm-dYRwQhHl0u5AfxcmmTU9 |

| | | |
|---|---|---|
| 76-176 | PP2A-B' | https://benchling.com/s/seq-vMWqBFiKweSC3lbfvTet?m=slm-C3zToe5IZgJanvOwg5Jb |
| 76-150 | mCherry | https://benchling.com/s/seq-kYMKyHRGdy2oTZvt2IU3?m=slm-xRUOuQLeTy6oTcayxgBM |
| 76-73 | Myc-DBD | https://benchling.com/s/seq-s86vJay1ws2nFE8d2svR?m=slm-SlShPTCEI2HTtX0nl5If |
| 76-166 | PP2A-B" | https://benchling.com/s/seq-BY4emeWMOUzOfwxXcfvt?m=slm-tcINDmZcOgsImkBKTvfx |
| 76-151 | MBP | https://benchling.com/s/seq-RP3yWG9oCdfyKJMPJdi7?m=slm-bdiXkPhY9MVs3uDPfc3c |
| 76-146 | Fibronectin | https://benchling.com/s/seq-ZzKvhno6Q626q81EjK1d?m=slm-qlkowg5flGJeA9Vk1DI0 |
| 76-78 | VHL | https://benchling.com/s/seq-e5ttFNgV64jbsyNZQ3PY?m=slm-jnoBlNbfrsKSaq2Ep8E9 |
| 76-159 | BTK (PH) | https://benchling.com/s/seq-cZ9gtCN49Kl6B2RBmoyP?m=slm-qIKp2sQWKCw2RvVRqM1X |
| 76-77 | GABARAP | https://benchling.com/s/seq-QWrUgcltA0dgKyKnI5fN?m=slm-TRc3OoEsRBgRJn0jMb2z |
| 76-160 | GTF2I | https://benchling.com/s/seq-wbAjXeo3iMuzOgsGCymv?m=slm-HZwpAi3kcEaOrU708mZN |
| 76-101 | 14-3-3z Monomeric (L12Q, R18E and S58E) | https://benchling.com/s/seq-rYnWfjvOKjhc9ak4KfD7?m=slm-P6aLDJAlg1nAvJWgnMIF |
| 73-165 | PCNA-AF | https://benchling.com/s/seq-NyXUSZ8TBJmIT3RRKy2Q?m=slm-QIapA5oxPlJqO1kuZv6V |
| 73-175 | TEV | https://benchling.com/s/seq-gtv7gSu5b2FGnMJJbnlD?m=slm-4fv9YIXApwoWPXu8mhkd |
| 76-172 | FnbpA | https://benchling.com/s/seq-IheXG0iYJlZxW30gL3Y0?m=slm-XvsmEjdyVf3S8μLrjM0E |
| 76-161 | IBTK | https://benchling.com/s/seq-duQlBjHihWV9fAG3GRnc?m=slm-ELq4y4tZ8zUtm72XAPUG |
| 73-171 | PIK3CB (3-115) | https://benchling.com/s/seq-M7YPl5R5qVffKYHqggRo?m=slm-kcbPs3f0DZT63IlNpyc4 |
| 76-104 | TNFR2 (33-205) | https://benchling.com/s/seq-IvN8nTYDCjqopYD1VWkE?m=slm-r3CzTPlvJvIKRsO8WfZb |
| 73-166 | CDKN1A | https://benchling.com/s/seq-istCEzGSqnP9JetiQCuN?m=slm-RISOjKarN6Ff2X2KVkaH |

| | | |
|---|---|---|
| 76-81 | SH3BP5 | https://benchling.com/s/seq-I1mxq7TOjVqvnYaXbq69?m=slm-S9kmHpbmaAehQ1Cif04r |
| 76-82 | OTUB1 (1-102) | https://benchling.com/s/seq-yBlNWiwMPwIP08fY6rTL?m=slm-Ek7d1P6R9a8umhclHnGY |
| 76-155 | PINK | https://benchling.com/s/seq-A2zauOogFx5uYT7qmcT0?m=slm-1lbMTuobO1B89f2kHsSu |
| 76-83 | RB1 | https://benchling.com/s/seq-Xpy5OSXeQ51g6hi8WiGl?m=slm-28UkfqgISqw1C5vZBdBU |
| 73-179 | FKBP | https://benchling.com/s/seq-EjiHC9vGIVQLxz16BYzG?m=slm-FlerSN3uC2FpJhXgIewN |
| 76-102 | HEWL | https://benchling.com/s/seq-vfQnr94eWLSHUc62TMCm?m=slm-xUA1Kr8fRnMd1cavpdlO |
| 73-164 | HSPB1-str (90-171) | https://benchling.com/s/seq-GUv6WsRBpj4gz6bKvn3W?m=slm-iXkRj7y0gZgfL1PG29pI |
| 76-103 | LMO2 (28-150) | https://benchling.com/s/seq-isKCMFIWJoigsZl8JsEJ?m=slm-5kBZFRPsk6xqvvslyub2 |
| 76-178 | HIF1a | https://benchling.com/s/seq-yEjxiXRm7O8Cvwc7ygFu?m=slm-jY0kKsGMQjCrXn1AumaJ |
| 76-158 | MXD1 | https://benchling.com/s/seq-SXFtHNpq5CNf3Y1GqBzz?m=slm-yAz9909ZIqdM8yxl3RbS |
| 76-84 | OmpA (ec, 84-224) | https://benchling.com/s/seq-gEac4hfWY7YSGwDZ2t1K?m=slm-FHCEGTHZ6TtM3BR38qLT |
| 73-168 | CD3z | https://benchling.com/s/seq-zhEuJgddfbTMl3LhVoeE?m=slm-xkjuBuJDo1ZVinKxxInh |
| 73-172 | PIK3CA (1-105) | https://benchling.com/s/seq-uarUD9PQsIuP4MamVl5m?m=slm-N3NhFTD6CqskaShokbqy |
| 73-176 | UEV | https://benchling.com/s/seq-1x3VYe3Rn6Cl2xHsf2My?m=slm-uBwK3p8e3BRgYpSLxbYu |
| 73-178 | FRB | https://benchling.com/s/seq-P2jkD6cGA69xaoxjgGiV?m=slm-cpQ3pvLLLgDsEorNynV0 |
| 73-182 | PD-1 | https://benchling.com/s/seq-yQzKo1Jh4I6HZq5rVG2V?m=slm-14baz2qEaANS3WMAKP5E |
| 76-79 | LC3B | https://benchling.com/s/seq-rlt5IzSBDshWhl9Rb2fu?m=slm-SμL56BpmJVP1Nz5nKQcW |
| 76-156 | ALKBH5 | https://benchling.com/s/seq-s6c48MGHowq9jMME85ff?m=slm-23tUDuPOmB1iHJnMrSFO |

| | | |
|---|---|---|
| 76-177 | HIF1b (1-474) | https://benchling.com/s/seq-4qbTb9Y8HldXECOCqQxH?m=slm-bH3t5YRzVtbYllKm7XGF |
| 76-72 | HPV-pE7 | https://benchling.com/s/seq-Dc5dfBI5LB2iEjmcv8vV?m=slm-dCd1RlzjxdwI3r8YpUXR |
| 77-54 | MST2 (313-437) | https://benchling.com/s/seq-3QXRIY0pRUN97KNELCgH?m=slm-eJW6pYZYT851VaVwyHhC |
| 76-144 | p53 (long) (1-293) | https://benchling.com/s/seq-Fu83Td8HuPIQwC2nRt64?m=slm-SWkrs4aEcLewx5C6B1oC |
| 73-183 | PD-L1 | https://benchling.com/s/seq-RvhD3N2PU1hj77T4fwoo?m=slm-FWNH2UNTgw5H7MTcHQFl |
| 77-60 | BAX (short) | https://benchling.com/s/seq-aINRskIXSYSJcJv8aURp?m=slm-zKz2j49bk5r1v2mVWmMc |
| 76-68 | p53 (short) (1-61) | https://benchling.com/s/seq-oboHu3wQGGrOxVZXR8mT?m=slm-CL2SXiwFRno7endZDLq8 |
| 76-69 | IFNG (24-156) | https://benchling.com/s/seq-Rxb7OuSxhwC1KxJykaPR?m=slm-qgjWG0Jvg76X7yQpLdCb |
| 76-153 | CRBN | https://benchling.com/s/seq-BAPh6kwfucKjYDmmPXFp?m=slm-i01HyK2nieJmwBXY5n6z |
| 76-100 | MAX | https://benchling.com/s/seq-GKeMoRGv4qd77TtfORFX?m=slm-i9MBG0ae66SnGxHPZgvF |
| 73-170 | PPP1R11 | https://benchling.com/s/seq-XHGhEcm7nvFEbuDNURBH?m=slm-SRI1vBi0GmF6ybMdXEbZ |
| 73-174 | PMP22 | https://benchling.com/s/seq-AFHDnBiEoYvLILOGoita?m=slm-Bi5fpdTxyr84R5wT0uow |
| 77-53 | BAX | https://benchling.com/s/seq-SxhwkmdaNisoVdvFR6IP?m=slm-pbu3gAKdt55C3yvTXplM |
| 73-186 | Keap1 | https://benchling.com/s/seq-c4pJiEFPzHNjGtroj130?m=slm-VCVLym1r9ip2BtvQwbWY |
| 73-163 | HSPB1 | https://benchling.com/s/seq-sOzF6JEREPkrjU54uVFv?m=slm-vhYiNPEDQpHkYFGytVSr |
| 73-177 | HMGB1 | https://benchling.com/s/seq-Hme2Y2BCw59yBZQevApH?m=slm-00Z5hzluKqBGFduBmlC0 |
| 73-169 | CD3d | https://benchling.com/s/seq-WcQwZtijlEoLB1KU2yBt?m=slm-XAGmL1pF0bAJlqi2L7qA |
| SL515 | TRIM63 (160-353) | https://benchling.com/s/seq-fwVYgi9unEvoEI8oXexV?m=slm-DOMbXkYqizg4IF4mLXVc |

| | | |
|---|---|---|
| SL572 | RHEB | https://benchling.com/s/seq-Ql4ELNdU367IodSPPmNh?m=slm-Y8YKNacV3lCzzYBW93GC |
| SL516 | RNF2 (113-226, DZn finger) | https://benchling.com/s/seq-8zSY9KpOqaM5rvajPKJc?m=slm-29976uY99HfcSVK1RvXa |
| SL517 | RNF8 (DZn finger) | https://benchling.com/s/seq-3TRNLC8OP20NAfMyNNtw?m=slm-9macfF4Sbe7vQOWEGvN5 |
| SL573 | BIRC7 | https://benchling.com/s/seq-EBsZSvur6XGc0lA10aTx?m=slm-0f1dmd30kLTaTpajNkqS |
| SL489 | XIAP | https://benchling.com/s/seq-VsMuzcUp9dV1yE3eLrkY?m=slm-Y1bXQMjCCaMyc8bYwLnP |
| --- | SIX1-DBD (1-167) | https://benchling.com/s/seq-HoMdI0SJUOMWO8TKFuZ4?m=slm-sHtZ9lKEIwvFXiy9NSNY |
| SL518 | WWP2 | https://benchling.com/s/seq-KAIt2gwSkaTesvzUk3ln?m=slm-RpbJdCFILRabGxxgisI4 |
| 78-24 | SIX1 | https://benchling.com/s/seq-4Wi3Lu9zyNxaood3z3o0?m=slm-atIv55Q5gWiSeCIrVrDa |
| 78-27 | ERG-IDR (1-112) | https://benchling.com/s/seq-8MvsYnusD8uNvsiQDY7H?m=slm-FWlJdsiXT5KL8DlaE4gr |
| --- | HAND2-DBD (99-151) | https://benchling.com/s/seq-yfKSi6c1a0ooB0ev2lDA?m=slm-tR8tuQR496RwK0ok5KHa |
| SL574 | HES1 | https://benchling.com/s/seq-WWTa0eCv2lkfSOjHBwni?m=slm-EhyInCRZSJ1bjNkQDG58 |
| --- | HOXA9 | https://benchling.com/s/seq-a8LNlAi11jNRTDGXLbid?m=slm-qvFJJFru2vO2VgNbjqFJ |
| --- | GATA2-DBD (295-373) | https://benchling.com/s/seq-9v79p9hoU2TvOG5Yd1e2?m=slm-1jUj5K3C3NCeBWFesrsj |
| --- | POU5F1-DBD (138-212) | https://benchling.com/s/seq-EOGXysYLLAcAAUJOLhxa?m=slm-WGx3osXoALfw5Egj8ZvU |
| --- | THAP1 | https://benchling.com/s/seq-iaT5rv1flt6WH5HZ1j4s?m=slm-VjdF9qNrVKEXmoBBbKTZ |
| --- | TBX20-DBD (109-288 | https://benchling.com/s/seq-iVdF6UslfqafSLnZSQxS?m=slm-Obs6BiW6cFKDg1u6bnI3 |
| --- | OLIG2-DBD (108-165) | https://benchling.com/s/seq-sfigXvXADvpi7qqOcr5z?m=slm-RkLeSk6JUjZuVWyrqcci |
| --- | ZBED2 | https://benchling.com/s/seq-NgOg3spjQIgy6PEKE10f?m=slm-DLQMy03cbB4cVW7w1yDk |

| | | |
|---|---|---|
| SL575 | GAPBA-DBD (168-251) | https://benchling.com/s/seq-yPzv7Phne6N7T0lE2Iq4?m=slm-vQJV9e8BjEqjqbp5ZjYh |
| --- | GATA4 | https://benchling.com/s/seq-MivMzu3zo4MNid8SjZev?m=slm-ziTE1iUP9GuVy9a5XNnM |

**Supplementary Table 12: Primers used in this work.**

| ID | Description | Sequence ($X_{12}$ indicates sequence specific to target) |
|---|---|---|
| Library cloning | | |
| MS-783 | Forward primer for Design A affibody | CATCATCATGCTAGCGTAGATAATAAATTTWMYAA<br>AGAAKKGNNYDMYGCADHYNNYSARATTNNYNNY<br>CTGCCGAACTTAAACSNSNNYCAGVDRNNYGCCT<br>TCVTHNNYAGCCTGNNYRAYGACCCGTCCCAAAG<br>CGCTAATTTGCTGGCGGAG |
| MS-1157 | Forward primer for Design B affibody | CATCATgctagcGTAGATAATAAATTTAATAAAGAAN<br>NYNNYNNYGCANNYNNYGAAATTNNYNNYCTGCC<br>GAACTTAAACNNYNNYCAGNNYNNYGCCTTCATC<br>NNYAGCCTGNNYGATGACCCGTCCCAAAGCGCTA<br>ATTTGCTGGCGG |
| MS-1472 | Forward primer for Design C affibody | GCTAGCGTAGATNNYNNYTTTNNYNNKNNYGTAN<br>NYNNKGCACGTNNKNNYATTNNYNNYCTGCCGAA<br>CTTAAACCGCGAGCAGGGCGG |
| MS-650 | Forward primer for Design D affibody | CATCATgctagcGTAGATAATAAATTTAATAAAGAAN<br>NKNNKNNKGCANNKNNKGAAATTNNKNNKCTGCC<br>GAACTTAAACNNKNNKCAGNNKNNKGCCTTCATC<br>NNKAGCCTGNNKGATGACCCGTCCCAAAGCGCTA<br>ATTTGCTGGCGG |
| MS-1158 | Forward primer for Design E affibody | CATCATgctagcGTAGATAATAAATTTAATAAAGAAV<br>BYVBYVBYGCAVBYVBYGAAATTVBYVBYCTGCCG<br>AACTTAAACVBYVBYCAGVBYVBYGCCTTCATCVB<br>YAGCCTGVBYGATGACCCGTCCCAAAGCGCTAAT<br>TTGCTGGCGG |
| MS-1159 | Forward primer for Design F affibody | CATCATgctagcGTAGATAATAAATTTAATAAAGAAK<br>MYKMYKMYGCAKMYKMYGAAATTKMYKMYCTGC<br>CGAACTTAAACKMYKMYCAGKMYKMYGCCTTCAT<br>CKMYAGCCTGKMYGATGACCCGTCCCAAAGCGC<br>TAATTTGCTGGCGG |
| MS-1067 | Forward primer for Design G affibody | CATCATGCTAGCGTAGATAATAAATTTWMYAAAGA<br>AKKGNNYDMYGCADHYNNYSARATTNNYNNYCTG<br>CCGAACTTAAACSNSNNYCAGVDRNNYGCCTTCV<br>THNNYAGCGCTNNYRAYGACCCGTCCCAAAGCGC<br>TAATTTGCTGGCGGAG |
| MS-1068 | Forward primer for Design H affibody | CATCATGCTAGCGTAGATAATAAATTTWMYAAAGA<br>AKKGNNYDMYGCADHYNNYSARATTNNYNNYCTG<br>CCGAACTTAAACSNSNNYCAGVDRNNYGCCTTCV<br>THNNYAGCCTGNNYRAYGACCCGTCCCAAAGCGC<br>TAATGGTCTGGCGGAGCTCAAAAAGTTG |
| MS-1069 | Forward primer for Design I affibody | CATCATGCTAGCGTAGATAATAAATTTWMYAAAGA<br>AKKGNNYDMYGCADHYNNYSARATTNNYNNYCTG<br>CCGAACGGTAACSNSNNYCAGVDRNNYGCCTTCV<br>THNNYAGCGCTNNYRAYGACCCGTCCCAAAGCGC<br>TAATTTGCTGGCGGAGGGGAAAAAG |
| MS-618 | Reverse primer for all affibody designs | CATCATgctagcGCCAGAGCCAGAGCCACCCTTGTA<br>GAC |
| MS-1658 | Forward primer for full random 30mer NNY Design J | catcatcatGCTAGCaacatgctccctcaatcggttgaatgtcgccc |
| MS-1659 | Reverse primer for full random 30mer NNY Design J | catcatcatGCTAGCgaaagatctccaTTARNNRNNRNNRN<br>NRNNRNNRNNRNNRNNRNNRNNRNNRNNRNNRN<br>NRNNRNNRNNRNNRNNRNNRNNRNNRNNRNNRN |

| | | |
|---|---|---|
| | | NRNNRNNRNNRNNgCTCGAGCCAGAGCCAGAGC CACC |
| qPCR | | |
| VC-525 | Forward primer for qPCR | ggttcagcaaggtgatgctt |
| VC-526 | Reverse primer for qPCR | accgcctcacctctgtttta |
| Luciferase Assay | | |
| --- | General Forward for insert for cloning scaffold into phage or Lux-N | tggctctggctctggctcgagcXXXXXXXXXXXX |
| --- | General Reverse for insert for cloning scaffold into phage | cgattgagggagcatgttgaaaatctccattaXXXXXXXXXXXX |
| --- | General Reverse for insert for cloning scaffold into Lux-N | gctgaggagtgccgttaattaagtttaXXXXXXXXXXXX |
| BR-305 | Forward for vector for cloning scaffold into phage | tggagattttcaacatgctccctcaatcg |
| MS-40 | Forward for vector for cloning scaffold into Lux-N | taaacttaattaacggcactcctcagcaaatataatgacc |
| KJ-14 | Reverse for vector for cloning scaffold into phage or Lux-N | CTCTGGCTCTGGCTCGAGC |
| --- | General Forward for insert for cloning target into +AP | ggatccctcgaaaggaggaaaaaaaATGXXXXXXXXXXXX |
| --- | General Forward for insert for cloning target into Lux-C | gtctgacataaatgaccgctATGXXXXXXXXXXXX |
| --- | General Reverse for insert for cloning target into +AP/Lux-C | ctttaccgcttccacccgacgtXXXXXXXXXXXX |
| KJ-19 | Forward for vector for cloning target into +AP or Lux-C | ACGTCGGGTGGAAGCGGT |
| MS-46 | Reverse for vector for cloning target into +AP | tttttttcctcctttcgagGGATCCtaggtag |
| BR-116 | Reverse for vector for cloning scaffold into Lux-C | ACAACTCAAGTCTGACATAAATGACCGCT |
| MS-659 | Forward for insert for subcloning binder variants into Lux-N | cacgtctacaagGGTGGCTCTG |
| JD-856 | Reverse for insert for subcloning binder variants into Lux-N | tcaatcggttgaatgtcgcccttt |
| MS-660 | Forward for vector for subcloning binder variants into Lux-N | aatcggttgaatgtcgcccttacttaattaacggcactcctcagc |
| MS-661 | Reverse for vector for subcloning binder variants into Lux-N | cacgtctacaagGGTGGCTCT |
| Next generation sequencing | | |
| MS-1156 | Reverse primer for Amplicon-EZ NGS for all barcodes (purple is adaptor, green is priming region) | gactggagttcagacgtgtgctcttccgatctgggcgacattcaaccgatt gaggg |
| MS-1144 | Forward primer for Amplicon-EZ NGS Barcode 1 (red is adaptor, blue is barcode, green is priming region) | acactctttccctacacgacgctcttccgatctAAGGTTgcgcgtaggg cacgtctacaag |
| MS-1145 | Forward primer for Amplicon-EZ NGS Barcode 2 | acactctttccctacacgacgctcttccgatctTAGGATgcgcgtaggg cacgtctacaag |
| MS-1146 | Forward primer for Amplicon-EZ NGS Barcode 3 | acactctttccctacacgacgctcttccgatctTGAGATgcgcgtaggg cacgtctacaag |
| MS-1147 | Forward primer for Amplicon-EZ NGS Barcode 4 | acactctttccctacacgacgctcttccgatctAGTGATgcgcgtaggg cacgtctacaag |

| | | |
|---|---|---|
| MS-1148 | Forward primer for Amplicon-EZ NGS Barcode 5 | acactctttccctacacgacgctcttccgatctAAGTGTgcgcgtagggcacgtctacaag |
| MS-1149 | Forward primer for Amplicon-EZ NGS Barcode 6 | acactctttccctacacgacgctcttccgatctAAGTTGgcgcgtagggcacgtctacaag |
| MS-1150 | Forward primer for Amplicon-EZ NGS Barcode 7 | acactctttccctacacgacgctcttccgatctAGATGTgcgcgtagggcacgtctacaag |
| MS-1151 | Forward primer for Amplicon-EZ NGS Barcode 8 | acactctttccctacacgacgctcttccgatctAGTAGTgcgcgtagggcacgtctacaag |
| MS-1152 | Forward primer for Amplicon-EZ NGS Barcode 9 | acactctttccctacacgacgctcttccgatctGAATTGgcgcgtagggcacgtctacaag |
| MS-1153 | Forward primer for Amplicon-EZ NGS Barcode 10 | acactctttccctacacgacgctcttccgatctGATATGgcgcgtagggcacgtctacaag |
| MS-1154 | Forward primer for Amplicon-EZ NGS Barcode 11 | acactctttccctacacgacgctcttccgatctGTAATGgcgcgtagggcacgtctacaag |
| MS-1155 | Forward primer for Amplicon-EZ NGS Barcode 12 | acactctttccctacacgacgctcttccgatctGTATAGgcgcgtagggcacgtctacaag |
| MS-1346 | Forward primer for Amplicon-EZ NGS Barcode 13 | acactctttccctacacgacgctcttccgatctAACCTTgcgcgtagggcacgtctacaag |
| MS-1347 | Forward primer for Amplicon-EZ NGS Barcode 14 | acactctttccctacacgacgctcttccgatctTACCATgcgcgtagggcacgtctacaag |
| MS-1348 | Forward primer for Amplicon-EZ NGS Barcode 15 | acactctttccctacacgacgctcttccgatctTCACATgcgcgtagggcacgtctacaag |
| MS-1349 | Forward primer for Amplicon-EZ NGS Barcode 16 | acactctttccctacacgacgctcttccgatctACTCATgcgcgtagggcacgtctacaag |
| MS-1350 | Forward primer for Amplicon-EZ NGS Barcode 17 | acactctttccctacacgacgctcttccgatctAACTCTgcgcgtagggcacgtctacaag |
| MS-1351 | Forward primer for Amplicon-EZ NGS Barcode 18 | acactctttccctacacgacgctcttccgatctAACTTCgcgcgtagggcacgtctacaag |
| MS-1352 | Forward primer for Amplicon-EZ NGS Barcode 19 | acactctttccctacacgacgctcttccgatctACATCTgcgcgtagggcacgtctacaag |
| MS-1353 | Forward primer for Amplicon-EZ NGS Barcode 20 | acactctttccctacacgacgctcttccgatctACTACTgcgcgtagggcacgtctacaag |
| MS-1354 | Forward primer for Amplicon-EZ NGS Barcode 21 | acactctttccctacacgacgctcttccgatctCAATTCgcgcgtagggcacgtctacaag |
| MS-1355 | Forward primer for Amplicon-EZ NGS Barcode 22 | acactctttccctacacgacgctcttccgatctCATATCgcgcgtagggcacgtctacaag |
| MS-1356 | Forward primer for Amplicon-EZ NGS Barcode 23 | acactctttccctacacgacgctcttccgatctCTAATCgcgcgtagggcacgtctacaag |
| MS-1357 | Forward primer for Amplicon-EZ NGS Barcode 24 | acactctttccctacacgacgctcttccgatctCTATACgcgcgtagggcacgtctacaag |

**Supplementary Note 1: Clustering, convergence and library design.**

Clustering of binders from high-fidelity selections likely provides evidence of the evolvability of variants for affinity maturation by identifying invariant positions (converged), possibly optimizable positions (partial convergence) and non-dependent positions (no convergence). When too few variants are identified for a target, the binder probably has a binding motif with a frequency near the sampling size, and its local fitness landscape has not been well explored in the selection from which it was derived (Supplementary Figs 7 and 13).

The motif frequency framework should also guide library randomization design. Binding motifs isolated from selections of moderate ($10^8$) sampling size are unlikely to converge at more than 5–6 randomized positions; randomizing many additional positions should therefore be limited to those that create new types of motifs (for example, two helices rather than one) or whose randomization has minimal impact on scaffold stability (for example, additional residues in nanobody CDR3 loops).

The machine-learning model can additionally learn features needed for binding that convergence analysis cannot see, including fixed scaffold residues that participate in the binding interface, and it can reveal broader structural determinants of binding consistent with the critical role of scaffold stability. Finally, although we focused on the affibody scaffold, we suspect that motifs broadly adapted by natural evolution for binding a wide range of proteins (for example, antibodies, WD40 motifs and C-type lectins) have been selected because they have, or evolved to have, both a higher average binder density and a selection strategy matched to the types of landscapes involved.
